# KRAS G12 mutant alleles differentially control glutamine metabolism via FOXO1

**DOI:** 10.1101/2024.10.19.618135

**Authors:** Suzan Ber, Ming Yang, Marco Sciacovelli, Shamith Samarajiwa, Khushali Patel, Efterpi Nikitopoulou, Annie Howitt, Simon J. Cook, Ashok R. Venkitaraman, Christian Frezza, Alessandro Esposito

## Abstract

Mutations in KRAS, particularly at codon 12, are frequent in adenocarcinomas of the colon, lungs and pancreas, driving carcinogenesis by altering cell signalling and reprogramming metabolism. However, the specific mechanisms by which different KRAS G12 alleles initiate distinctive patterns of metabolic reprogramming is unclear. Using isogenic panels of colorectal cell lines harbouring the G12A, G12C, G12D and G12V heterozygous mutations and employing transcriptomics, metabolomics, and extensive biochemical validation, we demonstrate distinctive features of each allele. We also demonstrate that cells harbouring the common G12D and G12V oncogenic mutations significantly alter glutamine metabolism and nitrogen recycling through FOXO1-mediated regulation compared to parental lines. Moreover, with a combination of small molecule inhibitors targeting glutamine and glutamate metabolism, we also identify a common vulnerability that eliminates mutant cells selectively. These results highlight a previously unreported mutant-specific effect of KRAS alleles on metabolism and signalling that could be potentially harnessed for cancer therapy.

**In brief:** Ber *et al.* reveal common and distinct features of oncogenic KRAS mutant isogenic cell lines. Mutant lines upregulate different nitrogen-recycling metabolic pathways and show sensitivity to combinatorial drugging of glutamine metabolism.

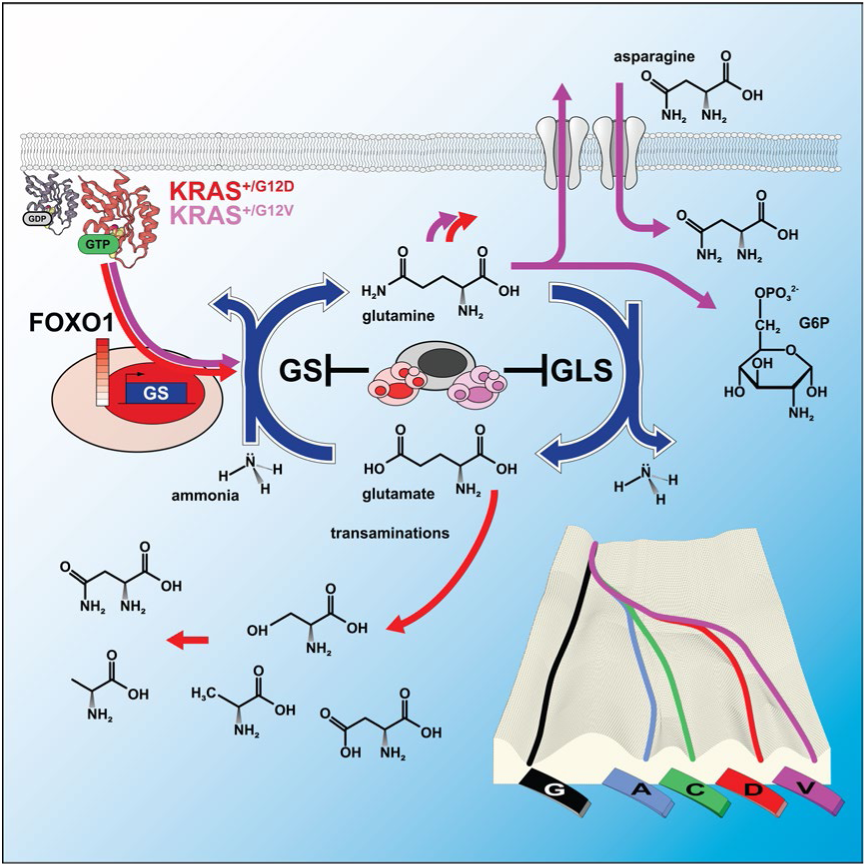

**Highlights:**

- Different KRAS mutations trigger distinct metabolic phenotypes in colorectal cell lines
- Glutamine metabolism and nitrogen recycling are upregulated in KRAS mutant cells
- FOXO1 is a key regulator of KRAS-induced rewiring of glutamine and nitrogen metabolism
- Inhibition of glutamine synthetase and glutaminase selectively kills mutant KRAS lines

## INTRODUCTION

The three RAS genes, HRAS, NRAS, and KRAS, code for membrane-bound small GTPases that play fundamental roles in development, adult tissue homeostasis, and disease. Mutations in RAS genes and deregulation of RAS-dependent signalling pathways drive several cancers associated with poor prognosis^1^. Missense mutations in RAS genes occur in 25-30% of all cancers, particularly at three hotspots, glycine-12 (G12), glycine-13 (G13), and glutamine-61 (Q61), with a tissue-dependent association. For instance, HRAS mutations occur more often in bladder cancer, NRAS in melanoma and KRAS in colon, lung and pancreatic adenocarcinomas^2,3^. The frequency of specific missense mutations within a hotspot also depends on the site of occurrence. For example, the missense KRAS mutations G12D and G12V are frequent in all KRAS-driven cancers, but G12R and G12C substitutions are very prevalent only in pancreatic (∼15-20%) and lung (∼60%) adenocarcinomas, respectively^4^.

The analysis of mutational signatures in patients suggests that the selection of mutation-dependent oncogenic signals might trigger distinct phenotypes in permissive tissues^1,5,6^. The mechanisms for this selection are largely undetermined, but different mutations at glycine-12 result in altered GTP hydrolysis mediated by KRAS, differential engagement of effector proteins and signalling^7–10^. Moreover, several studies have demonstrated that mutant KRAS also drives metabolic changes in cancers^11–14^ and that KRAS-driven metabolic reprogramming depends on the tissue of origin^15,16^, KRAS copy number^14^ and mutation^34^ (also reviewed in Kerk *et al.*^17^) These observations highlight the importance of understanding the mechanisms underpinning the pathogenicity of specific oncogenic KRAS alleles.

Here, we used hotspot mutations at the G12 codon of KRAS to investigate the consequences of different KRAS mutations on cellular phenotype in a panel of colorectal cell lines and to identify common vulnerabilities that could be targeted therapeutically. Transcriptomics and metabolomics revealed remarkable differences and commonalities between G12 mutant cells. More specifically, we identified significant differences in FOXO1 signalling, which regulates glutamine metabolism and provides proliferative advantages to KRAS mutant cells like G12D and G12V under limited nutrient conditions. We further show that the FOXO1-GLUL axis upregulated glutamine metabolism in these cells, leading to enhanced glutamine synthesis from extra-cellular glucose. At the same time, the upregulation of FOXO signalling – a pathway so far primarily associated with apoptosis in cancer^18–22^ enhances ammonia recycling via glutamine synthesis and transamination pathways, supporting the survival advantage of mutant cells. Notably, we identified that simultaneous targeting of glutamine synthesis and glutaminolysis could potently kill these mutants, suggesting a high dependency of KRAS mutant cells on nitrogen recycling and a possible new venue for therapeutic intervention.

## RESULTS

### KRAS missense mutations at glycine-12 perturb signalling and metabolic pathways

To investigate the effects of different KRAS G12 mutants, we first characterised the SW48 isogenic colorectal cancer cell line harbouring heterozygous mutations in KRAS at codon 12 (SW48^+/+^, SW48^+/G12A^, SW48^+/G12C^, SW48^+/G12D^, SW48^+/G12V^ hereafter also referred to as WT, G12A/C/D/V). We selected the G12D and G12V mutations because of their high frequency across all KRAS-driven cancers (including colorectal cancer) (Figure S1A), G12C for its high prevalence only in lung adenocarcinoma, and G12A because it is infrequently observed in colorectal adenocarcinoma yet biochemically indistinguishable from G12V^7^. Immunoprecipitation of GTP-bound KRAS confirmed that the mutant cell lines exhibit upregulated active KRAS (Figure S1B). However, these significant differences did not translate into clear upregulation of the well-characterised MAPK (mitogen-activated protein kinases) and PI3K (phosphoinositide 3-kinase) effector pathways as assessed by the phosphorylation of the ERK (extracellular signal-regulated kinase) and AKT kinases, respectively (Figure S1C). A modest upregulation of the ERK pathway was more apparent in serum-starved, low-nutrient conditions (1% FCS, 2 mM Glucose) for the G12V and G12D cell lines (Figure S1D), suggesting that these mutants are more capable of sustaining this signalling cascade than the others in limited nutrient and growth factor conditions.

We also characterised the panel by RNA sequencing to investigate the broader impact of the different KRAS mutant alleles. We found that about 2,000 genes are differentially regulated (false discovery rate less than 5% and log-2 fold change larger than 1) in at least one mutant cell line compared to parental SW48 cells (Figures 1A and S1E-F). Gene enrichment analysis (Table S1) highlighted differences in extracellular matrix receptor interactions (*e.g.*, laminins and integrins) and cell migration (semaphorin/plexin signalling) gene sets. Just around one hundred of these genes are similarly upregulated or downregulated in all mutant cell lines (Figures S1G (left) and S1H) and relate to transcriptional misregulation in cancer, including MAPK and mTOR signalling amongst other enriched KEGG pathways (File S1). DCLK1, MET and AKAP12 are some of the genes with significant changes in mutant lines that have been previously reported^23^. Most of the genes upregulated in G12D seem to be upregulated also in G12V, although to a lesser extent. As G12D and G12V are the most prevalent KRAS mutations in colorectal adenocarcinoma, we further analysed the differentially regulated genes in both mutant cell lines (Figure 1B-C and S1G-H, Table S1 and File S1). We identified around 300 genes enriched in key metabolic pathways (*e.g.*, *ALDH4A1*, *ALDH5A1*, *GAD1, GLUL* and *ABAT*, signalling (*e.g.*, *IL1R2, MET, SEMA6C*, *PPP3CA*, *EFNB3*, *ABLIM3*, *PLXNA2*, *SLIT1*, *MYL9*, *MET*, *NFATC4)*, and transcriptional regulation (*e.g.*, *MEF2C, CEBPB, MYCN, BCL6, FOXO1, DUSP6*). Notably, both PCA and gene enrichment analysis (Figure 1B,C) suggest an involvement of *FOXO1* in the differences observed in the G12D and G12V mutants.

**Figure 1.**
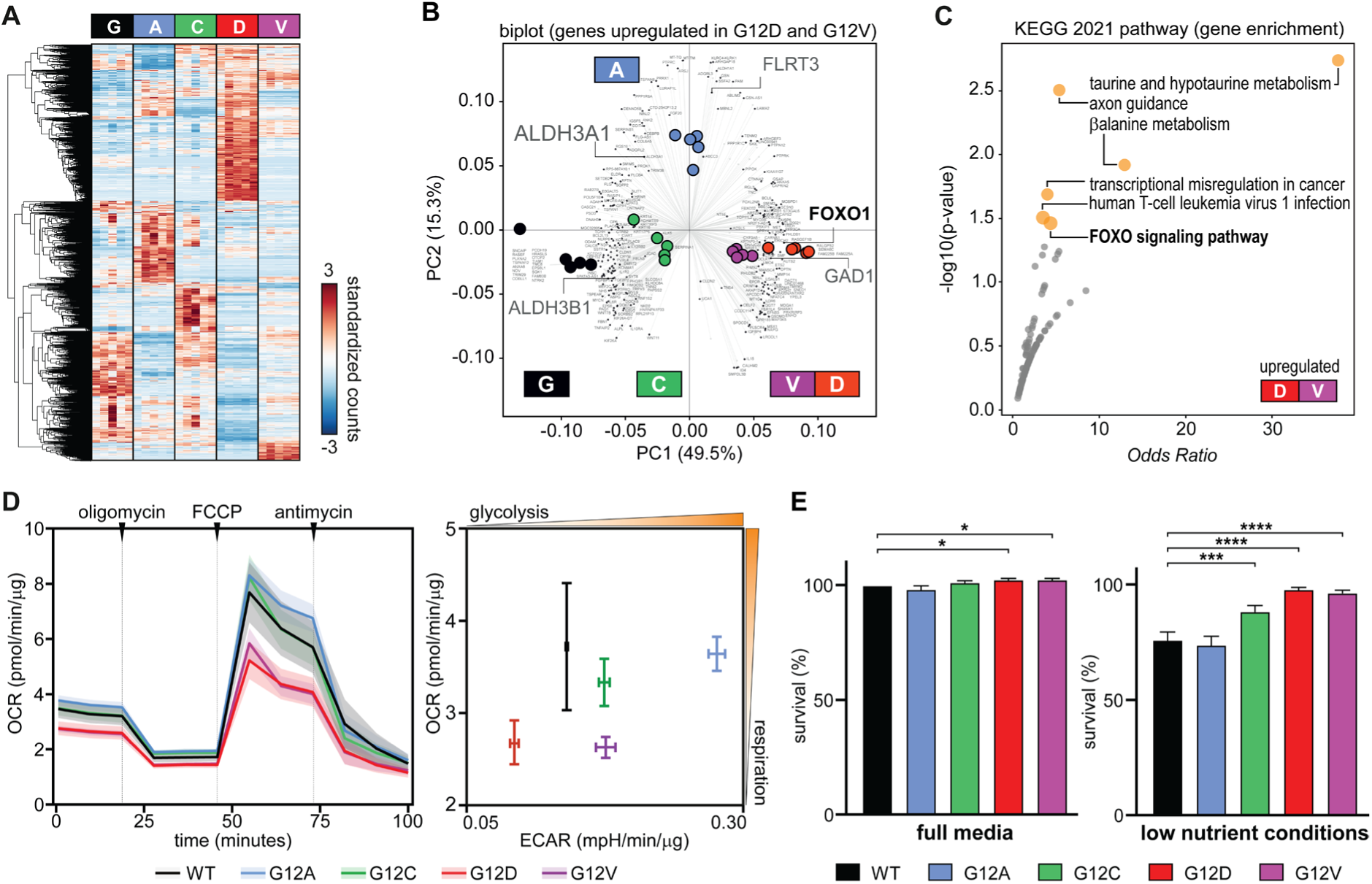
Isogenic SW48 cell lines reveal common and distinctive features of mutant KRAS alleles. (A) Hierarchical clustering showing genes that are differentially expressed in colorectal SW48 cancer cell lines harbouring heterozygous mutations (G12A, G12C, G12D and G12V) at glycine-12 of KRAS (n=5, FDR<5%, fold change > 2). Read counts were standardised for each gene, and each gene shown should be at least differentially regulated in one mutant line. Each cell line exhibits distinctive signatures. (B) Biplots related to PCA analysis with the gene loading in the background and the sample scores coloured in the foreground. The selected genes are upregulated in SW48 G12D and G12V (see also Figure S1E-H and Table S1). Some names of non-coding transcripts are omitted for clarity. (C) Example of gene enrichment analysis using EnrichR and KEGG 2021 pathways to analyse the genes indicated by the orange cluster in panel (Fig. S1G) suggesting that the G12D and G12V transcriptional signatures are related to metabolism and FOXO signalling. (D) Profiling of the oxygen consumption rate (OCR, left panel) and extracellular acidification rate (ECAR, right panel) obtained by the Seahorse extracellular flux analyser. OCR and ECAR were normalised to total protein content. Shaded areas are 95% confidence intervals, and error bars are standard deviations, evaluated over four independent repeats. (E) The growth rate of SW48 isogenic cells is similar in full media (left panel) but differs significantly in low nutrient, glucose and serum-reduced media conditions (right panel). Data represents an average of three independent experiments where each graph is normalised to WT in full media. p values are calculated using one-way ANOVA (*p ≤ 0.05, ***p ≤ 0.001, ****p ≤ 0.0001).

Since we observed significant changes in transcriptional and metabolic signatures, we verified whether these differences are reflected in mutant-specific alterations of cellular metabolism by characterising the mitochondrial function of SW48 cell lines (Figure 1D). The oxygen consumption rate (OCR) analysis revealed a lower basal and maximal respiration in G12D and G12V mutant cell lines. The extracellular acidification rate (ECAR) of G12D and G12V cells was lower or comparable to wild-type KRAS cells and the other mutants, suggesting that glycolysis does not compensate for the lower mitochondrial function of these cells. Differences in cellular respiration did not result in notable differences in cell viability as assessed by sulphorodamine B (SRB) colourimetric assay (Figure 1E, left panel) under standard culture conditions (RPMI with 11mM Glucose and 10% FCS). Given that growth media used for cell culture are formulated with supraphysiological concentrations of nutrients, we repeated the SRB assay with lower concentrations of serum (1%) and glucose (2 mM). These values are more representative of tissue interstitial levels^38^, and tumour microenvironment^24^. In these low-nutrient conditions, wild-type SW48 cells and the G12A line exhibited a substantial reduction in growth. In contrast, G12D and G12V cells exhibited high resilience to the change in nutrient conditions (Figure 1E, right panel).

### G12D and G12V KRAS mutants boost glutamine synthesis from glucose via FOXO1

To shed light on this unexpected metabolic behaviour of the G12D and G12V mutants, we traced glucose utilisation with ^13^C-glucose isotope and liquid chromatography-mass spectroscopy (LC-MS). Intracellular metabolomics showed that TCA (tricarboxylic acid) cycle metabolites like α-ketoglutarate and succinate are significantly lower in G12D and G12V mutant cells (Figure 2A), suggesting a lower TCA cycle activity. Surprisingly, G12D, G12V, and G12C to a lesser extent, exhibited a substantial flow of ^13^C-glucose carbon towards glutamine via glutamate, as indicated by the presence of *m*+2 isotopologues. The analysis of extracellular metabolites indicated that glutamine *m*+2 is also released in the media (Figure 2B). These results indicate that some KRAS mutants can generate glutamine from glucose.

**Figure 2.**
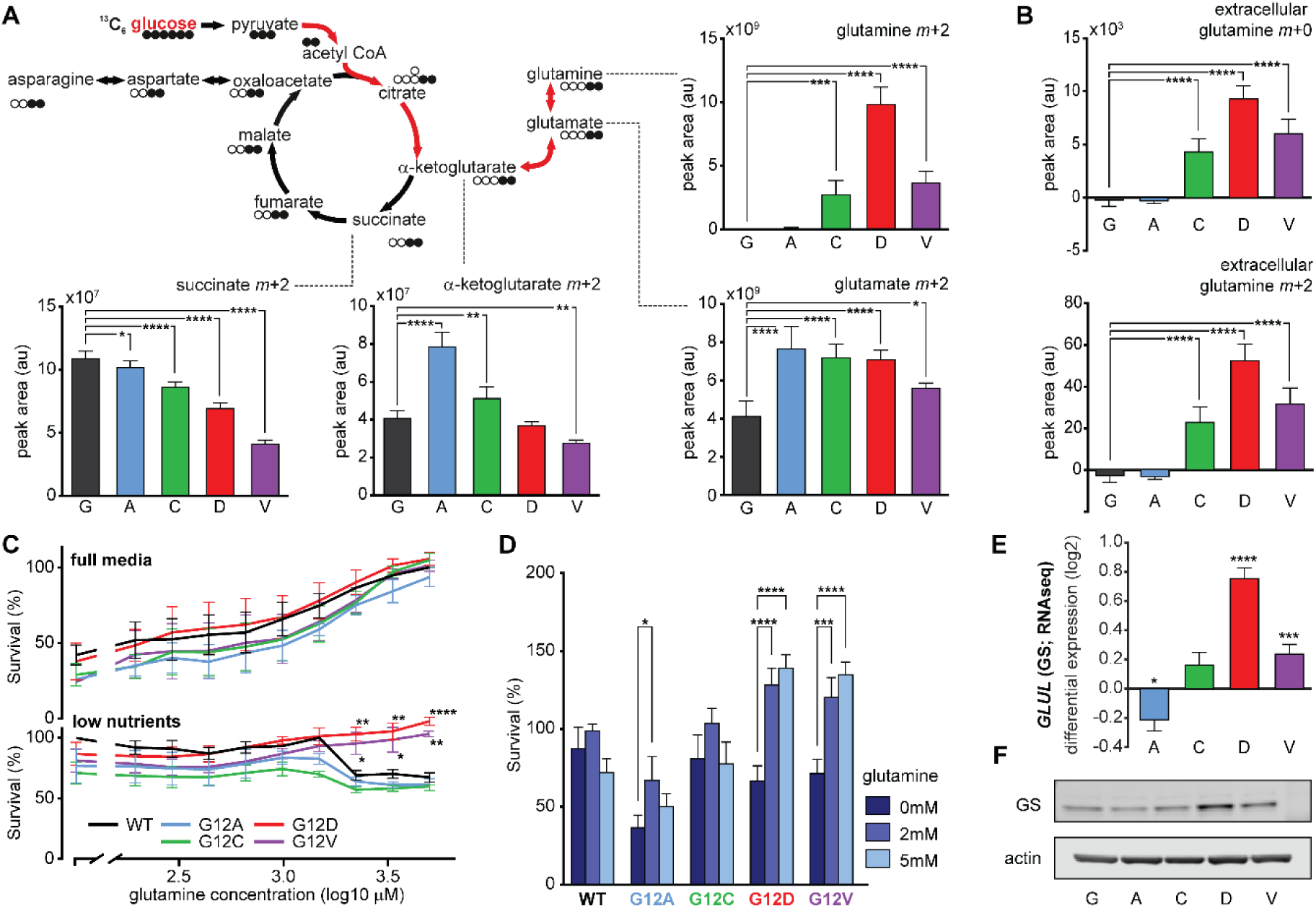
Oncogenic KRAS-allele dependent rewiring of cellular metabolism highlights differences in glutamine metabolism. (A) Metabolic flux analysis using ^13^C-labelled glucose. The diagram shows metabolite labelling patterns in the TCA cycle and glutamine pathway, depicting the incorporation of heavy carbon isotopes with full black circles. The red arrows highlight how the TCA cycle incorporates carbon from glucose into glutamine. The graphs show the intracellular abundance of glutamate, glutamine, succinate, and α-ketoglutaric acid (αKG) incorporating carbon derived from labelled glucose. Data represents the mean of 5 technical repeats and standard deviations. (B) Extracellular abundance of unlabelled (*m*+0) and labelled glutamine (*m*+2) determined by consumption-release measurements. Data represents the mean of 5 technical repeats and standard deviations. (C) Survival of isogenic SW48 cell lines at varying concentrations of glutamine in full media (up) and low nutrient media conditions (down) measured by SRB assay. Survival curves are the average of 3 independent experiments. The data is normalised to the highest viability of SW48 WT in each media condition. (D) Survival of SW48 isogenic cells in 0, 2 and 5 mM glutamine in low nutrient media conditions. The bar graph is representative of 4 independent experiments with standard errors. Values are normalised to the viability of WT at 2 mM glutamine in each experiment. (E) Differential expression of the *GLUL* gene coding for glutamine synthetase (GS) as measured by RNAseq and shown with standard errors. (F) Representative immunoblot showing glutamine synthetase (GS) levels in SW48 cell panel. Unless stated otherwise, data is shown as averages and standard deviations; all p-values (*p ≤ 0.05, **p ≤ 0.01, ***p ≤ 0.001, ****p ≤ 0.0001) were calculated with two-way ANOVA with corrections for multiple comparisons.

The synthesis of glutamine by some of the mutant cell lines prompted us to test the dependency of mutants on glutamine. In full growth media (10% FCS and 11 mM glucose), all SW48 cells of the isogenic panel increased viability with higher glutamine concentrations (Figure 2C, top panel). However, in low nutrient conditions (1% FCS and 2 mM glucose), the wild-type, G12A and G12C cell lines exhibited a significant decline in viability at a glutamine concentration above 1 mM (Figure 2C, bottom panel). On the contrary, the G12D and G12V mutant cells showed a further increase in viability at high concentrations of glutamine, thus resulting in a fitness advantage compared to the wild-type cells and the other mutants in nutrient conditions (*i.e.*, low glucose and high glutamine) reminiscent of the tumour microenvironment^24,25^ (Figure 2C,D). qPCR and western blotting (Figure 2E,F) reveal that glutamine synthetase (GS or *GLUL*) is expressed at a higher level in SW48^+/G12D^ and - to a lower extent - also in the G12V and G12C mutant cell lines. This observation is consistent with the increased synthesis of glutamine measured with metabolomics (Figure 2A,B).

Interestingly, *GLUL* is known to be under the transcriptional control of FOXO1^26^ and our trascriptomics analysis highlighted FOXO signalling as altered in the G12D and G12V mutant SW48 lines particularly (Figure 1B,C; upregulation of *BCL6*, *FOXO1* and *FOXO32*). We validated the higher expression of FOXO1 and its nuclear localisation by western blotting, qPCR, immunofluorescence and cell fractionation (Figures 3A-D and S3A). An increase of cytoplasmic FOXO1 was detectable in all mutant cells, although nuclear FOXO1 was upregulated more strongly in the G12D followed by G12V mutant lines.

**Figure 3.**
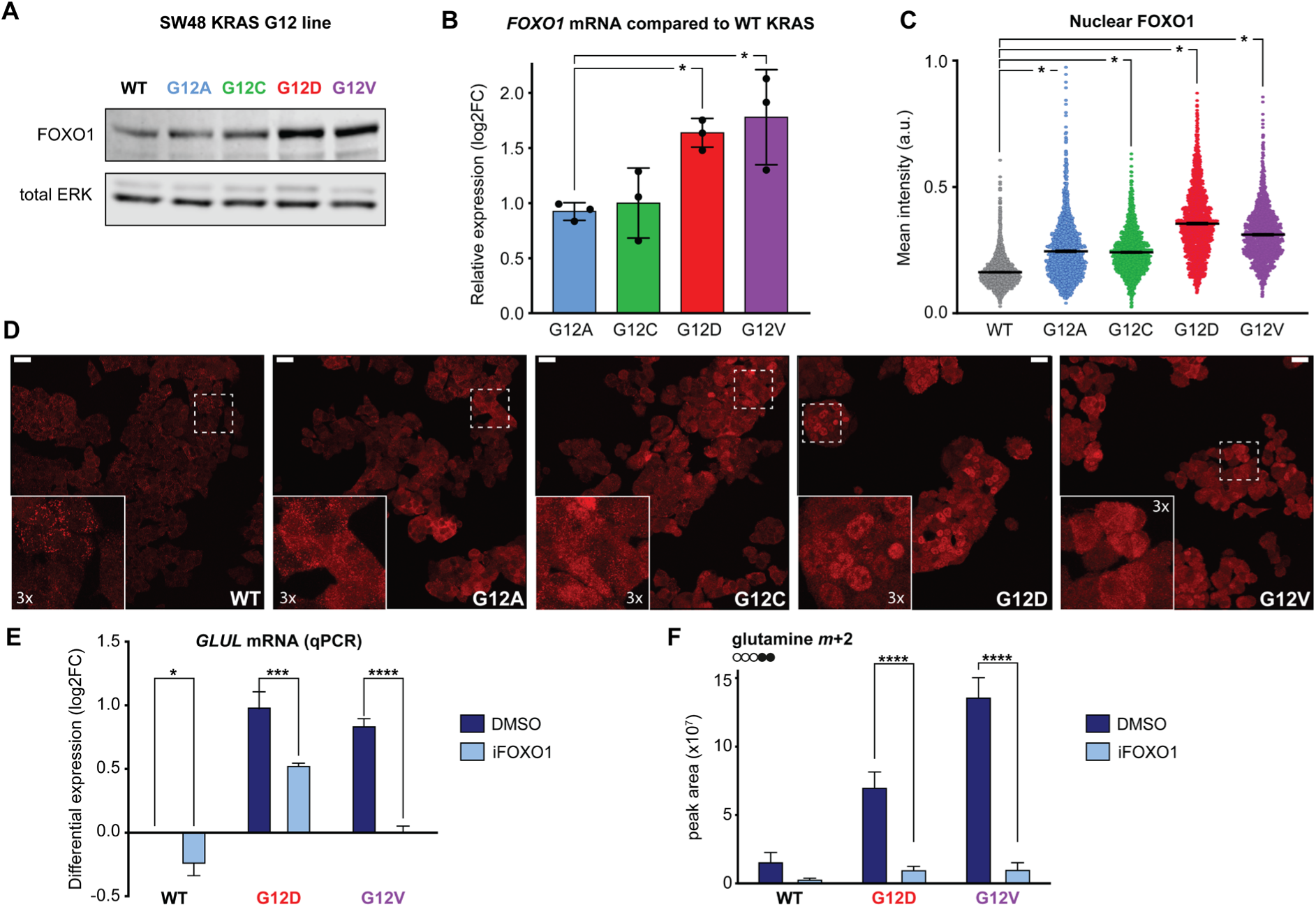
FOXO1 regulates glutamine synthesis upregulation in KRAS mutant cells. (A) Representative immunoblot (n=3) showing increased expression of FOXO1 in G12D and G12V mutant SW48 cells in full media. (B) mRNA levels of *FOXO1* detected by qPCR (n=3) compared by one-way ANOVA. (C-D) Immunostaining of FOXO1 in SW48 isogenic panel and quantification of mean nuclear intensity and standard errors (cells per sample ≥ 500, n=3) compared by one-way ANOVA. Scale bar: 25 µm; inserts: 3x magnification (E) qPCR analysis of *GLUL* mRNA levels upon FOXO1 inhibition. Values were normalised to SW48 WT DMSO of each experiment and shown with standard errors. (F) Changes in abundance of glutamine (*m*+2) upon inhibition of FOXO1 in a ^13^C-glucose labelling experiment. Data represents the mean of 5 technical repeats and standard deviations. Unless stated otherwise, data is shown as averages and standard deviations; all p-values (*p ≤ 0.05, **p ≤ 0.01, ***p ≤ 0.001, ****p ≤ 0.0001) were calculated with two-way ANOVA with corrections for multiple comparisons.

To test the hypothesis that the metabolic phenotype we observed depends on FOXO1, we tested gene expression and metabolic flux in these two mutant lines relative to the parental control when treated with the small molecule inhibitor AS1842856 (FOXOi). qPCR analysis (Figure 3E) confirmed that FOXO1 inhibition downregulates *GLUL* expression. Furthermore, transcriptomics (Figure S3C, Table S3 and File S1) permitted us to verify that FOXO1 inhibition downregulates several FOXO1 transcriptional targets and FOXO signalling more generally in addition to *GLUL*.

We also repeated ^13^C-glucose flux analysis in the presence of FOXO1 inhibition, demonstrating that the upregulation of glutamine synthesis from glucose (Figure 3F) in the G12D and G12V mutant lines depends on FOXO1. Taken together, this data indicates that FOXO1 is a critical factor in KRAS-dependent rewiring of glutamine metabolism.

### FOXO1 drives differential nitrogen recycling in G12D and G12V KRAS mutants

The synthesis of glutamine from glutamate is a major mechanism for ammonia detoxification, and FOXO1 has also been linked to nitrogen metabolism and ammonia assimilation^27^. Indeed, in addition to glutamine production, in our metabolomics experiment, FOXO1 inhibition significantly affected pathways related to ammonia recycling and nitrogen metabolism, such as the urea cycle, glutamate metabolism, arginine and proline metabolism, glycine and serine metabolism (Figure S3B). These results led us to investigate whether ammonia is recycled differently in the mutant lines (G12D and G12V) that exhibit high FOXO1 expression and increased glutamine synthesis.

To monitor ammonia incorporation, we cultured cells with labelled ammonia (^15^NH_3_ at 3 mM), confirming a significant *de novo* production of glutamine (*N*+1 and *N*+2 isotopologues) incorporating nitrogen from ammonia in G12V and G12D mutant cells (Figure 4A, first two graphs). Again, both mutant cells export more glutamine to the extracellular space compared to the parental line (Figure 4A, middle graph); however, G12D cells also incorporated the nitrogen derived from ammonia into several amino acids like aspartate, asparagine, serine, glycine and alanine (Figure 4B) via transamination reactions. The G12V cells preferentially produce and export glutamine *N*+1 and *N*+2 (Figure 4A) without upregulating transamination reactions. These cells also appear to accumulate aspartate (*N*+1) due to a lack of asparagine synthesis (Figure 4B). In addition, G12V showed enhanced ammonia incorporation into hexosamine biosynthesis (Figure S4A; see also File S2). We also noted that KRAS mutant cells substantially upregulated the synthesis of ophthalmic acid compared to the parental line. G12V cells were particularly efficient in incorporating labelled nitrogen from free ammonia into ophthalamate (Figure S4B).

**Figure 4.**
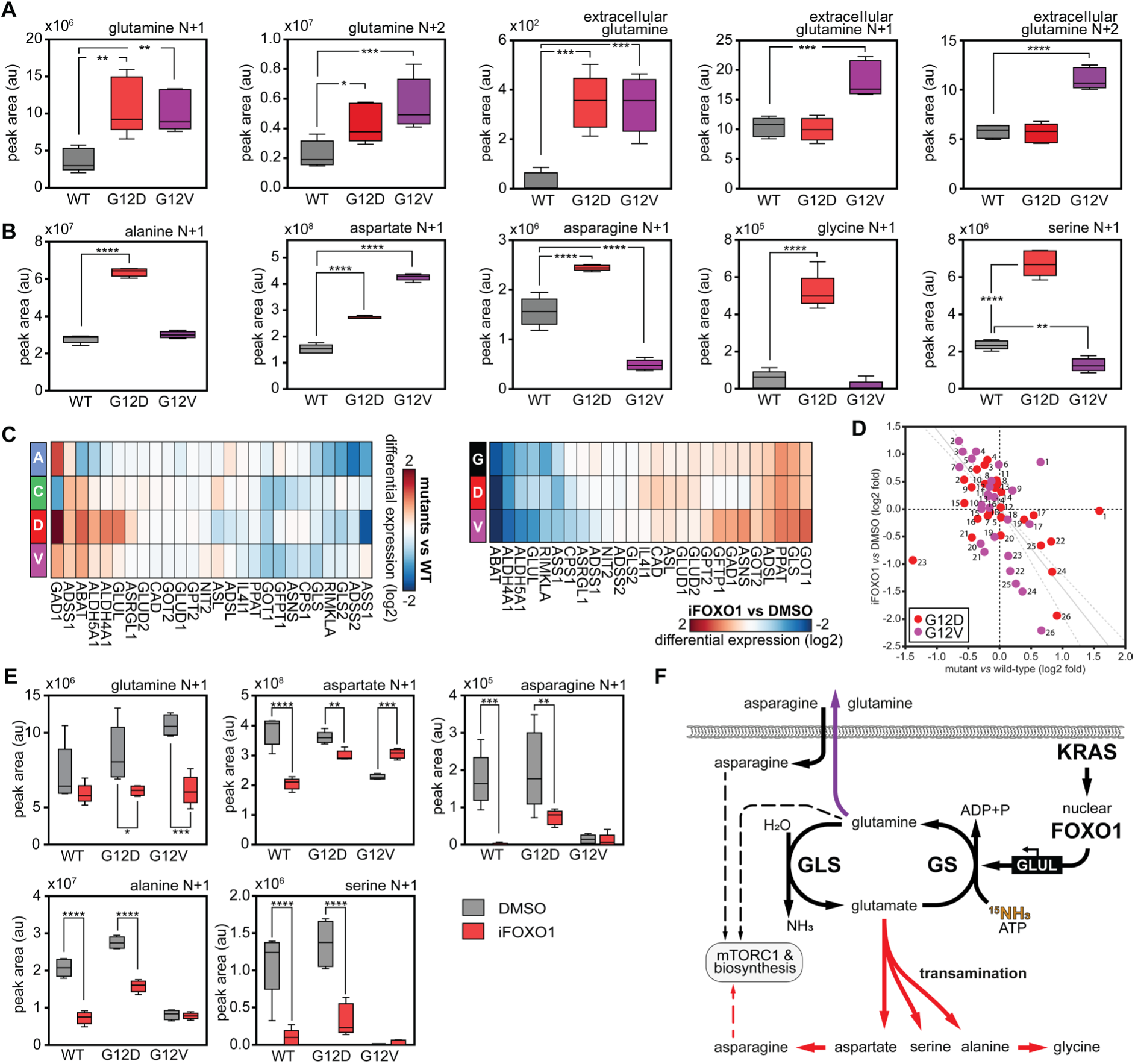
FOXO1 regulates differential nitrogen recycling in KRAS mutant cells. (A) Metabolic flux analysis using ^15^N-ammonia labelling shows elevated free ammonia incorporation into glutamine in G12D and G12V mutant cells. Synthesised ^15^N-labelled glutamine is released at higher rates in extracellular space in G12V mutant. (B) The abundance of ^15^N isotopologues in an ammonia labelling experiment showed increased ammonia incorporation into amino acids synthesis via transamination reactions, predominantly in G12D mutants. (C) Hierarchical clustering of gene differential expression (log-2 fold changes) in low-nutrient conditions showing the aspartate, alanine and glutamate metabolism pathway (KEGG pathway *hsa00250*). Left: SW48 mutant cells compared to wild-type cells; right: FOXO1 inhibition compared to DMSO control. We note that not every pathway gene is not detected in each experiment. (D) Correlation plot of the genes detected in both the experiments shown in (C); correlation coefficient of -0.4 and p-value∼0.001. Legend: (1) GAD1, (2) GOT1, (3) GLS, (4) PPAT, (5) ASNS, (6) ADSL, (7) GFPT1, (8) GOT2, (9) ASL, (10) IL4I1, (11) CAD, (12) GPT2, (13) GLUD1, (14) GLUD2, (15) GLS2, (16) ADSS2, (17) ADSS1, (18) NIT2, (19) ASRGL1, (20) CPS1, (21) RIMKLA, (22) GLUL, (23) ASS1, (24) ALDH4A1, (25) ALDH5A1, (26) ABAT. The solid and dashed grey lines represent the linear regression and the 95% confidence interval, respectively, with GAD1 and ASS1 outliers removed (correlation of -0.7 p-value<10^-7^). (E) Metabolic flux analysis using ^15^N-ammonia labelling with inhibition of FOXO1 reveals that the incorporation of ammonia-derived nitrogen in glutamine synthesis as well as in various transaminations depends on FOXO1. p-values were evaluated with two-way ANOVA. (F) Diagram showing the fate of ^15^N-labelled ammonia in KRAS mutant cells expressing high nuclear FOXO1. Unless otherwise stated, all metabolomics and transcriptomics data are shown as averages and standard deviations evaluated over 5 technical repeats. Adjusted p value was calculated using one-way ANOVA, (*p ≤ 0.05, **p ≤ 0.01, \*\*\**p* ≤ 0.001, \*\*\*\**p* ≤ 0.0001).

We complemented these experiments by labelling the amide and alpha nitrogens of glutamine. Efficient incorporation of nitrogen into aspartate, glycine, serine, and alanine in the G12D mutant and ophthalmic acid in both G12D and G12V mutants was seen predominantly from the alpha nitrogen of glutamine, suggesting an involvement of glutamate in these reactions (Figure S4C). In low-nutrient conditions, RNAseq analysis also shows that genes within the aspartate, alanine and glutamate metabolism pathway (KEGG2021) are differentially regulated in the mutant lines (Figure 4C, left panel). We repeated transcriptomics in the most different mutants (G12D and G12V) in the presence of FOXO1 inhibition or a DMSO control (Figure 4C, right panel). Correlation analysis between log2-fold changes shown in Figure 4C suggests that most changes in this metabolic pathway might be attributed to the expression of the oncogenic KRAS alleles and alterations mediated by FOXO1 (Figure 4D).

Metabolomics and transcriptomics thus suggest that KRAS G12D and G12V alleles upregulate nitrogen recycling pathways via FOXO signalling. To test this hypothesis, we repeated the metabolomic analysis of wild-type, G12D, and G12V mutant cells using labelled ammonia and the FOXO1 inhibitor. FOXO1 inhibition resulted in a substantial decrease in glutamine synthesis and transamination reactions (Figure 4E). Several independent lines of evidence thus suggest that oncogenic KRAS G12D and G12V lead to a higher expression of nuclear FOXO1, which in turn results in increased glutamine synthesis, ammonia detoxification via glutamine synthetase upregulation, and amino acid synthesis via transamination reactions in G12D. These mechanisms could maintain the proliferation potential of these mutant cells even in nutrient-limiting conditions (Figure 4F) and might thus represent valuable therapeutic targets.

### Combinatorial drugging of glutamine metabolic pathway as a therapeutic strategy

Therefore, we tested if inhibition of FOXO1 in low nutrient conditions would kill mutant cell lines selectively. However, FOXO1 inhibition reduced the viability of all tested cell lines, particularly cells with lower nuclear FOXO1, including KRAS wild-type cells (Figure S5A). Therefore, we investigated if targeting glutamine metabolism downstream of FOXO1 could offer selectivity for mutant cells (Figure 5C). First, we targeted the glutamine synthesis pathway with the well-characterized GLUL inhibitor methionine sulfoximine (MSO)^28–30^. While we observed a higher sensitivity of KRAS mutant cells relative to wild-type SW48 cells, the effects were only significant at high MSO concentrations (Figure 5A, left panel).

**Figure 5.**
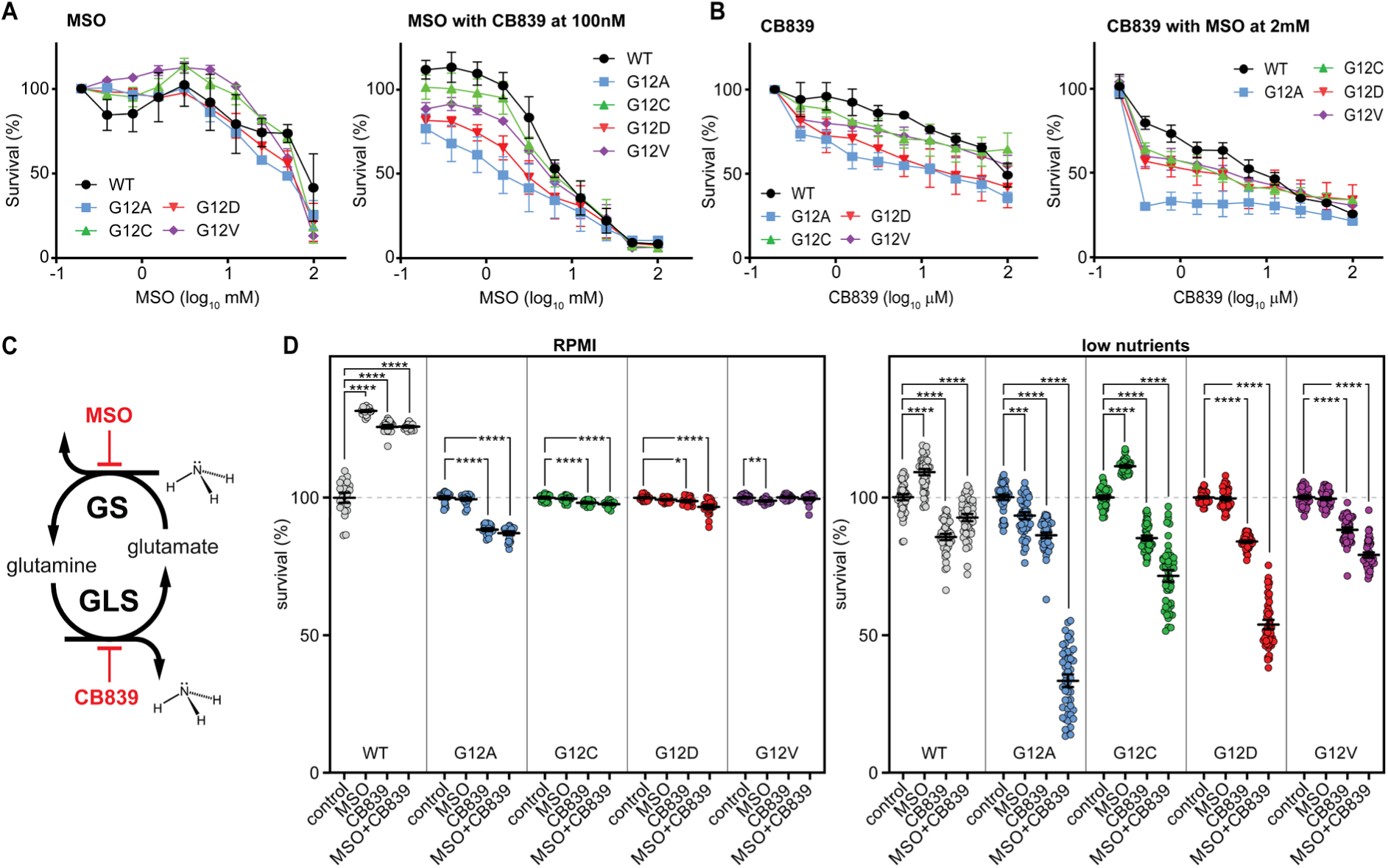
Inhibition of the glutamate-glutamine cycle sensitises KRAS mutant cells. (A) Viability curves showing titration of GS inhibitor, MSO, alone (left panel) or in combination with sub-lethal doses of GLS inhibitor, CB839, (right panel) is SW48 cells. (B) Viability curves showing titration of the GLS inhibitor, CB839, alone (left panel) or in combination with sub-lethal doses of the GS inhibitor, MSO (right panel), in SW48 cells. (C) Schematic representation of the glutamate-glutamine cycle and the drugs that inhibit the corresponding enzymes in this pathway. (D) Survival graphs of SW48 cells treated with single doses of GLS inhibitor (CB839) 100 nM, GS inhibitor (MSO) 2 mM alone or in combination in full media (left panel) and low nutrient media (right panel) conditions. Data represents the mean of 4 independent experiments. Adjusted p values were calculated using one-way ANOVA, (*p ≤ 0.05, **p ≤ 0.01, ***p ≤ 0.001, ****p ≤ 0.0001). Unless stated otherwise, the graphs show means and standard deviations of at least 3 independent assays.

All mutant cell lines exhibit high concentrations of glutamate, which could shift the balance of glutamine anabolism. It is thus conceivable that mutant cell lines might be more sensitive to the inhibition of glutaminase (GLS), the enzyme that converts glutamine to glutamate. We thus tested the effects of CB839, a GLS inhibitor that is already in early clinical trials in kidney and breast cancer^31,32^, on the SW48 isogenic panel. GLS inhibition does show some selectivity in the viability of mutant cell lines at the tested concentrations, particularly in G12A and G12D cells (Figure 5B, left panel). Notably, GLS is downregulated in all mutant lines (Figure 4C), particularly in the G12A mutant, which also exhibits the highest glutamate concentration within the SW48 panel.

Therefore, we tested whether MSO and CB839 - together - could further sensitise the mutant cell lines by targeting features common to all mutant cells (low *GLS* and high glutamate) and more specific to G12D and G12V mutant cells (high glutamine and high *GLUL*). When we titrated MSO and CB839 in the presence of sublethal concentrations of CB839 (100 nM) and MSO (2 mM), respectively, we observed a substantial reduction of the viability of KRAS mutant cells, including the most resilient mutants G12D and G12V (right panels of Figure 5A,B). Knocking down the two genes by siRNA also confirmed the sensitivity of G12D and G12V mutant cells to the inhibition of GLS and GS together (Figure S5G). The titration of both drugs reveals a strong synergistic effect of the GS and GLS inhibitors at all concentrations, particularly in G12D and G12V mutant cell lines, as tested by Bliss analysis^33^ (Figure S5B).

Finally, we investigated how robust our observations are in relation to nutrient conditions and cell types. We repeated the drug-sensitivity experiments at a fixed dose of CB839 (100 nM) and MSO (2 mM) in low nutrient media (1% FCS and 2 mM glucose) and full RPMI growth media conditions (Figure 5D). The viability data confirms that cells harbouring the mutant KRAS alleles are substantially more sensitive to the two drugs than the wild-type SW48 cell line. This sensitivity, however, disappeared once glucose and serum concentrations were restored to higher levels, suggesting that additional specificity could be achieved thanks to the tumour microenvironment that is usually deprived of glucose.

To mitigate the risk that the observed sensitivities entirely depend on the SW48 cell line background, we repeated crucial experiments using a second isogenic panel derived from the colorectal cancer cell line LIM1215 (Figure S5C-F). The LIM1215 panel harboured the same mutant alleles, also exhibited upregulation of FOXO1, and revealed the same sensitivity of mutant LIM1215 to combinatorial drugging of glutamine synthetase and glutaminase, as observed in SW48 isogenic panel.

Taken together, our results suggest that KRAS mutant cells, particularly those that exhibit upregulation of FOXO signalling, rely on glutamine synthesis and catabolism to survive in limited glucose conditions. Small-molecule inhibitors of clinical interest may be able to target this mechanism to achieve selective killing of oncogenic KRAS cells.

## DISCUSSION

Despite the vast literature on KRAS, the complex mechanisms underpinning the pathogenesis of specific KRAS mutant alleles have yet to be unravelled^3^. Therefore, we have carried out a systematic quantitative characterisation of four mutant isogenic cell lines comparing the two most common (G12D and G12V) and two rare (G12A and G12C) oncogenic mutations found in colorectal cancer with the parental SW48 line wild-type for KRAS.

While also reproducing results reported by other studies^34,35^, we observed that the differential gene expression and metabolic phenotypes we reported in Figure 1 do not translate into substantial differences in the viability of cell lines in full media conditions. Providing supraphysiological concentration of nutrients, full growth media is now known to cause significant differences between *in vitro* and *in vivo* studies^36–38^. Therefore, we utilised lower serum (1%) and glucose concentrations (2 mM) compared to full media (10% and 11 mM) to provide availability of nutrients more similar to interstitial fluids and tumour microenvironment^24,39^. SW48^+/G12D^ and SW48^+/G12V^ cell lines exhibit increased resilience in low nutrient conditions; moreover, these mutant lines demonstrate increased viability in high glutamine and low glucose concentrations (Figure 2), conditions that have been reported, for example, in a murine model of KRAS-driven pancreatic adenocarcinoma^24^.

Transcriptomics, metabolomics and validation data demonstrate a key role of FOXO1 signalling in KRAS-induced metabolic reprogramming (Figures 1, 3 and 4). To our knowledge, the implication of FOXO1 in KRAS-driven metabolic reprogramming is so far unreported. *FOXO* genes have been mostly regarded as tumour suppressors^20,22^ although recent work is changing this paradigm^19,40–42^. Stress pathways alter post-translational modifications of FOXO proteins, such as phosphorylation and acetylation^43–45^, regulating FOXO proteins’ diverse nuclear and cytoplasmic activities, thus establishing a balance between their function as tumour suppressors and as promoters of tumour growth^18,46,47^.

Our investigation shows that oncogenic KRAS induces FOXO1 expression and nuclear localisation, resulting in higher expression of its transcriptional target glutamine synthetase (GS), also known as glutamate-ammonia ligase or *GLUL*. The SW48^+/G12D^ and SW48^+/G12V^ cell lines share several transcriptional alterations, including the upregulation of FOXO1. In contrast, cells harbouring the G12C mutations exhibit a milder phenotype, and G12A cells are more similar to parental SW48 cells. This observation is intriguing because G12D and G12V are the most common mutations occurring in colorectal cancers. Metabolic profiling of the panel through oxygen consumption and media acidification confirmed the similarity between these mutants but – contrary to expectations – revealed lower mitochondrial respiration and glycolysis compared to wild-type and other mutants (Figure 1D). Metabolic flux analysis performed with ^13^C-glucose shows that all mutant cell lines avidly consume pyruvate to feed the TCA cycle, as shown by high levels of labelled citrate, which is eventually utilised to maintain high glutamate to support biogenesis in mutant cells (Figure 2A).

G12A cell line exhibits higher levels of α-ketoglutarate compared to all other cell lines and a higher flux through the TCA cycle as inferred from several isotopologues such as α-ketoglutarate *m*+1, *m*+2, *m*+3 and *m*+4 and glutamate *m*+4 and *m*+5 (Figure S2F) which are generated after several TCA cycles. Contrary to this, the KRAS- and FOXO1-dependent upregulation of glutamine synthetase seems to deplete α-ketoglutarate from the TCA cycle in G12D and G12V cells (and G12C to a lower extent). The high production of glutamine in the G12D, G12V and G12C mutants also results in high aspartate levels that, together with glutamate and glutamine, feed into biosynthesis, resulting in higher production of nucleotides (Figure S2B,D). The higher concentration of glutamine, uptake of asparagine and nucleotide synthesis, particularly in G12D and G12V, likely support mTOR activation and, in turn, increase proliferative signals in these cells. Indeed, we detect an apparent increase in phosphorylation of 4EBP1 (EIF4EBP1), a major target of the mTOR pathway, in SW48^+/G12D^ and SW48^+/G12V^ (Figure S1C,D).

While G12D and G12V are very similar, they also exhibit significant differences in nitrogen recycling, some of which are regulated by FOXO1 (Figure 4 and S4). For example, the high production of glutamine and upregulation of transamination reactions in SW48 G12D cells result in higher synthesis of amino acids such as alanine, serine, and glycine; instead, G12V cell lines exhibit a much higher production of aspartate as well as synthesis and export of glutamine. Nitrogen flux analysis using ^15^N-labelled ammonia and glutamine (Figures 4 and S4) revealed a striking increase in ophthalmic acid production, an analogue of glutathione, in mutant cells, G12V in particular. Instead of upregulating transaminations reactions, the SW48^+/G12V^ line further fixed nitrogen through glycogenesis and hexosamine pathways (*e.g.*, in UDP-glucose, GDP-glucose, UDP-N-acetylglucosamine and sialic acid), a nitrogen recycling pathway linked to colorectal cancer metastasis^48,49^.

We also observe that G12V and G12D cell lines exhibit higher intracellular ammonia than the parental line (Figure S5H). Ammonia has long been considered a metabolic waste product but several publications have now shown that metabolic recycling of ammonia supports the proliferation of cancer cells ^50–53^. For instance, Spinelli and colleagues have shown that free ammonia released through reductive amination by glutamate dehydrogenase (GLUD, also known as GDH) is utilised for amino acid synthesis by breast cancer cells, leading to tumour growth^51^. Our observations suggest that the oncogenic mutants commonly found in colorectal cancer exhibit a similar phenotype, *i.e.*, they can use excess ammonia and adapt nitrogen metabolism to fuel biosynthetic pathways. This phenotype is directly linked to changes in the expression or localisation of FOXO1 downstream of KRAS signalling, supporting the well-established role of FOXO1 in regulating glutamine, nitrogen metabolism and ammonia detoxification^27,54^. Moreover, these metabolic changes observed in G12D and G12V SW48 cells show striking similarities to the consensus molecular subtype 3 (CMS3) of colorectal cancer types, which show an association between increased KRAS mutations and elevated metabolic signature of glutamine and nitrogen metabolism^55^.

We note that the exchange of metabolites between different cell types can significantly impact cellular viability within tissues in physiological conditions and pathology. Notable examples are the glutamate-glutamine cycle between neurons and glia, but also exchanges of glutamine between cancer-associated fibroblasts and cancer cells that support further tumour progression^56–58^. It is thus intriguing to observe that differential regulation of nitrogen metabolism confers resilience to G12D and G12V SW48 clones and, at the same time, alters the exchange of metabolites to and from the extracellular space. Although beyond the scope of our work, these observations raise the possibility that the exchange of metabolites between mutant cells and the tissue microenvironment could favour the emergence of specific mutations in permissive tissues.

The involvement of FOXO1 in colorectal cancers and KRAS-induced metabolic reprogramming is not well characterised. Interestingly, gains and amplifications of FOXO1 are evident in colorectal cancers, and levels of FOXO1 show a moderate increase with advanced tumour stages of colorectal cancer (Figure S6). Our results suggest that oncogenic KRAS can upregulate FOXO1. More importantly, differences in the GTPase activity of KRAS or interaction with other proteins (*e.g.*, AKAP12-PKA axis^59,60^, Figure S1C-D) can alter the localisation of FOXO1 irrespective of its expression. We show a correlation between the nuclear localisation of FOXO1 and glutamine metabolism, and that inhibition of FOXO1 abrogates the KRAS-dependent upregulation of glutamine synthesis and transamination features. Our work suggests that FOXO1 inhibition might be a viable therapeutic route.

However, the inhibition of FOXO1 using AS1842856 is highly toxic for all cell lines we tested, including those wild-type for KRAS. We, therefore, targeted glutamine-glutamate metabolism, altered in KRAS mutants, using the glutamine synthetase inhibitor methionine sulfoximine (MSO) and the glutaminase inhibitor CB839 to avoid the toxicity that might result from targeting a key transcription factor such as FOXO1. Individually, MSO and CB839 show no or moderate selectivity towards mutant cells, respectively (Figure 5 and S5). This observation is largely unsurprising because isogenic panels of cell lines engineered from cancer cell lines are notoriously difficult to kill with specificity.

However, we show that MSO and CB839 work synergistically in the background of SW48 and LIM1215 colorectal cancer cells. In low nutrient conditions where mutant cell lines manifest enhanced fitness, low doses of MSO and CB839 are sufficient to abolish their growth advantage and selectively kill mutant lines (Figure 5 and S5). Differential regulation of glutamine metabolism and its importance in tumorigenesis is well-documented^11,61–63^, making it the optimal candidate for therapy. Treatment strategies like inhibition of GLUL^63^, treatment with glutamine analogue (DON)^64^, or using the GLS inhibitor CB839^65^ are promising in PDAC mouse models; however, alternative compensatory mechanisms have often led to the emergence of resistance. Metabolic pathways are very resilient and regulated to provide the appropriate metabolite levels utilising different pathways. Our results suggest that the pharmacological disruption of glutamine-glutamate metabolism at both sides of the glutamine-glutamate cycle is a promising strategy for the selective elimination of KRAS mutant cells. While CB839 is already clinically approved, MSO has limited clinical use because of its convulsive effects *in vivo*. In the absence, to our knowledge, of alternative inhibitors for glutamine synthetase that are clinically approved, our results indicate the necessity to investigate the use of low doses of MSO in conjunction with therapeutic doses of CB839 or at least the opportunity to direct drug discovery programmes in identifying new GS inhibitors.

In summary, our work suggests that the glutamine-glutamate cycle provides a growth advantage to KRAS mutant cells, thus representing a potentially attractive new target for drug discovery.

### Limitations of the study

Our study was performed by culturing monolayers of human cancer cells, which permitted us to control experimental conditions better and utilise an appropriate genetic background. However, studies carried out in cell culture cannot recapitulate aspects of three-dimensional tissues and *in vivo* experimentation. To partially mitigate these shortcomings, we have performed most of our experiments in nutrient conditions similar to interstitial fluid rather than typical media formulated to stimulate the indefinite growth of cell lines and compared our results to experiments done in full media.

The cell lines we used (SW48) are a commercially available panel of isogenic lines. SW48 cells are well-characterised and used as a platform to investigate differences between oncogenic alleles. However, cell lines in culture conditions can drift and adapt, manifesting phenotypes that might not represent genuine differences related to the specific mutations we analysed. To mitigate these issues, we have adopted several strategies. First, all our cell lines are not only STR profiled to avoid contamination with other cell types we culture in the laboratory but have been routinely genotyped to ensure contamination between different mutants does not occur. All replicates were done from different flasks and several stocks. RNA sequencing was performed with five technical replicates of RNA extracted from cultures in different flasks, including different passages. This permitted us to exclude any phenotype that was insufficiently robust to culturing conditions. More importantly, as it was impossible to procure independently generated and fully validated SW48 clones of the same mutants, we have repeated key experiments using a different isogenic panel in the background of LIM1215, substantiating the robustness of our observations.

## RESOURCE AVAILABILITY

### Lead contact

Further information and requests for resources should be directed to and will be fulfilled by the lead contact, Alessandro Esposito

### Materials availability

All the materials generated in this study are available to the lead contact upon reasonable request to the lead contact.

### Data and code availability

- All the RNA-seq results and analysis software is available in Supplemental File S1.
- All the LC-MS results are available in Supplemental File S2.
- Any additional information required to reanalyse the data reported in this paper is available from the lead contact upon request.

## ACKNOWLEDGEMENTS

A.E. acknowledges the financial support provided by the CRUK with a multi-disciplinary project award (OncoLive, C54674/A27487), pump-priming funds from the CRUK Cambridge Center (C9685/A25117, C9685/A28397). A.E. and A.R.V. also acknowledge financial support from Medical Research Council program grants (MC_UU_12022/1 and MC_UU_12022/8). A.E. also acknowledges funding from the European Union for HORIZON 2022 fLIMAGING3D (101073507) and HILIGHT (101135034, backed by Innovate UK as project number 10107542). C.F. was funded by the MRC Core award grant MRC_MC_UU_12022/6, and is funded by the CRUK Programme Foundation award C51061/A27453, the H2020 European Research Council Consolidator Grant (ERC819920), and by the Alexander von Humboldt Foundation in the framework of the Alexander von Humboldt Professorship endowed by the Federal Ministry of Education and Research. Work in Simon Cook’s laboratory was supported by Institute Strategic Programme Grants BB/J004456/1, BB/P013384/1 and BB/Y006925/1 from UKRI-BBSRC. We want to thank Dr Laura Tronci (CF group), Dr Pablo Oriol Valls and Dr Andrew Trinh (AE group) for performing experiments in relation to the study of KRAS mutant cell lines that have not been included in this work. The authors are indebted to the many colleagues in the professional and technical services at the MRC Cancer Unit for making this research possible. Also, the principal investigators express their gratitude towards all members of their respective teams and support staff who endured difficulties related to the COVID lockdowns and the eventual termination of the MRC Cancer Unit.

## AUTHOR CONTRIBUTIONS

Conceptualisation: S.B., A.E., A.R.V., C.F., and M.S.; software: S.S., A.E.; validation: S.B.; formal analysis: S.B., A.E., S.S., and M.Y.; investigation: S.B., K.P., A.H., M.Y., and E.N.; data curation: A.E.; writing – original draft: S.B. and A.E.; writing – review and editing: S.B., A.E., M.S., A.R.V. and C.F.; visualisation: S.B. and A.E.; supervision: S.B., A.E., A.R.V., and C.F.; project administration: A.E.; funding acquisition: A.E., A.R.V. and C.F.

## STAR★METHODS

### KEY RESOURCES TABLE

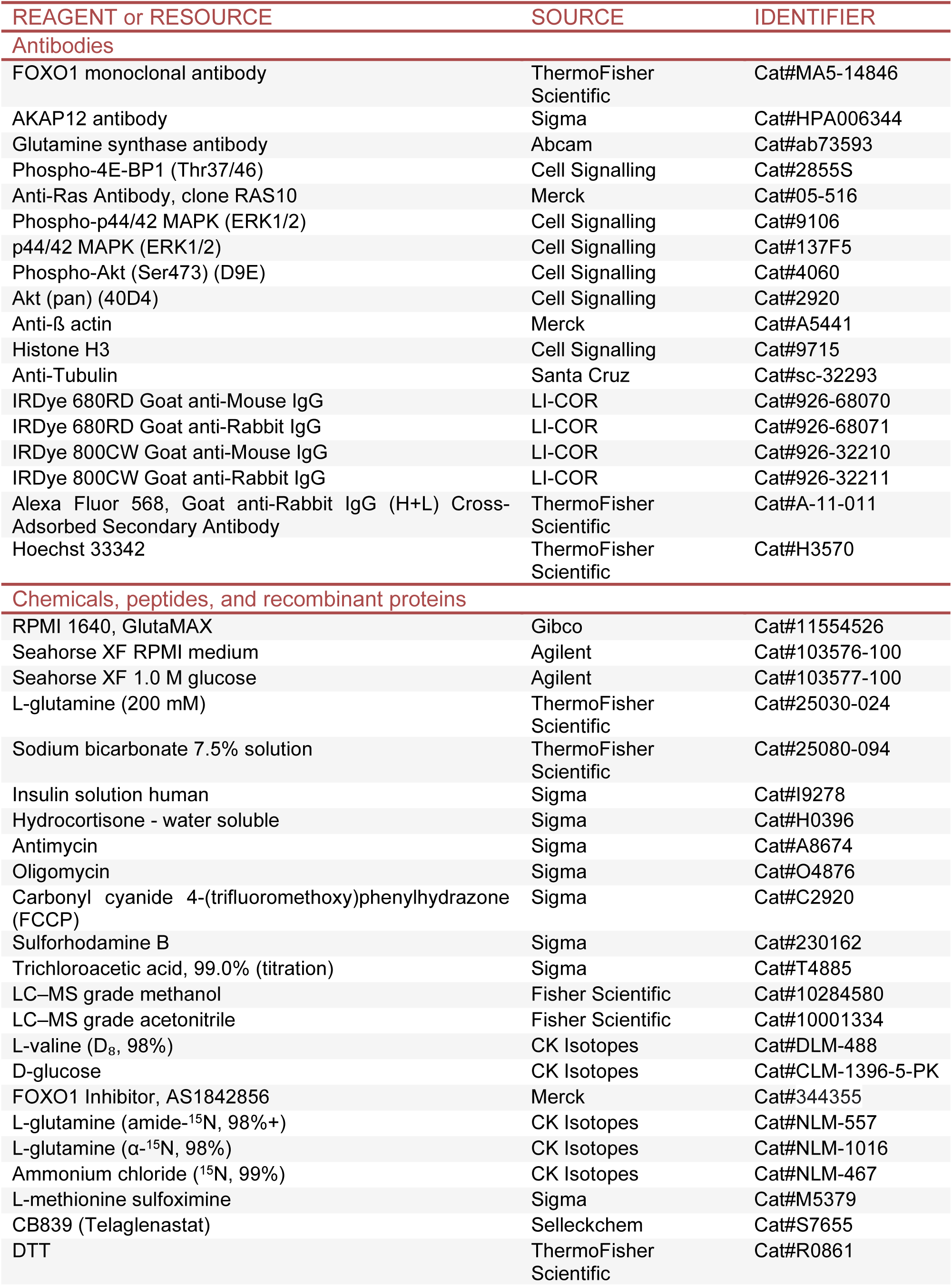
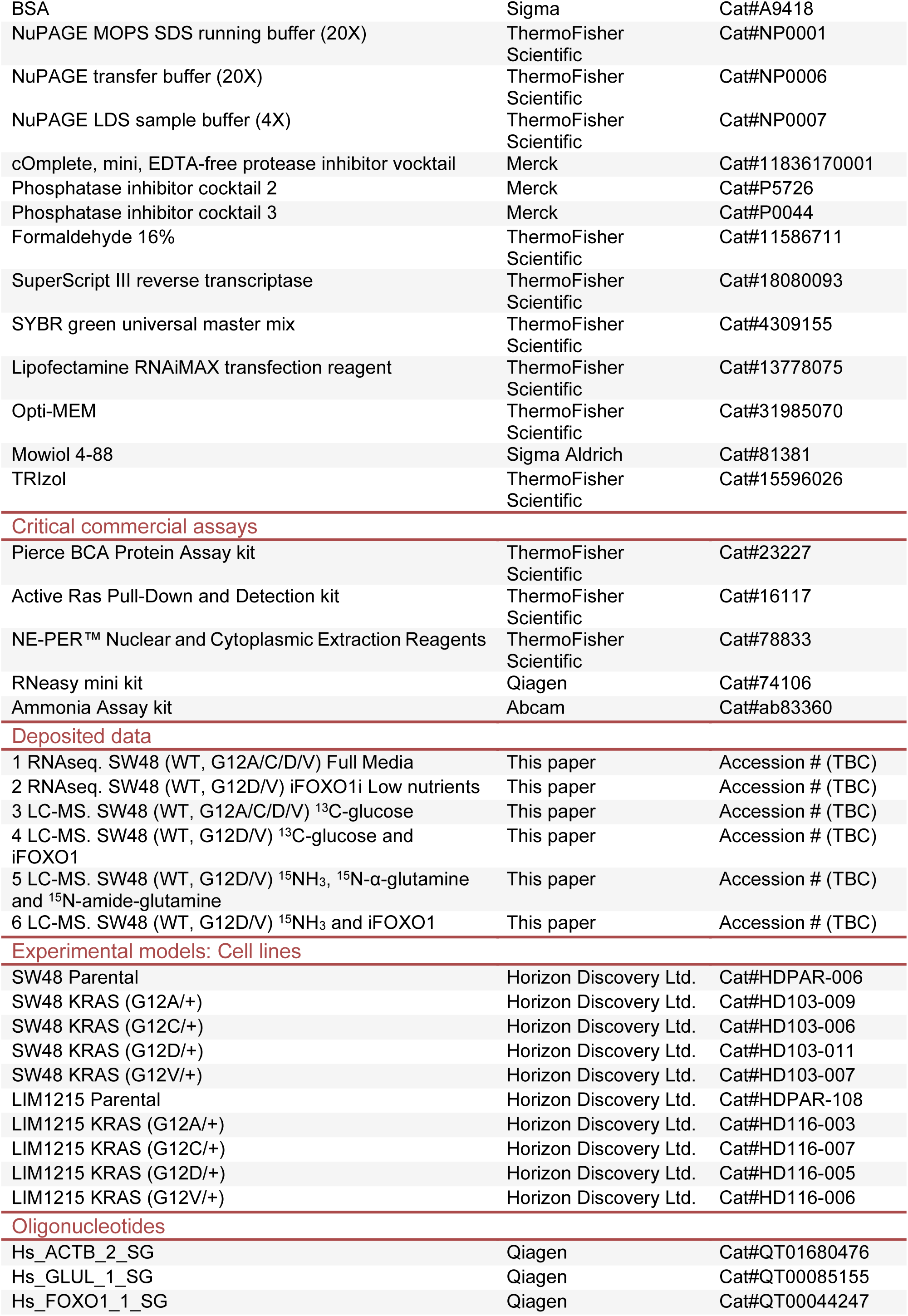
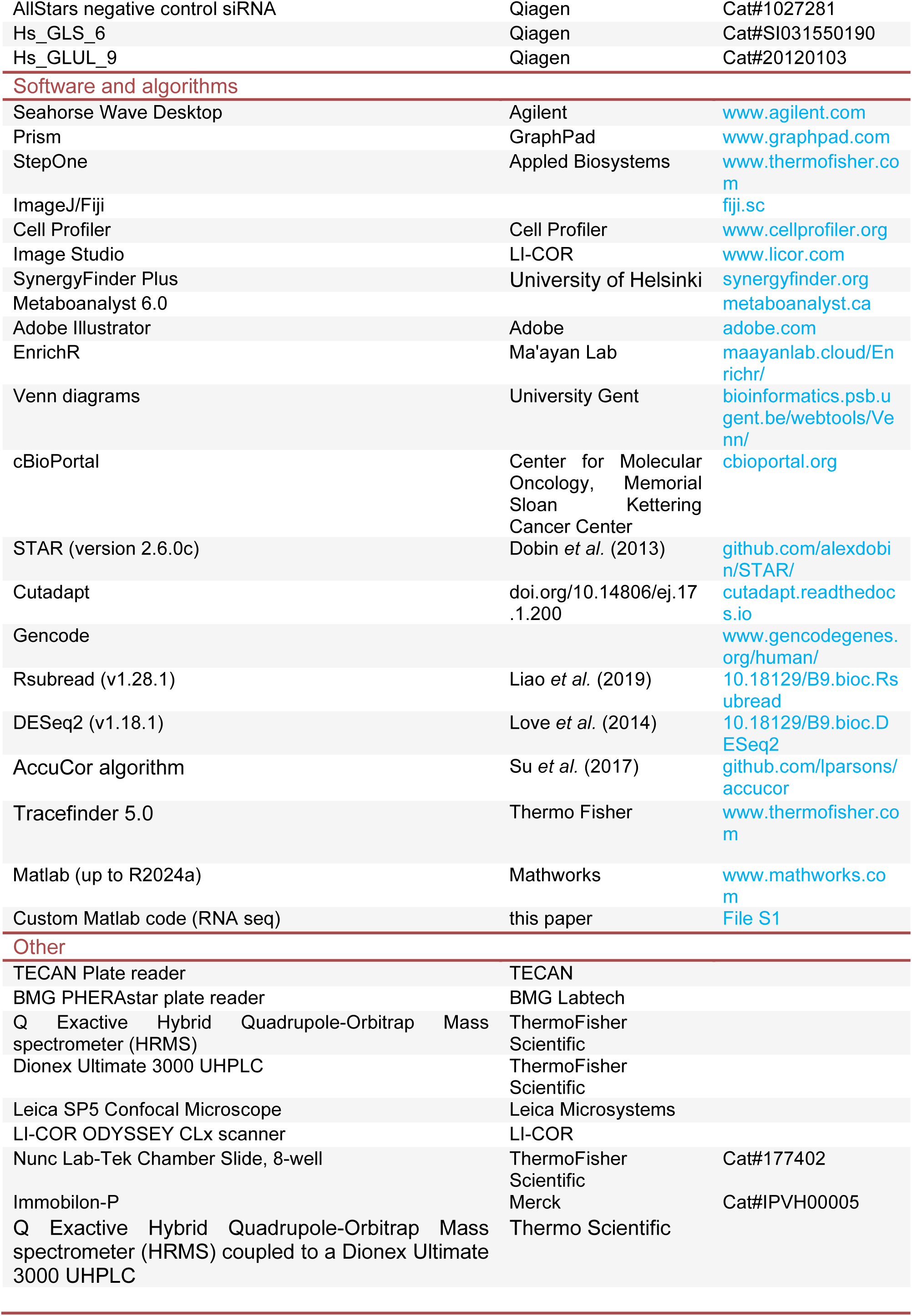

### METHOD DETAILS

#### Cell Culture

SW48 isogenic cells (SW48^+/+^, SW48^+/G12A^, SW48^+/G12C^, SW48^+/G12D^, SW48^+/G12V^), derived from colorectal adenocarcinoma were purchased from Horizon Discovery. Cells were cultured in their recommended conditions, also referred in the text as full media conditions, RPMI 1640 with GlutaMAX, HEPES (Gibco, # 11554526) and 10% foetal bovine serum (FBS; Gibco, #A5256701). LIM1215 isogenic cells (LIM1215^+/+^, LIM1215^+/G12A^, LIM1215^+/G12C^, LIM1215^+/G12D^, LIM1215^+/G12V^), derived from human colorectal carcinoma, were purchased from Horizon Discovery. Cells were cultured in their recommended media, RPMI 1640 with GlutaMAX, HEPES (Gibco, # 11554526) supplemented with 10% FBS, 1μg/ml insulin (Sigma, I9278), and 1μg/ml hydrocortisone (Sigma, H0396). Seahorse XF RPMI medium (Agilent, #103576-100) supplemented with 2 mM Glucose (Agilent, #103577-100), 2mM Glutamine (ThermoFisher ScientificTM, #25030-024), 0.2% sodium bicarbonate (ThermoFisher ScientificTM, #25080-094) and 1% FBS was used for experiments carried out in low nutrient conditions with both cell lines.

#### Seahorse Assay

For mitochondrial stress test, Agilent Seahorse XF Assays were performed in a 24-well XF Cell Culture Microplate using Seahorse XFe24 analyser (Agilent Technologies). Briefly, SW48 cells were plated at a density of 2×10^5^ cells/well in 24-well XF microplates (Agilent Technologies) to achieve a monolayer confluency. Next day, 1 hour prior to the experiment the media was replaced with 675μl/well Seahorse XF RPMI Medium (Agilent Technologies) supplemented with 1% FBS, 11mM Glucose and 2mM Glutamine, mimicking the original nutrient concentrations of full media. During the assay, the following compounds were injected sequentially, 75 μl Oligomycine (10 μM) (Sigma), 83 μl FCCP (5 μM) (Sigma), 92μl Antimycine (10 μM) (Sigma) and Rotenone (10 μM) (Sigma) together, and OCR and ECAR values were measured. At the end of the assay cells were harvested using 30 μl of RIPA buffer/well and total protein amount was measured using Pierce BCA Protein colorimetric assay (ThermoFisher Scientific). OCR and ECAR values were then normalized to total protein levels and graphs plotted using Graphpad.

#### Sulforhodamine-B (SRB) Viability Assay

For SRB assays both in full media or low nutrient conditions, SW48 isogenic cells were initially seeded on a 96-well plate ad a 7×10^3^ cells/well density in full media conditions. 24 hours after seeding and complete attachment, media was replaced with either full media or low nutrient media. After further 72 hours in culture, cells were used for the SRB assay. Briefly, cells were washed once with PBS and fixed with 100μl/well 1% (v/v) trichloroacetic acid (Sigma) for 2 hours at 4°C. Fixed cells were washed twice with deionized water and stained with 100μl/well Sulforhodamine B (Sigma) solution (0.057% w/v in 1% acetic acid) for 1 hour at room temperature. SRB was then removed, plates washed with 1% acetic acid 3 times and air dried. SRB stain was solubilized with 10 mM Tris base for 10 minutes. Absorbance was measured at 540 nm using a 96-well plate reader (BMG Pherastar Plus) and results calculated as percentage survival using Graphpad.

#### Liquid chromatography coupled to mass spectrometry (LC–MS)

HILIC chromatographic separation of metabolites was achieved using a Millipore Sequant ZIC-pHILIC analytical column (5 µm, 2.1 × 150 mm^2^) equipped with a 2.1 × 20 mm^2^ guard column (both 5 mm particle size) with a binary solvent system. Solvent A was 20 mM ammonium carbonate, 0.05% ammonium hydroxide; Solvent B was acetonitrile. The column oven and autosampler tray were held at 40 °C and 4 °C, respectively. The chromatographic gradient was run at a flow rate of 0.200 ml/minute as follows: 0–2 minutes: 80% B; 2–17 minutes: linear gradient from 80% B to 20% B; 17–17.1 minutes: linear gradient from 20% B to 80% B; 17.1–22.5 minutes: hold at 80% B. Samples were randomised and analyzed with LC–MS in a blinded manner with an injection volume was 5 µl. Pooled samples were generated from an equal mixture of all individual samples and analysed interspersed at regular intervals within the sample sequence as quality control.

Metabolites were measured with a Thermo Scientific Q Exactive Hybrid Quadrupole-Orbitrap Mass spectrometer (HRMS) coupled to a Dionex Ultimate 3000 UHPLC. The mass spectrometer was operated in full-scan, polarity-switching mode, with the spray voltage set to +4.5 kV/–3.5 kV, the heated capillary held at 320 °C, and the auxiliary gas heater held at 280 °C. The sheath gas flow was set to 25 units, the auxiliary gas flow to 15 units, and the sweep gas flow was set to 0 units. HRMS data acquisition was performed in a range of *m/z* = 70– 900, with the resolution set at 70,000, the AGC target at 1 × 10^6^, and the maximum injection time (Max IT) at 120 ms. Metabolite identities were confirmed using two parameters: (1) precursor ion m/z was matched within 5 ppm of theoretical mass predicted by the chemical formula; (2) the retention time of metabolites was within 5% of the retention time of a purified standard run with the same chromatographic method.

#### LC–MS metabolomics: tracing experiments

For the glucose tracing experiment, SW48 isogenic cells were plated on 6-well plates at a density of 8×10^5^ cells/well in full media conditions (5 replicates for each cell line). The next day, after a PBS wash, the media was changed into low nutrient media where 2 mM glucose was substituted with equimolar concentration of ^13^C-labelled glucose (D-glucose, CK Isotopes) and cultured for another 24 hours before extraction.

For extraction, as a first step cell numbers was determined for each cell line using a separate plate for counting. Cells were washed twice in PBS and incubated in PBS on cold bath (dry ice and methanol). PBS was then replaced with metabolite extraction buffer (MEB, 50% LC– MS grade methanol (Fisher Scientific), 30% LC–MS grade acetonitrile (Fisher Scientific) and 20% ultrapure water), 1 ml extraction buffer for 1×10^6^cells; after a couple of minutes on the cold bath, cells were stored at −80 °C overnight. The next day, extracts were scraped and mixed agitating for 15 minutes at 4 °C at maximum speed (3000rpm) .Finally, extracts were centrifuged for 10 minutes at maximum speed (14000rpm) at 4 °C and transferred into LC– MS vials for analysis. Valine-d8 5 μM (CK isotopes) was used as internal standard for the MEB.

For the ^13^C-glucose labelling experiments with the FOXO1 inhibitor (AS1842856) (Merck), SW48 isogenic cells were plated on 6-well plates at a density of 8×10^5^ cells/well in full media conditions (5 replicates for each cell line). 24 hours later cells were pre-treated with DMSO (0.003% v/v) control or FOXO1 inhibitor (1 µM) for 12 hours in full media conditions. Cells were then washed with PBS and media replaced with low nutrient media where 2mM glucose was substituted with equimolar concentration of ^13^C-labelled glucose (D-glucose, CK Isotopes) and DMSO (0.003% v/v) control and FOXO1 inhibitor (1 µM) were replenished for further 12 hours until extraction.

To assess the immediate metabolic fluxes, we used 4 hour tracing in all nitrogen tracing experiments. For nitrogen labelling experiments, SW48 isogenic cells were plated on 6-well plates at a density of 8×10^5^ cells/well in full media conditions (5 replicates for each cell line). 24 hours later, cells were washed once with PBS and cultured in low nutrient media where 2mM glutamine was substituted with equimolar concentrations of either L-glutamine amide-^15^N (CK Isotopes) or L-glutamine α-^15^N (CK Isotopes) for 4 hours before extraction.

For the ammonia labelling experiments, SW48 isogenic cells were plated on 6-well plates at a density of 8×10^5^ cells/well in full media conditions (5 replicates for each cell line). 24 hours later, cells were washed once with PBS and cultured in low nutrient media containing 3 mM of ^15^N-labelled ammonium chloride (CK Isotopes) for 4 hours before extraction. We repeated this experiment where 24 hours after seeding, cells were pre-treated with DMSO (0.003% v/v) control or FOXO1 inhibitor (1 µM) for 12 hours in full media conditions. Cells were then washed with PBS and media replaced with low nutrient media containing labelled ammonia with either DMSO control (0.003% v/v) or FOXO1 inhibitor (1 µM) for further 4 hours before extraction.

#### Cell treatments

LIM1215 and SW48 cells were seeded at 3×10^3^ cells/well and 7×10^3^ cells/well densities, respectively. 24 hours after seeding and complete attachment, media was replaced with either full media or low nutrient media containing the single drug or combined drug treatments below and cultured for an additional 72 hours.

For glutamine treatments, serial two-fold dilution of glutamine (maximum concentration 5mM) was performed in full media or low nutrient media.

For drug treatments in low nutrient conditions, serial two-fold dilution of GLS inhibitor (CB839, maximum concentration of 100 μM, and DMSO control of maximum 0.05% v/v) or GLUL inhibitor (MSO, maximum concentration of 100 mM dissolved in low nutrient media) was performed. For drug combination experiments, serial dilution of CB839 (Selleckchem) or MSO (Sigma) at concentrations above was performed in the presence or absence of sublethal doses of MSO (2 mM) or CB839 (100 nM), respectively. Water (1% v/v for MSO) or DMSO (0.05% v/v for CB839) were used as vehicle controls.

For single dose drug treatments, we used the FOXO1 inhibitor (AS1842856) at 100 nM, MSO at 2 mM, CB839 at 100 nM. Water (1% v/v for MSO), DMSO (0.05% v/v for CB839 and 0.003% v/v for AS1842856) were used as vehicle control.

The BLISS assay was also performed with the same protocol using 96-well plates using a two-fold dilution series of MSO (highest concentration of 25 mM) and CB839 (highest concentration 12.5 μM) in an 8 × 8 checkerboard pattern of combinations^66^. The BLISS assay analysis was performed with the online Synergy Finder tool (www.synergyfinder.org ).

At the end of the 72h growth assay, all cells were fixed and subjected to the SRB assay described in the methods. Cell viability is represented as the percentage survival normalised to all controls indicated above.

#### Western blots

Cells were extracted from 6-well plates using 80 µl/well RIPA buffer (300 mM NaCl, 1.0% NP-40, 0.5% sodium deoxycholate, 0.1% SDS (sodium dodecyl sulphate), 50 mM Tris-HCl, pH

8.0 with protease and phosphatase inhibitor 2/3 cocktails). Lysates were incubated on ice, agitating every 10 minutes for 30 minutes. Lysates were then centrifuged for 10 minutes at 14000rpm at 4°C and the supernatant processed for protein quantification using Pierce BCA protein colorimetric assay (ThermoFisher Scientific) following the manufacturer’s instructions. Absorbance was measured with a TECAN spectrophotometer at 562 nm. Proteins were diluted in 1x NuPAGE LDS Sample Buffer with final of 80 mM DTT (Thermo Scientific) and heated at 95°C for 5 minutes. Samples were ran on 4-12% NuPAGE Bis-Tris gels (Thermo Scientific) at constant 120V for 1 hour in 1x NuPAGE MOPS SDS Running Buffer (Thermo Scientific). Wet transfer of the proteins to a PVDF (polyvinylidene difluoride) membrane (Immobilon-P, Merck) was done using BioRad transfer system (Mini Trans-Blot® Cell, Bio-Rad). Membranes were then stained with Ponceau to assess protein transfer. Afterwards, the membranes were blocked for 1 hour with blocking buffer (5% BSA or 5% milk in TBST (Tris-Buffered Saline and 0.1% Tween). Primary antibody incubation was done overnight at 4°C.

Primary antibodies used are FOXO1 monoclonal antibody (ThermoFisher Scientific, 1:1000), AKAP12 (Sigma, 1:1000), glutamine synthase (Abcam, 1:1000), anti-RAS antibody (Merck, 1:1000), phospho-4E-BP1 Thr37/46 (Cell Signalling, 1:1000), phospho-p44/42 MAPK (ERK1/2; Cell Signalling, 1:1000), p44/42 MAPK (ERK1/2; Cell Signalling, 1:1000), phospho-Akt Ser473 (clone D9E; Cell Signalling, 1:1000), Akt (pan, clone 40D4; Cell Signalling, 1:1000), anti-ß actin (Merck, 1:1000), histone H3 (Cell Signalling, 1:1000), alpha-tubulin (Santa Cruz, 1:1000).

The next day, membranes were washed 4 times in TBST and incubated with LI-COR secondary antibodies (LI-COR, IRDye-680 or IRDye-800; anti-mouse and anti-rabbit) diluted 1:10000 in blocking buffer for 1 hour at room temperature. After 4 washes with TBST, membranes were imaged using LI-COR Odyssey CLx scanner.

#### Active RAS Pull-Down Assay

This assay was performed using Active RAS Pull-Down and Detection Kit (ThermoFisher Scientific) according to the manufacturer’s instructions. For this assay, 1×10^7^ SW48 cells were seeded on 10cm dishes. After 24h, cells were washed with 1xPBS and harvested using 500μl/dish 1X Lysis/Binding/Wash Buffer (provided in the kit). Lysates were kept on ice, agitating every 10 minutes for 30 minutes. Lysates were then centrifuged at 14000 rpm at 4°C, and the supernatant was processed for protein quantification using Pierce BCA colourimetric assay (ThermoFisher Scientific) following the manufacturer’s instructions. 100μl of Glutathione resins (provided in the kit) were used per sample, the supernatant was discarded, and 400 μl of 1X Lysis/Binding/Wash Buffer with 80 μg of GST-RAF1-RBD was added to the beads in each tube. An additional 700 μl of 1X Lysis/Binding/Wash Buffer with a total of 1.5 mg protein was added to the beads in each tube and incubated overnight at 4°C with gentle rotation. The next day, samples were washed 4 times with 1X Lysis/Binding/Wash Buffer using the columns provided in the kit. The protein was then eluted from the column with 50 μl of reducing sample buffer (2.5 μl ß-mercaptoethanol in 50 μl 2x SDS buffer (provided by the kit)). Eluted samples were then heated for 5 minutes at 95°C and run on 4-12% Bis-Tris gel (ThermoFisher Scientific) as described earlier. Samples were then blotted for total Ras (Merck) and Actin (Merck).

#### Nuclear and cytoplasmic fractionation assay

This assay was performed using NE-PERNuclear and Cytoplasmic Extraction kit (ThermoFisher Scientific). Briefly, SW48 cells were seeded on a 6-well plate (1×10^6^ cells/well). 24 hours later, cells were washed with 1x PBS and collected by scraping with 80µl/well CER-I buffer containing protease (Merck) and phosphatase (Merck) inhibitors. Cells were vortexed for 15 seconds and placed on ice for 15 minutes. 4.4 μl/sample CER-II buffer was added, vortexed for 5 seconds and placed on ice for 2 minutes. After another vortex step, samples were centrifuged for 5 minutes on a benchtop centrifuge at maximum speed (14000 rpm) at 4°C. The supernatant (the cytosolic fraction) was transferred into a clean tube, and the pellet (the nuclear fraction) was resuspended in 40μl of ice-cold RIPA buffer described earlier. Resuspended nuclear fractions were kept on ice for 1 hour during which samples were vortexed for 15 seconds every 10 minutes. Samples were then centrifuged at maximum speed (14000 rpm) for 10 minutes at 4°C on a benchtop centrifuge. Samples were diluted in 1x NuPAGE LDS Sample Buffer with a final of 80 mM DTT (ThermoFisher Scientific), heated at 95°C for 5 minutes, and processed further for western blotting.

#### Immunostaining

SW48 (4 10^4^ cells/well) or LIM1215 (2 10^4^ cells/well) cells were seeded on 8-well removable LabTek chambers (ThermoFisher Scientific). 24 hours later, cells were washed with PBS, and the media was changed to low-nutrient media. After another 24 hours, cells were fixed with 4% PFA (ThermoFisher Scientific) for 15 minutes at room temperature, washed with PBS and blocked with blocking buffer (1% BSA, 0.1% Triton-X, 5% Goat serum in PBS) for 30min. Cells were then stained with FOXO1 monoclonal antibody (ThermoFisher Scientific, 1:200) diluted in blocking buffer overnight at 4°C. The next day, cells were washed with PBS and incubated with a secondary antibody (AlexaFluor 568 conjugated goat anti-rabbit IgG (ThermoFisher Scientific, 1:500)) and Hoechst 33342 (ThermoFisher Scientific, 1:1000) in blocking buffer for 1 hour at room temperature. Cells were washed 3 times with PBS, wells were removed, and the sample was mounted with Mowiol (5% w/v Mowiol 4-88, 12% w/v glycerol in 2:3 dilution of 0.2M Tris buffer, pH 8.5, in water). Cells were imaged using 63X objective on Leica SP5 Confocal (SP5, Leica Microsystems). Nuclear quantification of FOXO1 was done using Cell Profiler software.

#### RNA extraction and real-time qPCR

SW48 isogenic cells were plated on 6-well dishes 8×10^5^ cells/well in full media conditions. For drug treatments, 24 hours after seeding, cells were treated with either DMSO (0.003% v/v) vehicle control or FOXO1 inhibitor (1 μM) in full media conditions. After 24 hours of treatment, RNA was extracted using RNeasy kit (Qiagen) following manufacturer’s instructions. 1 μg RNA was reverse-transcribed using SuperScript III Reverse Transcriptase kit (ThermoFisher Scientific). qPCR was performed using SYBR Green PCR Master Mix (ThermoFisher Scientific), and following primers Hs_ACTB_2_SG (Qiagen), Hs_GLUL_1_SG (Qiagen), Hs_FOXO1_1_SG (Qiagen). Data analysis was done using StepOne software.

#### siRNA

Lipofectamine RNAiMAX Transfection Reagent (ThermoFisher Scientific) via reverse transfection was used for knockdown experiments, following the manufacturer’s protocol. Briefly, master mix solution of Lipofectamine RNAiMAX, Opti-MEM (Thermo Scientific) and final 10 nM/well concentration of AllStars negative control siRNA (Qiagen), Hs_GLS_6 (Qiagen), Hs_GLUL_9 (Qiagen) siRNAs were prepared and after 15 minutes of incubation were aliquoted into a 96 well dish. SW48^+/+^, SW48^+/G12D^ and SW48^+/G12V^ cells were trypsinised, centrifuged and seeded in full media at a density of 7×10^3^ cells/well within the wells containing the siRNA mix. The next day, media was changed to low nutrient media and cells were kept for 72 hours in culture and viability was assessed using SRB assay as described above.

#### Ammonia assay

SW48^+/+^, SW48^+/G12D^ and SW48^+/G12V^ cells were seeded on a 6-well dish (5×10^5^ cells/well) in full media conditions. 48 hours later, the media was changed to low nutrient conditions for another 24 hours. Cells were then washed with PBS and lysed in 100 μl of ammonia assay buffer provided with the Ammonia assay kit (Abcam). Standards and samples were prepared together with the reaction mix and incubated for 1 hour. Absorbances were measured with a TECAN spectrophotometer at 570 nm. Ammonia production was calculated using the protocol provided by the manufacturer.

#### RNA sequencing

SW48 isogenic cells were thawed and cultured for 3 passages before sample preparation. 5 replicates from 3 different batches of each cell line were used for the RNAseq experiment. Briefly, 1.5 million cells were seeded on a 6 cm dish in full media conditions. 48 hours later, cells were washed once with cold PBS and collected by scraping with 1 ml PBS into 1.5 ml Eppendorf tubes. Samples were centrifuged for 3 minutes at 1,000 rpm and 4 °C. Pellets were stored at -80 °C until shipment. RNA extraction, library preparation and sequencing (20 million reads per sample) was performed by BGI Genomics (BGI Hong Kong Company Limited).

For samples treated with FOXO1 inhibitor, SW48^+/+^, SW48^+/G12D^ and SW48^+/G12V^ cells were thawed and cultured for 3 passages before sample preparation. 5 replicates from 3 different batches of each cell line were used for the RNAseq experiment. Cells were seeded at 1.5 million cells/well on 6cm dishes in full media conditions. 24 hours later, media was replaced with low nutrient media containing DMSO (0.003% v/v) vehicle control, FOXO1 inhibitor (1 μM) or just media control. After an additional 24 hours in low nutrient conditions, cells were scraped in 400 μl/well TRIzol (ThermoFisher Scientific) and stored at -80 °C until shipment. RNA extraction, library preparation and sequencing (20 million reads per sample) was performed by BGI Genomics (BGI Hong Kong Company Limited).

### QUANTIFICATION AND STATISTICAL ANALYSIS

#### RNA sequencing data analysis

Reads were mapped to the human reference genome GRCh38 with the STAR (version 2.6.0c) splice-aware aligner^67^ using annotation from Gencode (releases 24-35). Low-quality reads (mapping quality <20) as well as known adapter and artifact contaminations were filtered out using Cutadapt (version 1.10.0). Read counting was performed using Bioconductor package Rsubread (v1.28.1)^68^ and differential expression analysis with DESeq2 (v1.18.1)^69^.

RNA sequencing data was visualised with custom Matlab (Mathworks) scripts, which are available in File S1. The thresholds for log2 fold changes and false discovery rates are reported in the caption of each figure.

#### Metabolomics data analysis

The chromatograms generated by LC-MS were reviewed, and the peak area was integrated using the Thermo Fisher software Tracefinder 5.0. The peak area for each detected metabolite was normalised against the total ion count (TIC) of that sample to correct any variations introduced from sample handling through instrument analysis. The normalised areas were used as variables for further statistical data analysis. For ^13^C- and ^15^N-tracing analysis, the theoretical masses of ^13^C and ^15^N isotopes were calculated and added to a library of predicted isotopes. These masses were then searched with a five ppm tolerance and integrated only if the peak apex showed less than a 1% difference in retention time from the [U-^12^C and U-^14^N] monoisotopic mass in the same chromatogram. After analysis of the raw data, natural isotope abundances were corrected using the AccuCor algorithm^70^.

The resulting Microsoft Excel files are available in File S2.

#### Statistical analysis

Graphs and statistical tests were done using GraphPad Prism 8-10 (GraphPad, San Diego, CA). Details of data analysis are reported in figure captions. Statistical significance was evaluated by one-way ANOVA using Dunnets’s test, or two-way ANOVA using Sidak’s or Tukey’s tests for multiple comparisons. Technical and biological repeats were performed as indicated in figure legends, and results were reported as mean with standard deviation (SD) or standard error of the mean (SEM). All experiments, except for transcriptomics and metabolomics (see respective sections), were performed at least with three independent replicates.

## SUPPLEMENTAL INFORMATION

**Supplementary Table 1.** Differentially regulated gene sets in SW48 cell lines. Every gene set was analysed using thresholds for false discovery rates (FDR) of 5% and a minimum 2-fold change. Links to gene enrichment analysis carried out with *EnrichR*^71^ are embedded in the table, and data is available in File S1.

| Genes differentially regulated in | Number of genes | Jupyter file in File S1 (Table S1 subfolder) | Link |
| --- | --- | --- | --- |
| SW48 G12A | 851 | 1 | EnrichR |
| SW48 G12C | 482 | 2 | EnrichR |
| SW48 G12D | 1229 | 3 | EnrichR |
| SW48 G12V | 558 | 4 | EnrichR |
| at least one mutant cell line (Fig. 1A) | 1,935 | 5 | EnrichR |
| all mutant cell lines with similar trends (Fig. S1G left) | 97 | 6 | EnrichR |
| in G12D and G12V with similar trends (Fig. S1G right) | 334 | 7 | EnrichR |
| orange cluster in Fig. S1G right | 113 | 8 | EnrichR |

**Supplementary Table 2.** List of transcriptomics and metabolomics experiments. List of transcriptomics and metabolomics experiments with conditions and figures where the data is used. Data analysis and matbal scripts used to generate figures are provided in File S1-S2.

| Id | Cell lines | Drugs | Media | Type | Figures | Files |
| --- | --- | --- | --- | --- | --- | --- |
| 1 | SW48 (WT, G12A/C/D/V) | - | Full media | RNAseq | 1A-C and S1E-H, 2E | S1 |
| 2 | SW48 (WT, G12A/C/D/V) | - | Low nutrient | RNAseq | 4C | S1 |
| 3 | SW48 (WT, G12D and V) | DMSO and iFOXO1 | Low nutrient | RNAseq | 4D, S3C | S1 |
| 4 | SW48 (WT, G12A/C/D/V) | - | Low nutrient | <sup>13</sup>C-glucose LC-MS | 2A-B, S2A-B, S2D, S2F | S2 |
| 5 | SW48 (WT, G12D and V) | DMSO and iFOXO1 | Low nutrient | <sup>13</sup>C-glucose LC-MS | 3F, S3B | S2 |
| 6 | SW48 (WT, G12D and V) | - | Low nutrient | <sup>15</sup>N-ammonia, <sup>15</sup>N-alpha-glutamine and <sup>15</sup>N-amide-glutamine LC-MS | 4A-B, S4A-C | S2 |
| 7 | SW48 (WT, G12D and V) | DMSO and iFOXO1 | Low nutrient | <sup>15</sup>N-ammonia LC-MS | 4E | S2 |

**Supplementary Table 3.**
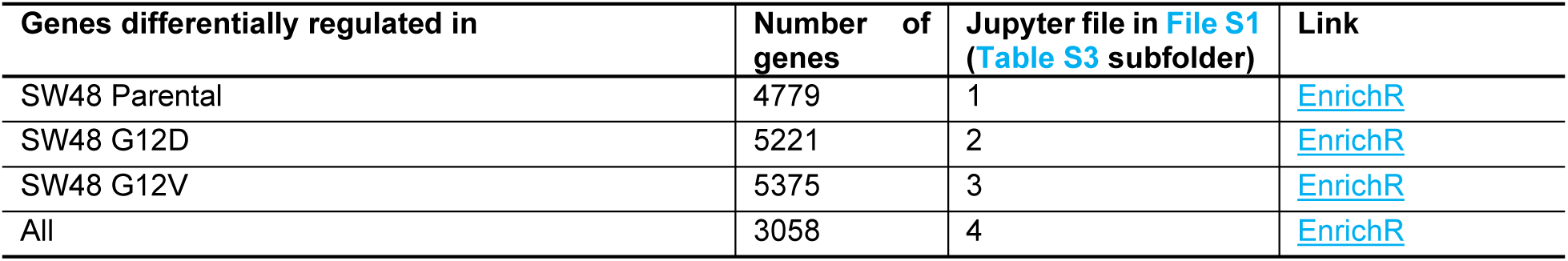
AS1842856 downregulates FOXO1 gene targets. List of transcripts downregulated (FDR<10% and log2 fold change less than -0.2) upon treatment with AS1842856. Gene enrichment analysis performed with EnrichR confirms that this small molecule inhibitor targets FOXO1 at the used concentration of 1 µM. Links to gene enrichment analysis carried out with *EnrichR*^71^ are embedded in the table, and data is available in File S1.

| Genes differentially regulated in | Number of genes | Jupyter file in File S1 (Table S3 subfolder) | Link |
| --- | --- | --- | --- |
| SW48 Parental | 4779 | 1 | EnrichR |
| SW48 G12D | 5221 | 2 | EnrichR |
| SW48 G12V | 5375 | 3 | EnrichR |
| All | 3058 | 4 | EnrichR |

**Figure S1.**
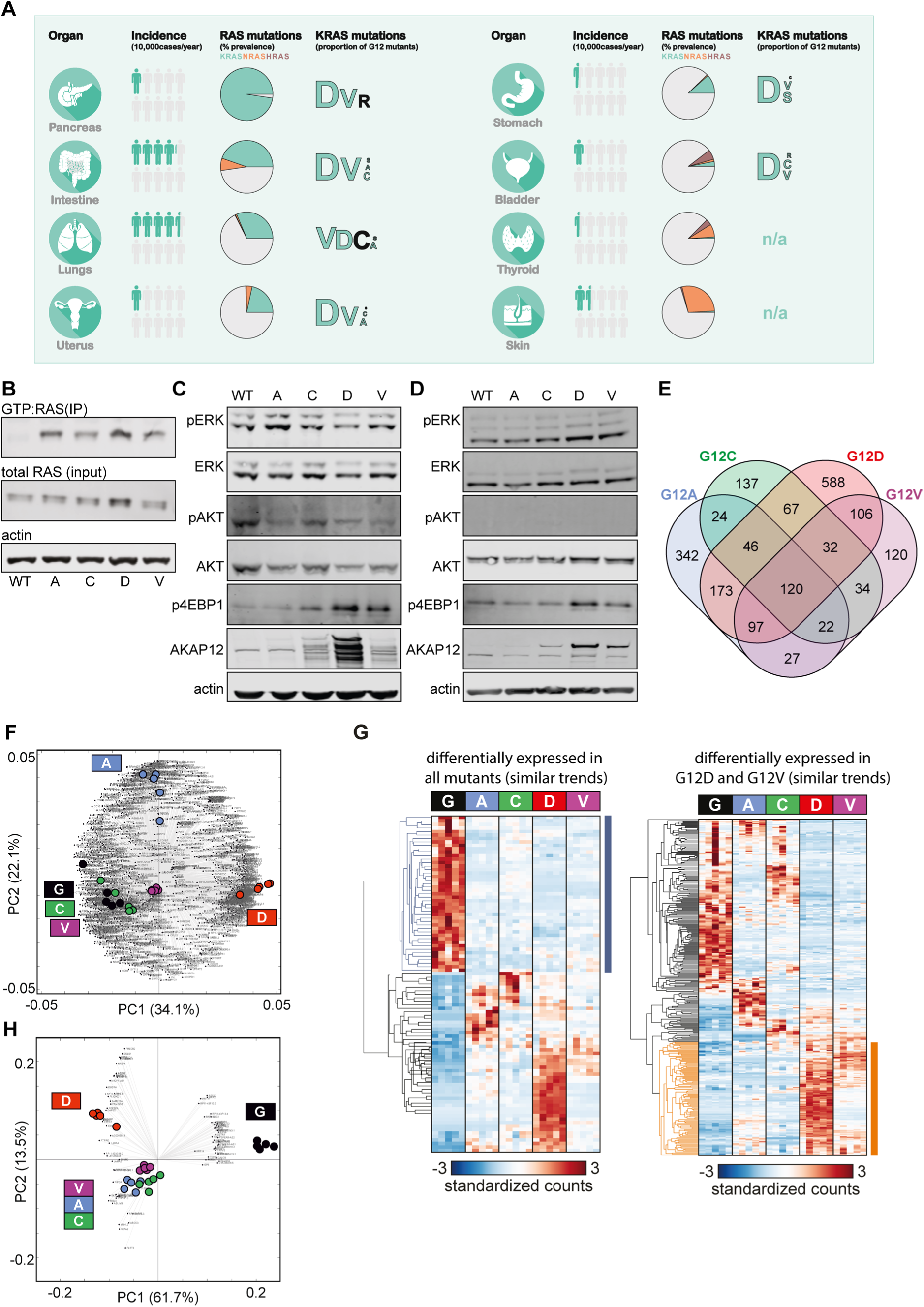
(related to Figure 1). Characterization of the SW48 isogenic panel. (A) Infographics summarizing tissue dependency of tumour incidence, prevalence of RAS mutations, and frequencies of glycine-12 codon missense mutations illustrated by the font size of the amino acid letter substituting glycine. Data from Cancer Research UK (2019) and Cox *et al.*^72^ (B) Immunoprecipitation of GTP-bound KRAS blotted tor RAS and actin. Lysates were prepared from SW48 cultured in full media. (C-D) Western blots show differences in downstream signalling amongst the different cell lines of the SW48 isogenic panel cultured in full media (10% FCS and 11 mM glucose) (C) or in low nutrient media (1% FCS and 2 mM glucose) conditions (D). (E) Venn diagram related to Figure 1A showing the number of genes that are differentially expressed in each mutant (n=5, FDR<5%, fold change > 2). For example, in these stringent statistical conditions, only 120 genes are differentially expressed in all mutants and G12D cells are those with the largest number (588) of genes to be uniquely regulated relative to WT. (F) Biplot related to Figure 1A. Panels show the results of PCA analysis with the gene loading in the background, and the sample scores coloured in the foreground. The code, data and vectorial files useful to inspect other genes are available as File S1 and in Table S1. (G) The same analysis reported in Figure 1A but all genes are constrained in exhibiting similar trends of up- or down-regulation relative to wild-type (left). All cell lines exhibit downregulation of common genes (blue cluster). The panel to the right shows a similar analysis but for those genes that are similarly up- or down-regulated in the G12D and G12V mutant lines. The orange cluster highlights genes that are particularly upregulated in G12D with a significant overlap in G12V. (H) Biplots related to the clustering analysis reported in panel G (left). The code, data and vectorial files useful to inspect other genes are available as File S1 and in Table S1.

**Figure S2.**
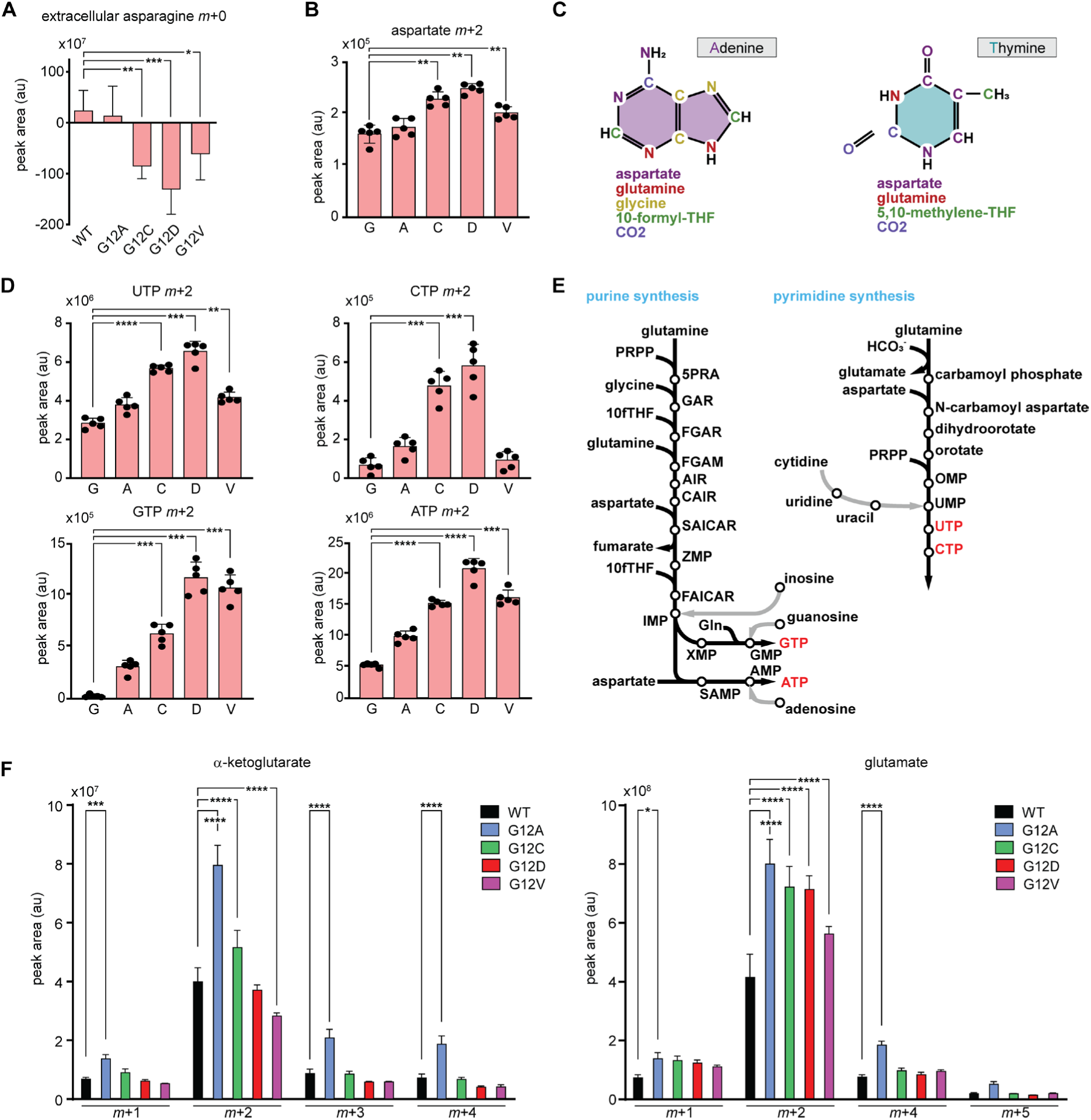
(related to Figure 2). Carbon flux from ^13^C-labelled glucose in SW48 cells. (A) Metabolic flux analysis using ^13^C-labelled glucose shows increased asparagine uptake in KRAS G12C, G12D and G12V mutants determined by consumption-release measurements. (B) Metabolic flux analysis using ^13^C-labelled glucose shows increased synthesis of aspartate (m+2) in KRAS G12C, G12D and G12V mutants. (C) Diagrammatic representations of adenine and thymine depicting the sources of each atom for purine and pyrimidine nucleotides. (D) Metabolic flux analysis using ^13^C-labelled glucose shows increased synthesis of ATP, CTP, GTP, UTP in KRAS G12C, G12D and G12V mutants. (E) Pathways showing the synthesis of purine (ATP, GTP) and pyrimidine (UTP, CTP) nucleotides. (F) Metabolic flux analysis using ^13^C-labelled glucose shows all detected isotopologues of α-ketoglutarate (left) and glutamate (right) in KRAS G12 mutants showing higher levels of all isotopologues in the G12A mutant cells. Adjusted p values are calculated using two-way ANOVA. Unless stated otherwise, data represents the mean of 5 technical repeats with standard deviations. Adjusted p values are calculated using one-way ANOVA, *p ≤ 0.05, \*\**p* ≤ 0.01, \*\*\**p* ≤ 0.001, \*\*\**p* ≤ 0.0001).

**Figure S3.**
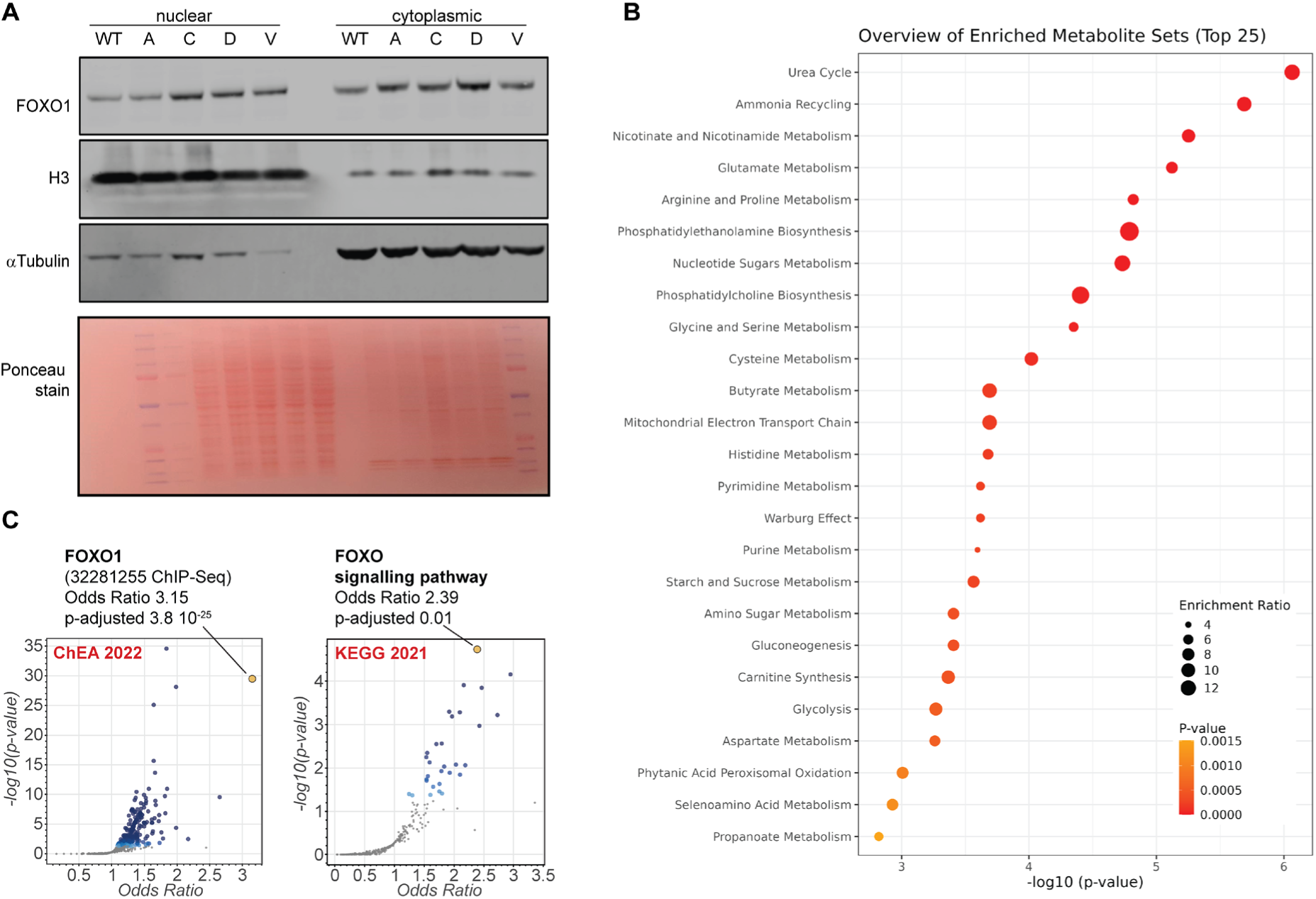
(related to Figure 3). FOXO1 nuclear and cytoplasmic fractions. (A) Representative immunoblots from nuclear-cytoplasmic fractionation experiments showing an increase in nuclear FOXO1 protein in KRAS mutants. (B) Enrichment analysis of metabolites changed upon FOXO1 inhibition in ^13^C-glucose carbon labelling experiment. Analysis was performed using Metaboanalyst 6.0^73^. (C) Volcano plots for all genes that are downregulated by FOXO1 inhibition (AS1842856 at 1 µM) in wild-type, G12D and G12V cell lines as measured by RNAseq. P-values and odd ratios for gene set enrichment was performed with EnrichR^71^ (see Table S3); here target genes of transcription factors (ChEA 2022) and KEGG pathways (KEGG 2021) are shown, both confirming that AS1842856 inhibits FOXO1 (see also File S1).

**Figure S4.**
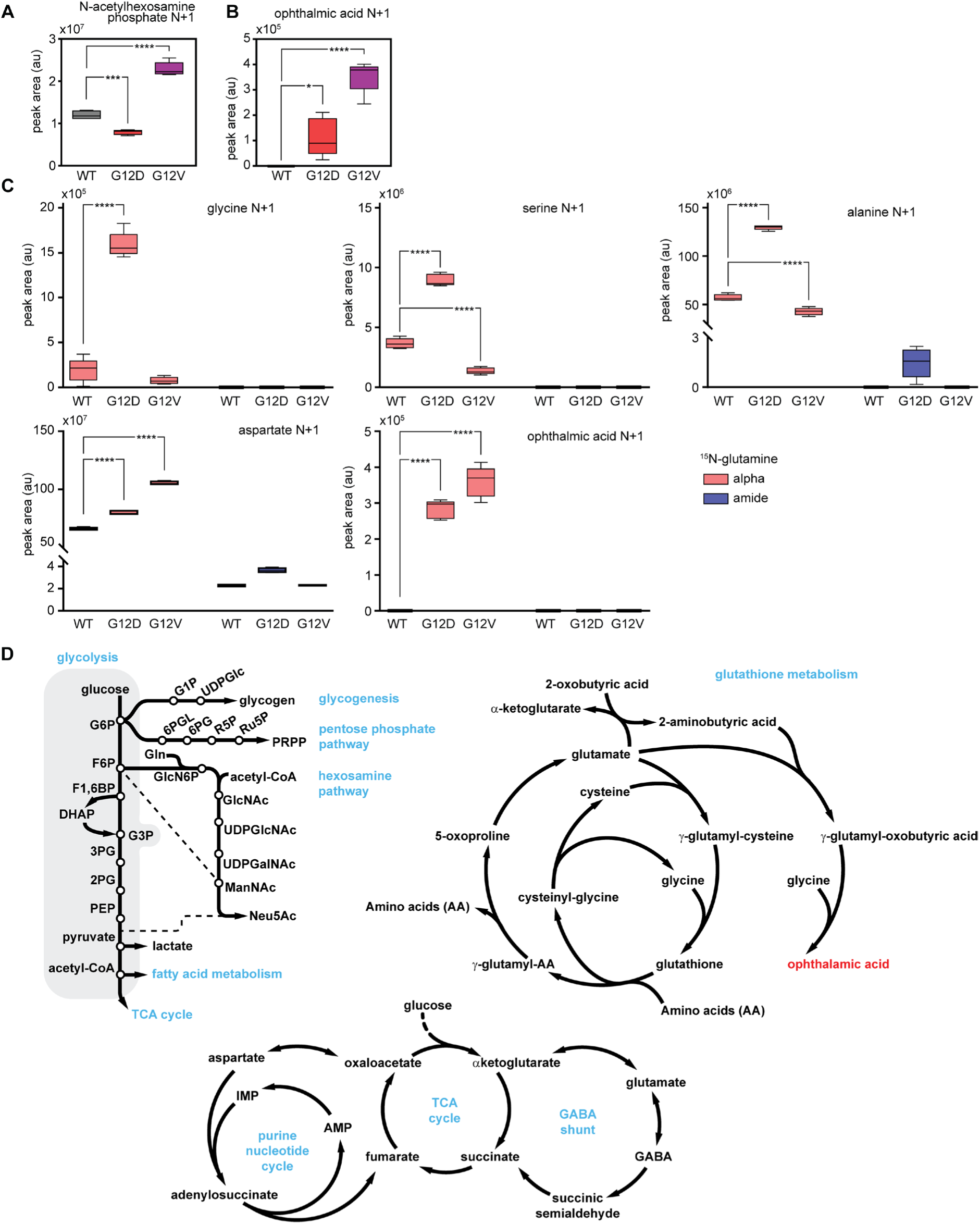
(related to Figure 4). Nitrogen metabolism in G12D and G12V SW48 mutant cells. (A) Nitrogen tracing using ^15^NH_3_ shows that the synthesis of N-acetylhexosamine, a metabolite within the hexosamine pathway (see also panel D), is significantly upregulated in the SW48 G12V mutant line. (B) Nitrogen tracing using ^15^NH_3_ shows that the synthesis of ophthalmic acid, a metabolite produced by the same enzymes that synthesize glutathion (see also panel D), is significantly upregulated in the mutant cell lines, SW48 G12V in particular. (C) Nitrogen tracing with ^15^N-amide-glutamine and ^15^N-alpha-glutamine. The nitrogen at the glutamine alpha-carbon is incorporated in the backbone of several amino acids through transamination reactions that are particularly upregulated in G12D cells. Higher integration of nitrogen from glutamate into aspartate and ophthalmic acid is observed in both G12D and G12V mutants. (D) Diagrammatic representations of various pathways discussed in the paper.

**Figure S5.**
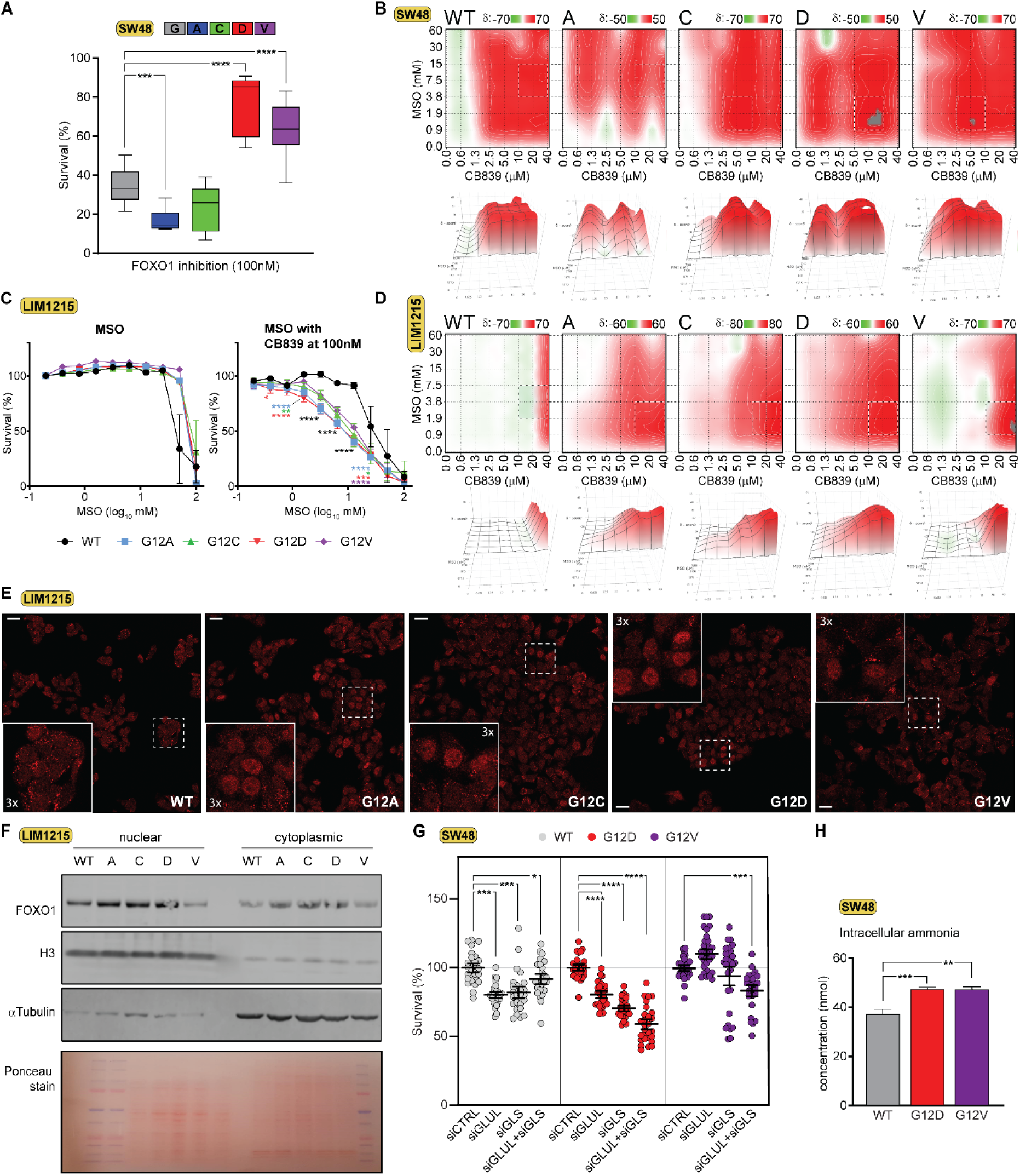
(related to Figure 5). Synergy analysis and validation. (A) G12D and G12V KRAS mutant cells exhibit resistance to inhibition of FOXO1 (AS1842856, 100 nM). Data represents the mean of 3 independent experiments and standard deviations. Adjusted p-values were computed by one-way ANOVA (***p ≤ 0.001, ****p ≤ 0.0001). (B) Synergistic effect of GS inhibitor, MSO, and GLS inhibitor, CB839, on KRAS mutant SW48 cells. BLISS analysis was performed with Synergy Finder^74^. (C) Viability curves showing titration of GS inhibitor, MSO, alone (left panel) or in combination with sub-lethal doses of GLS inhibitor, CB839, (right panel) in LIM1215 cells. The graphs show the means and standard deviations of 3 independent SRB assays. Adjusted p-values were computed by two-ways ANOVA (**p ≤ 0.05, **p ≤ 0.01,***p ≤ 0.001, ****p ≤ 0.0001). (D) Synergistic effect of GS inhibitor, MSO, and GLS inhibitor, CB839, on KRAS mutant LIM1215 cells. BLISS analysis was performed with Synergy Finder^74^. (E) Immunostaining for FOXO1 in LIM1215 cells. Scale bar: 25 µm; inserts: 3x magnification (F) Representative immunoblots for FOXO1 labelling from nuclear-cytoplasmic fractionation experiments in LIM1215 cells. (G) The combined knockdown of GLS and GLUL genes in SW48 cells sensitises G12D and G12V mutants. Data shows means with standard errors from 3 independent experiments. (H) Intracellular ammonia levels in SW48 cells were measured using an ammonia colourimetric assay. Graph shows means and standard deviations of 3 independent experiments. Adjusted p values are calculated using one-way ANOVA, (**p ≤ 0.01, ***p ≤ 0.001).

**Figure S6.**
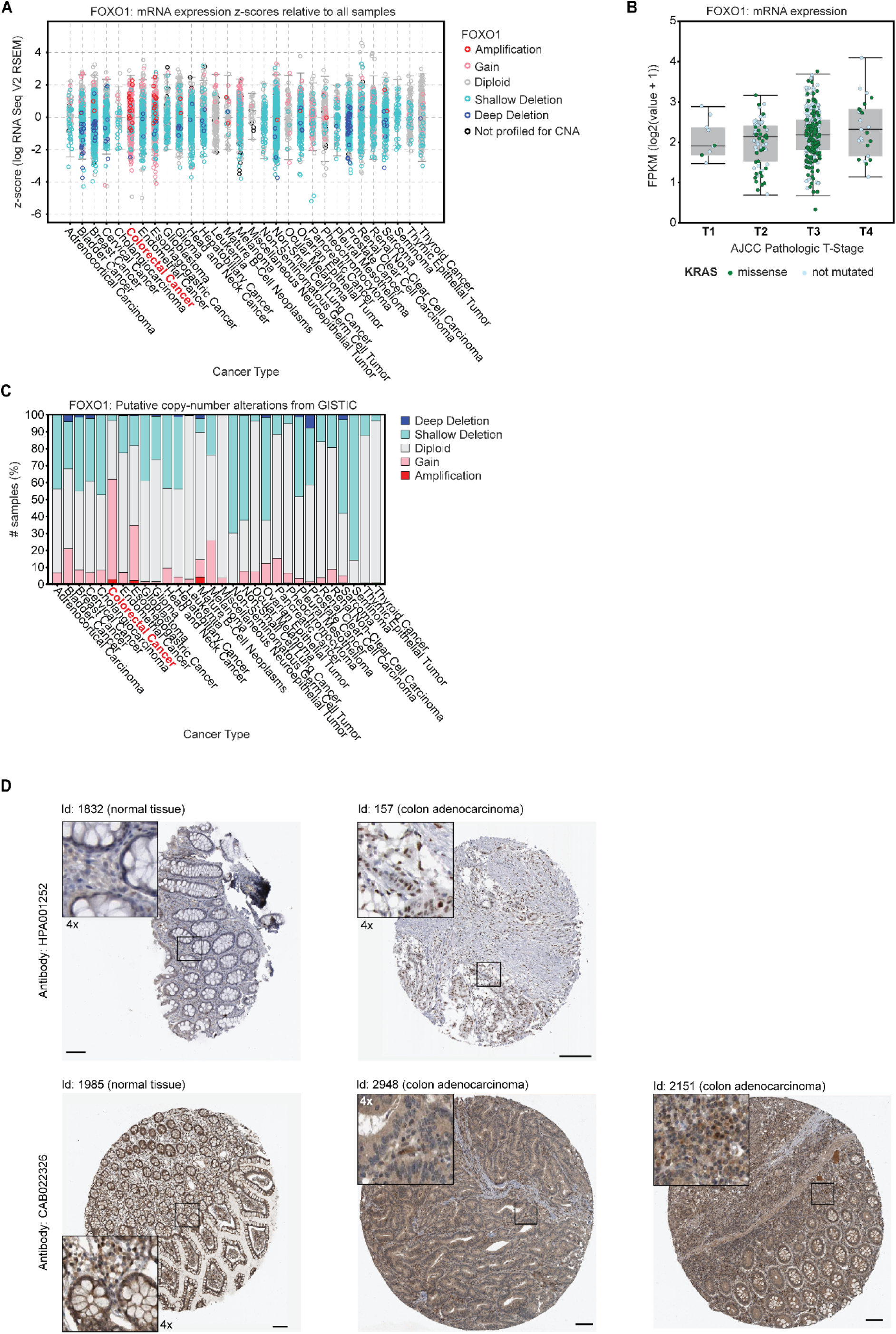
FOXO1 expression in cancer. (A) FOXO1 mRNA expression across different cancer types. The graph was generated with cBioPortal.org using the TCGA PanCancer Atlas Studies. FOXO1 mRNA does not seem to correlate with cancer type. The colour of each measurement indicates copy number variations in the FOXO1 gene. Gains and amplifications seems to be particularly prevalent in colorectal cancers. (B) FOXO1 mRNA expression in colorectal adenocarcinoma as a function of the American Joint Committee on Cancer (AJCC) tumour stage. The graph was generated with cBioPortal.org using the Colon Adenocarcinoma (TCGA, GDC) dataset. FOXO1 exhibits a mild upregulation during tumour progression. While exhibiting a strong correlation (R^2^=0.95) with differences that are statistically different (one-way ANOVA, p∼0.01), no statistical significance was identified between different groups, or KRAS mutational status. (C) The dataset shown in (A) was reanalysed for putative copy-number alterations showing that the colorectal cancers exhibit the highest frequency for FOXO1 gene gain and amlifications across all cancers included in the TCGA PanCancer Atlas Studies. (D) Immunohistopathology of colon sections comparing the staining of two different antibodies between normal tissue and colon adenocarcinoma. FOXO1 protein expression is very heterogeneous between patients and cells. Images were downloaded from the Human Protein Atlas (proteinatlas.org). The stainings were classified either as not detected, both cytoplasmic and nuclear, or nuclear.

## REFERENCES

1. Ostrow, S.L., Simon, E., Prinz, E., Bick, T., Shentzer, T., Nagawkar, S.S., Sabo, E., Ben-Izhak, O., Hershberg, R., and Hershkovitz, D. (2016). Variation in KRAS driver substitution distributions between tumor types is determined by both mutation and natural selection. Sci Rep 6, 21927. 10.1038/srep21927.

2. Cox, A.D., Fesik, S.W., Kimmelman, A.C., Luo, J., and Der, C.J. (2014). Drugging the undruggable RAS: Mission possible? Nat Rev Drug Discov 13, 828–851. 10.1038/nrd4389.

3. Haigis, K.M. (2017). KRAS Alleles: The Devil Is in the Detail. Trends Cancer 3, 686– 697. 10.1016/j.trecan.2017.08.006.

4. Cox, A.D., Fesik, S.W., Kimmelman, A.C., Luo, J., and Der, C.J. (2014). Drugging the undruggable RAS: Mission possible? Nat Rev Drug Discov 13, 828–851. 10.1038/nrd4389.

5. Haigis, K.M., Cichowski, K., and Elledge, S.J. (2019). Tissue-specificity in cancer: The rule, not the exception. Science (1979) 363, 1150–1151. 10.1126/science.aaw3472.

6. Li, S., Balmain, A., and Counter, C.M. (2018). A model for RAS mutation patterns in cancers: finding the sweet spot. Nat Rev Cancer 18, 767–777. 10.1038/s41568-018-0076-6.

7. Hunter, J.C., Manandhar, A., Carrasco, M.A., Gurbani, D., Gondi, S., and Westover, K.D. (2015). Biochemical and Structural Analysis of Common Cancer-Associated KRAS Mutations. Mol Cancer Res 13, 1325–1335. 10.1158/1541-7786.MCR-15-0203.

8. Ihle, N.T., Byers, L.A., Kim, E.S., Saintigny, P., Lee, J.J., Blumenschein, G.R., Tsao, A., Liu, S., Larsen, J.E., Wang, J., et al. (2012). Effect of KRAS oncogene substitutions on protein behavior: implications for signaling and clinical outcome. J Natl Cancer Inst 104, 228–239. 10.1093/jnci/djr523.

9. Munoz-Maldonado, C., Zimmer, Y., and Medova, M. (2019). A Comparative Analysis of Individual RAS Mutations in Cancer Biology. Front Oncol 9, 1088. 10.3389/fonc.2019.01088.

10. Yuan, T.L., Amzallag, A., Bagni, R., Yi, M., Afghani, S., Burgan, W., Fer, N., Strathern, L.A., Powell, K., Smith, B., et al. (2018). Differential Effector Engagement by Oncogenic KRAS. Cell Rep 22, 1889–1902. 10.1016/j.celrep.2018.01.051.

11. Son, J., Lyssiotis, C.A., Ying, H., Wang, X., Hua, S., Ligorio, M., Perera, R.M., Ferrone, C.R., Mullarky, E., Shyh-Chang, N., et al. (2013). Glutamine supports pancreatic cancer growth through a KRAS-regulated metabolic pathway. Nature 496, 101–105. 10.1038/nature12040.

12. Kim, J., Lee, H.M., Cai, F., Ko, B., Yang, C., Lieu, E.L., Muhammad, N., Rhyne, S., Li, K., Haloul, M., et al. (2020). The hexosamine biosynthesis pathway is a targetable liability in KRAS/LKB1 mutant lung cancer. Nat Metab 2, 1401–1412. 10.1038/s42255-020-00316-0.

13. Ying, H., Kimmelman, A.C., Lyssiotis, C.A., Hua, S., Chu, G.C., Fletcher-Sananikone, E., Locasale, J.W., Son, J., Zhang, H., Coloff, J.L., et al. (2012). Oncogenic Kras maintains pancreatic tumors through regulation of anabolic glucose metabolism. Cell 149, 656–670. 10.1016/j.cell.2012.01.058.

14. Kerr, E.M., Gaude, E., Turrell, F.K., Frezza, C., and Martins, C.P. (2016). Mutant Kras copy number defines metabolic reprogramming and therapeutic susceptibilities. Nature 531, 110–113. 10.1038/nature16967.

15. Mayers, J.R., Torrence, M.E., Danai, L. V, Papagiannakopoulos, T., Davidson, S.M., Bauer, M.R., Lau, A.N., Ji, B.W., Dixit, P.D., Hosios, A.M., et al. (2016). Tissue of origin dictates branched-chain amino acid metabolism in mutant Kras-driven cancers. Science 353, 1161–1165. 10.1126/science.aaf5171.

16. Gwinn, D.M., Lee, A.G., Briones-Martin-Del-Campo, M., Conn, C.S., Simpson, D.R., Scott, A.I., Le, A., Cowan, T.M., Ruggero, D., and Sweet-Cordero, E.A. (2018). Oncogenic KRAS Regulates Amino Acid Homeostasis and Asparagine Biosynthesis via ATF4 and Alters Sensitivity to L-Asparaginase. Cancer Cell 33, 91–107.e6. 10.1016/j.ccell.2017.12.003.

17. Kerk, S.A., Papagiannakopoulos, T., Shah, Y.M., and Lyssiotis, C.A. (2021). Metabolic networks in mutant KRAS-driven tumours: tissue specificities and the microenvironment. Nat Rev Cancer 21, 510–525. 10.1038/s41568-021-00375-9.

18. Zhao, Y., Yang, J., Liao, W., Liu, X., Zhang, H., Wang, S., Wang, D., Feng, J., Yu, L., and Zhu, W.G. (2010). Cytosolic FoxO1 is essential for the induction of autophagy and tumour suppressor activity. Nat Cell Biol 12, 665–675. 10.1038/ncb2069.

19. Kim, J.H., Kim, M.K., Lee, H.E., Cho, S.J., Cho, Y.J., Lee, B.L., Lee, H.S., Nam, S.Y., Lee, J.S., and Kim, W.H. (2007). Constitutive phosphorylation of the FOXO1A transcription factor as a prognostic variable in gastric cancer. Modern Pathology 20, 835–842. 10.1038/modpathol.3800789.

20. Paik, J.H., Kollipara, R., Chu, G., Ji, H., Xiao, Y., Ding, Z., Miao, L., Tothova, Z., Horner, J.W., Carrasco, D.R., et al. (2007). FoxOs Are Lineage-Restricted Redundant Tumor Suppressors and Regulate Endothelial Cell Homeostasis. Cell 128, 309–323. 10.1016/j.cell.2006.12.029.

21. Zhang, X., Tang, N., Hadden, T.J., and Rishi, A.K. (2011). Akt, FoxO and regulation of apoptosis. Preprint, 10.1016/j.bbamcr.2011.03.010 https://doi.org/10.1016/j.bbamcr.2011.03.010.

22. Xie, L., Ushmorov, A., Leithä, F., Guan, H., Steidl, C., Fä, J., Pelzer, C., Vogel, M.J., Maier, H.J., Gascoyne, R.D., et al. (2012). LYMPHOID NEOPLASIA FOXO1 is a tumor suppressor in classical Hodgkin lymphoma. 10.1182/blood.

23. Hammond, D.E., Mageean, C.J., Rusilowicz, E. V, Wickenden, J.A., Clague, M.J., and Prior, I.A. (2015). Differential reprogramming of isogenic colorectal cancer cells by distinct activating KRAS mutations. J Proteome Res 14, 1535–1546. 10.1021/pr501191a.

24. Sullivan, M.R., Danai, L. V, Lewis, C.A., Chan, S.H., Gui, D.Y., Kunchok, T., Dennstedt, E.A., Vander Heiden, M.G., and Muir, A. (2019). Quantification of microenvironmental metabolites in murine cancers reveals determinants of tumor nutrient availability. Elife 8. 10.7554/eLife.44235.

25. Cruzat, V., Macedo Rogero, M., Noel Keane, K., Curi, R., and Newsholme, P. (2018). Glutamine: Metabolism and Immune Function, Supplementation and Clinical Translation. Nutrients 10, 1564. 10.3390/nu10111564.

26. Van Der Vos, K.E., Eliasson, P., Proikas-Cezanne, T., Vervoort, S.J., Van Boxtel, R., Putker, M., Van Zutphen, I.J., Mauthe, M., Zellmer, S., Pals, C., et al. (2012). Modulation of glutamine metabolism by the PI(3)K-PKB-FOXO network regulates autophagy. Nat Cell Biol 14, 829–837. 10.1038/ncb2536.

27. Kamei, Y., Hattori, M., Hatazawa, Y., Kasahara, T., Kanou, M., Kanai, S., Yuan, X., Suganami, T., Lamers, W.H., Kitamura, T., et al. (2014). FOXO1 activates glutamine synthetase gene in mouse skeletal muscles through a region downstream of 3=-UTR: possible contribution to ammonia detoxification. 10.1152/ajpendo.00177.2014.-Skeletal.

28. Bruce Rowe, W., Ronzio, R.A., and Meister, A. (1965). Inhibition of Glutamine Synthetase by Methionine Sulfoximine. Studies on Methionine Sulfoximine Phosphate" (Meister, A).

29. Demarco, V., Dyess, K., Strauss, D., West, C.M., and Neu, J. (1999). Biochemical and Molecular Roles of Nutrients Inhibition of Glutamine Synthetase Decreases Proliferation of Cultured Rat Intestinal Epithelial Cells 1,2.

30. Ghoddoussi, F., Galloway, M.P., Jambekar, A., Bame, M., Needleman, R., and Brusilow, W.S.A. (2010). Methionine sulfoximine, an inhibitor of glutamine synthetase, lowers brain glutamine and glutamate in a mouse model of ALS. J Neurol Sci 290, 41–47. 10.1016/j.jns.2009.11.013.

31. Gross, M.I., Demo, S.D., Dennison, J.B., Chen, L., Chernov-Rogan, T., Goyal, B., Janes, J.R., Laidig, G.J., Lewis, E.R., Li, J., et al. (2014). Antitumor activity of the glutaminase inhibitor CB-839 in triple-negative breast cancer. Mol Cancer Ther 13, 890–901. 10.1158/1535-7163.MCT-13-0870.

32. Raczka, A.M., and Reynolds, P.A. (2019). Glutaminase inhibition in renal cell carcinoma therapy. Cancer Drug Resistance. 10.20517/cdr.2018.004.

33. Zheng, S., Wang, W., Aldahdooh, J., Malyutina, A., Shadbahr, T., Tanoli, Z., Pessia, A., and Tang, J. (2022). SynergyFinder Plus: Toward Better Interpretation and Annotation of Drug Combination Screening Datasets. Genomics Proteomics Bioinformatics 20, 587–596. 10.1016/j.gpb.2022.01.004.

34. Varshavi, D., Varshavi, D., McCarthy, N., Veselkov, K., Keun, H.C., and Everett, J.R. (2020). Metabolic characterization of colorectal cancer cells harbouring different KRAS mutations in codon 12, 13, 61 and 146 using human SW48 isogenic cell lines. Metabolomics 16. 10.1007/s11306-020-01674-2.

35. Hammond, D.E., Mageean, C.J., Rusilowicz, E. V, Wickenden, J.A., Clague, M.J., and Prior, I.A. (2015). Differential reprogramming of isogenic colorectal cancer cells by distinct activating KRAS mutations. J Proteome Res 14, 1535–1546. 10.1021/pr501191a.

36. Gui, D.Y., Sullivan, L.B., Luengo, A., Hosios, A.M., Bush, L.N., Gitego, N., Davidson, S.M., Freinkman, E., Thomas, C.J., and Vander Heiden, M.G. (2016). Environment Dictates Dependence on Mitochondrial Complex I for NAD+ and Aspartate Production and Determines Cancer Cell Sensitivity to Metformin. Cell Metab 24, 716–727. 10.1016/j.cmet.2016.09.006.

37. Muir, A., and Vander Heiden, M.G. (2018). The nutrient environment affects therapy Nutrient availability affects cancer cell metabolism and therapeutic responses. Science (1979) 360, 962–963. 10.1126/science.aar5986.

38. Muir, A., Danai, L. V, Gui, D.Y., Waingarten, C.Y., Lewis, C.A., and Vander Heiden, M.G. (2017). Environmental cystine drives glutamine anaplerosis and sensitizes cancer cells to glutaminase inhibition. Elife 6. 10.7554/eLife.27713.

39. Nightingale, A.M., Leong, C.L., Burnish, R.A., Hassan, S. ul, Zhang, Y., Clough, G.F., Boutelle, M.G., Voegeli, D., and Niu, X. (2019). Monitoring biomolecule concentrations in tissue using a wearable droplet microfluidic-based sensor. Nat Commun 10. 10.1038/s41467-019-10401-y.

40. Feng, X., Wu, Z., Wu, Y., Hankey, W., Prior, T.W., Li, L., Ganju, R.K., Shen, R., and Zou, X. (2011). Cdc25A Regulates Matrix Metalloprotease 1 through Foxo1 and Mediates Metastasis of Breast Cancer Cells. Mol Cell Biol 31, 3457–3471. 10.1128/mcb.05523-11.

41. Trinh, D.L., Scott, D.W., Morin, R.D., Mendez-Lago, M., An, J., Jones, S.J.M., Mungall, A.J., Zhao, Y., Schein, J., Steidl, C., et al. (2013). Analysis of FOXO1 mutations in diffuse large B-cell lymphoma. Blood 121, 3666–3674. 10.1182/blood-2013.

42. Hornsveld, M., Dansen, T.B., Derksen, P.W., and Burgering, B.M.T. (2018). Re-evaluating the role of FOXOs in cancer. Preprint at Academic Press, 10.1016/j.semcancer.2017.11.017 https://doi.org/10.1016/j.semcancer.2017.11.017.

43. Greer, E.L., and Brunet, A. (2005). FOXO transcription factors at the interface between longevity and tumor suppression. Preprint, 10.1038/sj.onc.1209086 https://doi.org/10.1038/sj.onc.1209086.

44. Essers, M.A.G., Weijzen, S., De Vries-Smits, A.M.M., Saarloos, I., De Ruiter, N.D., Bos, J.L., and Burgering, B.M.T. (2004). FOXO transcription factor activation by oxidative stress mediated by the small GTPase Ral and JNK. EMBO Journal 23, 4802–4812. 10.1038/sj.emboj.7600476.

45. Brunet, A., Sweeney, L.B., Sturgill, J.F., Chua, K.F., Greer, P.L., Lin, Y., Tran, H., Ross, S.E., Mostoslavsky, R., Cohen, H.Y., et al. (2004). Stress-Dependent Regulation of FOXO Transcription Factors by the SIRT1 Deacetylase. Science (1979) 303, 2011–2015. 10.1126/science.1094637.

46. Van Der Heide, L.P., Hoekman, M.F.M., and Smidt, M.P. (2004). The ins and outs of FoxO shuttling: mechanisms of FoxO translocation and transcriptional regulation.

47. Cheng, Z. (2022). FoxO transcription factors in mitochondrial homeostasis. Preprint at Portland Press Ltd, 10.1042/BCJ20210777 https://doi.org/10.1042/BCJ20210777.

48. Jiang, M., Xu, B., Li, X., Shang, Y., Chu, Y., Wang, W., Chen, D., Wu, N., Hu, S., Zhang, S., et al. (2019). O-GlcNAcylation promotes colorectal cancer metastasis via the miR-101-O-GlcNAc/EZH2 regulatory feedback circuit. Oncogene 38, 301–316. 10.1038/s41388-018-0435-5.

49. Paneque, A., Fortus, H., Zheng, J., Werlen, G., and Jacinto, E. (2023). The Hexosamine Biosynthesis Pathway: Regulation and Function. Genes (Basel) 14, 933. 10.3390/genes14040933.

50. Kim, J., Hu, Z., Cai, L., Li, K., Choi, E., Faubert, B., Bezwada, D., Rodriguez-Canales, J., Villalobos, P., Lin, Y.F., et al. (2017). CPS1 maintains pyrimidine pools and DNA synthesis in KRAS/LKB1-mutant lung cancer cells. Nature 546, 168–172. 10.1038/nature22359.

51. Spinelli, J.B., Yoon, H., Ringel, A.E., Jeanfavre, S., Clish, C.B., and Haigis, M.C. (2017). Metabolic recycling of ammonia via glutamate dehydrogenase supports breast cancer biomass. Science (1979) 358, 941–946. 10.1126/science.aam9305.

52. Lie, S., Wang, T., Forbes, B., Proud, C.G., and Petersen, J. (2019). The ability to utilise ammonia as nitrogen source is cell type specific and intricately linked to GDH, AMPK and mTORC1. Sci Rep 9. 10.1038/s41598-018-37509-3.

53. Cheng, C., Geng, F., Li, Z., Zhong, Y., Wang, H., Cheng, X., Zhao, Y., Mo, X., Horbinski, C., Duan, W., et al. (2022). Ammonia stimulates SCAP/Insig dissociation and SREBP-1 activation to promote lipogenesis and tumour growth. Nat Metab 4, 575–588. 10.1038/s42255-022-00568-y.

54. Karkoutly, S., Takeuchi, Y., Mehrazad Saber, Z., Ye, C., Tao, D., Aita, Y., Murayama, Y., Shikama, A., Masuda, Y., Izumida, Y., et al. (2024). FoxO transcription factors regulate urea cycle through Ass1. Biochem Biophys Res Commun 739. 10.1016/j.bbrc.2024.150594.

55. Guinney, J., Dienstmann, R., Wang, X., de Reyniès, A., Schlicker, A., Soneson, C., Marisa, L., Roepman, P., Nyamundanda, G., Angelino, P., et al. (2015). The consensus molecular subtypes of colorectal cancer. Nat Med 21, 1350–1356. 10.1038/nm.3967.

56. Li, X., Zhu, H., Sun, W., Yang, X., Nie, Q., and Fang, X. (2021). Role of glutamine and its metabolite ammonia in crosstalk of cancer-associated fibroblasts and cancer cells. Preprint at BioMed Central Ltd, 10.1186/s12935-021-02121-5 https://doi.org/10.1186/s12935-021-02121-5.

57. Mestre-Farrera, A., Bruch-Oms, M., Peña, R., Rodríguez-Morato, J., Alba-Castellon, L., Comerma, L., Quintela-Fandino, M., Duñach, M., Baulida, J., Pozo, O.J., et al. (2021). Glutamine-Directed Migration of Cancer-Activated Fibroblasts Facilitates Epithelial Tumor Invasion. Cancer Res 81, 438–451. 10.1158/0008-5472.CAN-20-0622.

58. Yang, L., Achreja, A., Yeung, T.L., Mangala, L.S., Jiang, D., Han, C., Baddour, J., Marini, J.C., Ni, J., Nakahara, R., et al. (2016). Targeting Stromal Glutamine Synthetase in Tumors Disrupts Tumor Microenvironment-Regulated Cancer Cell Growth. Cell Metab 24, 685–700. 10.1016/j.cmet.2016.10.011.

59. Ozcan, L., Wong, C.C.L., Li, G., Xu, T., Pajvani, U., Park, S.K.R., Wronska, A., Chen, B.-X., Marks, A.R., Fukamizu, A., et al. (2012). Calcium Signaling through CaMKII Regulates Hepatic Glucose Production in Fasting and Obesity. Cell Metab 15, 739– 751. 10.1016/j.cmet.2012.03.002.

60. Wu, Y., Pan, Q., Yan, H., Zhang, K., Guo, X., Xu, Z., Yang, W., Qi, Y., Guo, C.A., Hornsby, C., et al. (2018). Novel Mechanism of Foxo1 Phosphorylation in Glucagon Signaling in Control of Glucose Homeostasis. Diabetes 67, 2167–2182. 10.2337/db18-0674.

61. Gaglio, D., Metallo, C.M., Gameiro, P.A., Hiller, K., Danna, L.S., Balestrieri, C., Alberghina, L., Stephanopoulos, G., and Chiaradonna, F. (2011). Oncogenic K-Ras decouples glucose and glutamine metabolism to support cancer cell growth. Mol Syst Biol 7. 10.1038/msb.2011.56.

62. Ko, Y.-H., Lin, Z., Flomenberg, N., Pestell, R.G., Howell, A., Sotgia, F., Lisanti, M.P., and Martinez-Outschoorn, U.E. (2011). Glutamine fuels a vicious cycle of autophagy in the tumor stroma and oxidative mitochondrial metabolism in epithelial cancer cells. Cancer Biol Ther 12, 1085–1097. 10.4161/cbt.12.12.18671.

63. Bott, A.J., Shen, J., Tonelli, C., Zhan, L., Sivaram, N., Jiang, Y.P., Yu, X., Bhatt, V., Chiles, E., Zhong, H., et al. (2019). Glutamine Anabolism Plays a Critical Role in Pancreatic Cancer by Coupling Carbon and Nitrogen Metabolism. Cell Rep 29, 1287–1298.e6. 10.1016/j.celrep.2019.09.056.

64. Encarnación-Rosado, J., Sohn, A.S.W., Biancur, D.E., Lin, E.Y., Osorio-Vasquez, V., Rodrick, T., González-Baerga, D., Zhao, E., Yokoyama, Y., Simeone, D.M., et al. (2024). Targeting pancreatic cancer metabolic dependencies through glutamine antagonism. Nat Cancer 5, 85–99. 10.1038/s43018-023-00647-3.

65. Biancur, D.E., Paulo, J.A., Małachowska, B., Del Rey, M.Q., Sousa, C.M., Wang, X., Sohn, A.S.W., Chu, G.C., Gygi, S.P., Harper, J.W., et al. (2017). Compensatory metabolic networks in pancreatic cancers upon perturbation of glutamine metabolism. Nat Commun 8. 10.1038/ncomms15965.

66. Lin, Y., Richards, F.M., Krippendorff, B.-F., Bramhall, J.L., Harrington, J.A., Bapiro, T.E., Robertson, A., Zheleva, D., and Jodrell, D.I. (2012). Paclitaxel and CYC3, an aurora kinase A inhibitor, synergise in pancreatic cancer cells but not bone marrow precursor cells. Br J Cancer 107, 1692–1701. 10.1038/bjc.2012.450.

67. Dobin, A., Davis, C.A., Schlesinger, F., Drenkow, J., Zaleski, C., Jha, S., Batut, P., Chaisson, M., and Gingeras, T.R. (2013). STAR: ultrafast universal RNA-seq aligner. Bioinformatics 29, 15–21. 10.1093/bioinformatics/bts635.

68. Liao, Y., Smyth, G.K., and Shi, W. (2019). The R package Rsubread is easier, faster, cheaper and better for alignment and quantification of RNA sequencing reads. Nucleic Acids Res 47, e47–e47. 10.1093/nar/gkz114.

69. Love, M.I., Huber, W., and Anders, S. (2014). Moderated estimation of fold change and dispersion for RNA-seq data with DESeq2. Genome Biol 15, 550. 10.1186/s13059-014-0550-8.

70. Su, X., Lu, W., and Rabinowitz, J.D. (2017). Metabolite Spectral Accuracy on Orbitraps. Anal Chem 89, 5940–5948. 10.1021/acs.analchem.7b00396.

71. Chen, E.Y., Tan, C.M., Kou, Y., Duan, Q., Wang, Z., Meirelles, G.V., Clark, N.R., and Ma’ayan, A. (2013). Enrichr: interactive and collaborative HTML5 gene list enrichment analysis tool. BMC Bioinformatics 14, 128. 10.1186/1471-2105-14-128.

72. Cox, A.D., Fesik, S.W., Kimmelman, A.C., Luo, J., and Der, C.J. (2014). Drugging the undruggable RAS: Mission possible? Nat Rev Drug Discov 13, 828–851. 10.1038/nrd4389.

73. Pang, Z., Lu, Y., Zhou, G., Hui, F., Xu, L., Viau, C., Spigelman, A.F., MacDonald, P.E., Wishart, D.S., Li, S., et al. (2024). MetaboAnalyst 6.0: towards a unified platform for metabolomics data processing, analysis and interpretation. Nucleic Acids Res 52, W398–W406. 10.1093/nar/gkae253.

74. Zheng, S., Wang, W., Aldahdooh, J., Malyutina, A., Shadbahr, T., Tanoli, Z., Pessia, A., and Tang, J. (2022). SynergyFinder Plus: Toward Better Interpretation and Annotation of Drug Combination Screening Datasets. Genomics Proteomics Bioinformatics 20, 587–596. 10.1016/j.gpb.2022.01.004.

