## Supplemental File S1 for "KRAS G12 mutant alleles differentially control glutamine metabolism via FOXO1": fig_process_2024.m

```
% option structure must contain the following fields:
%
% trend:   '+': upregulated
%          '-': downregulated
%          '' : differentially regulated, 
%          '=': differentially regulated but similarly across mutants (all + or all -)
%
% fdr:              false discovery rate threshold
% hits:             in how many mutants the gene should be differentially expressed?
% mutants:          index of which mutant to test
% foldchange:       threhold on log2 foldchange (|log2| > threshold)
% name:             name of the folder that will be created to store the data
% kegg:             KEGG pathway to search geneset from (e.g. hsa00250)
% highlightgenes:   gene list of genes that should be hilighted in vulcano plots and biplots
% destroy:          true/false - close all figures when finished
% stats:            filter genes that are statistically significant
% filetypes        output filetypes (e.g. emf, pdf, ...)

function fig_process_2024(cnt, qv, fc, lbl, gene_names, opts)

    % Init clustergram
    stat_dist = 'euclidean';
    stat_link = 'average';

    slbl = {'G12A','G12C','G12D','G12V'};
    
    highlight_idx = [];
    cDFE_SW = (1:4);
    gn = numel(gene_names);

    if ~isempty(opts.kegg)
        kegg_genes = kegg_api(opts.kegg);  
    else
        kegg_genes = '';
    end

    if ~isempty(opts.highlightgenes)
        kegg_genes = opts.highlightgenes;
        pathway_name = 'not KEGG';
    end

    if isempty(opts.lut)
        LUT = colormap(hot);
    else
        LUT = opts.lut;
    end

    smp_idx = (1:25);
    dfe_idx = cDFE_SW(opts.mutants);
    
    if opts.stats % filter genes
        
        % Statistical filtering of genes 
    
        gen_up = (qv(dfe_idx,:)<=opts.fdr) & (fc(dfe_idx,:)>= opts.foldchange);  
        gen_dw = (qv(dfe_idx,:)<=opts.fdr) & (fc(dfe_idx,:)<=-opts.foldchange);  
        gen_de = gen_up | gen_dw;
    
        switch opts.trend
            case '+'
                gen_idx = find( sum(gen_up,1)>=opts.hits );
            case '-'
                gen_idx = find( sum(gen_dw,1)>=opts.hits );
            case '='
                gen_idx = unique([find( sum(gen_up,1)>=opts.hits) find( sum(gen_dw,1)>=opts.hits )]);
            otherwise 
                gen_idx = find( sum(gen_de,1)>=opts.hits );
        end     
        
    else % use gene list and plot all
        gen_idx = find(ismember(gene_names,kegg_genes));
    end
        
    
    if ~isempty(kegg_genes) & opts.stats
        highlight_idx = find(ismember(gene_names(gen_idx), kegg_genes));
    end
    
    %% clustering of counts
    
    cgo = clustergram( cnt(smp_idx,gen_idx)',...
                      'Standardize',2,...                   % 3: no, 2: genes, 1: mutants
                      'cluster',1,...                       % 1: genes, 2: samples, 3: both
                      'colormap',LUT,...            
                      'displayrange',3,...
                      'rowpdist',stat_dist,...
                      'columnpdist',stat_dist,...
                      'linkage',stat_link,...
                      'RowLabels', gene_names(gen_idx),...
                      'ColumnLabels',lbl(smp_idx));
    h_cgo  = plot(cgo);
    hf_cgo = gcf;
    set(hf_cgo,'WindowState','maximized','renderer','painters');  
    colorbar(h_cgo);
    
    
    
    %% clustering fold changes
    
    cgo2 = clustergram( fc([1 2 3 4],gen_idx)',...
                      'Standardize',3,...                   % 3: no, 2: genes, 1: mutants
                      'cluster',1,...                       % 1: genes, 2: samples, 3: both
                      'colormap',LUT,...            
                      'displayrange',2,...
                      'rowpdist',stat_dist,...
                      'columnpdist',stat_dist,...
                      'linkage',stat_link,...
                      'RowLabels', gene_names(gen_idx),...
                      'ColumnLabels',slbl);
   
    h_cgo2  = plot(cgo2);
    hf_cgo2 = gcf;
    set(hf_cgo2,'WindowState','maximized','renderer','painters');  
    colorbar(h_cgo2);
     
    %% PCA analysis of counts
    hf_pca = rnaseq_pca(cnt(smp_idx,gen_idx), gene_names(gen_idx), highlight_idx);
    
    %% vulcano plot
    hf_vul = figure('WindowState','maximized','renderer','painters');
    dfe_idx = (1:4);
    
    for si=1:4
        subplot(1,4,si)
        
        % plot markers of every gene
        hs=scatter(fc(dfe_idx(si),:),-log10(qv(dfe_idx(si),:)),'.');   
        % gene markers are gray by default
        scol = repmat([.8 .8 .8],[gn 1]);
        % or red if statistically significant
        scol(gen_idx,:)= repmat([1 0 0],[numel(gen_idx) 1]);
        set(hs,'CData',scol);
        
        % plot labels only for the statistically significant genes. 
        ht = text(fc(dfe_idx(si),gen_idx)+.1,-log10(qv(dfe_idx(si),gen_idx))+2,gene_names(gen_idx),'fontsize',6);
        % gene labels are black by default 
        % but if genes need to be highlighted, those not highlighted turn grey
    
        if ~isempty(kegg_genes)
            %kidx = find(ismember(gene_names(gen_idx),kegg_genes));
            if ~isempty(highlight_idx)
                set(ht,'color',[.8 .8 .8],'fontsize',6);
                set(ht(highlight_idx),'color',[1 0 0 ],'fontsize',8,'fontweight','bold');
                scol(gen_idx,:)= repmat([0 0 0],[numel(gen_idx) 1]);
            %    scol(kidx,:)= repmat([1 0 0],[numel(gen_idx) 1]);
                set(hs,'CData',scol);
                uistack(hs,'top')
                uistack(ht(highlight_idx),'top')
     
            else
               % title(replace([folder_name ' (no kegg gene found from ' pathway_name ')'],'_',' '))
            end
        end
        
        % plot axes and guidelines
        
        box on
        line(get(gca,'xlim'),[-log10(opts.fdr) -log10(opts.fdr)])
        line( [opts.foldchange opts.foldchange],get(gca,'ylim'))
        line(-[opts.foldchange opts.foldchange],get(gca,'ylim'))
        xlabel('fold change (log2)')
        ylabel('fdr (-log10)')
        title(slbl{si})
    end
    
    %% plot individual gene expression (counts) 
    
    if ~isempty(opts.kegg)
        hf_kegg = figure('name',opts.kegg,'WindowState','maximize','renderer','painters');
        ggn = numel(highlight_idx);
        for ggi=1:ggn
            subplot(round(sqrt(ggn)),ceil(sqrt(ggn)),ggi)
            boxplot(cnt(smp_idx,gen_idx(highlight_idx(ggi)))',reshape(repmat([1 2 3 4 5]',[1 5])',1,[]),'plotstyle','compact','Labels',{'G','A','C','D','V'},'colors','kbgrm')
            title(gene_names{gen_idx(highlight_idx(ggi))})
            ylabel('counts')
            axis square
        end
    end
    %% save output
    mkdir(opts.name)
    writecell(gene_names(gen_idx),[opts.name '/gene_list.txt']); 
    for fi=1:numel(opts.filetypes)
        saveas(hf_vul,[opts.name '/vulcano'],opts.filetypes{fi});
        saveas(hf_pca,[opts.name '/pca_biplot'],opts.filetypes{fi});
        saveas(h_cgo.Parent,[opts.name '/clustergram'],opts.filetypes{fi});
        saveas(h_cgo2.Parent,[opts.name '/clustergramDE'],opts.filetypes{fi});
        if ~isempty(kegg_genes)
            saveas(hf_kegg,[opts.name '/kegg' opts.kegg],opts.filetypes{fi});
        end
    end
    
    display(['Saved info about ' num2str(numel(gen_idx)) ' genes'])
    
    if opts.destroy
        delete(cgo)
        delete(cgo2)
        close all
    end
    
    beep
```
