## Supplemental File S1 for "KRAS G12 mutant alleles differentially control glutamine metabolism via FOXO1": generate_data_for_figures_2024.m

```
% Initialize

load('rnaseqdb2024.mat')                    % Load dataset

%% Plot GLUL RNAseq expression (Fig. 2E)
plot_gene_expression('GLUL',gene_names, fc, qv, se, cnt);

%% Hierarcical clustering of genes that are different across all mutants - example code
exp_def = struct('name','test', ...
                 'trend','', ...
                 'fdr',0.1, ...
                 'foldchange',.5, ...
                 'mutants',[cMUT_A cMUT_C cMUT_D cMUT_V], ...
                 'hits',1,'kegg','', ...
                 'highlightgenes','', ...                
                 'destroy',0, ...                
                 'stats',1, ...
                 'filetypes','emf', ...
                 'lut',LUT);                       

fig_process_2024(cnt, qv, fc, lbl, gene_names, exp_def)


return

%% Hierarcical clustering of genes that are different across all mutants - example code
exp_def = struct('name','hsa00250_nofilter', ...
                 'trend','', ...
                 'fdr',0.1, ...
                 'foldchange',.5, ...
                 'mutants',[cMUT_A cMUT_C cMUT_D cMUT_V], ...
                 'hits',1,...
                 'kegg','hsa00250', ...
                 'highlightgenes','', ...                
                 'destroy',0, ...                
                 'stats',0, ...
                 'filetypes',{{'emf'}});                       

fig_process_2024(cnt, qv, fc, lbl, gene_names, exp_def)


%%  Common parameters to Fig. 1A-D
bDestroy    = true;                         % Set to true if the script is run at once. Figures will be closed but data will be available from harddrive
fChange     = 1;                            % Fold change threshold log2(1.5) or 1
fDr         = 0.05;                         % FDR threshold
sPrefix     = '2024_fdr005_fc1';            % Prefix to name to identify run
file_types  = {{'pdf','png','fig','emf'}};    % file format Matlab will save figure files to
bStats      = 1;                            %

%% Gene differentially expressed in at least one mutant (Fig.1A)
exp_def = struct('name',[ sPrefix 'atleastone_fig1A'],'trend','','fdr',fDr,'foldchange',fChange,'mutants',[cMUT_A cMUT_C cMUT_D cMUT_V],'hits',1,'kegg','','highlightgenes','','destroy',bDestroy,'stats',bStats,'filetypes',file_types);
fig_process_2024(cnt, qv, fc, lbl, gene_names, exp_def);

%% Gene differentially expressed in all mutants with similar (up/down) trends (Fig.1B)
exp_def = struct('name',[ sPrefix 'all_diff_equal_fig1B'],'trend','=','fdr',fDr,'foldchange',fChange,'mutants',[cMUT_A cMUT_C cMUT_D cMUT_V],'hits',4,'kegg','','highlightgenes','','destroy',bDestroy,'stats',bStats,'filetypes',file_types);
fig_process_2024(cnt, qv, fc, lbl, gene_names, exp_def);

%% Gene differentially expressed in G12V and G12D with similar (up/down) trends (Fig.1C)
exp_def = struct('name',[ sPrefix 'VD_diff_equal_fig1C'],'trend','=','fdr',fDr,'foldchange',fChange,'mutants',[cMUT_D cMUT_V],'hits',2,'kegg','','highlightgenes','','destroy',bDestroy,'stats',bStats,'filetypes',file_types);
fig_process_2024(cnt, qv, fc, lbl, gene_names, exp_def);

%% Gene differentially expressed in G12A (supp table 1)
exp_def = struct('name',[ sPrefix 'topgenes_A'],'trend','','fdr',fDr,'foldchange',fChange,'mutants',[cMUT_A],'hits',1,'kegg','','highlightgenes','','destroy',bDestroy,'stats',bStats,'filetypes',file_types);
fig_process_2024(cnt, qv, fc, lbl, gene_names, exp_def);

%% Gene differentially expressed in G12C (supp table 1)
exp_def = struct('name',[ sPrefix 'topgenes_C'],'trend','','fdr',fDr,'foldchange',fChange,'mutants',[cMUT_C],'hits',1,'kegg','','highlightgenes','','destroy',bDestroy,'stats',bStats,'filetypes',file_types);
fig_process_2024(cnt, qv, fc, lbl, gene_names, exp_def);

%% Gene differentially expressed in G12D (supp table 1)
exp_def = struct('name',[ sPrefix 'topgenes_D'],'trend','','fdr',fDr,'foldchange',fChange,'mutants',[cMUT_D],'hits',1,'kegg','','highlightgenes','','destroy',bDestroy,'stats',bStats,'filetypes',file_types);
fig_process_2024(cnt, qv, fc, lbl, gene_names, exp_def);

%% Gene differentially expressed in G12V (supp table 1) 
exp_def = struct('name',[ sPrefix 'topgenes_V'],'trend','','fdr',fDr,'foldchange',fChange,'mutants',[cMUT_V],'hits',1,'kegg','','highlightgenes','','destroy',bDestroy,'stats',bStats,'filetypes',file_types);
fig_process_2024(cnt, qv, fc, lbl, gene_names, exp_def);

%% Gene differentially expressed in G12V and G12D up trends (orange cluster in supp table 1 and Fig. 1)
exp_def = struct('name',[ sPrefix 'VD_UP_orangecluster'],'trend','+','fdr',fDr,'foldchange',fChange,'mutants',[cMUT_D cMUT_V],'hits',2,'kegg','','highlightgenes','','destroy',bDestroy,'stats',bStats,'filetypes',file_types);
fig_process_2024(cnt, qv, fc, lbl, gene_names, exp_def);

%%  GLS clustering
ggg = {'GLS';'FOXO1';'GLUL';'GAD1'};
exp_def = struct('name','GLS','trend','','fdr',1,'foldchange',.1,'mutants',[cMUT_D cMUT_V],'hits',2,'kegg','','highlightgenes',{ggg},'destroy',1,'stats',true,'filetypes',file_types);
fig_process_2024(cnt, qv, fc, lbl, gene_names, exp_def);
    
    
    
```
