## Supplemental File S1 for "KRAS G12 mutant alleles differentially control glutamine metabolism via FOXO1": kegg_api.m

```
function kegg_genes = kegg_api(pathway_name)

    webdata_genes  = webread(['http://rest.kegg.jp/get/' pathway_name]);
    data_genes     = webdata_genes(findstr(webdata_genes,'GENE')+12 : (min([ findstr(webdata_genes,'COMPOUND'),...
                                                                             findstr(webdata_genes,'REFERENCE')])-1));
    [idx0, idx1] = regexp(data_genes,'\D*\s\w*;');
    for gi=1:numel(idx0)
        kegg_genes{gi} = data_genes(idx0(gi)+2:idx1(gi)-1);
    end

    display(['Loaded ' num2str(numel(kegg_genes)) ' gene names from KEGG server']);
```
