## Supplemental File S1 for "KRAS G12 mutant alleles differentially control glutamine metabolism via FOXO1": kegg_cluster.m

```
function kegg_cluster(kegg_pathway, gene_names, fc, qv, se, cnt

    % Init clustergram
    LUT = flipud(cbrewer2('div','RdBu',256));
    stat_dist = 'euclidean';
    stat_link = 'average';

    slbl = {'G12A','G12C','G12D','G12V'};
    
    cDFE_SW = (1:4);
    gn = numel(gene_names);
    smp_idx = (1:25);
    dfe_idx = cDFE_SW(opts.mutants);

    kegg_genes = kegg_api(kegg_pathway); 

    
    %% clustering fold changes
    
    cgo2 = clustergram(fc([1 2 3 4],gen_idx)',...
                      'Standardize',3,...                   % 3: no, 2: genes, 1: mutants
                      'cluster',1,...                       % 1: genes, 2: samples, 3: both
                      'colormap',LUT,...            
                      'displayrange',2,...
                      'rowpdist',stat_dist,...
                      'columnpdist',stat_dist,...
                      'linkage',stat_link,...
                      'RowLabels', gene_names(gen_idx),...
                      'ColumnLabels',slbl);
   
    h_cgo2  = plot(cgo2);
    hf_cgo2 = gcf;
    set(hf_cgo2,'WindowState','maximized','renderer','painters');  
    colorbar(h_cgo2);
     

    
    %% plot individual gene expression (counts) 
    
    if ~isempty(kegg_genes)
        hf_kegg = figure('name',opts.kegg,'WindowState','maximize','renderer','painters');
        ggn = numel(highlight_idx);
        for ggi=1:ggn
            subplot(round(sqrt(ggn)),ceil(sqrt(ggn)),ggi)
            boxplot(cnt(smp_idx,gen_idx(highlight_idx(ggi)))',reshape(repmat([1 2 3 4 5]',[1 5])',1,[]),'plotstyle','compact','Labels',{'G','A','C','D','V'},'colors','kbgrm')
            title(gene_names{gen_idx(highlight_idx(ggi))})
            ylabel('counts')
            axis square
        end
    end
```
