## Supplemental File S1 for "KRAS G12 mutant alleles differentially control glutamine metabolism via FOXO1": plot_gene_expression.m

```
function h = plot_gene_expression(search_gene, gene_names, fc, qv, se, cnt)
    
    h = figure;
    
    % search gene
    idx = find(ismember(gene_names,search_gene));

    % plot fold changes
    b = bar(fc(:,idx));

    % plot errror bars (SE)
    hold on
    errorbar(fc(:,idx),se(:,idx),'LineStyle','none');

    % plot p-value adjusted
    xtips2 = b.XEndPoints;
    ytips2 = b.YEndPoints;
    text(xtips2,ytips2,num2str(qv(:,idx)),'HorizontalAlignment','center', 'VerticalAlignment','bottom')


    % format the plot
    ylabel('differential expression (log2)');
    title(search_gene);
    set(gca,'xlim',[0.5 4.5]);
    set(gca,'xticklabel',{'A','C','D','V'});
    axis square
```
