## Supplemental File S1 for "KRAS G12 mutant alleles differentially control glutamine metabolism via FOXO1": rnaseq_pca.m

```
function hf_pca = rnaseq_pca(vals, gene_names, highlight_idx)

    scol = 'kbgrm';
    Z    = zscore(vals);            
    
    [coeff, score, latent, ts, explained] = pca(Z);
    
    hf_pca = figure('WindowState','maximized','renderer','painters');
    hb     = biplot(coeff(:,1:2),'Scores',score(:,1:2),'VarLabels',gene_names);
    h_smp  = findobj(hb,'Tag','obsmarker');
    h_lin  = findobj(hb,'Tag','varline');
    h_mrk  = findobj(hb,'Tag','varmarker');
    h_lbl  = findobj(hb,'Tag','varlabel');
    
    for si=1:5
        set(h_smp((1:5)+(si-1)*5),'color', scol(si),'markersize',30);
    end
    set([h_lin h_mrk ],'color',[.9 .9 .9]);
    axis square
    box on
    set(h_lbl,'fontsize',6,'color',[.4 .4 .4]);
    xlabel(['principal component 1 (' num2str(explained(1),'%2.1f') '%)'])
    ylabel(['principal component 2 (' num2str(explained(2),'%2.1f') '%)'])

    % highlight genes
    if ~isempty(highlight_idx)
        set(h_lbl(highlight_idx),'color',[1 0 0 ],'fontsize',10,'fontweight','bold');
        set(h_mrk(highlight_idx),'color',[1 .5 .5 ],'markersize',20);
        uistack(h_lbl(highlight_idx),'top')
        uistack(h_mrk(highlight_idx),'top')
    end
```
