## Supplemental File S1 for "KRAS G12 mutant alleles differentially control glutamine metabolism via FOXO1": readme.docx

Related to Supporting Table 1, Supporting Figure 1, and Figure 1.

In this compressed folder you will find the following subfolders **ACDV_Venn_FigS1A, TableS1** and **MATLAB_KRAS_SW48**.

Matlab code

The folder **MATLAB_KRAS_SW48** stores Matlab code and results used to generate Fig. 1A-C, Fig. 2E… with **generate_data_for_figures_2024.m** script. All data generated by the Samarajiwa at Imperial College London (formerly at the MRC Cancer Unit, University of Cambridge, UK) was consolidated into a matlab workspace **rnaseqdb2024.mat**.

**plot_gene_expression.m** codes for a function that plots the differential expression of genes, providing standard errors and adjusted p-values. The following Matlab command generates a bar plot for GLUL. Matlab figures were exported as EMF files and imported into Adobe Illustrator to generate panes at publication quality.

plot_gene_expression('GLUL',gene_names, fc, qv, se, cnt);

**fig_process_2024.m** codes for a function that can plot the following graphs:

- Hierarchical clustering of differentially expressed genes using gene counts
- Hierarchical clustering of differentially expressed genes using the differential expression contrasts relative to the wild-type cell line
- Vulcano plot showing genes that have significant fold changes and adjusted p-values (or FDR)
- Biplot showing how individual RNAseq experiment and sample clusters using Principal Component Analysis

The function should be called as follows:

fig_process_2024(cnt, qv, fc, lbl, gene_names, exp_def)

The variable that can be changed by the user is the structure defining the specific analysis (or experiment) that should be run (exp_def). This structure requires the following fields to be defined:

- **.trend** defines specific gene sets according to their upregulation or downregulation relative to wild-type. It supports four values:

‘+’	upregulated genes

‘-’	downregulated genes

‘’	(empty string) for any differentially regulated gene

‘=’	differentially regulated genes that exhibit the same trend across mutants

- **.fdr** defines the maximum accepted false discovery rate (or p-adjusted values) to create cohort of differentially regulated genes.
- **.foldchange** defines the minimum accepted fold change expression (in log2) to create cohort of differentially regulated genes.
- **.hits** defines the minimum number of mutant that show differences for a gene to be considered into a new gene set. For example, if analysing all four mutant cell lines, .hits=1 requires that a gene should be differentially regulated significantly in at least one mutant, while .hits=4 requires the gene to be differentially expressed in all mutants
- **.mutants** defines which mutant is analysed statistically. [1 2 3 4] or the array of constants [cMUT_A cMUT_C cMUT_D cMUT_V] requests the analysis of all mutants. However, if we want to plot only those genes altered in G12D, we would use .mutants=cMUT_D.
- **.name** simply define the folder name were to store all output files.
- **.kegg** defines a KEGG pathway (*e.g.*, hsa00250) that should be analysed. The code will restrict any analysis only on the gene sets defined by KEGG.
- **.highlightgenes** defines a list of genes that should be highlighted in the Biplots or Vulcano plots. For example .highlightgenes={‘FOXO1’,’GAD1’} will highlight the FOXO1 and GAD1 genes.
- **.filetypes** defines which output the code will return, for example {{‘emf’,’png’}} will export both vectorial and raster images.
- **.destroy** is a Boolean flag the triggers the deletion of all figures if set to true. It is a useful option when multiple figures are generated with a single script. Data will be still accessible through the saved files.
- **.stats** is a Boolean flag, usually set to true, that activate statistical filtering that uses .fdr, .foldchange and .hits. If set to false, the script will not filter out any gene.

Licences: we are using cbrewer2 that can be downloaded from Matlab Exchange to implement Cynthia Brewer [http://colorbrewer.org](http://colorbrewer.org/) lookuptables.

The script **generate_data_for_figures_2024.m** contains several examples on how to use the code.

Venn diagram

The results used to plot the Venn diagram in Fig. S1A are available in the folder **ACDV_Venn_FigS1A**. The Venn diagram was generated with the webtool <https://bioinformatics.psb.ugent.be/webtools/Venn/> freely available at the webpage of the Bioinformatics & Evolutionary Genomics group, University of Gent, Belgium.

In the folder, a text file with the gene sets corresponding to the intersections shown below is available.

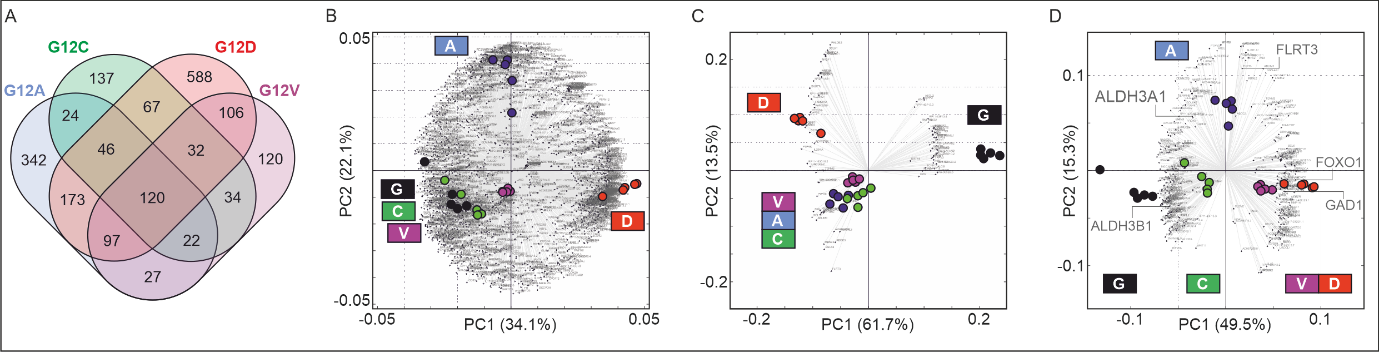

Enrichment analysis

Examples of data analysis utilized for Figure 1 and described also in Table S1 are available in the folder **TableS1**.

We provide the Jupyter notebooks generated with EnrichR. Link to the datasets stored on EnrichR are provided in Table S1. The data is one example of enrichment analysis using KEGG 2021 pathways. In this compressed folder we also include the comma separated files providing p- and q- values generated by EnrichR, and the plots related to the relevant clusters. The data is summarized in the figure below.

Each panel shows a volcano plot generated with EnrichR’s Appyter included in the Jupyter notebooks and generated using the -log10(p-values) plotted versus the odds ratios. The blue circles represent the KEGG 2021 pathways that are identified as enriched of differentially regulated genes in G12A, G12C, G12D, and G12V, in at least one mutant cell line, and in all cell lines with similar trends (*i.e.*, upregulated or downregulated in each mutant cell line).

The last two plots shows the pathways enriched of genes differentially regulated with similar trends in G12D and G12V mutant cells, and those genes that are upregulated in these two mutants that represent the cluster shown in Fig. 1D with the orange bar.

The lists below the plot show the pathways identified by EnrichR as statistically significant. The results and methods are discussed in the main manuscript. These files are provided to ensure long-term access to the analyses in the case EnrichR will become unavailable in the future.

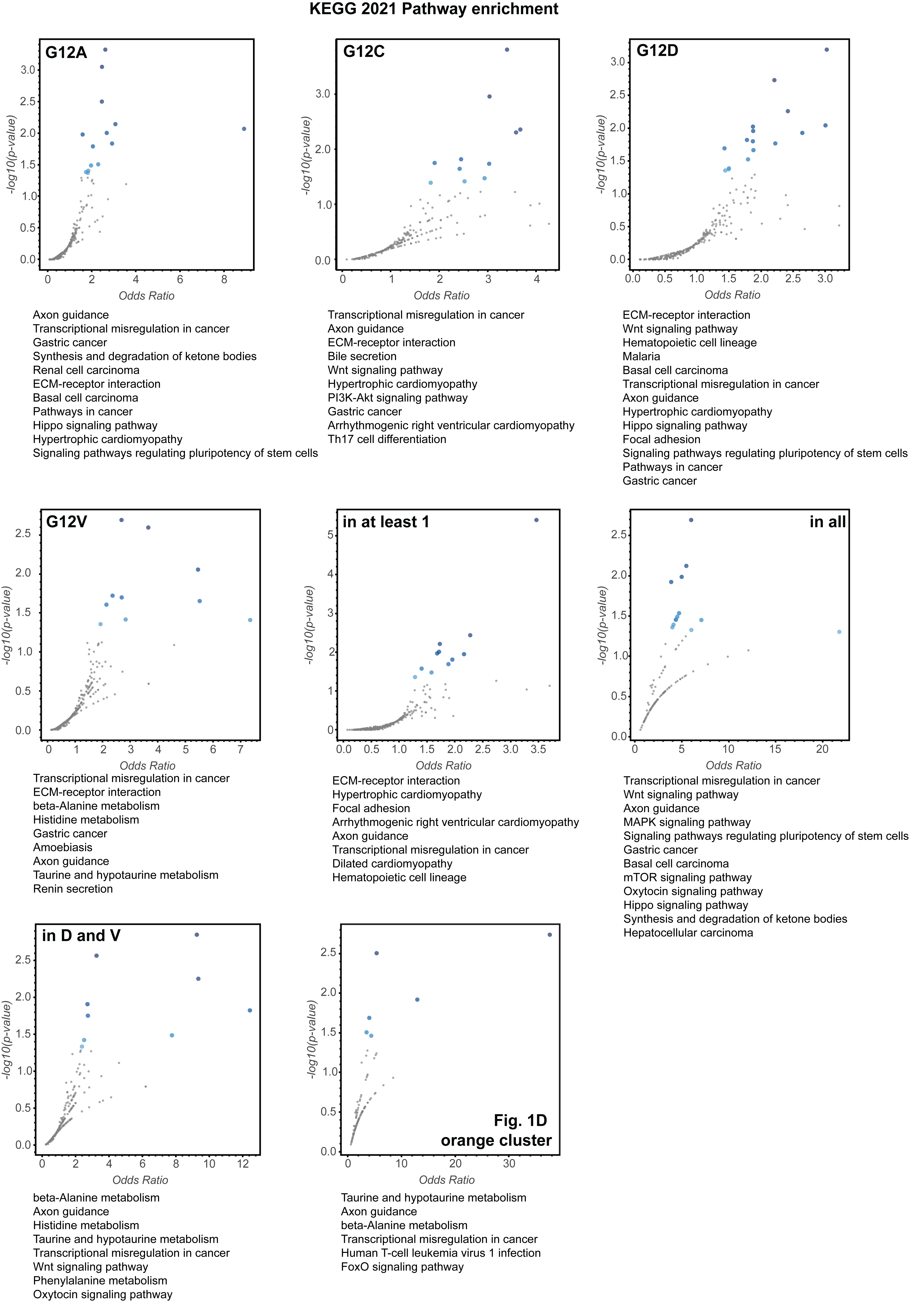
