## Supplemental File S1 for "KRAS G12 mutant alleles differentially control glutamine metabolism via FOXO1": readme.pdf

The script **generate_data_for_figures_2024.m** contains several examples on how to use the code.

### Venn diagram

The results used to plot the Venn diagram in Fig. S1A are available in the folder **ACDV_Venn_FigS1A**. The Venn diagram was generated with the webtool https://bioinformatics.psb.ugent.be/webtools/Venn/ freely available at the webpage of the Bioinformatics & Evolutionary Genomics group, University of Gent, Belgium.

In the folder, a text file with the gene sets corresponding to the intersections shown below is available.

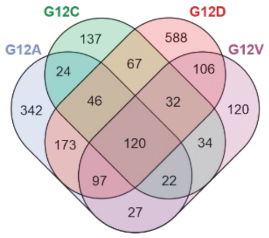


### Enrichment analysis

Examples of data analysis utilized for Figure 1 and described also in Table S1 are available in the folder **TableS1**.

We provide the Jupyter notebooks generated with EnrichR. Link to the datasets stored on EnrichR are provided in Table S1. The data is one example of enrichment analysis using KEGG 2021 pathways. In this compressed folder we also include the comma separated files providing p- and q- values generated by EnrichR, and the plots related to the relevant clusters. The data is summarized in the figure below.

Each panel shows a volcano plot generated with EnrichR's Appyter included in the Jupyter notebooks and generated using the -log10(p-values) plotted versus the odds ratios. The blue circles represent the KEGG 2021 pathways that are identified as enriched of differentially regulated genes in G12A, G12C, G12D, and G12V, in at least one mutant cell line, and in all cell lines with similar trends (*i.e.*, upregulated or downregulated in each mutant cell line).

The last two plots shows the pathways enriched of genes differentially regulated with similar trends in G12D and G12V mutant cells, and those genes that are upregulated in these two mutants that represent the cluster shown in Fig. 1D with the orange bar.

The lists below the plot show the pathways identified by EnrichR as statistically significant. The results and methods are discussed in the main manuscript. These files are provided to ensure long-term access to the analyses in the case EnrichR will become unavailable in the future.

### KEGG 2021 Pathway enrichment

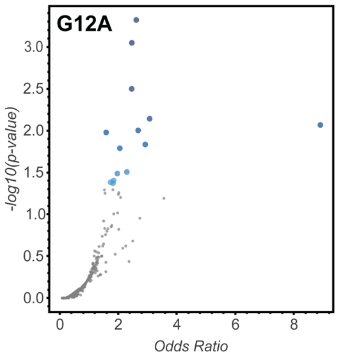


Axon guidance
Transcriptional misregulation in cancer
Gastric cancer
Synthesis and degradation of ketone bodies
Renal cell carcinoma
ECM-receptor interaction
Basal cell carcinoma
Pathways in cancer
Hippo signaling pathway
Hypertrophic cardiomyopathy
Signaling pathways regulating pluripotency of stem cells

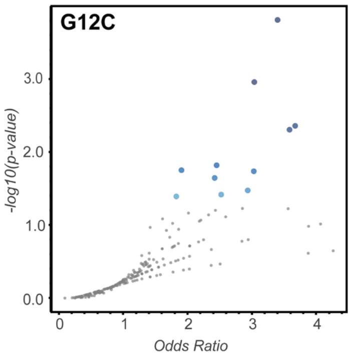


Transcriptional misregulation in cancer
Axon guidance
ECM-receptor interaction
Bile secretion
Wnt signaling pathway
Hypertrophic cardiomyopathy
PI3K-Akt signaling pathway
Gastric cancer
Arrhythmogenic right ventricular cardiomyopathy
Th17 cell differentiation

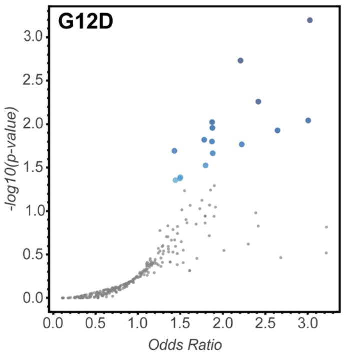


ECM-receptor interaction
Wnt signaling pathway
Hematopoietic cell lineage
Malaria
Basal cell carcinoma
Transcriptional misregulation in cancer
Axon guidance
Hypertrophic cardiomyopathy
Hippo signaling pathway
Focal adhesion
Signaling pathways regulating pluripotency of stem cells
Pathways in cancer
Gastric cancer

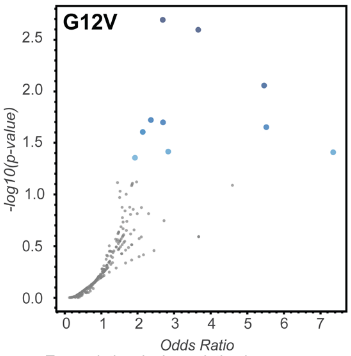


Transcriptional misregulation in cancer
ECM-receptor interaction
beta-Alanine metabolism
Histidine metabolism
Gastric cancer
Amoebiasis
Axon guidance
Taurine and hypotaurine metabolism
Renin secretion

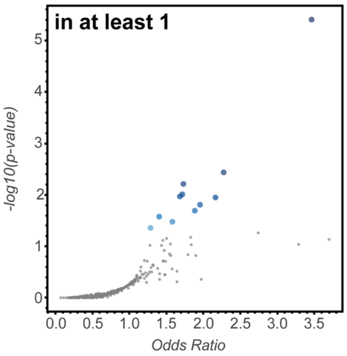


ECM-receptor interaction
Hypertrophic cardiomyopathy
Focal adhesion
Arrhythmogenic right ventricular cardiomyopathy
Axon guidance
Transcriptional misregulation in cancer
Dilated cardiomyopathy
Hematopoietic cell lineage

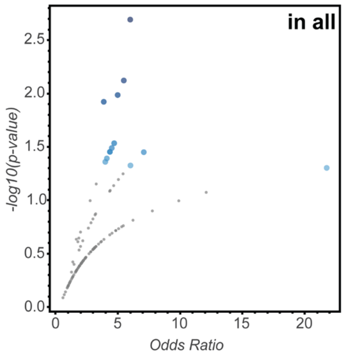


Transcriptional misregulation in cancer
Wnt signaling pathway
Axon guidance
MAPK signaling pathway
Signaling pathways regulating pluripotency of stem cells
Gastric cancer
Basal cell carcinoma
mTOR signaling pathway
Oxytocin signaling pathway
Hippo signaling pathway
Synthesis and degradation of ketone bodies
Hepatocellular carcinoma

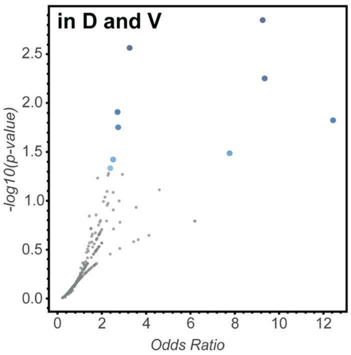


beta-Alanine metabolism
Axon guidance
Histidine metabolism
Taurine and hypotaurine metabolism
Transcriptional misregulation in cancer
Wnt signaling pathway
Phenylalanine metabolism
Oxytocin signaling pathway

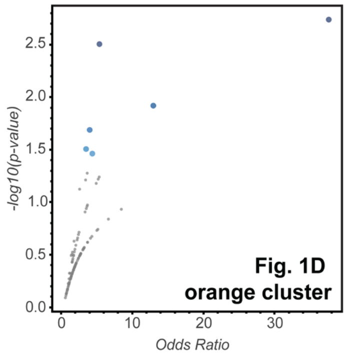


Taurine and hypotaurine metabolism
Axon guidance
beta-Alanine metabolism
Transcriptional misregulation in cancer
Human T-cell leukemia virus 1 infection
FoxO signaling pathway
