## Supplemental File S1 for "KRAS G12 mutant alleles differentially control glutamine metabolism via FOXO1": 4_g12v_KEGG2021.ipynb

```
# %% [markdown]
# Enrichment Analysis Visualizer

This appyter creates a variety of visualizations for enrichment analysis results for one selected Enrichr library, and may be run as either a standalone appyter from the [Appyter Catalog](https://appyters.maayanlab.cloud/#/Enrichment_Analysis_Visualizer) or programmatically from the [Enrichr](https://maayanlab.cloud/Enrichr/) results page. 

For simplicity, the only inputs for this appyter are a gene list and one library. Other parameters are set to default values in the cell below. You can download the notebook, change these parameters, and rerun it if you wish.

The pre-processed libraries used to create the scatter plot and hexagonal canvas visualizations can be found [here](https://github.com/MaayanLab/Enrichr-Viz-Appyter/tree/master/Enrichr-Processed-Library-Storage). 

**A link to the full analysis results on the Enrichr website can be found at the bottom of this page.**

# %% [code]
# Scatter Plot Imports
from maayanlab_bioinformatics.enrichment import enrich_crisp
import matplotlib as mpl

# Bar Chart Imports
import pandas as pd 
import numpy as np
import json
import requests
import matplotlib.pyplot as plt
import matplotlib.colors as colors
import seaborn as sns
import time
from matplotlib.ticker import MaxNLocator
from IPython.display import display, FileLink, HTML, Markdown

# Hexagonal Canvas Imports
import math
import uuid
import urllib
from textwrap import dedent
from string import Template
from operator import itemgetter

# Bokeh
from bokeh.io import output_notebook
from bokeh.plotting import figure, show
from bokeh.models import HoverTool, ColumnDataSource
from bokeh.palettes import Category20
output_notebook()

# %% [code]
gene_list_input = ['CASC21', 'RP11-845C23.3', 'KIAA1462', 'TSPEAR', 'COL1A1', 'AL163953.2', 'RP11-296L22.8', 'MYL9', 'MT-TM', 'PCDHB9', 'RP11-531A24.3', 'OPTN', 'ATF3', 'USP6NL', 'KRT16P3', 'SAMD9', 'ASCL2', 'CLDN2', 'CLDN1', 'SSFA2', 'APOL6', 'NUDT16P1', 'RP11-1055B8.1', 'FLRT3', 'LURAP1L', 'AC010980.2', 'ST3GAL6', 'PROK1', 'AFAP1L2', 'STARD13', 'CYP4F3', 'AC107016.1', 'NR2F1', 'HRNR', 'FOSL2', 'FAM111B', 'PLSCR4', 'FOXL2NB', 'KCNMA1', 'GPR183', 'FAT1', 'LINC01515', 'PRKCQ', 'RP1-69D17.3', 'PRR15', 'FLG', 'ZNF493', 'SLC30A10', 'EPS8', 'PLAC9', 'KIAA1107', 'ZC3H12D', 'PLAC8', 'RP11-307N16.6', 'TDO2', 'PCDHA1', 'ZC3H12C', 'AF127936.9', 'IL20RA', 'RP11-685M7.3', 'PCDHA6', 'PCDHA5', 'TPRG1', 'LINC00659', 'PCDHA8', 'POU5F1B', 'PCDHA7', 'RPL21P13', 'GSDMD', 'FGFBP1', 'DUSP6', 'FGFBP2', 'KCNB2', 'ANXA5', 'NR2F1-AS1', 'SEC16B', 'TESC', 'IL10RA', 'NFATC4', 'ABCA17P', 'RP11-13A1.1', 'TMC8', 'GP6', 'PLXDC1', 'DCBLD2', 'MIR612', 'SOCS2-AS1', 'GPRC5C', 'MMP14', 'AC098823.3', 'RP11-231N3.1', 'ADGRL2', 'PAM', 'ADGRL3', 'AC007255.8', 'SERPINA1', 'TRIM2', 'PEAR1', 'RP11-624L4.1', 'LDHD', 'AC005237.4', 'MIR4800', 'TERT', 'TCP11L2', 'RNF152', 'ZNF467', 'MBNL2', 'MEF2C', 'SLC2A12', 'CDKN2A', 'ODAM', 'SERPINB1', 'FLJ20021', 'GPR1', 'CD180', 'WNT7B', 'CTC-550B14.7', 'DHRS2', 'FAM225B', 'AC068724.1', 'FAM225A', 'RNF125', 'CH17-360D5.2', 'PLCB4', 'MAP1B', 'SLCO3A1', 'RP11-589N15.2', 'RP11-428K3.1', 'LGR6', 'MT-TQ', 'CDKN1C', 'RP11-114H23.3', 'AC007277.3', 'RP11-114H23.2', 'AOAH', 'RP11-114H23.1', 'RP11-554D20.1', 'PRSS22', 'SNCAIP', 'CHN2', 'RP11-342M3.5', 'CARNS1', 'ANXA8', 'GPAT3', 'GSN', 'RP11-64K12.4', 'S100A3', 'PRRX1', 'DISP2', 'PVRL4', 'MIR210HG', 'PARK2', 'PHLDB2', 'PHLDB1', 'VTCN1', 'TNC', 'AHR', 'PAPSS2', 'FOXO1', 'CTGF', 'HMGN5', 'RP1-244F24.1', 'ABLIM3', 'HAS2', 'PIPOX', 'SLC16A9', 'IL12RB2', 'KRTAP3-1', 'SLC16A4', 'MAP3K5', 'BCL2L15', 'SEMA6C', 'AC005776.1', 'IGFBP4', 'IGFBP3', 'DYRK1B', 'B3GALT5', 'CYP4F12', 'C1QL4', 'FBXO32', 'CTA-363E6.5', 'ZNF419', 'ZNF772', 'RP11-883A18.3', 'TGFB2-AS1', 'PCDHAC1', 'RIMS2', 'RPTN', 'RP11-427J23.1', 'RP11-353N14.4', 'SESN3', 'RP11-823E8.3', 'NDP', 'PLXNA2', 'BMF', 'AC012507.3', 'LINC01468', 'PCDHAC2', 'KRT18P10', 'HMGCS2', 'SETDB2', 'RP11-460I13.2', 'AVPR2', 'MACC1', 'PTPN12', 'RGMB-AS1', 'FAM27E2', 'BCL6', 'PCCA', 'MYO3B', 'LANCL3', 'DLX2', 'CNTNAP2', 'AC067956.1', 'SHC2', 'RP11-331F4.5', 'CELF2', 'RP11-1223D19.2', 'RP11-1223D19.3', 'CYP3A5', 'GS1-358P8.4', 'RP11-64C12.6', 'TGFB2-OT1', 'RP11-528I4.2', 'MROH7', 'SKIL', 'TNFRSF14', 'NTRK2', 'SMIM14', 'MYO16', 'NDP-AS1', 'CTRB2', 'LINC00202-1', 'CPT1C', 'AC093843.1', 'ASPG', 'ZNF615', 'MET', 'TRIM38', 'KCTD12', 'EVL', 'SMPDL3B', 'CEBPB', 'SEMA3A', 'DRAXIN', 'MDGA1', 'CH17-232I21.1', 'AC007405.4', 'AC007405.6', 'TSPAN12', 'PBXIP1', 'THSD4', 'PSD3', 'TRIM29', 'PSD2', 'DHX58', 'TNNI2', 'NHS', 'CEP152', 'PCDH7', 'ITGB7', 'ADAMTS9', 'LINC00698', 'CTHRC1', 'KCNJ4', 'RP11-54O7.11', 'CTC-467M3.1', 'IL1R2', 'FBXO2', 'ACSL5', 'PLEKHA4', 'OTOP2', 'SEPP1', 'APLN', 'DCLK1', 'MALL', 'CTD-2280E9.1', 'RP11-100G15.7', 'AC005592.1', 'AC005592.2', 'NINJ2', 'MT1G', 'GSAP', 'AC144831.1', 'HRASLS', 'PLEKHB1', 'TRANK1', 'ANTXR2', 'APCDD1', 'RASIP1', 'RP11-429J17.5', 'MGC45922', 'AC113949.1', 'CHAC1', 'SUSD4', 'RPRML', 'RP5-887A10.1', 'APOBEC3D', 'FER1L4', 'APOBEC3F', 'EGR3', 'BCL11B', 'HAS2-AS1', 'RAB27B', 'IGFL2', 'ANK2', 'TSPEAR-AS2', 'PUM3', 'TSPEAR-AS1', 'RP5-1120P11.1', 'BBS12', 'PARP10', 'KLHDC8A', 'PPP1R1C', 'RGL3', 'CAPS2', 'APOBEC3C', 'RP11-324O2.3', 'RP11-506B6.6', 'RP11-392O17.1', 'SPON1', 'MAGED1', 'RP11-43F13.3', 'RP11-43F13.4', 'IFI35', 'PCDH19', 'SLC6A20', 'FLG-AS1', 'LGALS1', 'INSC', 'TSPAN8', 'ARSJ', 'ZBED2', 'LGALS9', 'TSPAN1', 'TNS4', 'RALGPS2', 'CDSN', 'IL15', 'CCDC114', 'IZUMO1', 'SIGLEC10', 'KRT14', 'BTN3A3', 'RP3-523K23.2', 'ID2-AS1', 'PTPRC', 'RP11-346D6.4', 'NMU', 'KRT18', 'KRT16', 'ID4', 'VWA2', 'ARHGEF3', 'MGC32805', 'SGK1', 'RP11-90L1.8', 'TENM2', 'CSRNP3', 'IGSF5', 'PTPRO', 'KLK5', 'KRT23', 'PTPRK', 'PPP3CA', 'IGSF22', 'RP3-395M20.8', 'SOCS2', 'PAG1', 'DMRT2', 'DMRT1', 'GAD1', 'SORBS2', 'ACTL8', 'RP11-30K9.5', 'RP11-5O17.1', 'ANO1', 'KIF26A', 'NOV', 'AC022101.1', 'TJP3', 'NRP2', 'PTGER4', 'SLC43A2', 'TAF9P3', 'AC064874.1', 'RP11-734K21.5', 'SIPA1L2', 'AKAP12', 'AC078883.3', 'AC078883.4', 'CREG2', 'CTNNA2', 'MGAT5', 'AP006621.9', 'TGFB2', 'AP000692.10', 'CTD-2540F13.2', 'LMO4', 'TRPV3', 'CRIM1', 'RP11-712B9.2', 'PCDHA13', 'PCDHA12', 'PCDHA11', 'MYO16-AS1', 'MYCN', 'SPOCD1', 'CYP24A1', 'DOK7', 'TRPV6', 'UCA1', 'CCDC152', 'YPEL3', 'PRDM8', 'GDA', 'MIR3074', 'LRCOL1', 'CTD-2325A15.5', 'RGS16', 'NAPG', 'NKD2', 'GSN-AS1', 'PSORS1C2', 'FBLN5', 'EFNB3', 'ATOH8', 'NT5E', 'DNAJC22', 'NLRP6', 'CAPN3', 'FGF20', 'SLIT1', 'MSX1', 'DENND5B', 'NLRP1', 'RP11-553K8.5', 'GGT7', 'PHGR1', 'AUTS2', 'HNRNPA1P33', 'MIR27B', 'MR1', 'RP11-103C16.2', 'SYT8', 'TMEM2', 'ALDH3A1', 'CAPRIN2', 'AL591893.1', 'CTC-448F2.6', 'COL6A5', 'RASA4DP', 'FBN1', 'MXD4', 'LINC00941', 'CALCB', 'RASEF', 'RP11-164J13.1', 'RPP30', 'DOCK8', 'SLC2A3', 'PLAU', 'ARHGAP18', 'ETS2', 'PPP1R9A', 'ARHGAP23', 'CALHM2', 'DEAF1', 'ALDH3B1', 'COBLL1', 'MUC13', 'IL6R', 'RP11-442H21.2', 'RASA4B', 'ABCC3', 'MYRIP', 'SSTR1', 'WNT16', 'SHANK1', 'MANSC4', 'RP11-58O9.2', 'ALPL', 'PDE3A', 'IRF7', 'RP11-334C17.5', 'RP1-261G23.7', 'SMIM3', 'SMIM5', 'RP11-297D21.2', 'DNAH3', 'RP4-742J24.2', 'MAST4', 'AC011747.6', 'LAMC3', 'SLC40A1', 'TNFAIP2', 'MOSPD1', 'DPP4', 'ENHO', 'MIR6724-3', 'KLRC4-KLRK1', 'MIR6724-4', 'PDLIM1', 'SCIN', 'MIR6724-1', 'RP11-311F12.2', 'WNT11', 'ELDR', 'RASGEF1B', 'ERICH2', 'NRIP1', 'GPER1', 'PKMP1', 'GGACT', 'RP4-798P15.3', 'CCDC186', 'DACT1', 'FAM83B', 'DAPK1', 'MIR23B', 'PDE4D', 'ZIC5', 'SPATA3-AS1', 'EPOR', 'TIAM1', 'SGPP2', 'RP11-596C23.6', 'CPED1', 'PDE10A', 'HSH2D', 'FAM167A', 'ALDH1A1', 'RP11-1105G2.3', 'DDIT4', 'ADGRB1', 'AC004510.3', 'SPRY4', 'CTD-2223O18.1', 'RP11-485O10.2', 'RP11-553L6.5', 'TRIB3', 'SNED1', 'KRT81', 'AC019117.1', 'LAMA2', 'TREH', 'LAMA4', 'ATP2A3', 'TCHH', 'PRKRIRP3', 'CIART', 'KRT17P6', 'ZIC2', 'EPS8L1', 'EPS8L2', 'CASC19', 'RP11-246E12.2', 'CRYM']
enrichr_library = 'KEGG_2021_Human'

# %% [code]
genes = [x.strip() for x in gene_list_input]

# %% [code]
# Error handling
class NoResults(Exception):
    pass 
class APIFailure(Exception):
    pass

# %% [code]
# Enrichr API Function for Manhattan Plot and Bar Chart
# Takes a gene list and Enrichr libraries as input
def Enrichr_API(enrichr_gene_list, all_libraries):

    all_terms = []
    all_pvalues =[] 
    all_adjusted_pvalues = []

    for library_name in all_libraries : 
        ENRICHR_URL = 'https://maayanlab.cloud/Enrichr/addList'
        genes_str = '\n'.join(enrichr_gene_list)
        description = ''
        payload = {
            'list': (None, genes_str),
            'description': (None, description)
        }

        response = requests.post(ENRICHR_URL, files=payload)
        if not response.ok:
            raise APIFailure

        data = json.loads(response.text)
        time.sleep(0.5)
        ENRICHR_URL = 'https://maayanlab.cloud/Enrichr/enrich'
        query_string = '?userListId=%s&backgroundType=%s'
        user_list_id = data['userListId']
        short_id = data["shortId"]
        gene_set_library = library_name
        response = requests.get(
            ENRICHR_URL + query_string % (user_list_id, gene_set_library)
         )
        if not response.ok:
            raise APIFailure

        data = json.loads(response.text)

        if len(data[library_name]) == 0:
            raise NoResults

        short_results_df  = pd.DataFrame(data[library_name][0:10])
        all_terms.append(list(short_results_df[1]))
        all_pvalues.append(list(short_results_df[2]))
        all_adjusted_pvalues.append(list(short_results_df[6]))
        
        results_df  = pd.DataFrame(data[library_name])
        # adds library name to the data frame so the libraries can be distinguished
        results_df['library'] = library_name.replace('_', '')

    return [results_df, short_results_df, all_terms, all_pvalues, all_adjusted_pvalues, str(short_id)]

# %% [code]
# Scatter Plot Parameters
significance_value = 0.05

# Bar Chart Parameters
figure_file_format = ['png', 'svg']
output_file_name = 'Enrichr_results_bar'
color = 'lightskyblue'
final_output_file_names = ['{0}.{1}'.format(output_file_name, file_type) for file_type in figure_file_format]

# Hexagonal Canvas Parameters
canvas_color = 'Blue'
num_hex_colored = 10

# Manhattan Plot Parameters
manhattan_colors = ['#003f5c', '#7a5195', '#ef5675', '#ffa600']

# %% [markdown]
# Scatter Plot

# %% [markdown]
The scatterplot is organized so that simliar gene sets are clustered together. Larger, black-outlined points represent significantly enriched terms. 

Hovering over points will display the associated gene set name and the p-value. You may have to zoom in using the toolbar next to the plot in order to see details in densely-populated portions. Plots can also be downloaded as an svg using the save function on the toolbar.

Clusters are computed using the Leiden algorithm. Points are plotted on the first two UMAP dimensions. 

# %% [code]
### Scatter Plot Functions
def download_library(library_name):
    try: 
        df = pd.read_csv(f"https://raw.githubusercontent.com/MaayanLab/Enrichr-Viz-Appyter/master/Enrichr-Processed-Library-Storage/Clustered_Scatterplots/{library_name}.csv")
        df = df.dropna(subset=['term', 'genes'])
        return df, True
    except:
        display(HTML("<div style='font-size:1.5rem; padding:1.5rem 0;'>Failed to retrieve the selected pre-processed library.</div>"))
        return None, False

def get_library_iter(libdf):
    for row in libdf.itertuples():
        term = row.term
        gene_set = row.genes.split(' ')
        yield term, gene_set

def get_enrichment_results(genes, libdf):
    results = enrich_crisp(genes, get_library_iter(libdf), 20000, True)
    return {res[0]: res[1].pvalue for res in results}

def get_pvalue(row, enrich_results):
    if row['term'] in enrich_results.keys():
        return enrich_results[row['term']]
    else:
        return 1

def get_colors(df):
    clusters = pd.unique(df['cluster']).tolist()
    colors = list(Category20[20])[::2] + list(Category20[20])[1::2]
    color_mapper = {clusters[i]: colors[i % 20] for i in range(len(clusters))}
    return color_mapper

def get_top_10(df):
    top_indices = df[df['pvalue'] < significance_value].nsmallest(10, 'pvalue')['term'].tolist()
    return top_indices   

def get_plot(library_name):
    df, df_exists = download_library(library_name)
    if not df_exists:
        return -1, -1
    
    enrich_results = get_enrichment_results(genes, df)
    
    # add p value to the dataframe
    df['pvalue'] = df.apply(lambda row: get_pvalue(row, enrich_results), axis=1)

    # get appropriate color and size attributes
    color_mapper = get_colors(df)

    # normalize pvalues for coloring
    norm_pval = colors.Normalize(-1, 1)

    # get top 5 for each cluster
    top_terms = get_top_10(df)

    # add attributes to df
    df['color'] = df['cluster'].apply(lambda x: color_mapper[x])
    df['lw'] = df['term'].apply(lambda x: 2 if x in top_terms else 0.5)
    df['la'] = df['term'].apply(lambda x: 1 if x in top_terms else 0)
    df['size'] = df['pvalue'].apply(lambda x: 12 if x < significance_value else 5)
    df['fa'] = df['pvalue'].apply(lambda x: norm_pval(-np.log10(x)))

    if len(enrich_results.keys()) == 0:
        display(HTML(f"<div style='font-size:1.5rem; padding:1rem 0;'>There are no enriched terms for your input gene set in the <b>{library_name}</b> library.</div>"))
        tooltips = [
            ("Gene Set", "@gene_set"),
            ("Cluster", "@label")
        ]
    
    else: 
        tooltips = [
            ("Gene Set", "@gene_set"),
            ("p-value", "@p_value"),
            ("Cluster", "@label")
        ]
        
    hover_emb = HoverTool(tooltips=tooltips)
    tools_emb = [hover_emb, 'pan', 'wheel_zoom', 'reset', 'save']

    plot_emb = figure(
        plot_width=900, 
        plot_height=700, 
        tools=tools_emb
    )

    source = ColumnDataSource(
        data=dict(
            x = df['x'],
            y = df['y'],
            gene_set = df['term'],
            p_value = df['pvalue'],
            sizes = df['size'],
            colors = df['color'],
            fill_alphas = df['fa'],
            label = df['cluster'],
            line_widths = df['lw'],
            line_alphas = df['la'],
            line_colors = ['#000000']*df.shape[0]
        )
    )

    # hide axis labels and grid lines
    plot_emb.xaxis.major_tick_line_color = None
    plot_emb.xaxis.minor_tick_line_color = None
    plot_emb.yaxis.major_tick_line_color = None
    plot_emb.yaxis.minor_tick_line_color = None
    plot_emb.xaxis.major_label_text_font_size = '0pt'
    plot_emb.yaxis.major_label_text_font_size = '0pt' 

    plot_emb.output_backend = "svg"    
    
    plot_emb.xaxis.axis_label = "UMAP_1"
    plot_emb.yaxis.axis_label = "UMAP_2"
    
    s = plot_emb.scatter(
            'x', 
            'y', 
            size = 'sizes', 
            source = source, 
            color = 'colors', 
            fill_alpha = 'fill_alphas',
            line_alpha = 'line_alphas',
            line_width = 'line_widths',
            line_color = 'line_colors',
            legend_group = 'label'
        )

    plot_emb.add_layout(plot_emb.legend[0], 'right')
    
    return plot_emb, df

# %% [code]
# Display Scatter Plot
caption1 = f"**Figure 1. Scatterplot of all terms in the {enrichr_library} gene set library.** Each point represents a term in the library. \
    Term frequency-inverse document frequency (TF-IDF) values were computed for the gene set corresponding to each term, and UMAP was  \
    applied to the resulting values. The terms are plotted based on the first two UMAP dimensions. Generally, terms with more similar \
    gene sets are positioned closer together. Terms are colored by automatically identified clusters computed with the Leiden algorithm \
    applied to the TF-IDF values. The darker and larger the point, the more significantly enriched the term. Hovering over points \
    will display the term, the p-value from the enrichment calculation, and the automatically assigned cluster."

plot, df = get_plot(enrichr_library)
if plot == -1:
    display(HTML("<div style='font-size:1rem;'>Unable to create scatter plot visualization.</div>"))
else:
    display(HTML(f"<div style='font-size:1.5rem;'>Scatter plot visualization for {enrichr_library}.</div>"))
    show(plot)
    display(Markdown(caption1))

# %% [markdown]
# Bar Chart

# %% [markdown]
The bar chart shows the top 10 enriched terms in the chosen library, along with their corresponding p-values. Colored bars correspond to terms with significant p-values (<0.05). An asterisk (*) next to a p-value indicates the term also has a significant adjusted p-value (<0.05).

The bar chart can be downloaded as an image using the links below the figure. 

For creating customized bar charts for multiple libraries at once, use the standalone [Bar Chart Appyter](https://appyters.maayanlab.cloud/Enrichr_compressed_bar_chart_figure/).

# %% [code]
# Bar Chart Functions
# Takes all terms, all p-values, all adjusted p-values, plot title, Enrichr libraries, and specified figure format
def enrichr_figure(all_terms, all_pvalues, all_adjusted_pvalues, plot_names, all_libraries, bar_color): 
    # Bar colors
    if bar_color != 'lightgrey':
        bar_color_not_sig = 'lightgrey'
        edgecolor=None
        linewidth=0
    else:
        bar_color_not_sig = 'white'
        edgecolor='black'
        linewidth=1    

    plt.figure(figsize=(24, 12))
    
    i = 0
    bar_colors = [bar_color if (x < 0.05) else bar_color_not_sig for x in all_pvalues[i]]
    fig = sns.barplot(x=np.log10(all_pvalues[i])*-1, y=all_terms[i], palette=bar_colors, edgecolor=edgecolor, linewidth=linewidth)
    fig.axes.get_yaxis().set_visible(False)
    fig.set_title(all_libraries[i].replace('_', ' '), fontsize=26)
    fig.set_xlabel('−log₁₀(p‐value)', fontsize=25)
    fig.xaxis.set_major_locator(MaxNLocator(integer=True))
    fig.tick_params(axis='x', which='major', labelsize=20)
    if max(np.log10(all_pvalues[i])*-1)<1:
        fig.xaxis.set_ticks(np.arange(0, max(np.log10(all_pvalues[i])*-1), 0.1))
    for ii,annot in enumerate(all_terms[i]):
        if all_adjusted_pvalues[i][ii] < 0.05:
            annot = '  *'.join([annot, str(str(np.format_float_scientific(all_pvalues[i][ii], precision=2)))]) 
        else:
            annot = '  '.join([annot, str(str(np.format_float_scientific(all_pvalues[i][ii], precision=2)))])

        title_start= max(fig.axes.get_xlim())/200
        fig.text(title_start, ii, annot, ha='left', wrap = True, fontsize = 26)

    fig.spines['right'].set_visible(False)
    fig.spines['top'].set_visible(False)

    for plot_name in plot_names:
        plt.savefig(plot_name, bbox_inches = 'tight')
    
    # Show plot 
    plt.show()  

# %% [code]
# Display Bar Chart
caption2 = f"**Figure 2. Bar chart of top enriched terms from the {enrichr_library} gene set library.** The top 10 \
    enriched terms for the input gene set are displayed based on the -log10(p-value), with the actual p-value \
    shown next to each term. The term at the top has the most significant overlap with the input query gene set."

try:
    results = Enrichr_API(genes, [enrichr_library])
    enrichr_figure(results[2], results[3], results[4], final_output_file_names, [enrichr_library], color)
    display(Markdown(caption2))
    # Download Bar Chart
    for i, file in enumerate(final_output_file_names):
        display(FileLink(file, result_html_prefix=str('Download ' + figure_file_format[i] + ': ')))
except APIFailure:
    results = []
    display(HTML("<div style='font-size:1.5rem; padding:1rem 0;'><b>Unable to display plot because of a bad response from the Enrichr API</b></div>"))
    display(HTML("<div style='font-size:1rem; padding:1rem 0;'>Please try again later."))
except NoResults:
    results = []
    display(HTML("<div style='font-size:1.5rem; padding:1rem 0;'><b>Unable to display plot because there were no results returned from Enrichr</b></div>"))
    display(HTML("<div style='font-size:1rem; padding:1rem 0;'>Please try again with a different Enrichr library, or a different query gene list."))

# %% [markdown]
# Hexagonal Canvas

# %% [markdown]
Each hexagon in the hexagonal canvas plot represents one gene set from the selected library. The hexagons are colored based on the Jaccard similarity index between the input gene list and the gene set represented by the hexagon, with brighter color indicating higher similarity. Hexagons that are grouped together represent similar gene sets. 

Hovering over a hexagon will display the name of the gene set and the associated similarity index. 

For creating customized hexagonal canvas plots for up to two libraries at once, use the standalone [Hexagonal Canvas Appyter](https://appyters.maayanlab.cloud/#Enrichr_Canvas_Appyter).

# %% [code]
# Hexagonal Canvas Functions

def library_processing():
    # Downloads library data for the hexagonal canvas
    # Library data is pre-annealed so the canvas will have the most similar gene sets closest together
    raw_library_data = []

    try:
        library_name = enrichr_library
        with urllib.request.urlopen('https://raw.githubusercontent.com/MaayanLab/Enrichr-Viz-Appyter/master/Enrichr-Processed-Library-Storage/Annealing/Annealed-Libraries/' + enrichr_library + '.txt') as f:
            for line in f.readlines():
                raw_library_data.append(line.decode('utf-8').split("\t\t"))
        name = []
        gene_list = []
    except:
        display(HTML("<div style='font-size:1.5rem; padding:1.5rem 0;'>Failed to retrieve the selected annealed library.</div>"))
        return [], -1, -1

    for i in range(len(raw_library_data)):
        name += [raw_library_data[i][0]]
        raw_genes = raw_library_data[i][1].split('\t')
        gene_list += [raw_genes[:-1]]

    library_data = [list(a) for a in zip(name, gene_list)]

    # raw_library_data: a 2D list where the first element is the name and the second element is a list of genes associated with that name

    jaccard_indices = []
    indices = []

    for gene_set in library_data:
        intersection = [value for value in gene_set[1] if value in genes]
        index = len(intersection)/(len(gene_set[1]) + len(genes))
        jaccard_indices += [[gene_set[0], index]]
        indices += [round(index, 5)]

    # determine the dimensions of the canvas
    x_dimension = math.ceil(math.sqrt(len(indices)))
    y_dimension = math.ceil(math.sqrt(len(indices)))

    # zip name, gene_list, indices, and blank list for neighbor score then add dummy entries to the zipped list
    anneal_list = list(zip(name, gene_list, indices))

    return anneal_list, x_dimension, y_dimension

def unzip_list(anneal_list):
    unzipped_list = zip(*anneal_list)
    return list(unzipped_list)

# define a list of colors for the hexagonal canvas
def get_color(anneal_list, cut_off_value, x_dimension, y_dimension):

    # Deal with cut_off_value (only color the most significant 10/20 hexagons)
    if cut_off_value == 2.0:
        sort_list = sorted(anneal_list, key=itemgetter(2), reverse=True)
        cut_off_value = sort_list[int(num_hex_colored)-1][2]

    r_value = 0
    g_value = 0
    b_value = 0

    if canvas_color == 'Red':
        r_value = 0.0
        g_value = 0.8
        b_value = 0.8
    if canvas_color == 'Yellow':
        r_value = 0.0
        g_value = 0.3
        b_value = 1.0
    if canvas_color == 'Purple':
        r_value = 0.5
        g_value = 1.0
        b_value = 0.0
    if canvas_color == 'Pink':
        r_value = 0.0
        g_value = 1.0
        b_value = 0.2
    if canvas_color == 'Orange':
        r_value = 0.0
        g_value = 0.45
        b_value = 1.0
    if canvas_color == 'Green':
        r_value = 1.0
        g_value = 0.0
        b_value = 1.0
    if canvas_color == 'Blue':
        r_value = 1.0
        g_value = 0.9
        b_value = 0.0

    color_list = []

    unzipped_anneal_list = unzip_list(anneal_list)

    max_index = max(unzipped_anneal_list[2])

    if max_index != 0:
        scaled_list = [i/max_index for i in unzipped_anneal_list[2]]
    else:
        scaled_list = unzipped_anneal_list[2]

    for i in range(x_dimension*y_dimension):
        if i < len(unzipped_anneal_list[2]) and float(unzipped_anneal_list[2][i]) >= cut_off_value:
            color_list += [mpl.colors.to_hex((1-scaled_list[i]*r_value, 
            1-scaled_list[i]*g_value, 1-scaled_list[i]*b_value))]
        elif i < len(unzipped_anneal_list[2]):
            color_list += [mpl.colors.to_hex((1-scaled_list[i], 
            1-scaled_list[i], 1-scaled_list[i]))]
        else:
            color_list += ["#FFFFFF"]
    return color_list, max_index, cut_off_value

def init_chart():
  chart_id = 'mychart-' + str(uuid.uuid4())
  display(HTML('<script src="/static/components/requirejs/require.js"></script>'))
  display(HTML(Template(dedent('''
  <script>
  require.config({
    paths: {
      'd3': 'https://cdnjs.cloudflare.com/ajax/libs/d3/5.16.0/d3.min',
      'd3-hexbin': 'https://d3js.org/d3-hexbin.v0.2.min',
    },
    shim: {
      'd3-hexbin': ['d3']
    }
  })

  // If we configure mychart via url, we can eliminate this define here
  define($chart_id, ['d3', 'd3-hexbin'], function(d3, d3_hexbin) {
    return function (figure_id, numA, numB, colorList, libraryList, indices) {
      var margin = {top: 50, right: 20, bottom: 20, left: 50},
        width = 850 - margin.left - margin.right,
        height = 350 - margin.top - margin.bottom;

      // append the svg object to the body of the page
      var svG = d3.select('#' + figure_id)
                  .attr("width", width + margin.left + margin.right)
                  .attr("height", height + margin.top + margin.bottom)
                  .append("g")
                  .attr("transform", "translate(" + margin.left + "," + margin.top + ")");
      
      //The number of columns and rows of the heatmap
      var MapColumns = numA,
          MapRows = numB;

      //The maximum radius the hexagons can have to still fit the screen
      var hexRadius = d3.min([width/((MapColumns + 0.5) * Math.sqrt(3)), height/((MapRows + 1/3) * 1.5)]);

      //Calculate the center position of each hexagon
      var points = [];
      for (var i = 0; i < MapRows; i++) {
          for (var j = 0; j < MapColumns; j++) {
              var x = hexRadius * j * Math.sqrt(3)
              //Offset each uneven row by half of a "hex-width" to the right
              if(i%2 === 1) x += (hexRadius * Math.sqrt(3))/2
              var y = hexRadius * i * 1.5
              points.push([x,y])
          }
      }

      //Set the hexagon radius
      var hexbin = d3_hexbin.hexbin().radius(hexRadius);

      svG.append("g")
        .selectAll(".hexagon")
        .data(hexbin(points))
        .enter().append("path")
        .attr("class", "hexagon")
        .attr("d", function (d) {
            return "M" + d.x + "," + d.y + hexbin.hexagon();
        })
        .attr("stroke", "black")
        .attr("stroke-width", "1px")
        .style("fill", function (d,i) { return colorList[i]; })
        .on("mouseover", mover)
        .on("mouseout", mout)
        .append("svg:title")
        .text(function(d,i) { return libraryList[i].concat(" ").concat(indices[i]); });

      // Mouseover function
      function mover(d) {
      d3.select(this)
        .transition().duration(10)  
        .style("fill-opacity", 0.3)
      };

      // Mouseout function
      function mout(d) { 
      d3.select(this)
        .transition().duration(10)
        .style("fill-opacity", 1)
      };

  }

  })
  </script>
  ''')).substitute({ 'chart_id': repr(chart_id) })))
  return chart_id

def Canvas(numA, numB, colorList, libraryList, indices):
  chart_id = init_chart()
  display(HTML(Template(dedent('''
  <svg id=$figure_id></svg>
  <script>
  require([$chart_id], function(mychart) {
    mychart($figure_id, $numA, $numB, $colorList, $libraryList, $indices)
  })
  </script>
  ''')).substitute({
      'chart_id': repr(chart_id),
      'figure_id': repr('fig-' + str(uuid.uuid4())),
      'numA': repr(numA),
      'numB': repr(numB),
      'colorList': repr(colorList),
      'libraryList': repr(libraryList),
      'indices': repr(indices)
  })))

# %% [code]
# Display Hexagonal Canvas
caption3 = f"**Figure 3. Hexagonal canvas plot of terms from the {enrichr_library} gene set library.** Each \
    hexagon in the plot represents a single term. The brighter the color, the higher the Jaccard similarity \
    between the term gene set and the input gene set. Similar gene sets are generally grouped together. \
    The terms highlighted in blue are the most significantly overlapping with the input query gene set."

anneal_list, x_dimension, y_dimension = library_processing()
if x_dimension < 0:
    display(HTML("<div style='font-size:1rem;'>Unable to create hexagonal canvas visualization.</div>"))
else:
    color_list, scaling_factor, cut_off_value = get_color(anneal_list, 2.0, x_dimension, y_dimension)
    unzipped_anneal_list = unzip_list(anneal_list)
    if len(set(unzipped_anneal_list[2])) > 1:
        Canvas(x_dimension, y_dimension, color_list, list(unzipped_anneal_list[0]), list(unzipped_anneal_list[2]))
        display(Markdown(caption3))
    else:
        display(HTML("<div style='font-size:1.5rem; padding:1rem 0;'><b>Unable to display plot because there were no results returned from Enrichr</b></div>"))
        display(HTML("<div style='font-size:1rem; padding:1rem 0;'>Please try again with a different Enrichr library, or a different query gene list."))

# %% [markdown]
# Manhattan Plot

# %% [markdown]
In the Manhattan plot below, each line on the x-axis denotes a single gene set from the selected library, while the y-axis measures the −log₁₀(p‐value) for each gene set. 

Hovering over a point will display the name of the gene set and the associated p-value. You can also zoom, pan, and save the plot as an svg using the toolbar on the right.

For creating customized static and dynamic Manhattan plots to compare multiple libraries at once, use the standalone [Manhattan Plot Appyter](https://appyters.maayanlab.cloud/#Enrichr_Manhattan_Plot).

# %% [code]
# Manhattan Plot Functions

# Processes Enrichr data for Manhattan plots
def get_data(genes):
    # Process Enrichr data
    sorted_data = pd.DataFrame({"Gene Set": [], "-log(p value)": [], "Library": []})

    # get enrichr results from the library selected
    results_df = Enrichr_API(genes, [enrichr_library])[0]

    all_terms = []
    all_pvalues = []
    library_names = []

    all_terms.append(list(results_df[1]))
    all_pvalues.append(list(results_df[2]))
    library_names.append(list(results_df['library']))

    x=np.log10(all_pvalues[0])*-1
    sorted_terms = list(zip(all_terms[0], x, library_names[0]))
    sorted_terms = sorted(sorted_terms, key = itemgetter(0))
    unzipped_sorted_list = list(zip(*sorted_terms))

    data = pd.DataFrame({"Gene Set": unzipped_sorted_list[0], "-log(p value)": unzipped_sorted_list[1], "Library": unzipped_sorted_list[2]})

    sorted_data = pd.concat([sorted_data, data])

    # group data by library
    groups = sorted_data.groupby("Library")
    return sorted_data, groups

# Create Manhattan Plots
def manhattan(sorted_data):
    # split data frame into smaller data frames by library
    list_of_df = []
    for library_name in [enrichr_library]:
        library_name = library_name.replace('_', '')
        df_new = sorted_data[sorted_data['Library'] == library_name]
        list_of_df += [df_new]

    list_of_xaxis_values = []
    for df in list_of_df:  
        list_of_xaxis_values += df["Gene Set"].values.tolist()

    # define the output figure and the features we want
    p = figure(x_range = list_of_xaxis_values, plot_height=300, plot_width=750, tools='pan, box_zoom, hover, reset, save')

    # loop over all libraries
    r = []
    color_index = 0
    for df in list_of_df:
        if color_index >= len(manhattan_colors):
            color_index = 0 

        # calculate actual p value from -log(p value)
        actual_pvalues = []
        for log_value in df["-log(p value)"].values.tolist():
            actual_pvalues += ["{:.5e}".format(10**(-1*log_value))]

        # define ColumnDataSource with our data for this library
        source = ColumnDataSource(data=dict(
            x = df["Gene Set"].values.tolist(),
            y = df["-log(p value)"].values.tolist(),
            pvalue = actual_pvalues,
        ))
    
        # plot data from this library
        r += [p.circle(x = 'x', y = 'y', size=5, fill_color=manhattan_colors[color_index], line_color = manhattan_colors[color_index], line_width=1, source = source)]
        color_index += 1

    p.background_fill_color = 'white'
    p.xaxis.major_tick_line_color = None 
    p.xaxis.major_label_text_font_size = '0pt'
    p.y_range.start = 0
    p.yaxis.axis_label = '-log(p value)'

    p.hover.tooltips = [
        ("Gene Set", "@x"),
        ("p value", "@pvalue"),
    ]
    p.output_backend = "svg"
    
    # returns the plot
    return p

# %% [code]
# Display Manhattan Plot
caption4 = f"**Figure 4. Manhattan plot of terms from the {enrichr_library} gene set.** Each point \
    represents a single term along the x-axis. The y-values represent the -log10(p-value) corresponding \
    to the enrichment of the input gene set for the term gene set. Hovering over the point will display \
    the gene set term and the enrichment analysis p-value."

try:
    sorted_data, groups = get_data(genes)
    show(manhattan(sorted_data))
    display(Markdown(caption4))
except APIFailure:
    display(HTML("<div style='font-size:1.5rem; padding:1rem 0;'><b>Unable to display plot because of a bad response from the Enrichr API</b></div>"))
    display(HTML("<div style='font-size:1rem; padding:1rem 0;'>Please try again later."))
except NoResults:
    display(HTML("<div style='font-size:1.5rem; padding:1rem 0;'><b>Unable to display plot because there were no results returned from Enrichr</b></div>"))
    display(HTML("<div style='font-size:1rem; padding:1rem 0;'>Please try again with a different Enrichr library, or a different query gene list."))

# %% [markdown]
# Volcano Plot

# %% [markdown]
The volcano plot shows the significance of each gene set from the selected library versus its odds ratio. Each point represents a single geneset; the x-axis measures the odds ratio (0, inf) calculated for the gene set, while the y-axis gives the -log(p-value) of the gene set. 

Larger blue points represent significant terms (p-value < 0.05); smaller gray points represent non-significant terms. The darker the blue color of a point, the more significant it is. 

Hovering over points will display the corresponding gene set term, the p-value, and the odds ratio. You may have to zoom in using the toolbar next to the plot in order to see details in densely-populated portions. Plots can also be downloaded as an svg using the save function on the toolbar.

# %% [code]
def get_library(lib_name):
    '''
    Returns a dictionary mapping each term from the input library to 
    its associated geneset. 
    '''
    raw_lib_data = []

    with urllib.request.urlopen('https://maayanlab.cloud/Enrichr/geneSetLibrary?mode=text&libraryName=' + lib_name) as f:
        for line in f.readlines():
            raw_lib_data.append(line.decode("utf-8").split("\t\t"))

    name = []
    gene_list = []
    lib_data = {}

    for i in range(len(raw_lib_data)):
        name += [raw_lib_data[i][0]]
        raw_genes = raw_lib_data[i][1].replace('\t', ' ')
        gene_list += [raw_genes[:-1]]
    
    lib_data = {a[0]:a[1].split(' ') for a in zip(name, gene_list)}
    return lib_data


def volcano_plot(library_name, lib):
    '''
    Make volcano plot of odds ratio vs. significance for input library.
    '''
    enrich_results = enrich_crisp(genes, lib, 21000, True)

    res_df = pd.DataFrame(
        [ [
            term, 
            res.pvalue, 
            res.odds_ratio
        ] for (term, res) in enrich_results ], 
        columns=['term', 'pvalue', 'odds_ratio']
    )

    res_df['log_pval'] = np.negative(np.log10(res_df['pvalue']))

    cmap = mpl.cm.get_cmap('Blues_r')
    cnorm = mpl.colors.Normalize(vmin = res_df['pvalue'].min(), vmax = 0.1)

    my_colors = []
    my_sizes = []
    for row in res_df.itertuples():
        if row.pvalue < 0.05:
            my_colors += [mpl.colors.to_hex(cmap(cnorm(row.pvalue)))]
            my_sizes += [12]
        else:
            my_colors += ['#808080']
            my_sizes += [6]

    source = ColumnDataSource(
        data=dict(
            x = res_df['odds_ratio'],
            y = res_df['log_pval'],
            gene_set = res_df['term'],
            p_value = res_df['pvalue'],
            odds_r = res_df['odds_ratio'],
            colors = my_colors,
            sizes = my_sizes
        )
    )

    hover_emb = HoverTool(
        names=["res_df"], 
        tooltips="""
        <div style="margin: 10">
            <div style="margin: 0 auto; width:200px;">
                <span style="font-size: 12px; font-weight: bold;">Term:</span>
                <span style="font-size: 12px">@gene_set<br></span>
                <span style="font-size: 12px; font-weight: bold;">P-Value:</span>
                <span style="font-size: 12px">@p_value<br></span>
                <span style="font-size: 12px; font-weight: bold;">Odds Ratio:</span>
                <span style="font-size: 12px">@odds_r<br></span>
            </div>
        </div>
        """
    )

    tools_emb = [hover_emb, 'pan', 'wheel_zoom', 'reset', 'save']

    plot_emb = figure(
        plot_width = 700, 
        plot_height = 700,
        tools=tools_emb
    )

    plot_emb.circle(
        'x', 'y', size = 'sizes', 
        alpha = 0.7, line_alpha = 0, 
        line_width = 0.01, source = source, 
        fill_color = 'colors', name = "res_df"
    )

    plot_emb.xaxis.axis_label = "Odds Ratio"
    plot_emb.yaxis.axis_label = "-log10(p-value)"

    plot_emb.output_backend = "svg"
    
    return plot_emb

# %% [code]
lib_data = get_library(enrichr_library)
caption5 = f"**Figure 5. Volcano plot of terms from the {enrichr_library} gene set.** Each point \
    represents a single term, plotted by the corresponding odds ratio (x-position) and -log10(p-value) \
    (y-position) from the enrichment results of the input query gene set. The larger and darker-colored \
    the point, the more significantly enriched the input gene set is for the term. Hovering over \
    the point will display the gene set term, the odds ratio, and the enrichment analysis p-value."

if lib_data == {}:
    display(HTML("<div style='font-size:1.5rem; padding:1.5rem 0;'>Failed to access Enrichr library API, please try again later.</div>"))
else:
    try:
        plot = volcano_plot(enrichr_library, lib_data)
        show(plot)
        display(Markdown(caption5))
    except NoResults:
        display(HTML("<div style='font-size:1.5rem; padding:1rem 0;'><b>Unable to display plot because there were no results returned from Enrichr</b></div>"))
        display(HTML("<div style='font-size:1rem; padding:1rem 0;'>Please try again with a different Enrichr library, or a different query gene list."))

# %% [markdown]
# Table of significant p-values

# %% [markdown]
A downloadable table displaying the names, p-values, and q-values of significant terms in the selected library. The `q-value` is an adjusted p-value calculated using the Benjamini-Hochberg method for correction for multiple hypotheses testing. Only the top 10 significant results are displayed in this notebook, but the full table containing all significant results can be downloaded using the *Download CSV file of the full table of significant results* link.

# %% [code]
# Output a table of significant p-values and q-values

def create_download_link(df, title = "Download CSV file of the full table of significant results", filename = "Enrichment_Analysis_Visualizer_data.csv"):  
    csv = df.to_csv(filename, index = False)
    html = f'<a href="{filename}" target=_blank>{title}</a>'
    return HTML(html)

if len(results) > 0:
    res_df = pd.DataFrame(results[0]).rename(columns={
        0: 'rank',
        1: 'term',
        2: 'p-value',
        3: 'zscore',
        4: 'combined_score',
        5: 'overlap_genes',
        6: 'q-value'
    })
    sorted_res_df = res_df.sort_values(by='p-value', ascending=True)
    filtered_res_df = sorted_res_df[sorted_res_df['p-value'] <= significance_value].reset_index()
    display(HTML(f"<strong>Table of top 10 significant p-values and q-values for {enrichr_library.replace('_', ' ')}</strong>"))
    display(HTML(filtered_res_df[['term', 'p-value', 'q-value', 'overlap_genes']][:10].to_html(index = False)))
    display(create_download_link(filtered_res_df[['term', 'p-value', 'q-value']]))
else:
    display(HTML("<div style='font-size:1.5rem; padding:1rem 0;'><b>Unable to display table</b></div>"))
    display(HTML("<div style='font-size:1rem; padding:1rem 0;'>Please try again later, or start over with a new input list and/or library."))

# %% [markdown]
# Link to Enrichr

# %% [code]
# Get complete enrichment analysis results from Enrichr 
if len(results) > 0:
    url = 'https://maayanlab.cloud/Enrichr/enrich?dataset=' + results[5]
    display(HTML(f"<div style='font-size:1.25rem; padding:1rem 0;'><a href='https://maayanlab.cloud/Enrichr/enrich?dataset={results[5]}'>Access the complete enrichment analysis results on the Enrichr website. </a></div>"))
else:
    display(HTML("<div style='font-size:1.5rem; padding:1rem 0;'><b>No Enrichr results available for the current query</b></div>"))
    display(HTML("<div style='font-size:1rem; padding:1rem 0;'>Please try again with a different input list and/or library."))

# %% [code]
```
