## Supplemental File S1 for "KRAS G12 mutant alleles differentially control glutamine metabolism via FOXO1": KRAS_FOXO1_cluster.m

```
genes = {};

webdata_genes  = webread(['http://rest.kegg.jp/get/' pathway_name]);
data_genes     = webdata_genes(findstr(webdata_genes,'GENE')+12 : (min([ findstr(webdata_genes,'COMPOUND'),...
                                                                         findstr(webdata_genes,'REFERENCE')])-1));
[idx0, idx1] = regexp(data_genes,'\D*\s\w*;');
for gi=1:numel(idx0)
    genes{gi} = data_genes(idx0(gi)+2:idx1(gi)-1);
end

display(['Loaded ' num2str(numel(genes)) ' gene names from KEGG server']);

gen_idx = find(ismember(gene_names, genes));
cgo1 = cluster_folds(fld(gen_idx,:), gene_names(gen_idx), smp_lbl((numel(controls)+1):end)); 
cgo2 = cluster_counts(cnt(gen_idx,:), gene_names(gen_idx), smp_lbl);

addTitle(cgo1,['KEGG entry: ' pathway_name],'color','red');
addTitle(cgo2,['KEGG entry: ' pathway_name],'color','red');

figure
plot(cgo1)
set(gcf, 'Units', 'Normalized', 'OuterPosition', [0 0 1 1]);
saveas(gcf,[fld_sv  '/' pathway_name num2str(case_study) '_fld.png'])

figure
plot(cgo2)
set(gcf, 'Units', 'Normalized', 'OuterPosition', [0 0 1 1]);
saveas(gcf,[fld_sv  '/' pathway_name  num2str(case_study) '_cnt.png'])
```
