## Supplemental File S1 for "KRAS G12 mutant alleles differentially control glutamine metabolism via FOXO1": KRAS_FOXO1_import.m

```
%% Import data
gene_names = {};

eval(['file_names = file_names_cls' num2str(case_study) ';'])
eval(['smp_lbl    = sample_lbl'     num2str(case_study) ';'])
eval(['controls   = ctrl'           num2str(case_study) ';'])

fn = numel(file_names);
cn = numel(controls);
sn = numel(smp_lbl);
                                                                    
% Fetching gene names from file
for fi=1:fn
   display(['Fetching gene names from ''' file_names{fi} ''' ...' ]);
   import_counts = readtable([folder_import file_names{fi}],'filetype','text');
   gene_names = unique([gene_names import_counts.NAME']); 
   display(['Fetching gene names from ''' [strrep(file_names{fi}(1:end-3),'gsea','de2') 'csv'] ''' ...' ]);
   import_de = readtable([folder_import strrep(file_names{fi}(1:end-3),'gsea','de2') 'csv'],'filetype','text');
   gene_names = unique([gene_names import_de.gene_name']); 
end

% Importing data
gn  = numel(gene_names);
cnt = NaN(gn, n_repeats*(fn+1)); %+1 because of controls
[fld pdj] = deal(NaN(gn, fn)); 
typ = cell(gn,1); 
for fi=1:fn
    
   % IMPORT COUNTS
   display(['Importing raw counts from GSEA file  ''' file_names{fi} ''' ...' ]);
   import_counts = readtable([folder_import file_names{fi}],'filetype','text');
   [isgene gene_position] = ismember(import_counts.NAME,gene_names);
   if ~isempty(nonzeros(~isgene))
       error('Gene not found');
   end
   
   if ismember(fi, controls) % load control
       cnt(gene_position,(fi-1)*n_repeats+(1:n_repeats)) = table2array(import_counts(:,(3+n_repeats):end));
   end
   cnt(gene_position,n_repeats*(cn+fi-1)+(1:n_repeats)) = table2array(import_counts(:,3:2+n_repeats));
   
   % IMPORT DE
   display(['Importing differential expression values from DE2 file  ''' [strrep(file_names{fi}(1:end-3),'gsea','de2') 'csv'] ''' ...' ]);
   import_de =  readtable([folder_import strrep(file_names{fi}(1:end-3),'gsea','de2') 'csv'],'TreatAsEmpty','NA');
   [isgene gene_position] = ismember(import_de.gene_name,gene_names);
   if ~isempty(nonzeros(~isgene)), error('Gene not found');  end
   fld(gene_position,fi) = import_de.log2FoldChange;       
   pdj(gene_position,fi) = import_de.padj;       
   typ(gene_position) = import_de.gene_type;       
   
end
```
