## Supplemental File S1 for "KRAS G12 mutant alleles differentially control glutamine metabolism via FOXO1": KRAS_LN_FOXO1_FIG4.m

```
clear all

% initialize analysis

% set paths
folder_import = '.\data\';
fld_sv        = '.\output\';

% 1-	All SW48 cells in LN
file_names_cls1 = {'sw_gsea_G12A_LN___G12G_LN.gct',...
                   'sw_gsea_G12C_LN___G12G_LN.gct',...
                   'sw_gsea_G12D_LN___G12G_LN.gct',...
                   'sw_gsea_G12V_LN___G12G_LN.gct'};
sample_lbl1     = {'G','A','C','D','V'};
ctrl1           = [1];

% 2-	SW48 Par, G12D and G12V in DMSO and iFOXO1. 
file_names_cls2 = {'sw_gsea_G12G_LN_iFOXO1___G12G_LN_DMSO.gct',...
                   'sw_gsea_G12D_LN_iFOXO1___G12D_LN_DMSO.gct',...
                   'sw_gsea_G12V_LN_iFOXO1___G12V_LN_DMSO.gct'};
sample_lbl2     = {'G-DMSO','D-DMSO','V-DMSO','G-iFOXO1','D-iFOXO1','V-iFOXO1'};
ctrl2 = [1 2 3];

% number of repeats
n_repeats = 5;    


% initialize clustering function
%% Define clustering functions

stat_dist = 'euclidean';                    % define distance for clustergram
stat_link = 'complete';                     % define linkage for clustergrum

clsimpute = @(x)arrayfun(@(y)isnan(y)*0 + ~isnan(y)*nanmean([0 y]),x);

cluster_folds = @(vals, genes, smp) clustergram( vals,...
                                                      'Standardize',3,...                   % 3: no, 2: genes, 1: mutants
                                                      'cluster',1,...                       % 1: genes, 2: samples, 3: both
                                                      'colormap',redbluecmap,...                    % 
                                                      'displayrange',2,...
                                                      'rowpdist',stat_dist,...
                                                      'columnpdist',stat_dist,...
                                                      'linkage',stat_link,...
                                                      'RowLabels', genes',...
                                                      'ImputeFun', @clsimpute,...
                                                      'ColumnLabels',smp);
                                                  
                                                  
cluster_counts  = @(vals, genes, smp) clustergram( vals,...
                                                      'Standardize',2,...                   % 3: no, 2: genes, 1: mutants
                                                      'cluster',1,...                       % 1: genes, 2: samples, 3: both
                                                      'colormap',redbluecmap,...                   
                                                      'displayrange',2,...
                                                      'rowpdist',stat_dist,...
                                                      'columnpdist',stat_dist,...
                                                      'linkage',stat_link,...
                                                      'RowLabels', genes',...
                                                      'ImputeFun', @clsimpute,...
                                                      'ColumnLabels',reshape(repmat(smp,[n_repeats 1]),1,[]));


%% INPUT 

% 1: SW48 in LN
case_study = 1; 
pathway_name = 'hsa00250';

KRAS_FOXO1_import;
KRAS_FOXO1_cluster;

ln_folds = fld(gen_idx,[3 4]);
ln_genes = gene_names(gen_idx); 

%% 2: FOXO1 inhibition in SW48
case_study = 2; 
KRAS_FOXO1_import;
KRAS_FOXO1_cluster;   

fx_folds = fld(gen_idx,[2 3]);
fx_genes = gene_names(gen_idx);

%% comparison

% check genes detected in both experiments
gene_names = unique([ln_genes fx_genes]);

% reindex fold values
ln_folds = ln_folds(find(ismember(gene_names, ln_genes)),:);
fx_folds = fx_folds(find(ismember(gene_names, fx_genes)),:);

%%
% correlation coefficient

outliers = {''};
% use this to remove outlier genes, otherwise comment out
outliers = {'GAD1','ASS1'};
idx = find(~ismember(gene_names, outliers));
[R,P,RL, RU]=corrcoef(ln_folds(idx,:),fx_folds(idx,:))


figure
plot(ln_folds(:,1),fx_folds(:,1),'r.','MarkerSize',40)
hold on
text(ln_folds(:,1),fx_folds(:,1),gene_names)
plot(ln_folds(:,2),fx_folds(:,2),'m.','MarkerSize',40)
text(ln_folds(:,2),fx_folds(:,2),gene_names)

 plot(ln_folds(:,1),ln_folds(:,1)/RU(1,2),'r')
 plot(ln_folds(:,1),ln_folds(:,1)/R(1,2),'r')
 plot(ln_folds(:,1),ln_folds(:,1)/RL(1,2),'r')


axis square
xlabel('D (red) and V vs WT in low nutrient')
ylabel('D and V - iFOXO1 vs DMSO')

set(gcf, 'Units', 'Normalized', 'OuterPosition', [0 0 1 1]);
saveas(gcf,[fld_sv  '/' pathway_name 'comparison_fld.png'])
saveas(gcf,[fld_sv  '/' pathway_name 'comparison_fld.emf'])
```
