## Supplemental File S1 for "KRAS G12 mutant alleles differentially control glutamine metabolism via FOXO1": readme.docx

Related to Supporting Table 3 and Supporting Figure 3

AS1842856 is a well-characterized inhibitor for FOXO1. To validate that AS1842856 inhibits FOXO1 at the sub-lethal concentration of 1 µM, we selected all genes downregulated upon FOXO1 treatment (log2<-0.2 and fdr<0.1). The lists of genes for SW48 parental, G12D and G12V were obtained by filtering the excel file “SW48_transcriptomics_iFOXO1.xlsx”.

A Venn diagram was generated with the webtool <https://bioinformatics.psb.ugent.be/webtools/Venn/> freely available at the webpage of the Bioinformatics & Evolutionary Genomics group, University of Gent, Belgium.

In the folder, a text file with the gene sets corresponding to the intersections shown below is available.

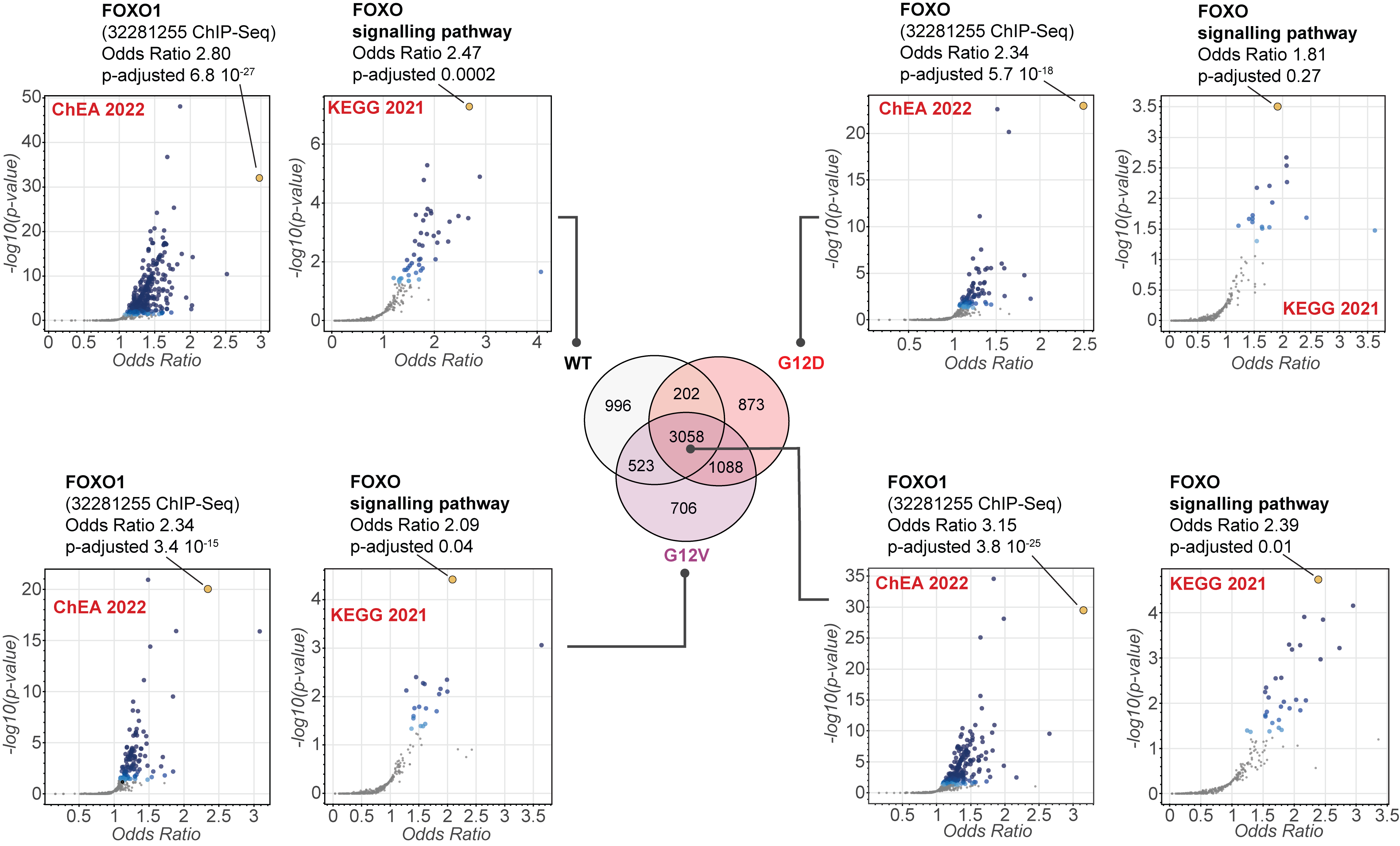

Data analysis utilized for Figure S3 and described also in Table S3. We provide the Jupyter notebooks generated with EnrichR^1^. Link to the datasets stored on EnrichR are provided in Table S3. Each panel shows a volcano plot generated with EnrichR’s Appyter included in the Jupyter notebooks and generated using the -log10(p-values) plotted versus the odds ratios. The blue circles represent either the KEGG 2021 pathways of the ChEA 2022 transcription factor target analysis that are identified as enriched of down-regulated genes upon treatment of cells with FOXO1 inhibitor.

This data was used to validate FOXO1 inhibition.

References

1.	Chen, E.Y., Tan, C.M., Kou, Y., Duan, Q., Wang, Z., Meirelles, G.V., Clark, N.R., and Ma’ayan, A. (2013). Enrichr: interactive and collaborative HTML5 gene list enrichment analysis tool. BMC Bioinformatics 14, 128. https://doi.org/10.1186/1471-2105-14-128.
