## Supplementary figures and images for "KRAS G12 mutant alleles differentially control glutamine metabolism via FOXO1"

### 13C labelling_DMSO_FOXOi_Fractions and total.pdf

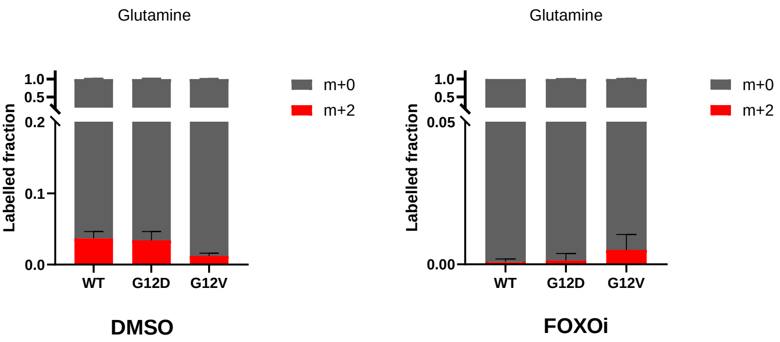

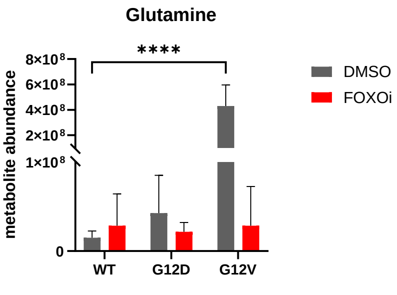

### 13C labelling_Fractions and total.pdf

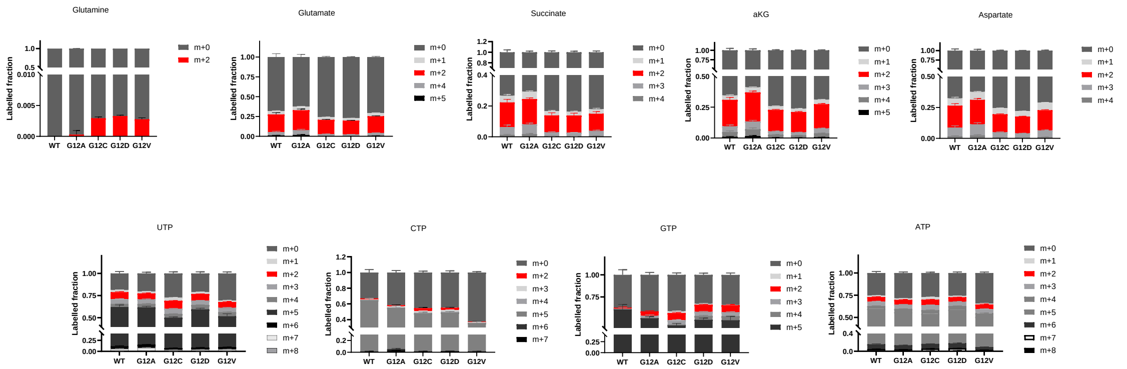

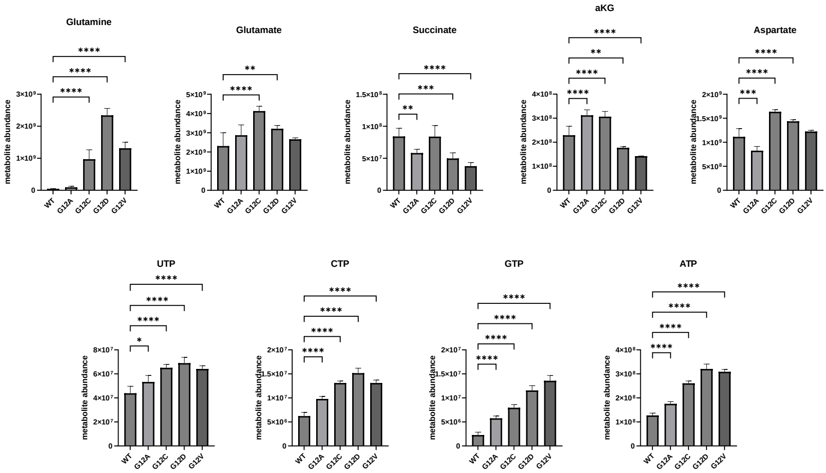

### 15_Alpha_Fractions and total.pdf

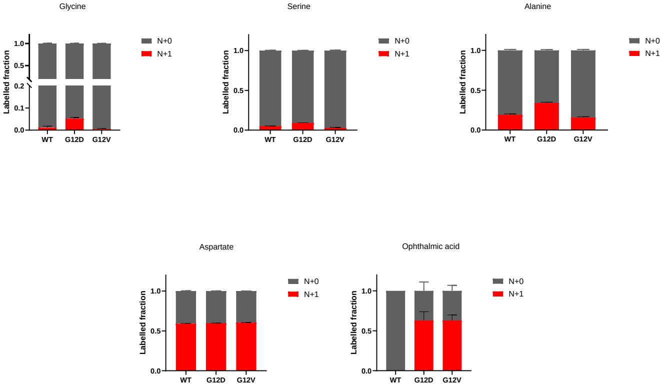

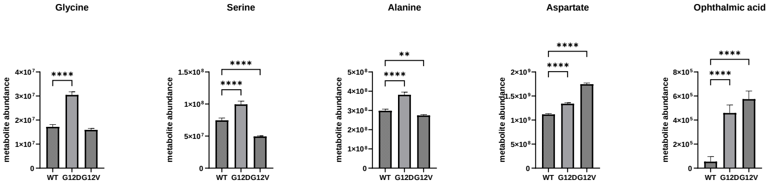

### 15_Amide_Fractions and total.pdf

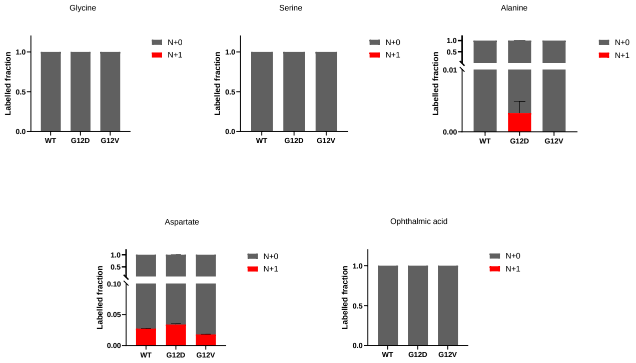

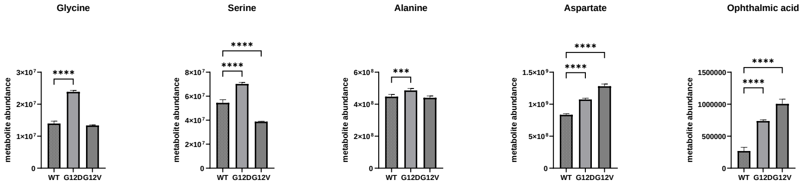

### 15N_Ammonia_Fractions and total.pdf

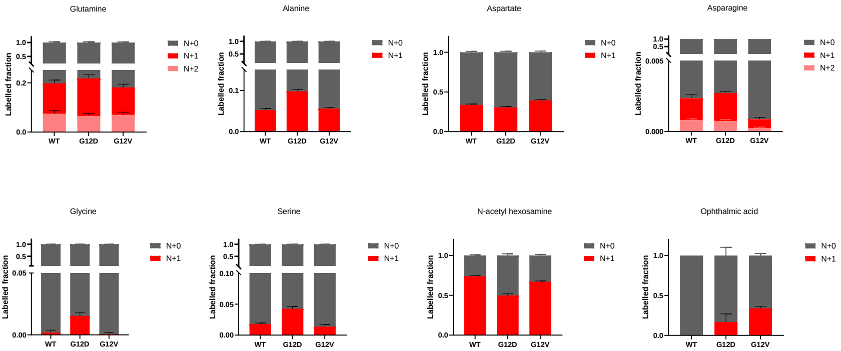

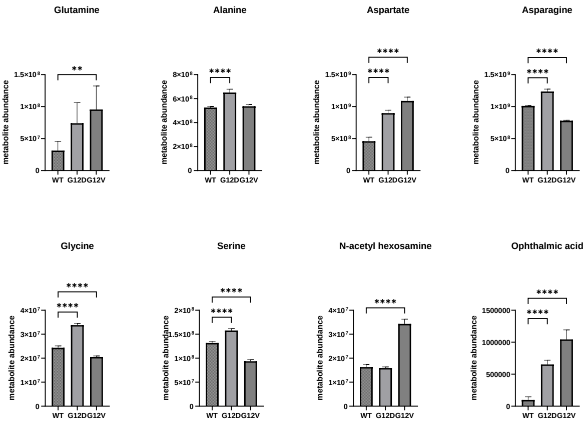

### 15N_DMSO_FOXOi_Ammonia_Fractions and total.pdf

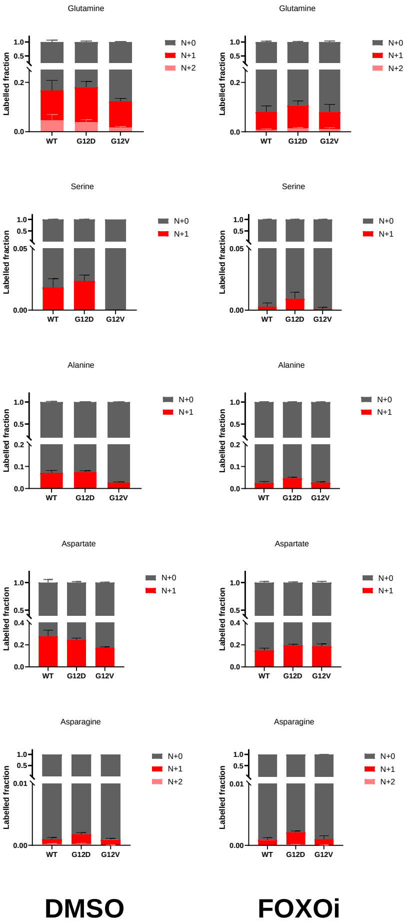

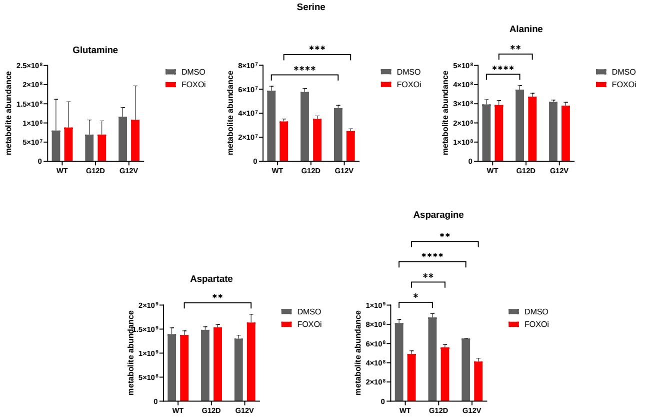

### clustergram.pdf

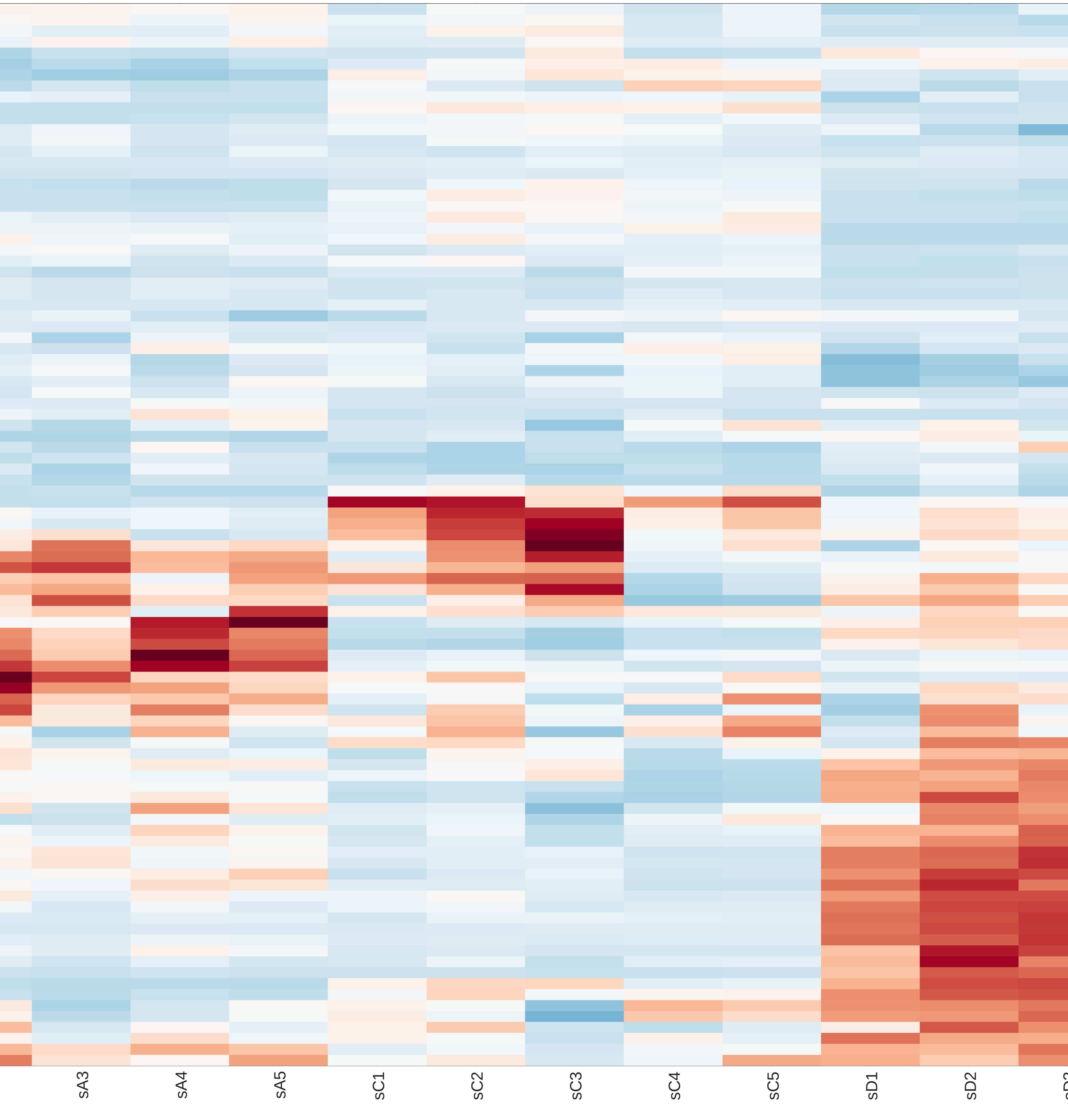

### clustergram.pdf

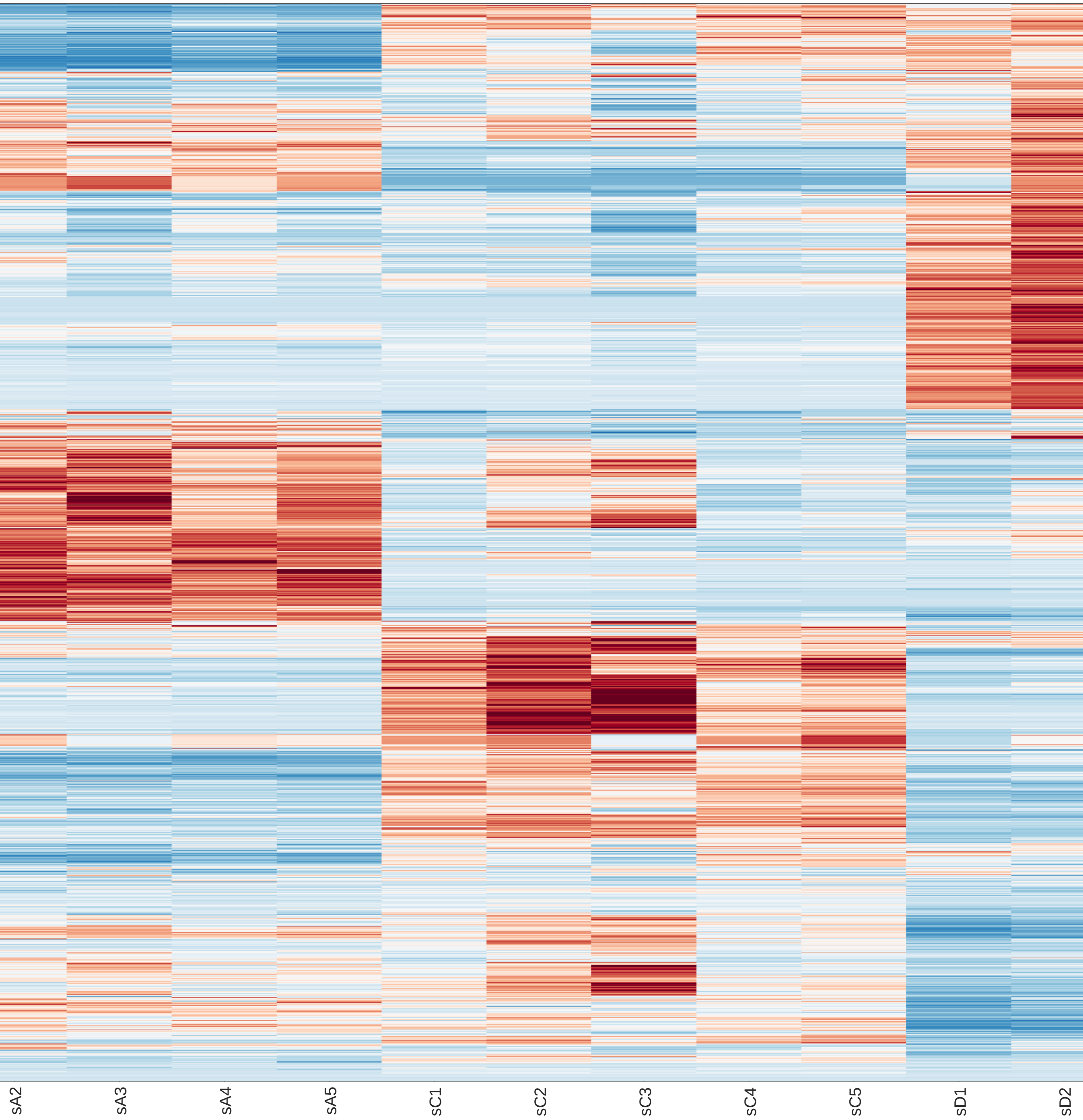

### clustergram.pdf

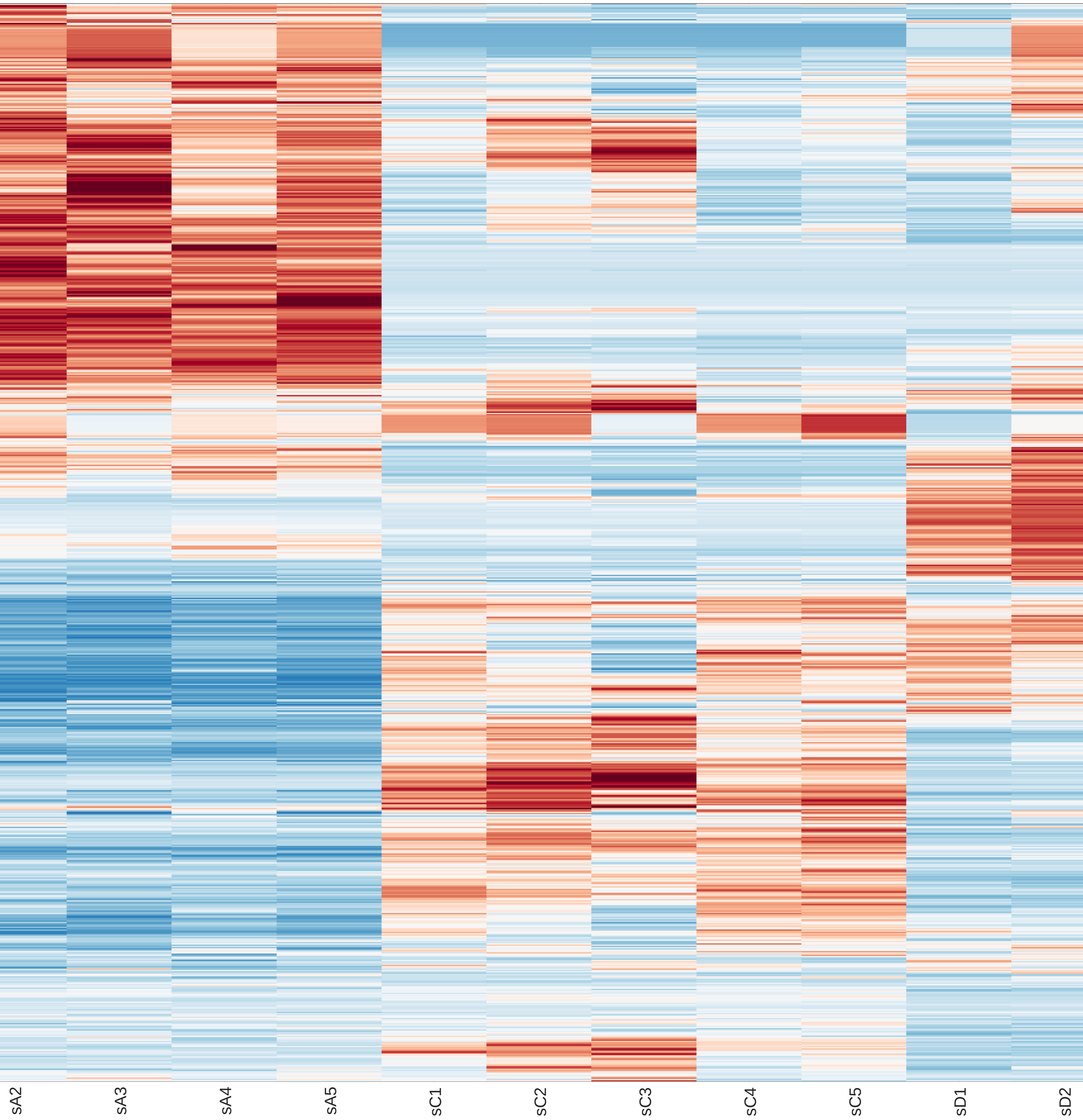

### clustergram.pdf

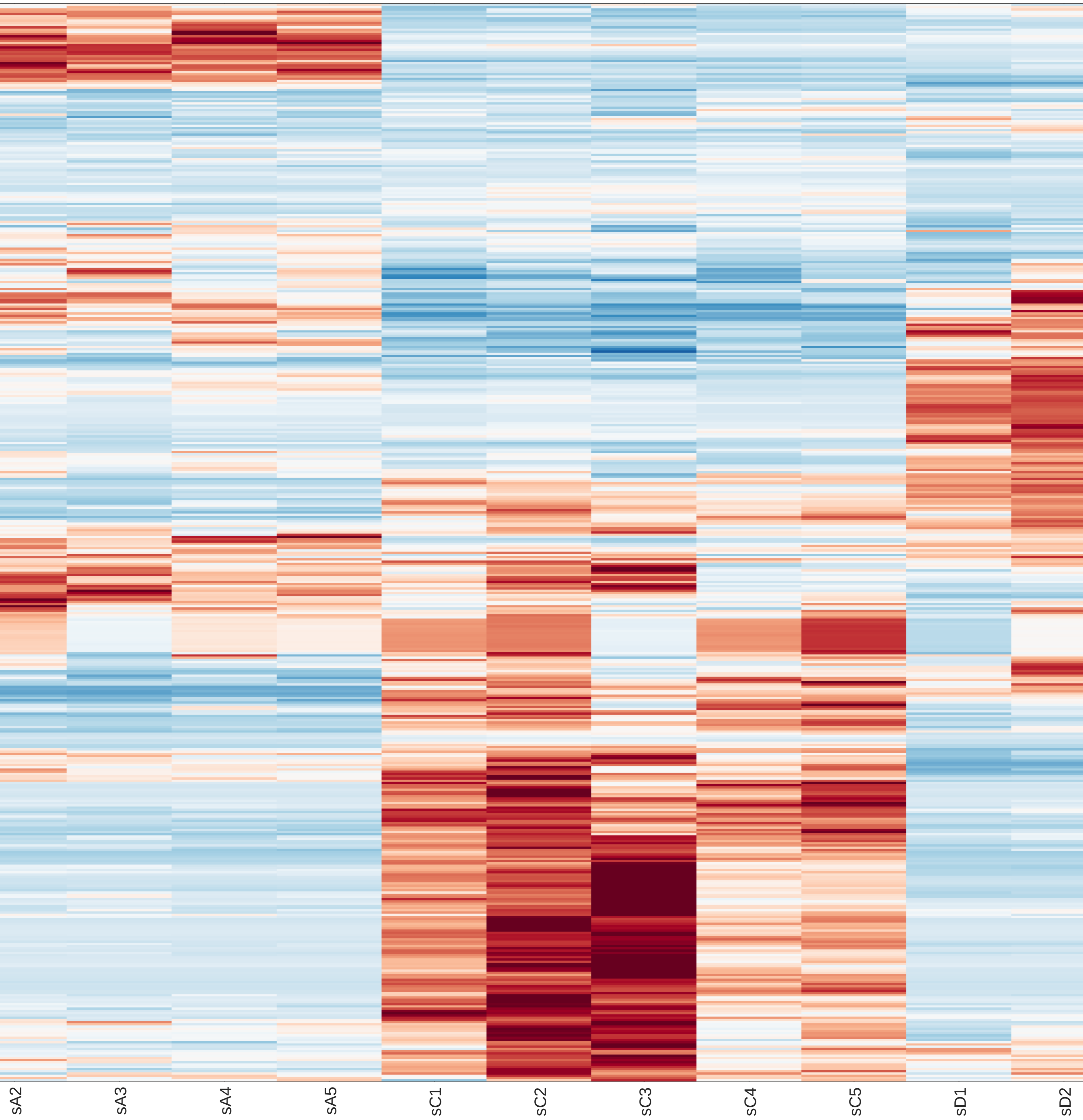

### clustergram.pdf

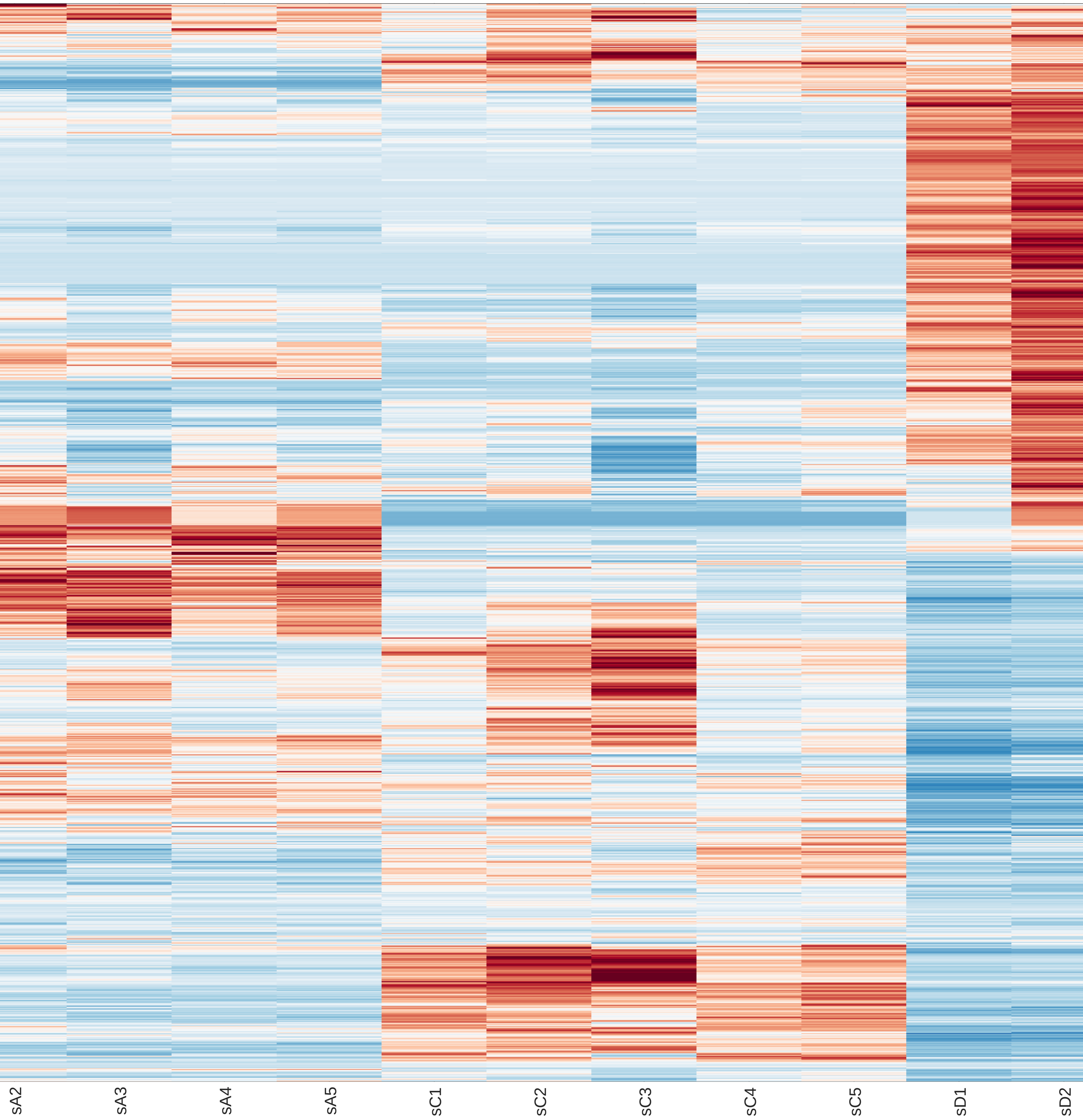

### clustergram.pdf

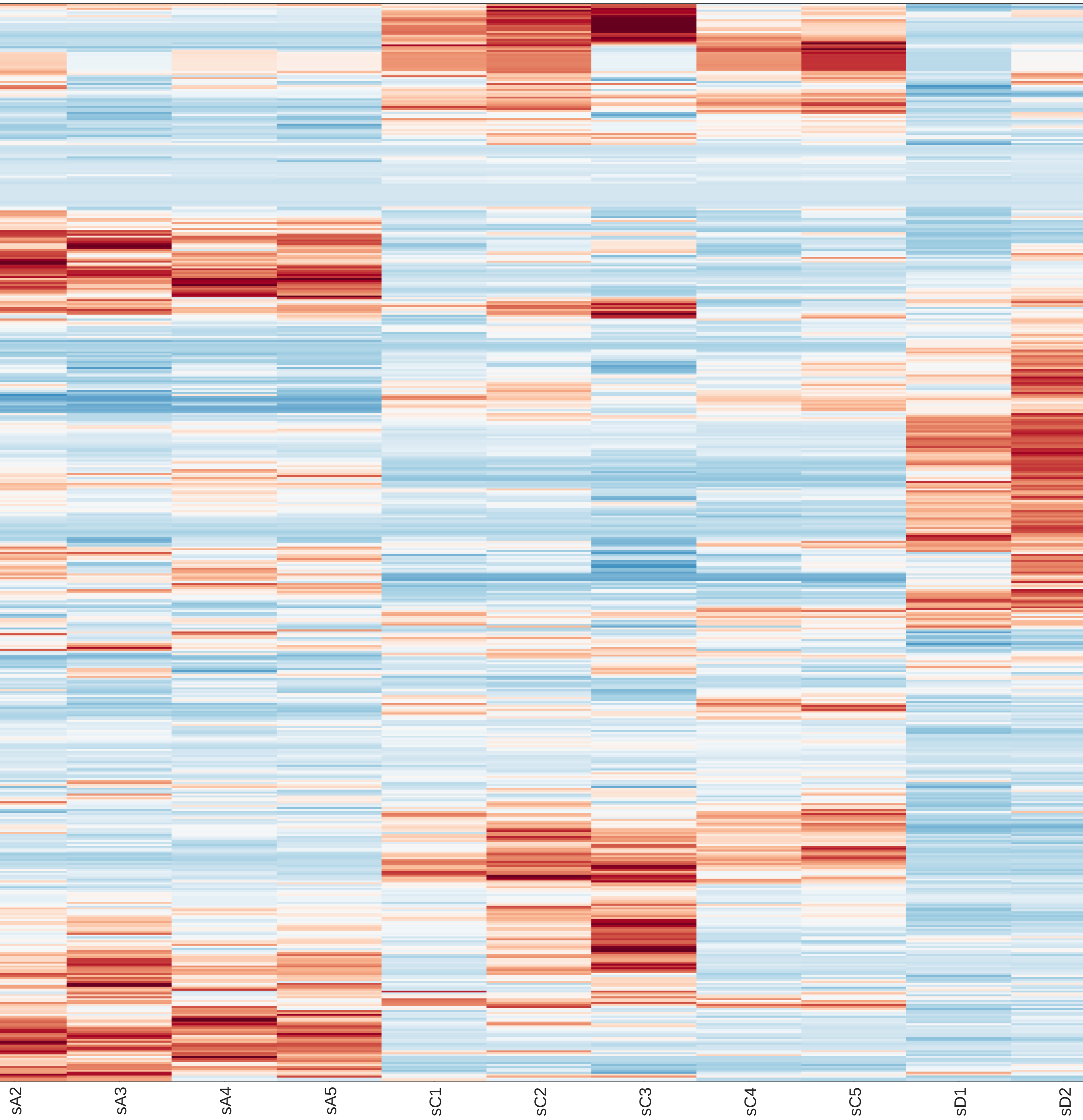

### clustergram.pdf

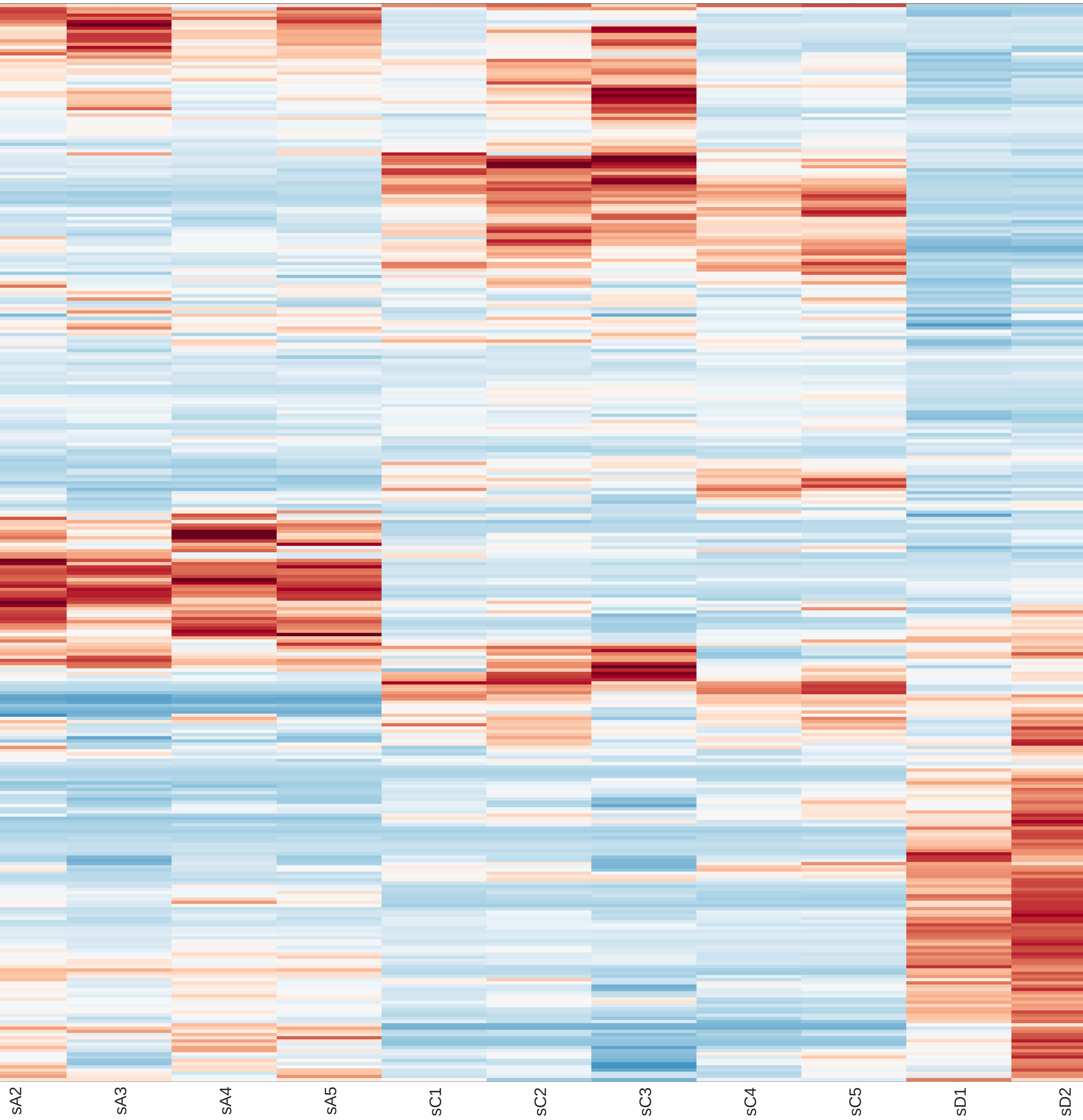

### clustergram.pdf

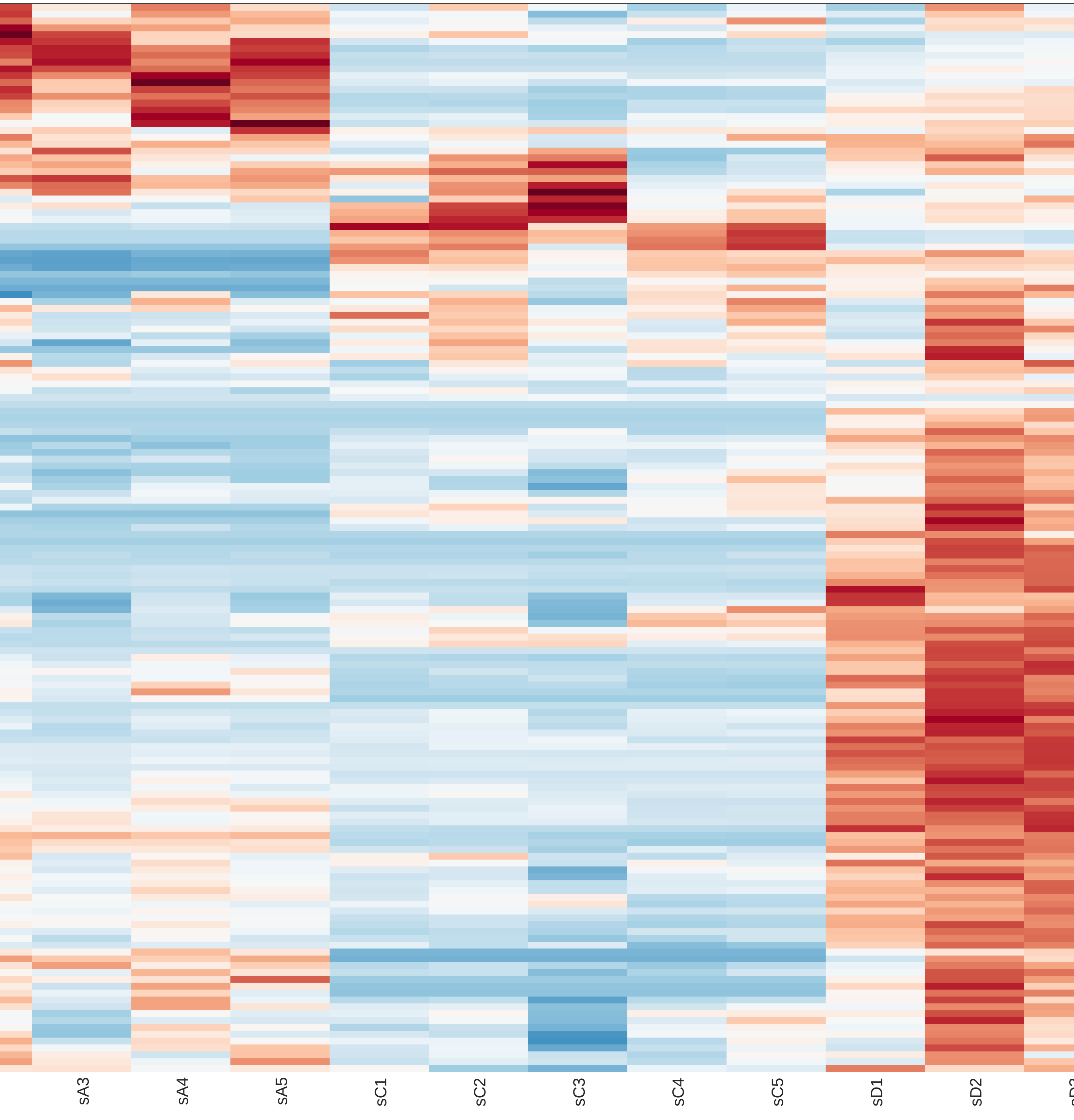

### clustergram.png

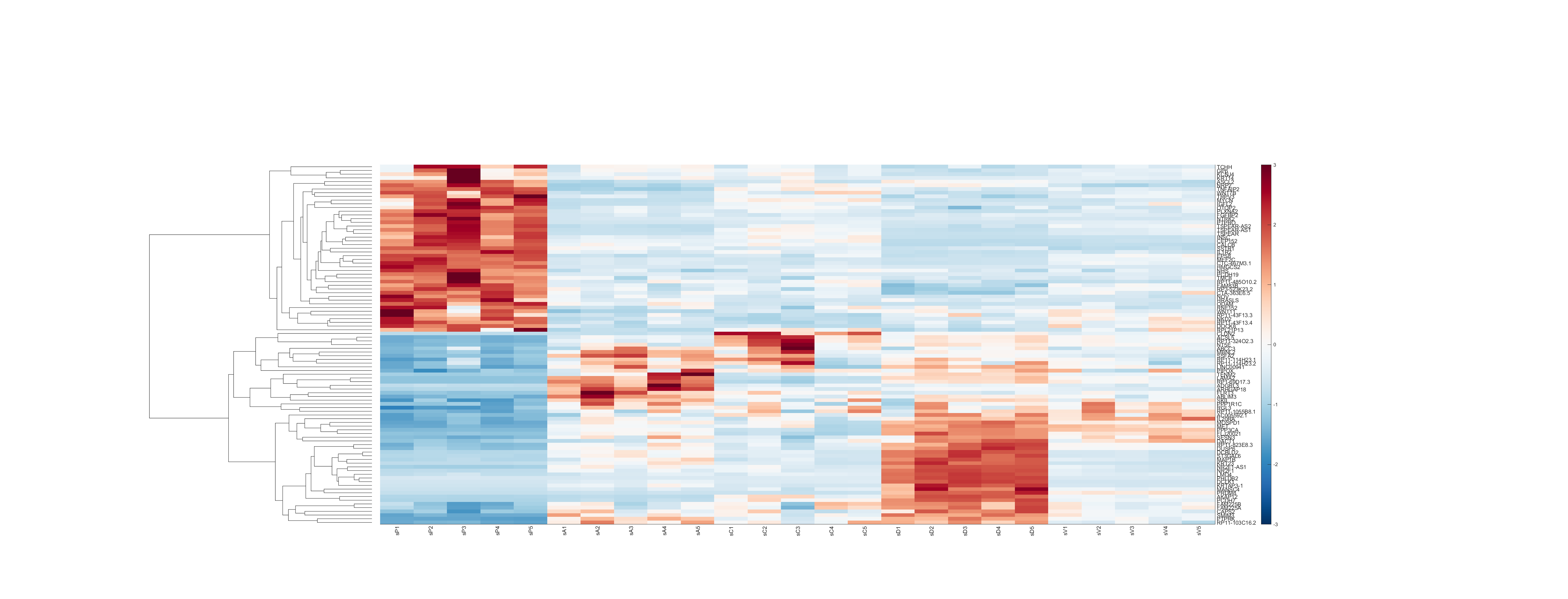

### clustergram.png

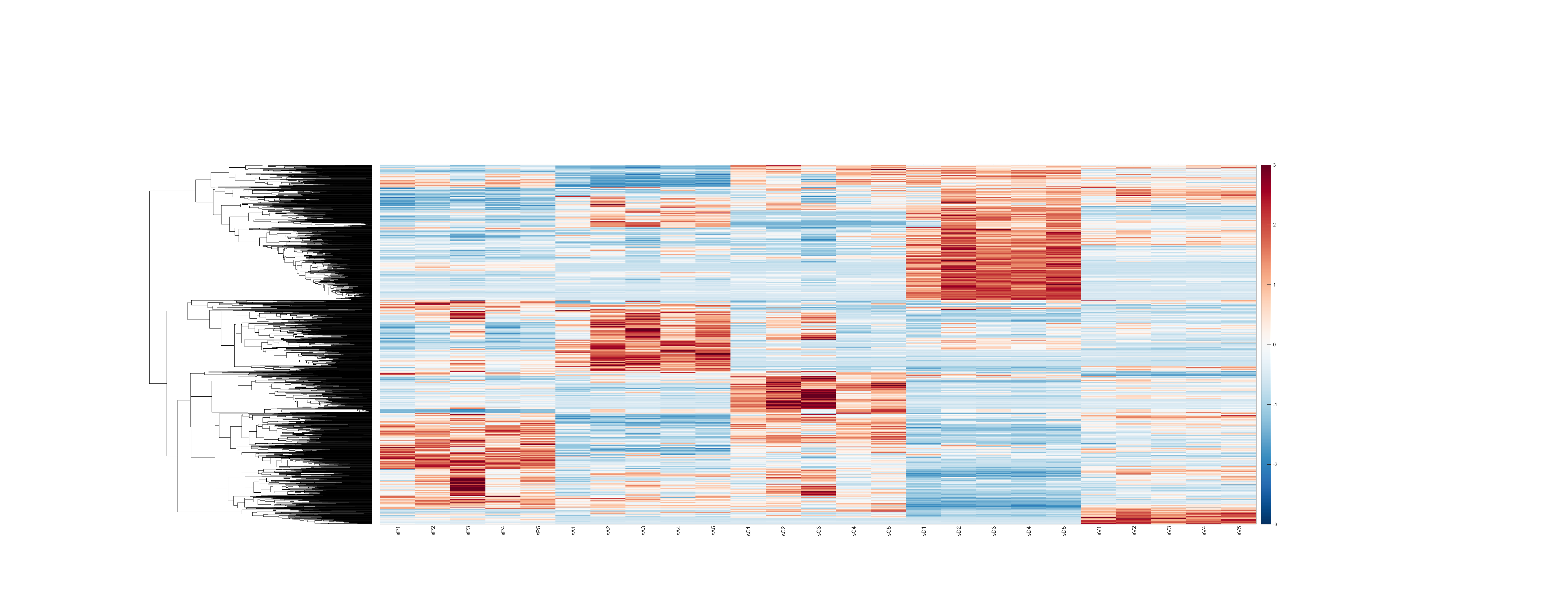

### clustergram.png

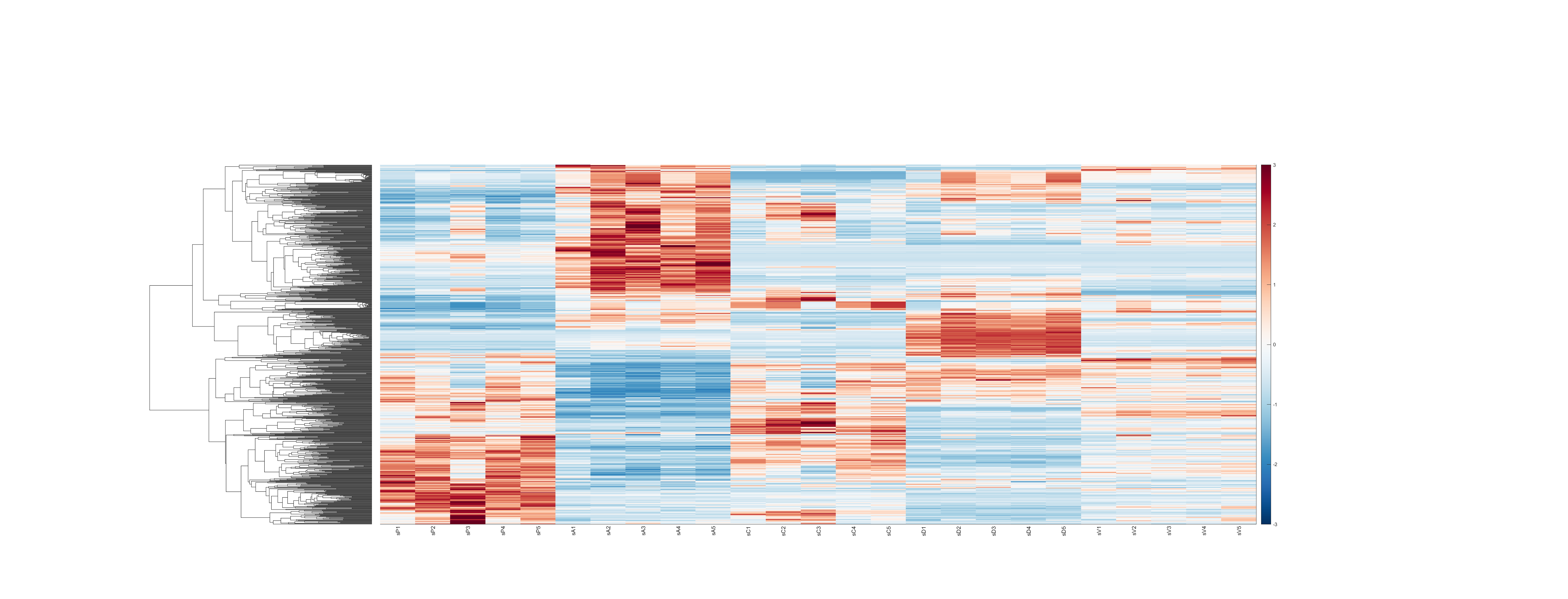

### clustergram.png

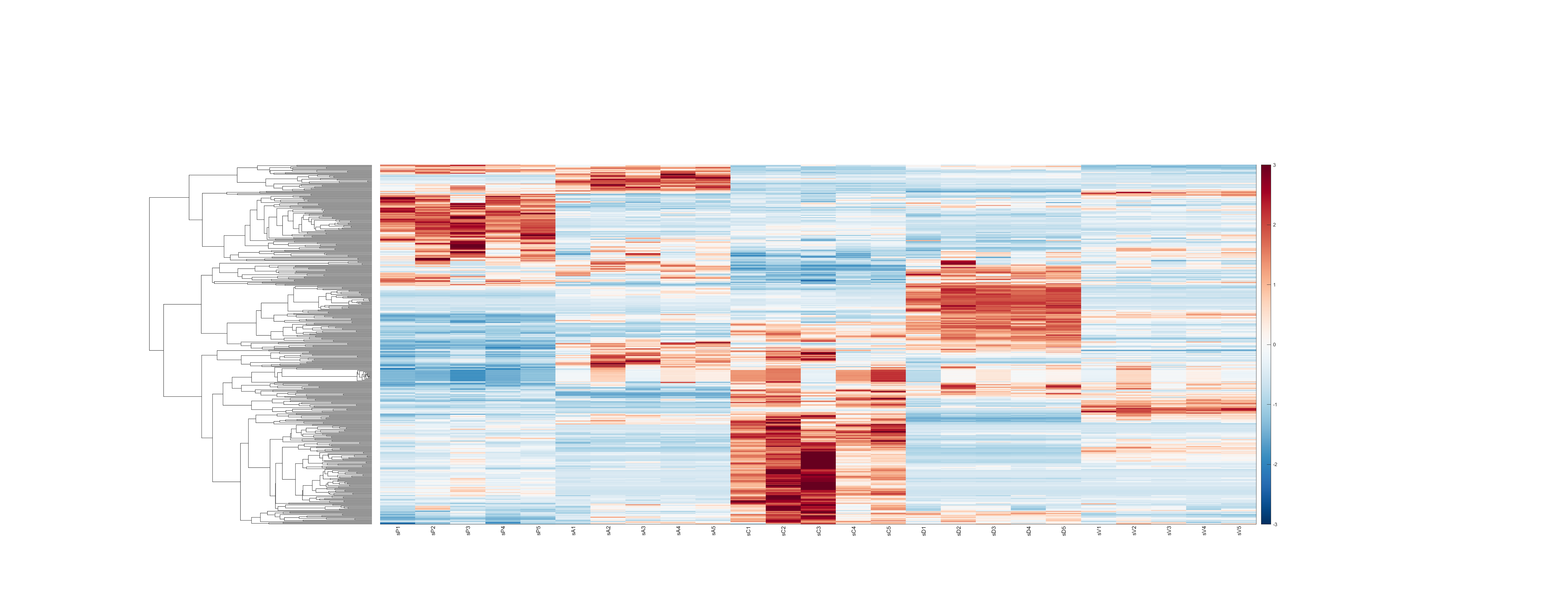

### clustergram.png

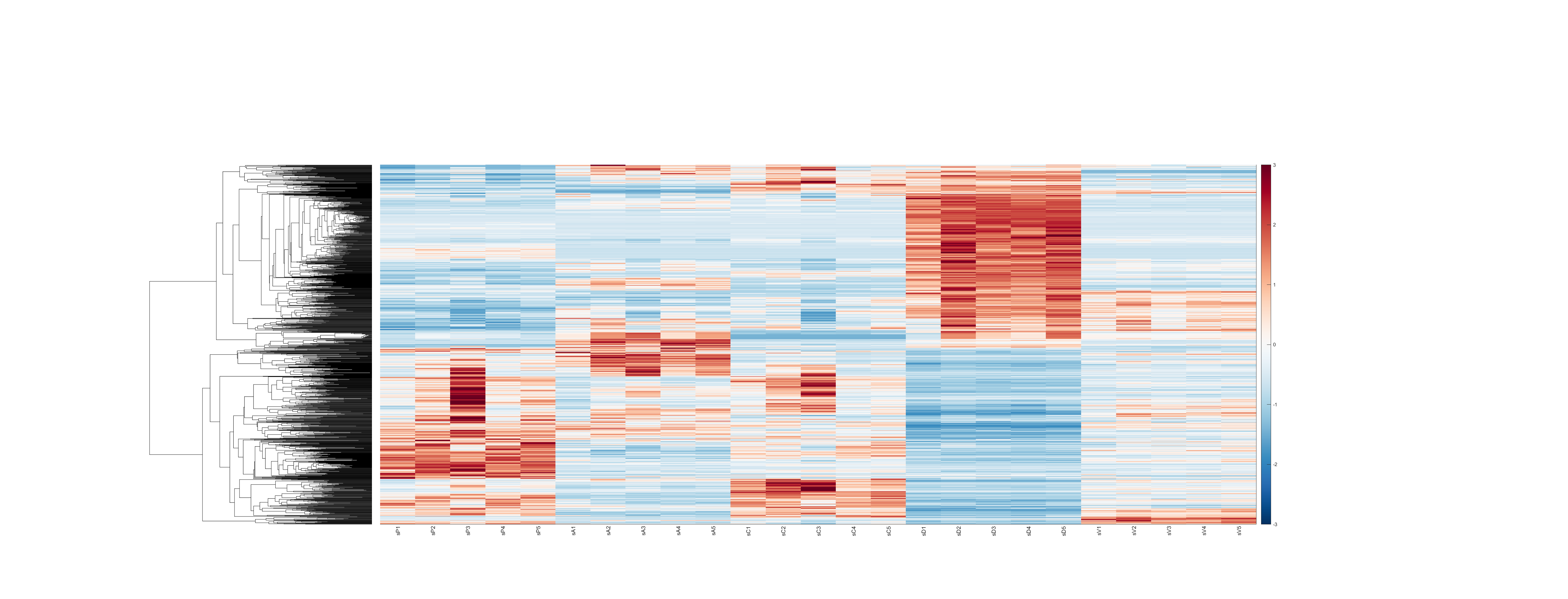

### clustergram.png

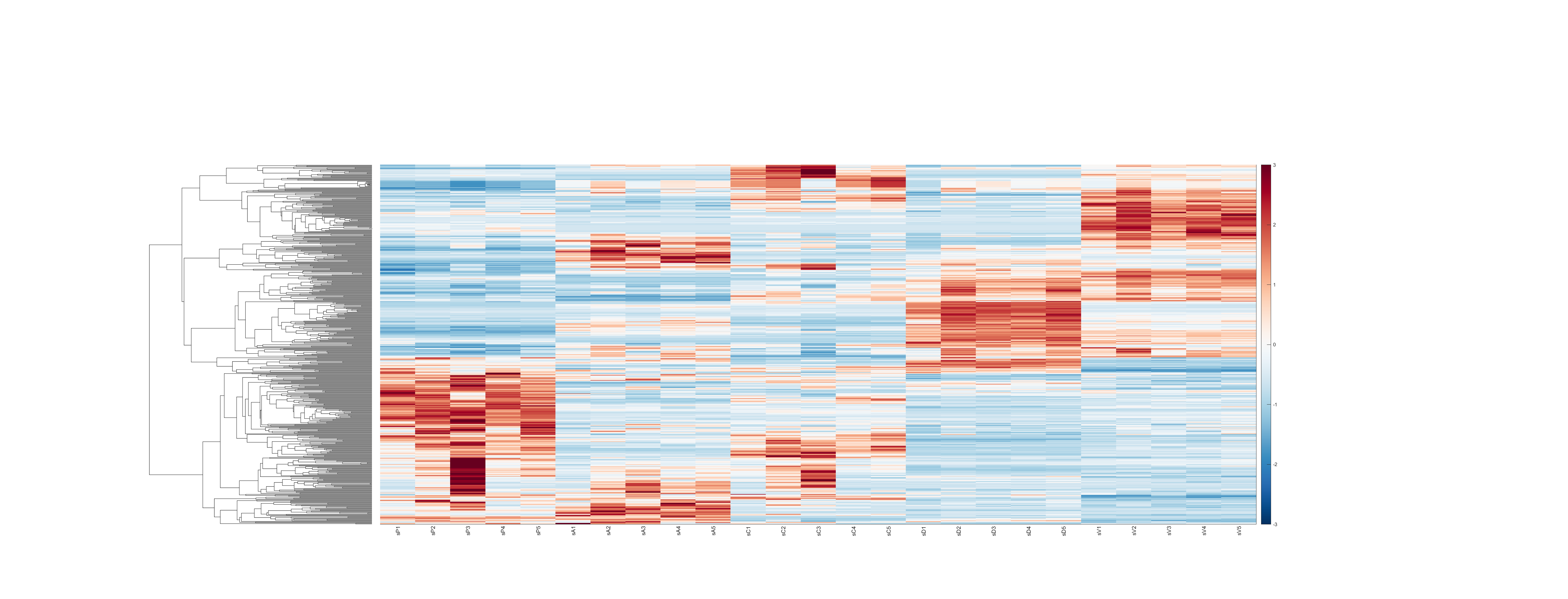

### clustergram.png

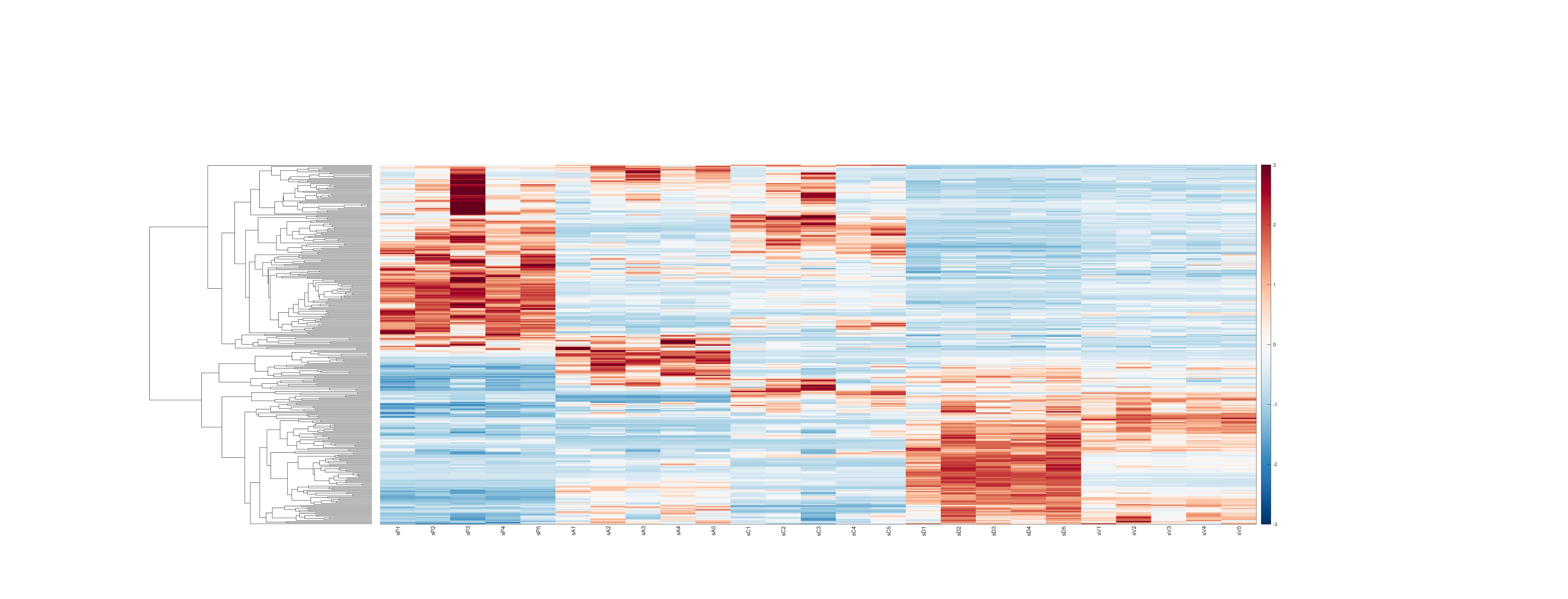

### clustergram.png

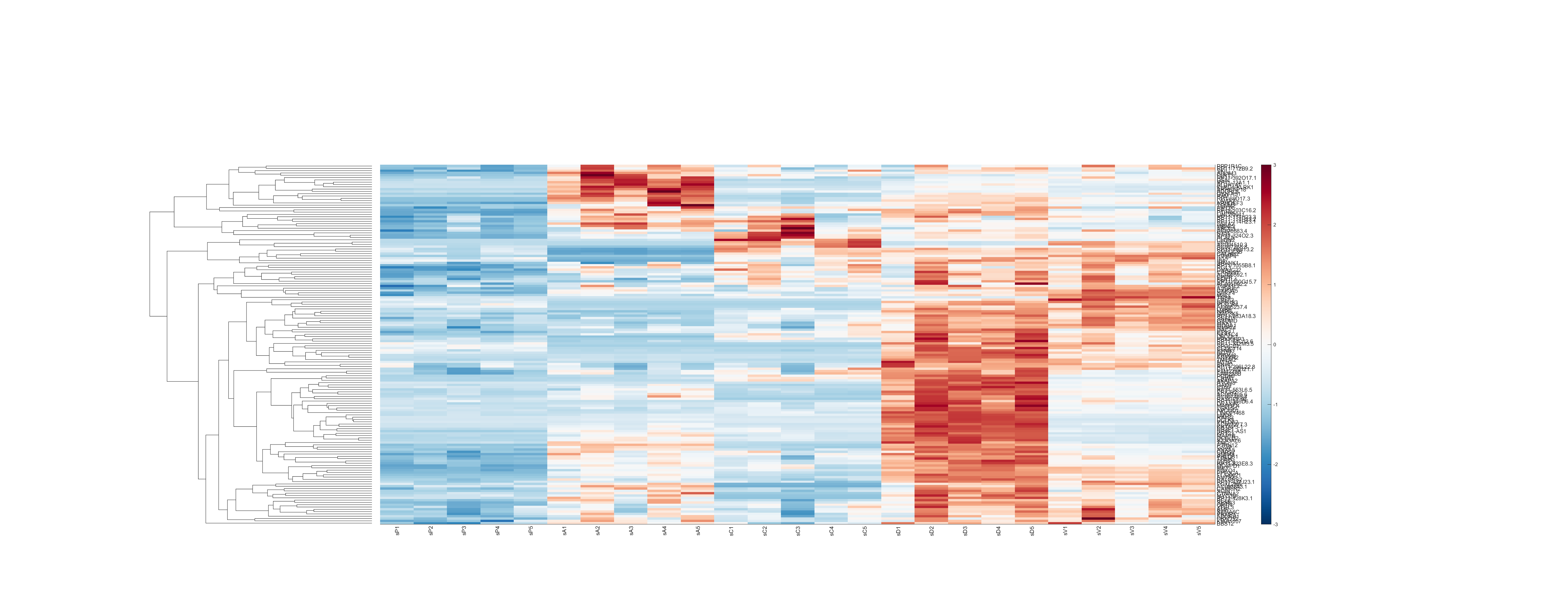

### clustergramDE.pdf

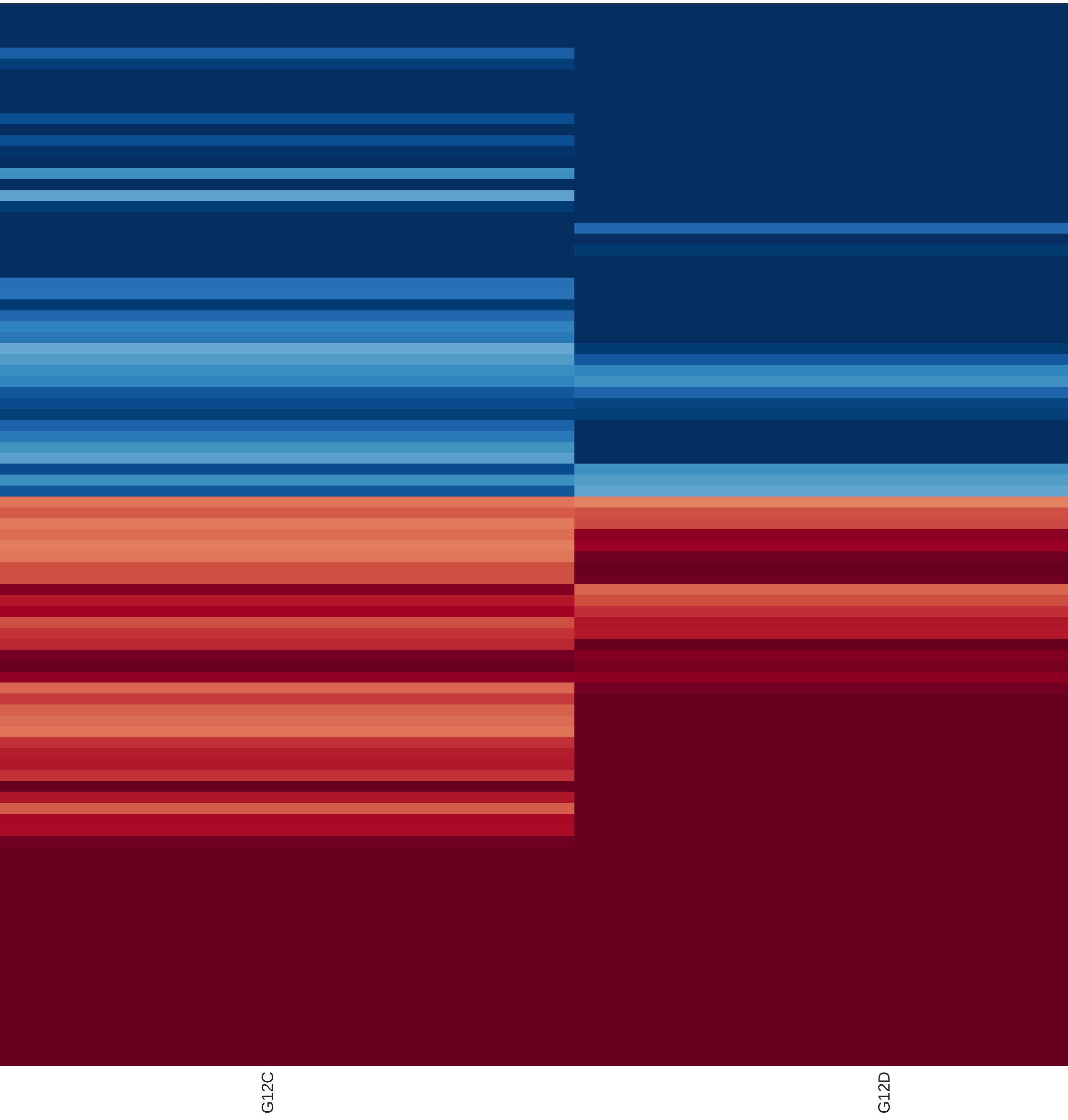

### clustergramDE.pdf

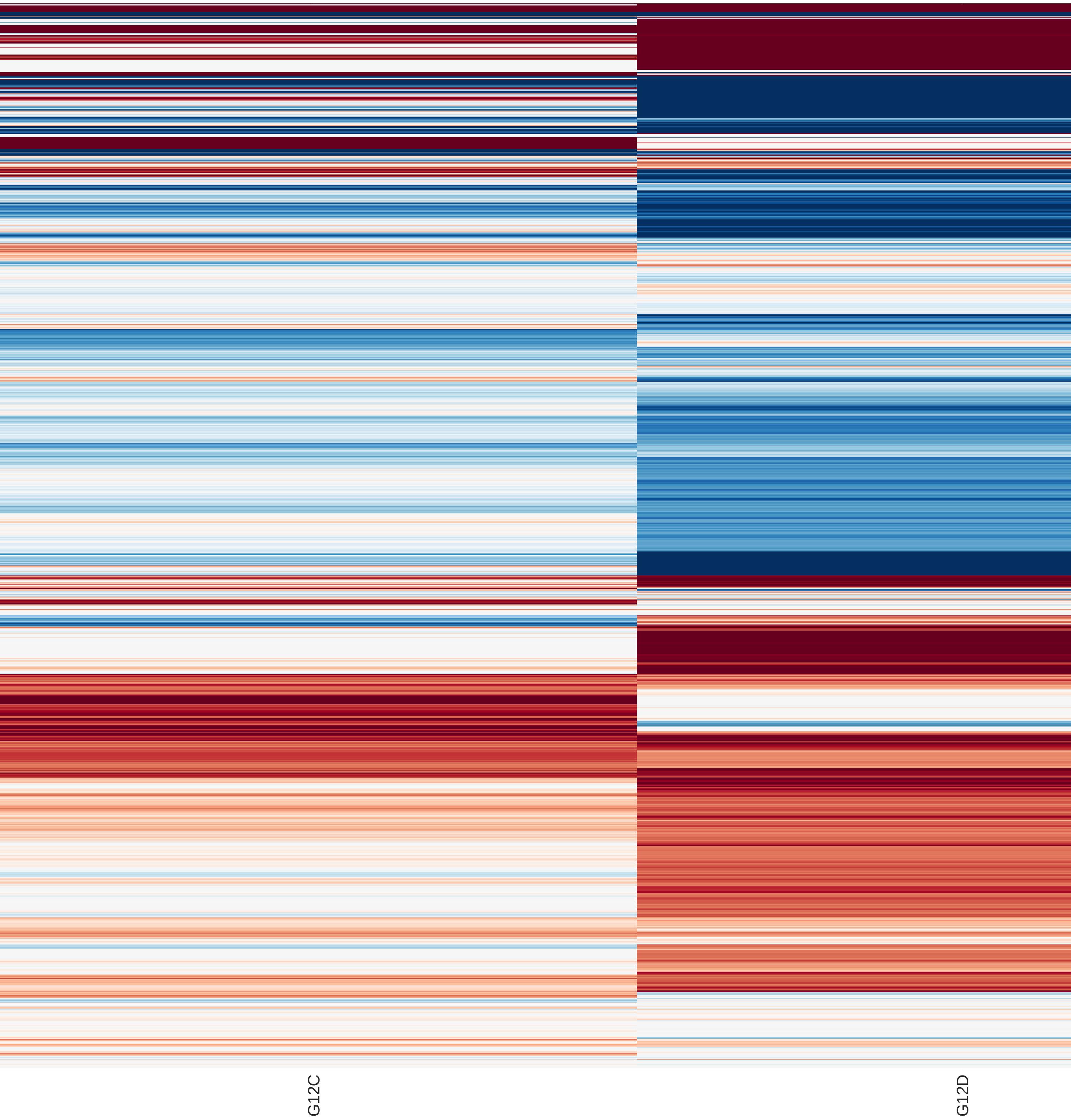

### clustergramDE.pdf

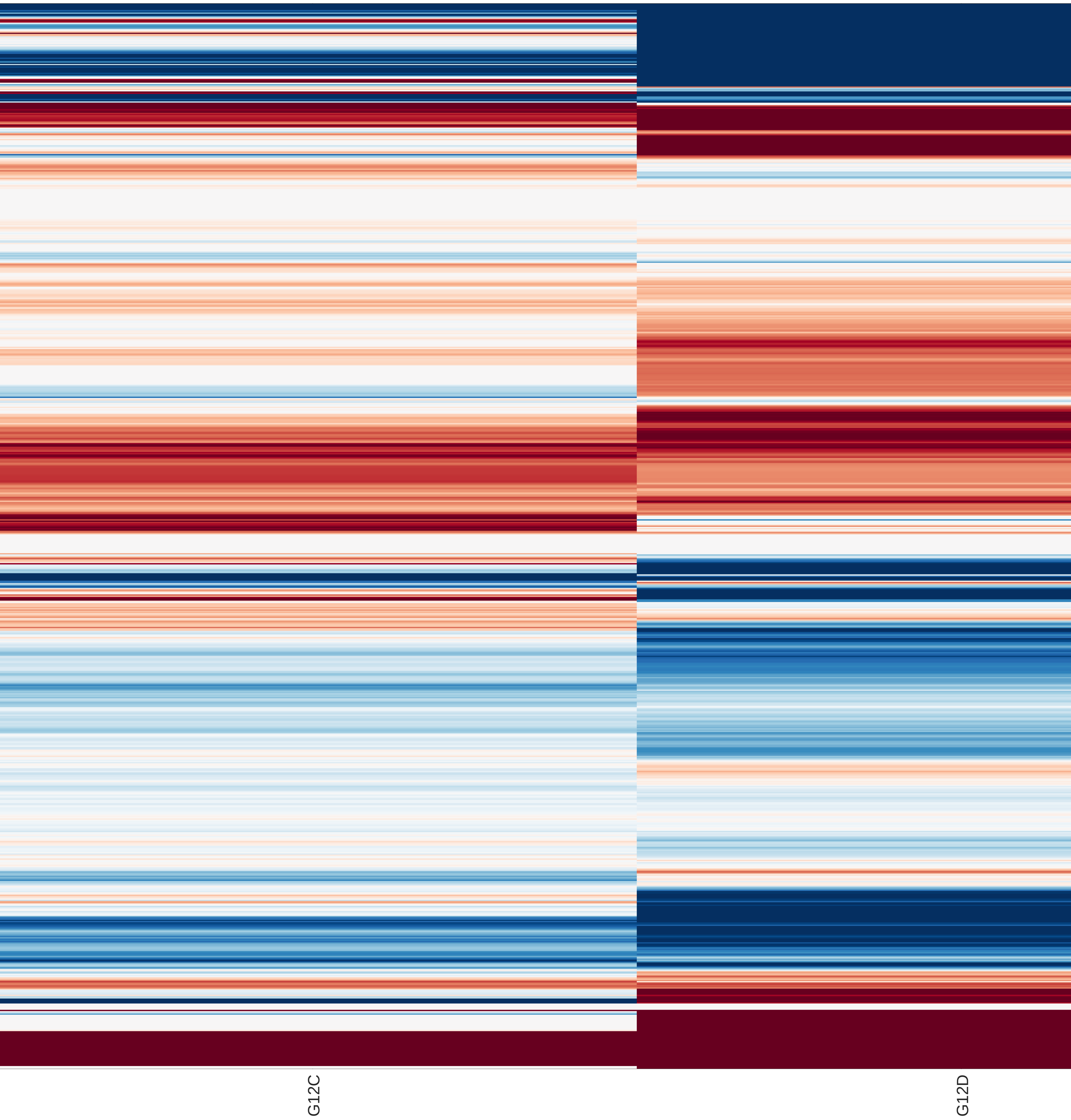

### clustergramDE.pdf

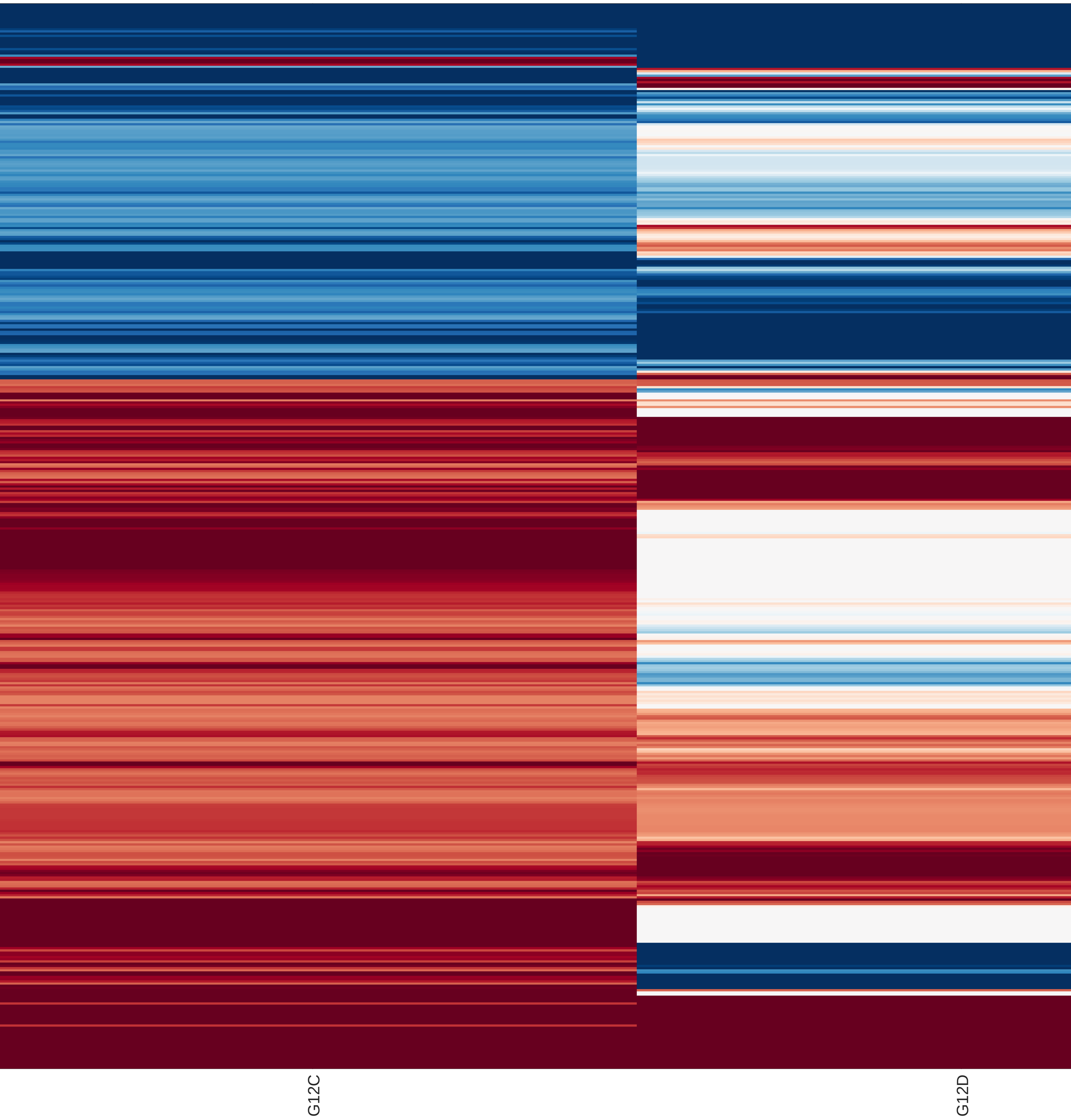

### clustergramDE.pdf

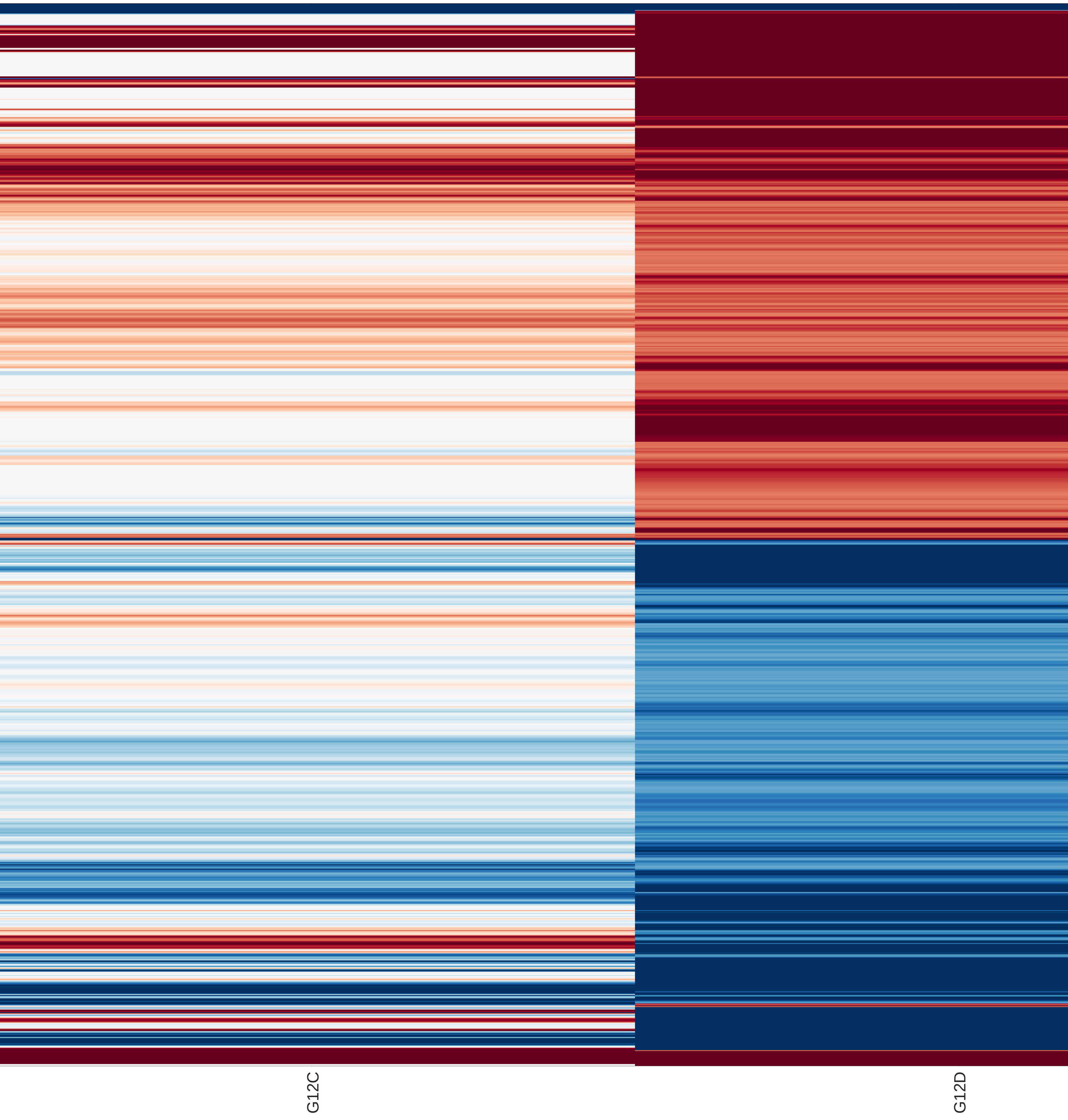

### clustergramDE.pdf

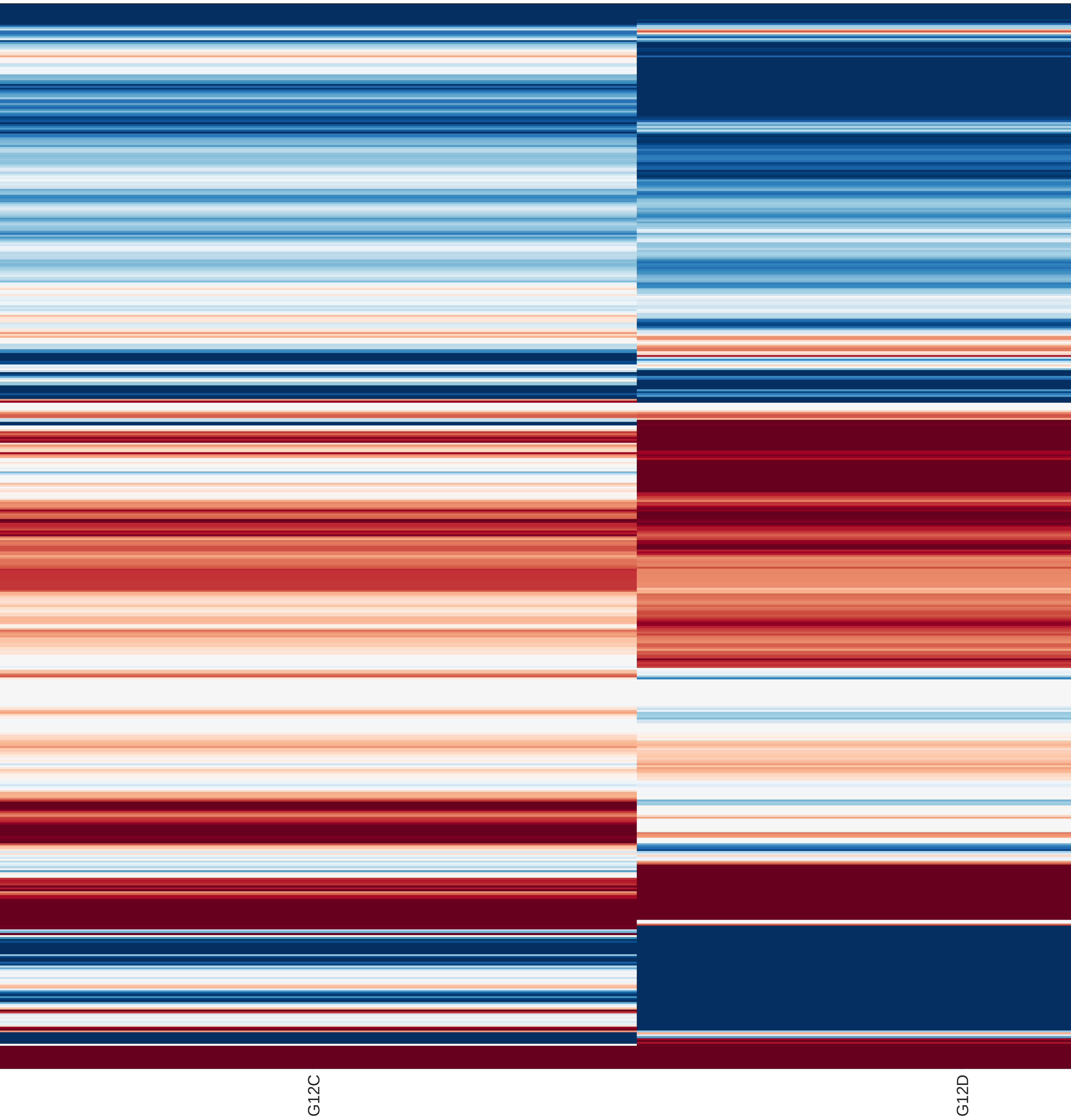

### clustergramDE.pdf

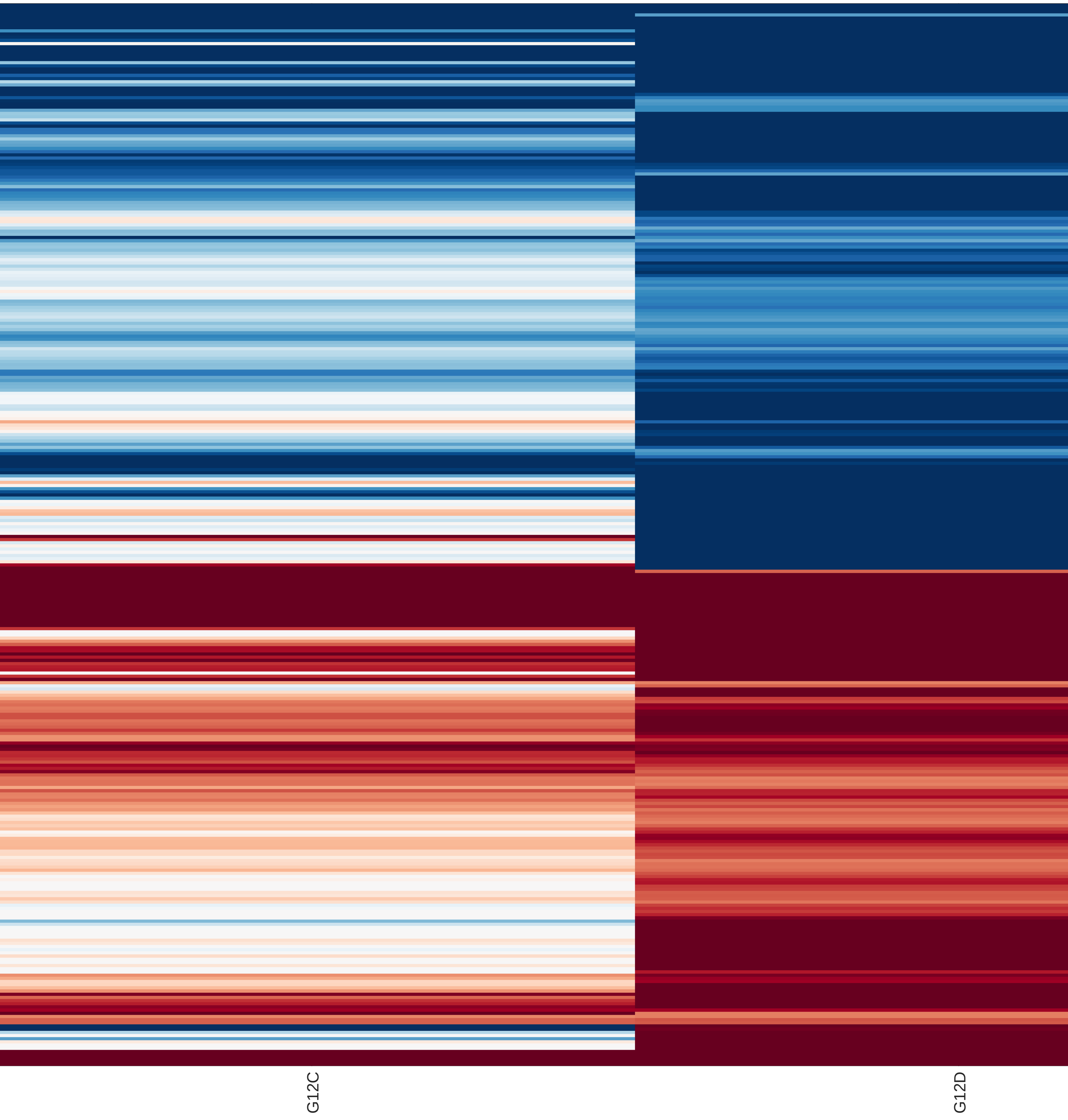

### clustergramDE.pdf

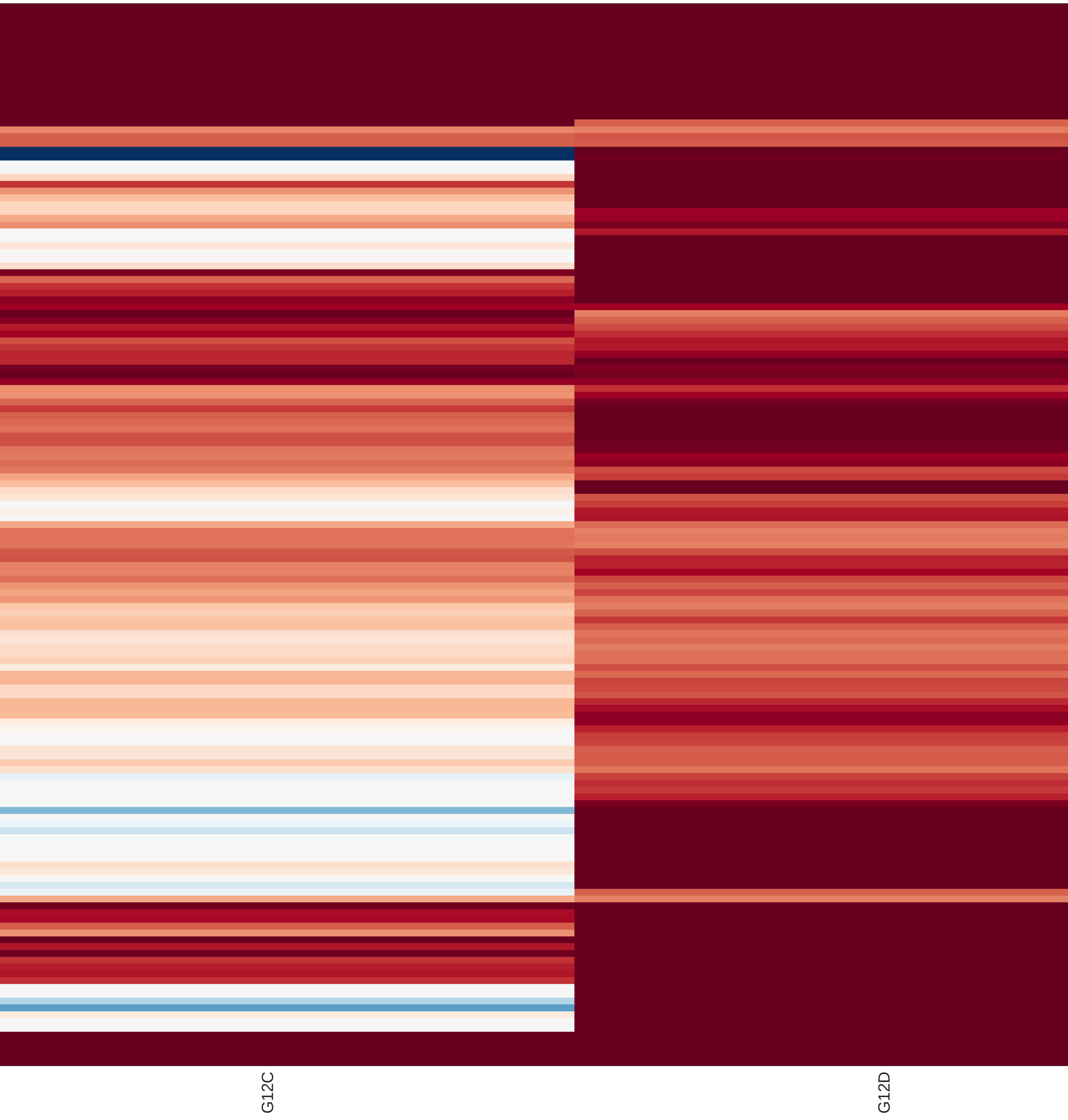

### clustergramDE.png

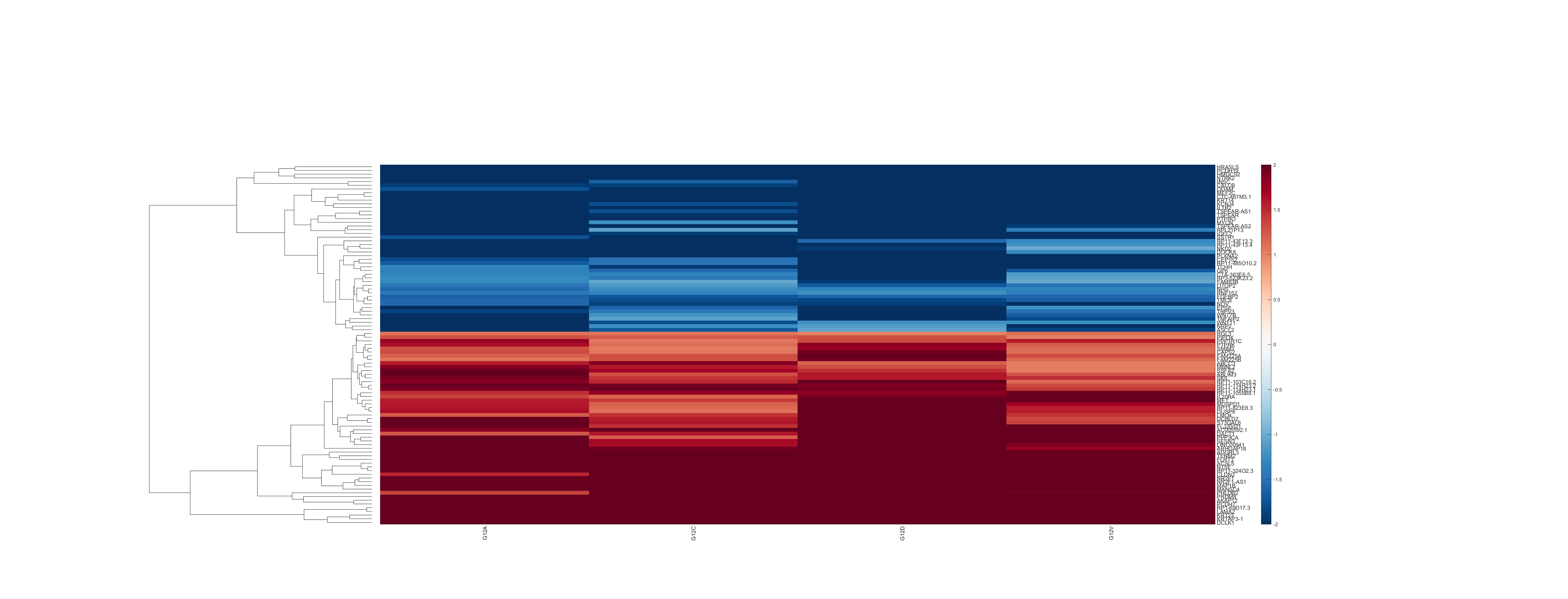

### clustergramDE.png

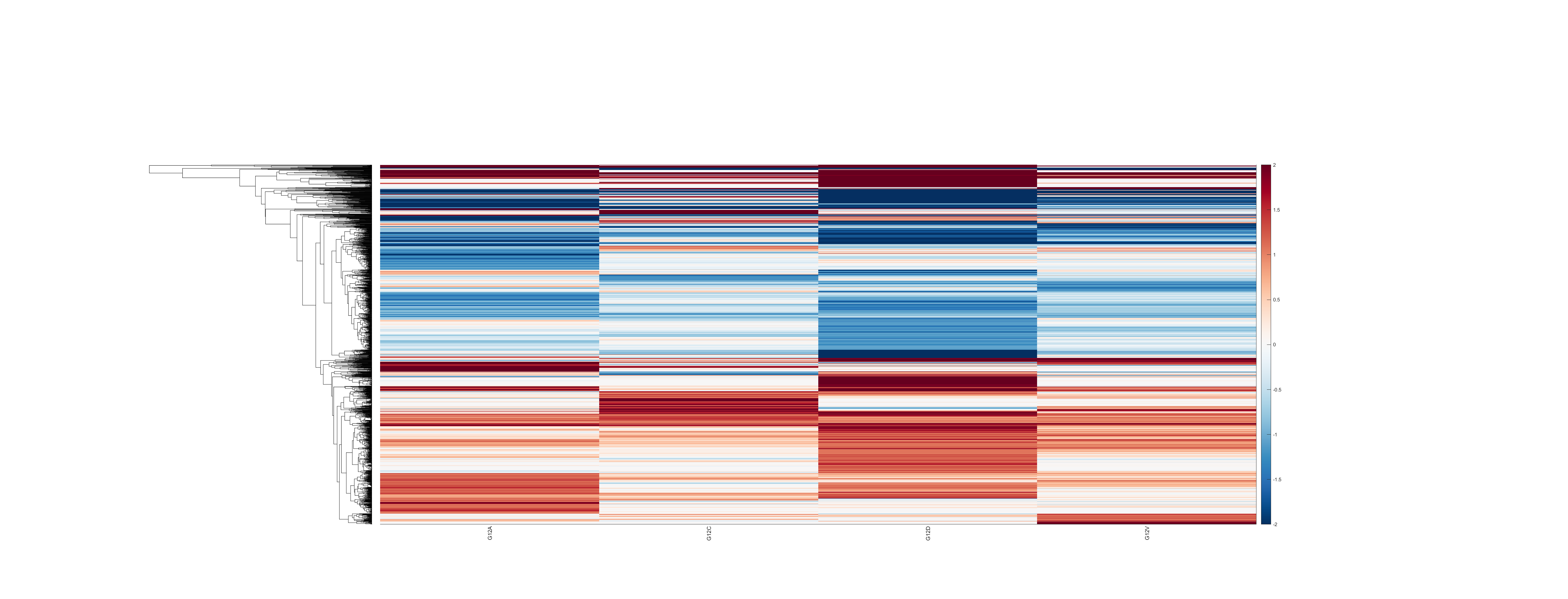

### clustergramDE.png

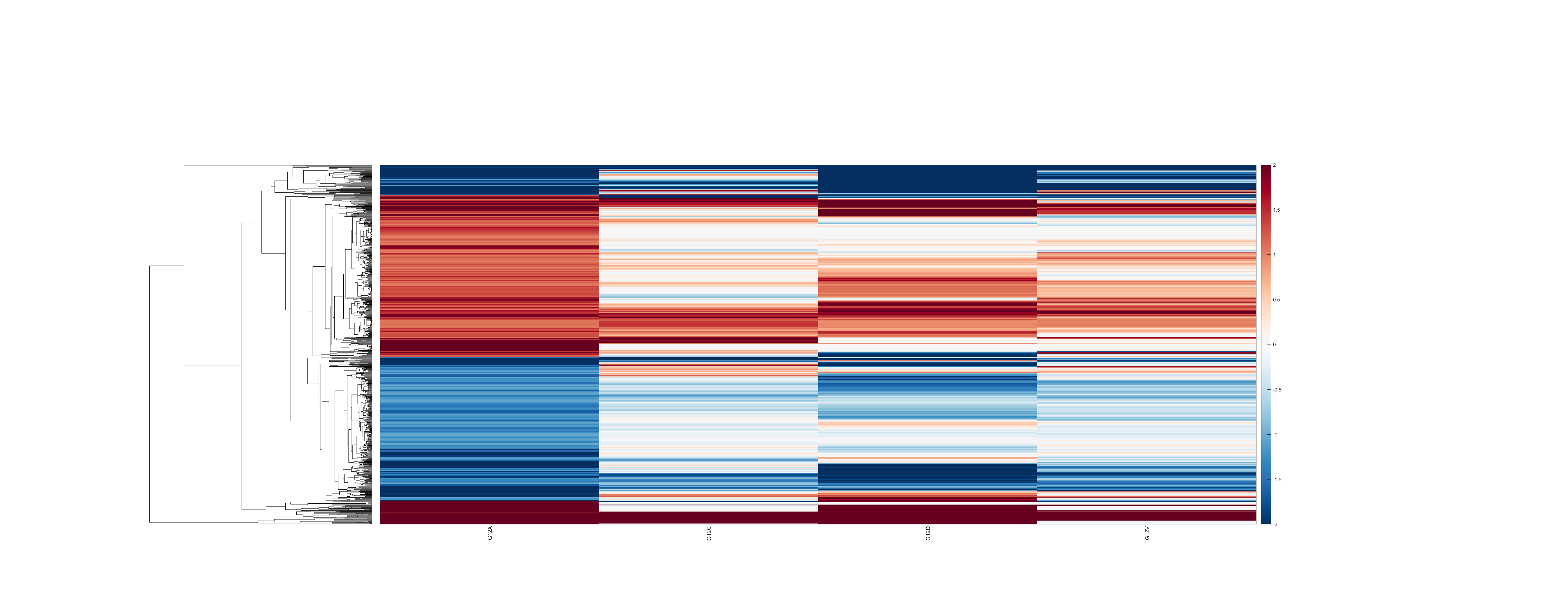

### clustergramDE.png

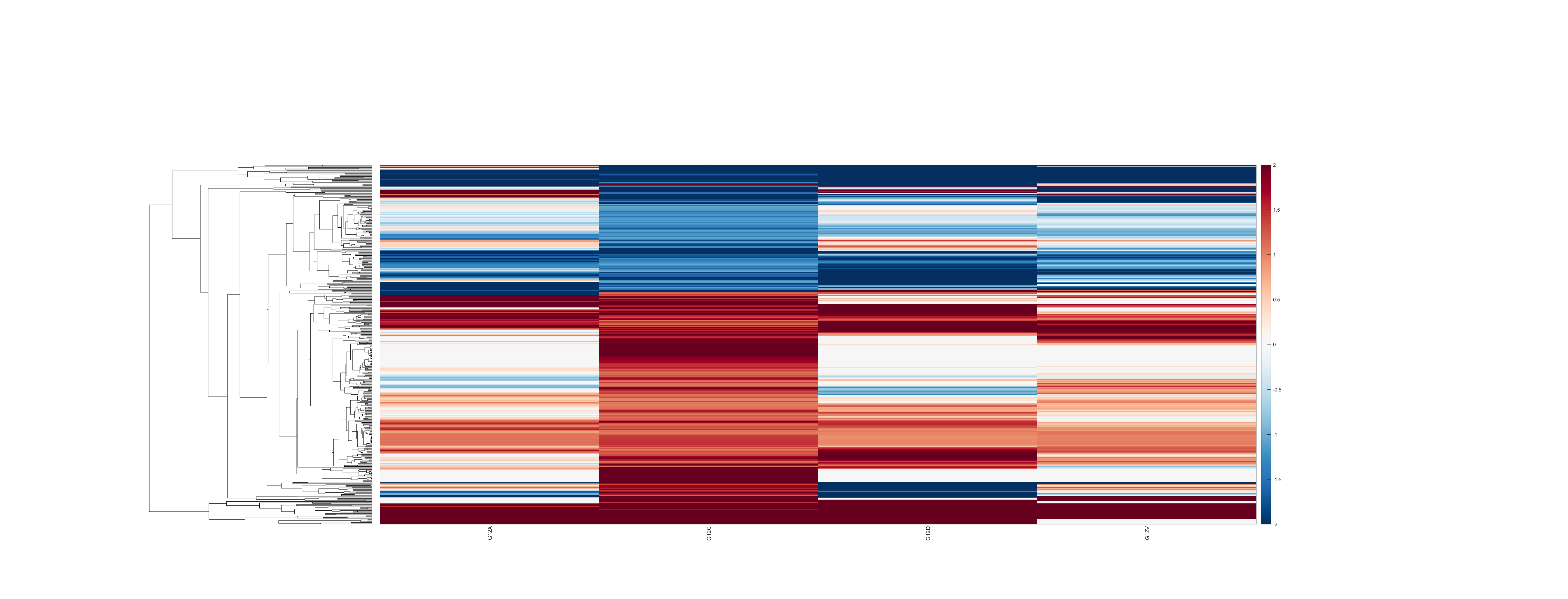

### clustergramDE.png

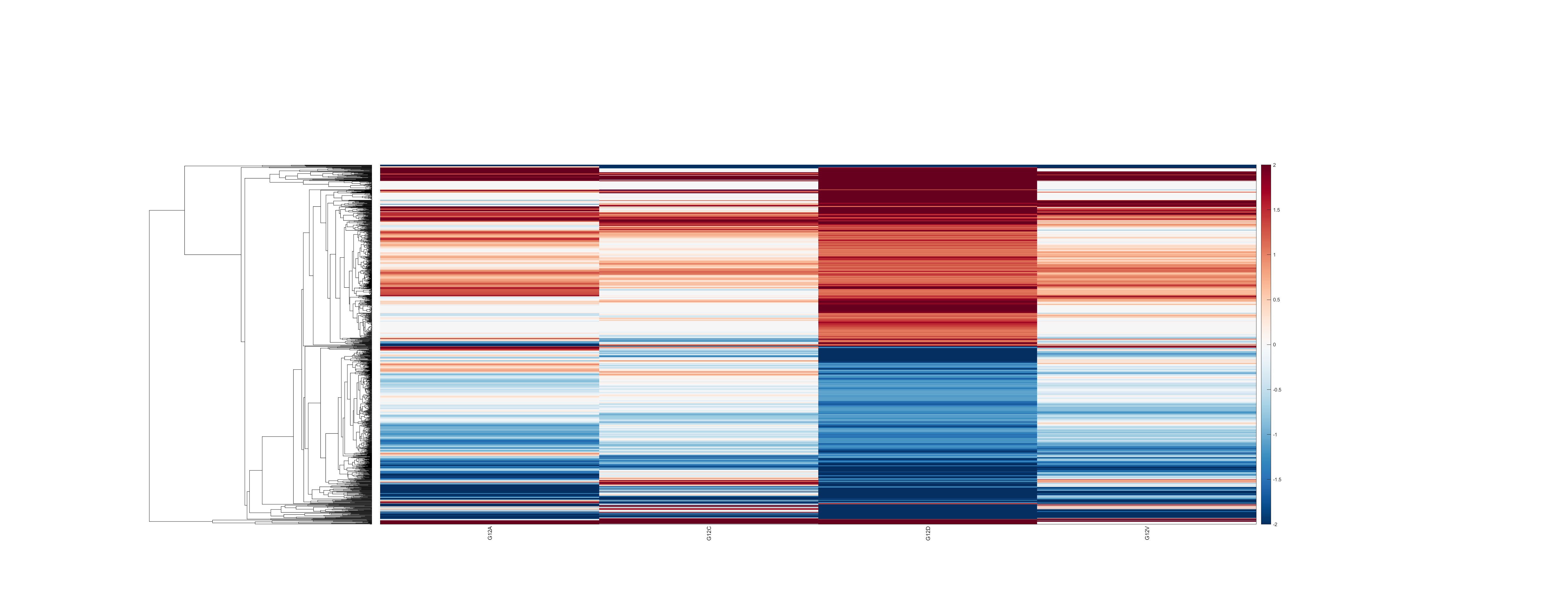

### clustergramDE.png

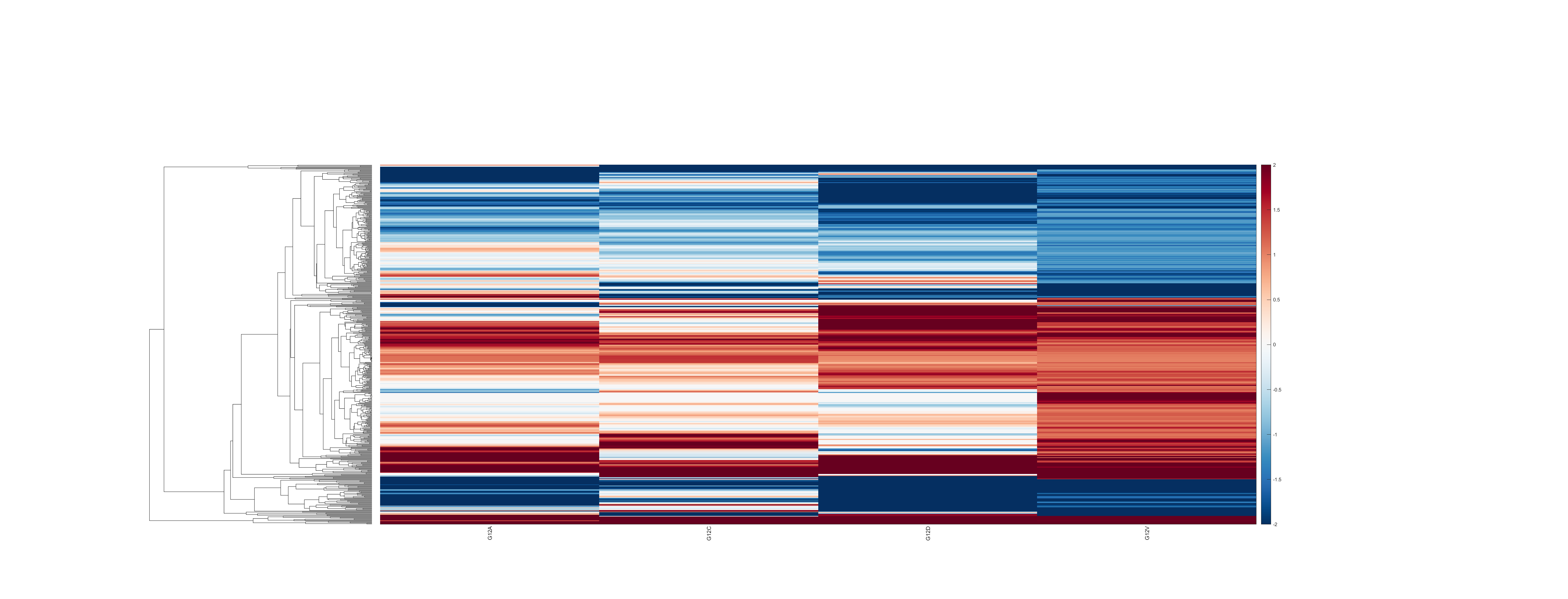

### clustergramDE.png

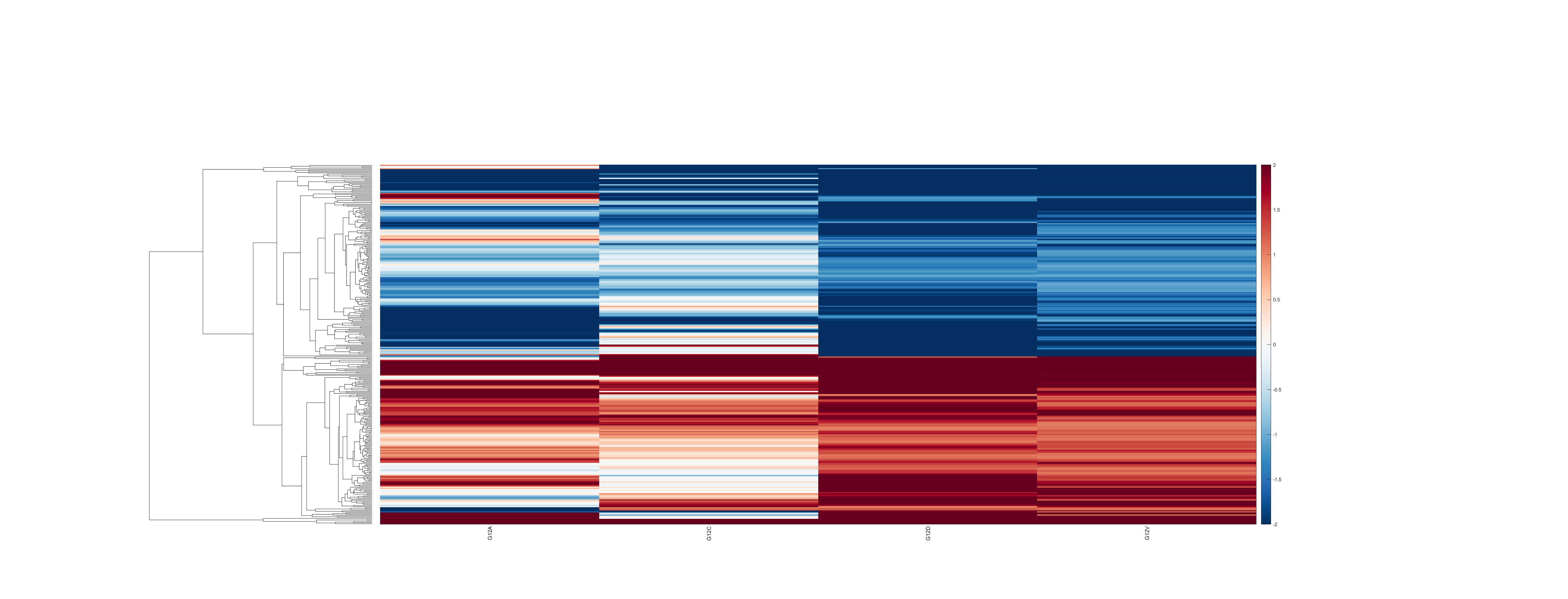

### clustergramDE.png

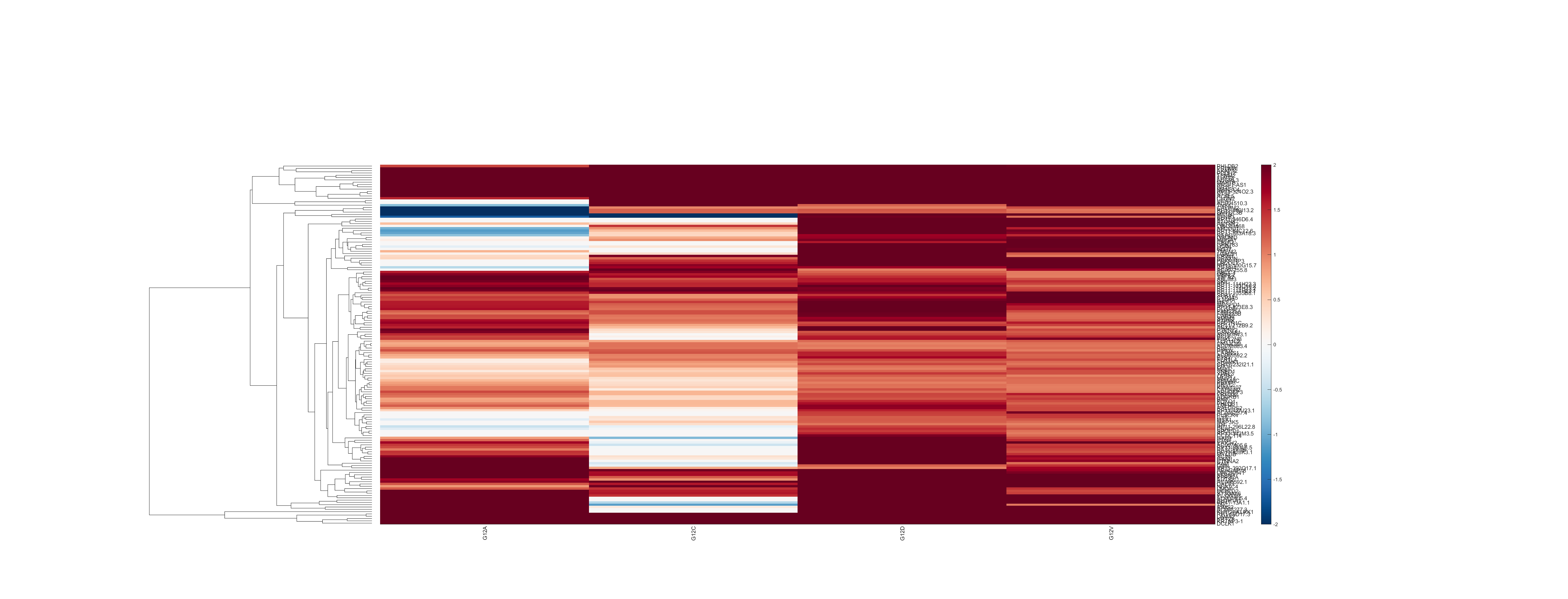

### hsa00250_cnt.png

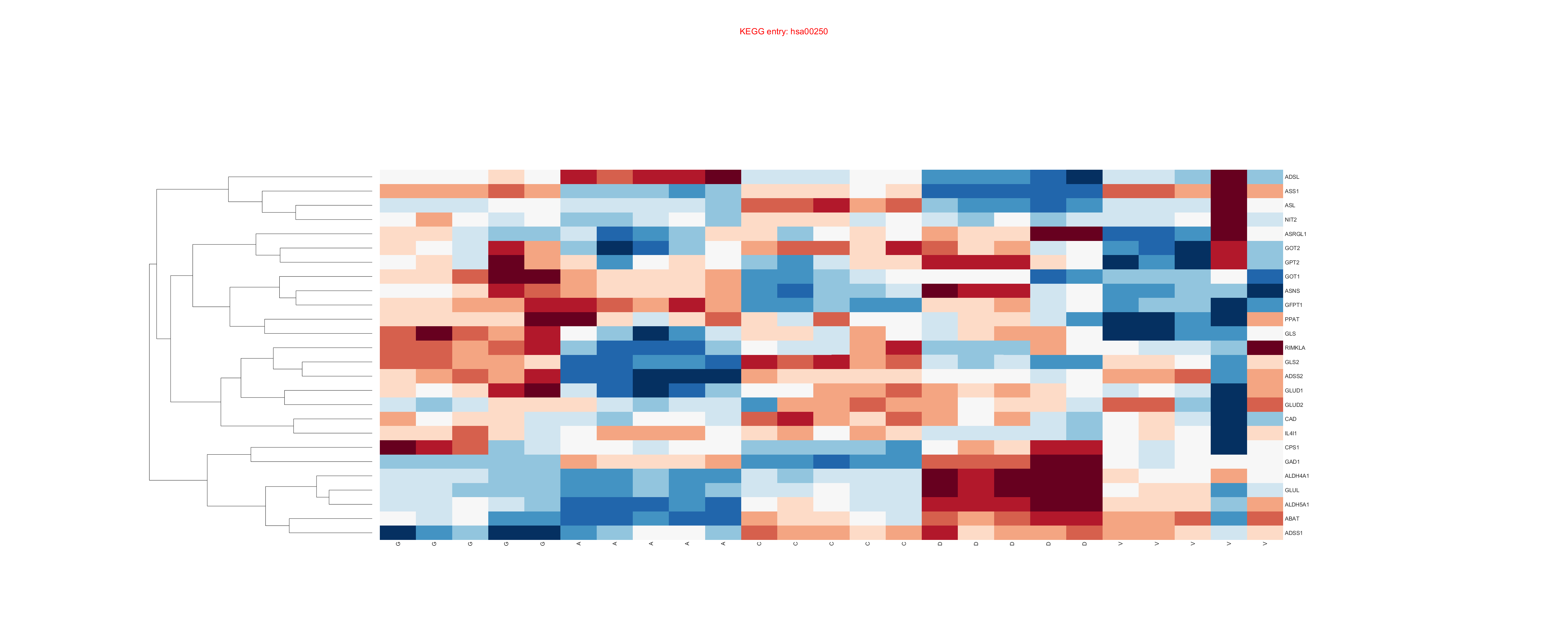

### hsa00250_fld.png

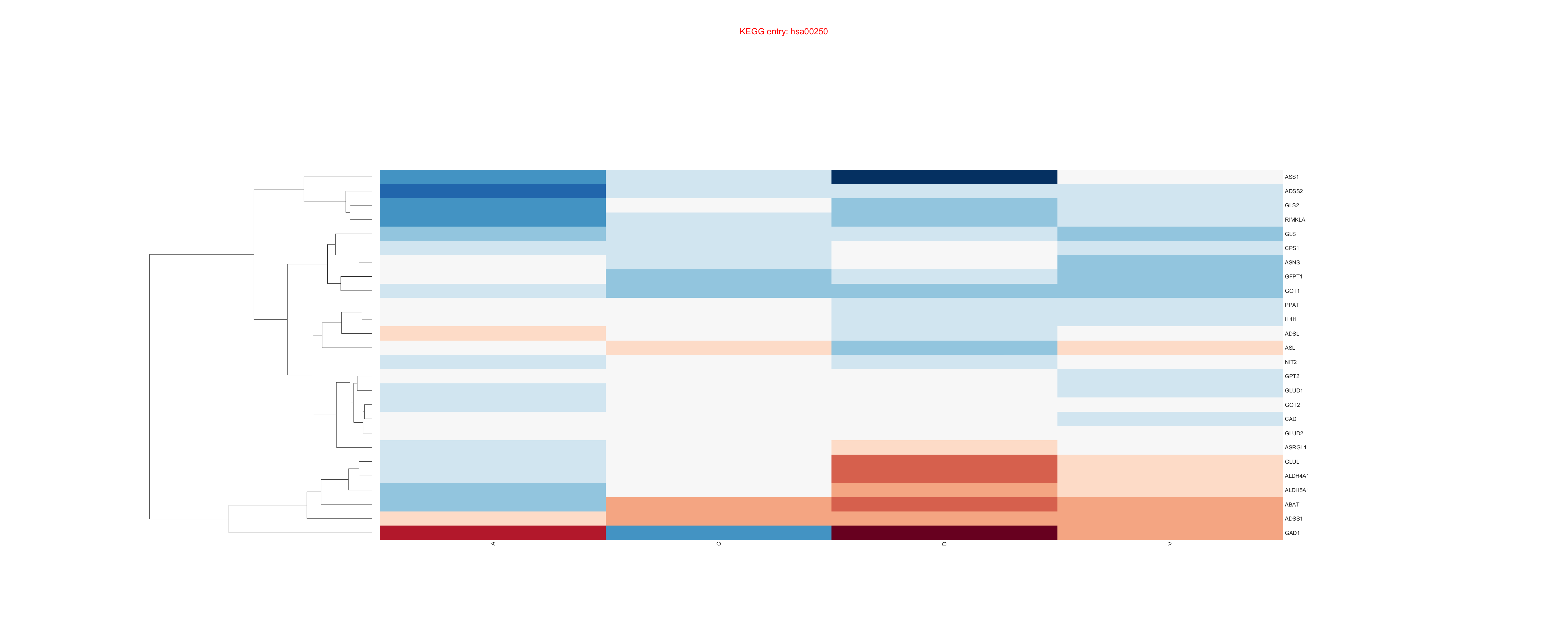

### hsa00250comparison_fld.png

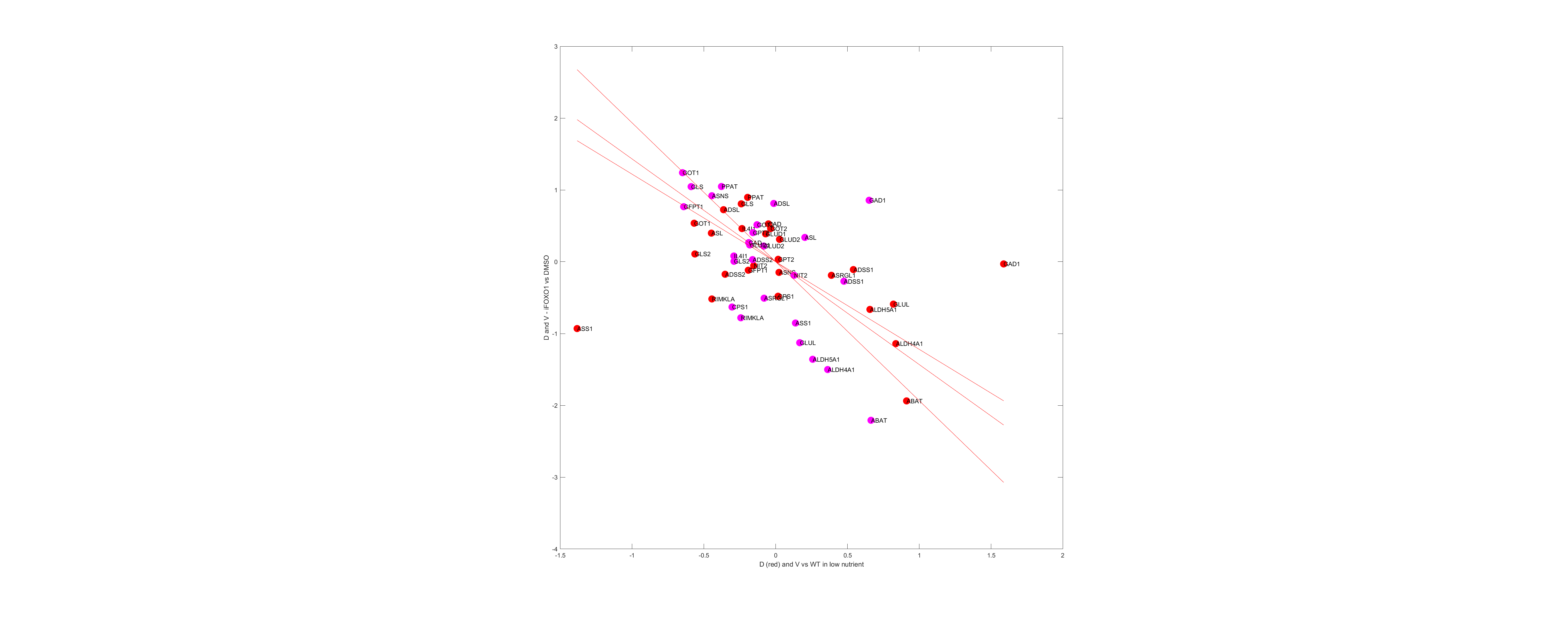

### hsa002501_cnt.png

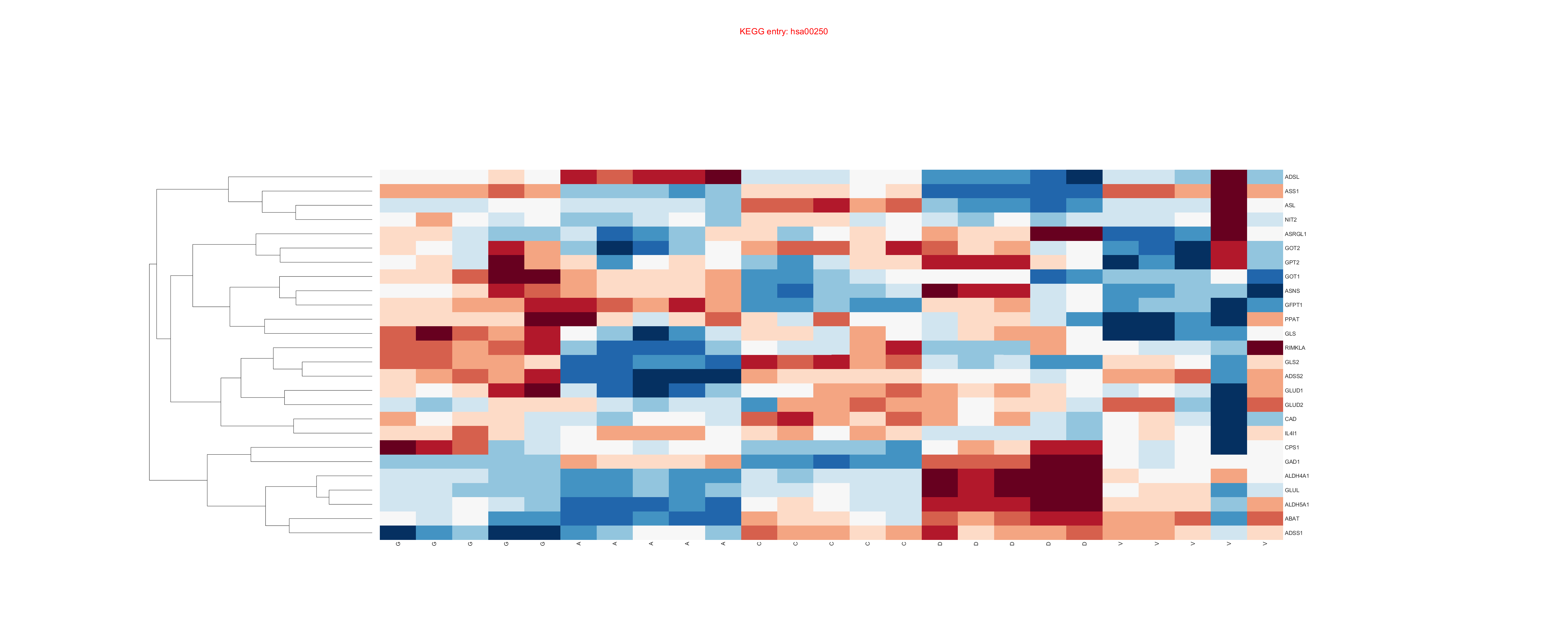

### hsa002501_fld.png

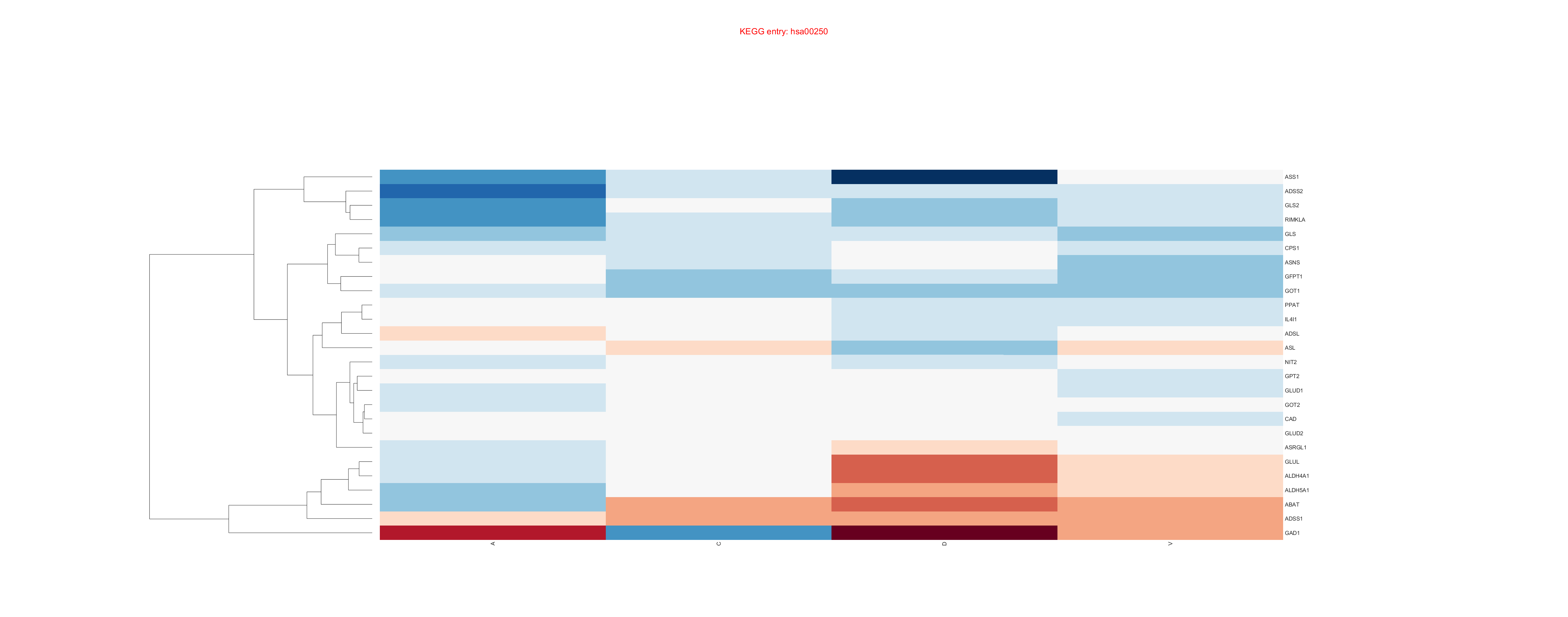

### pca_biplot.pdf

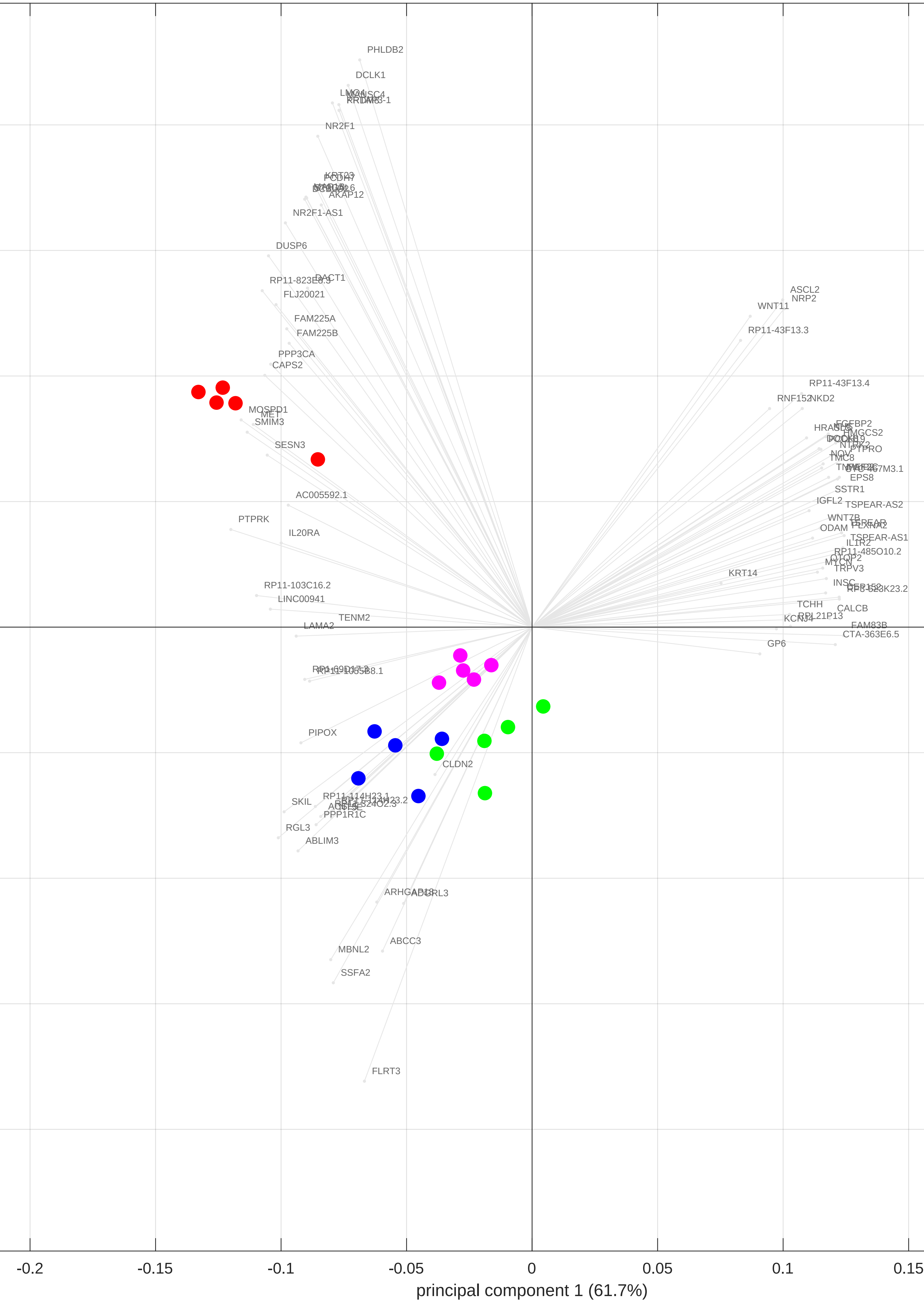

### pca_biplot.pdf

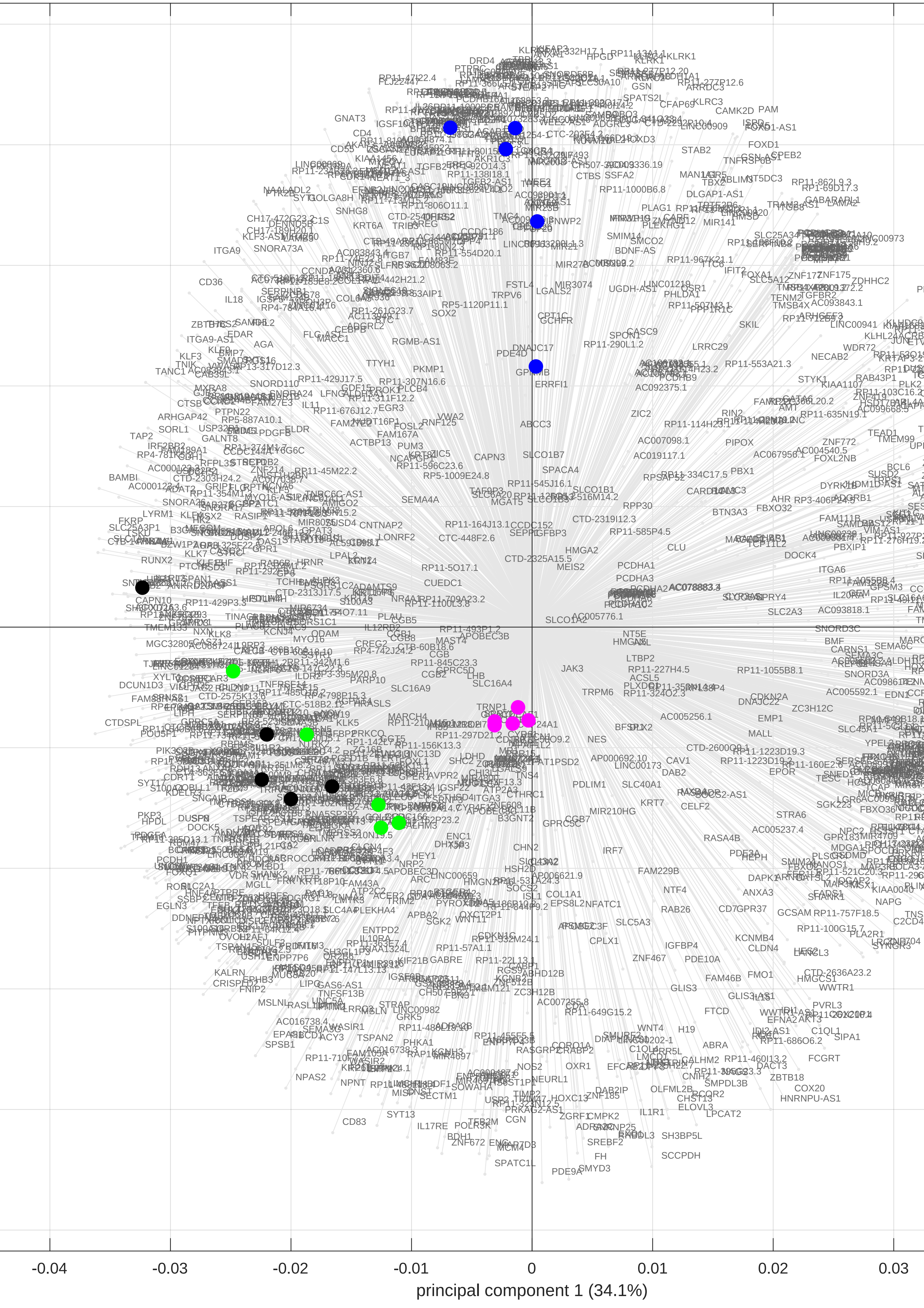

### pca_biplot.pdf

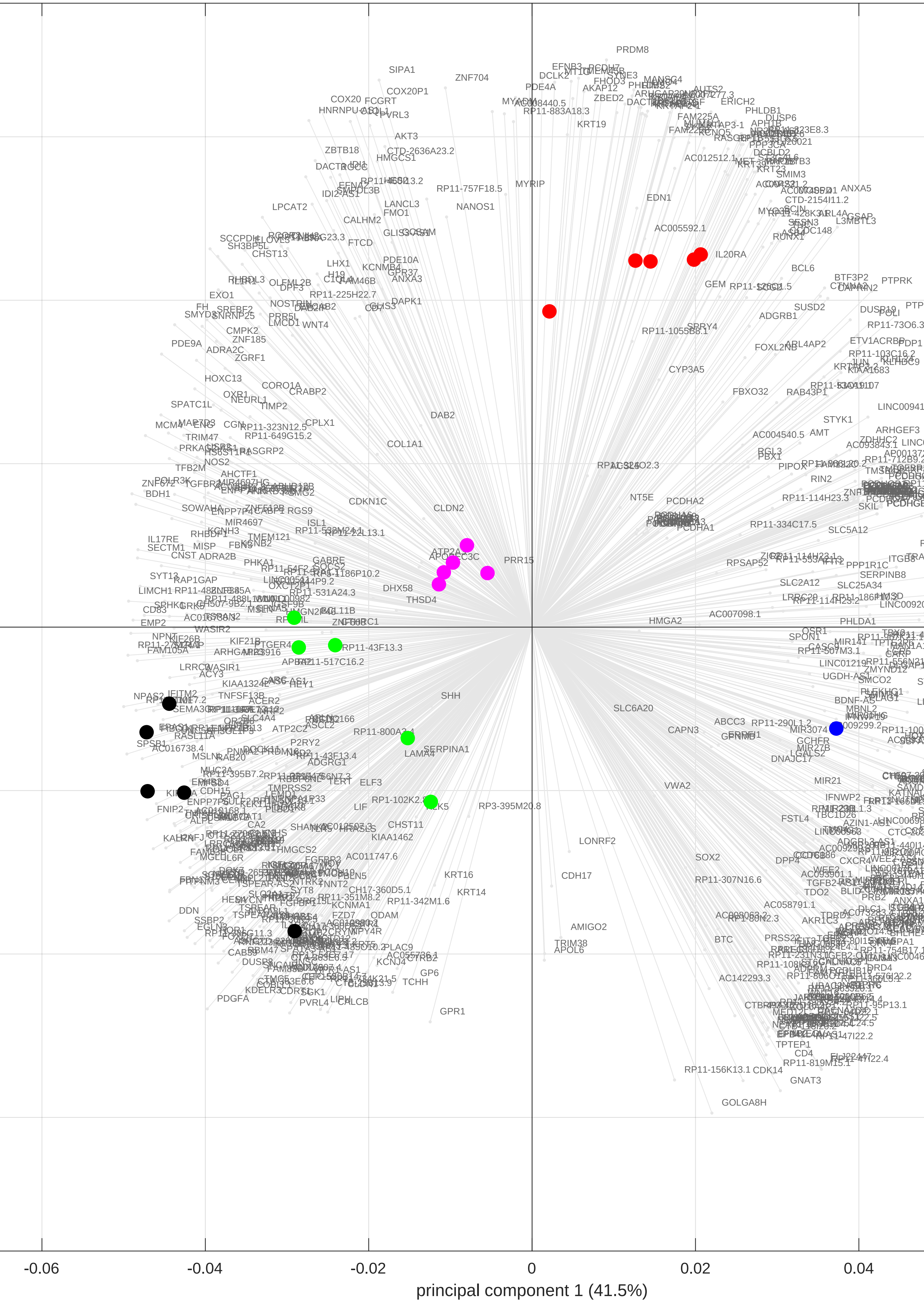

### pca_biplot.pdf

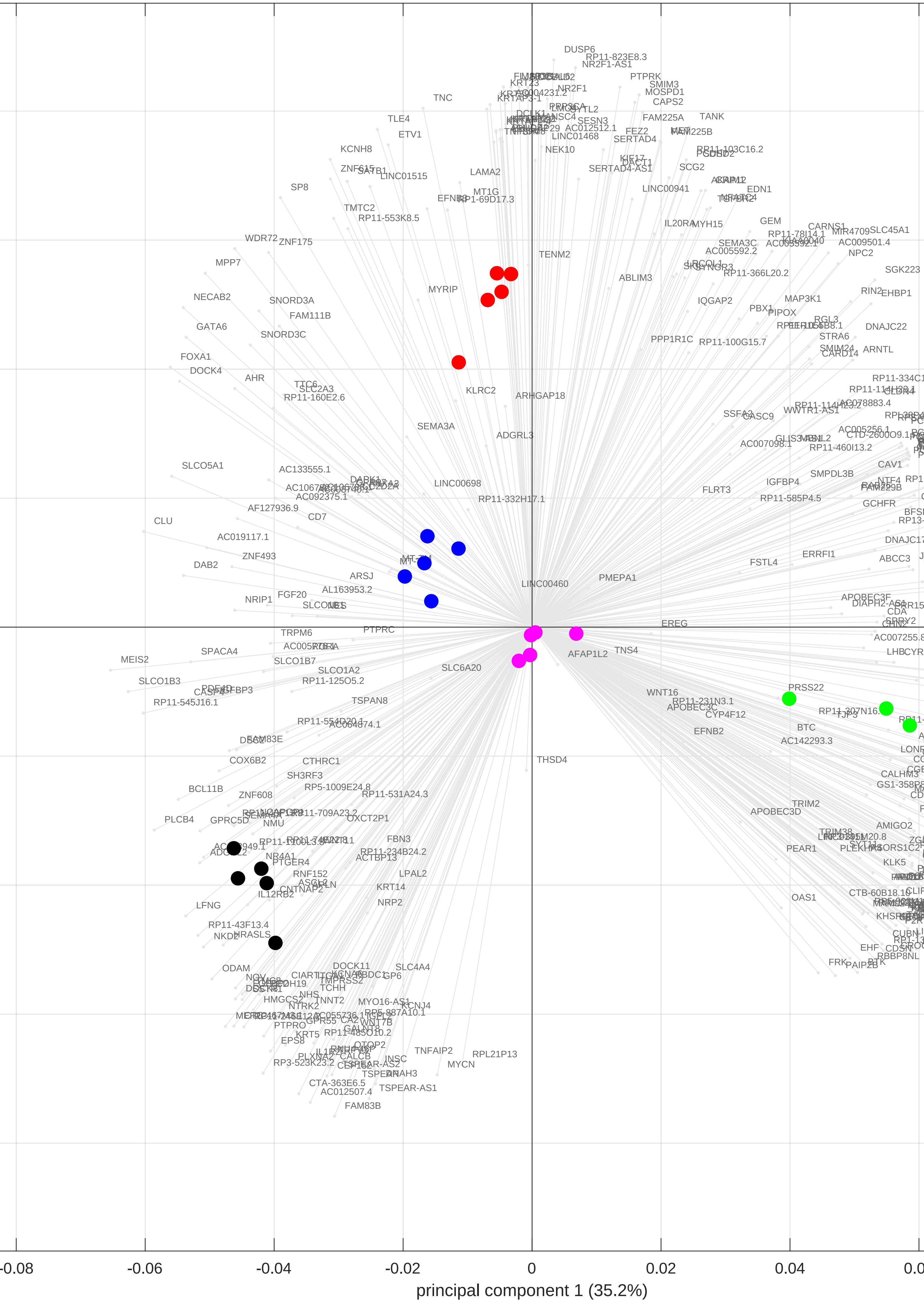

### pca_biplot.pdf

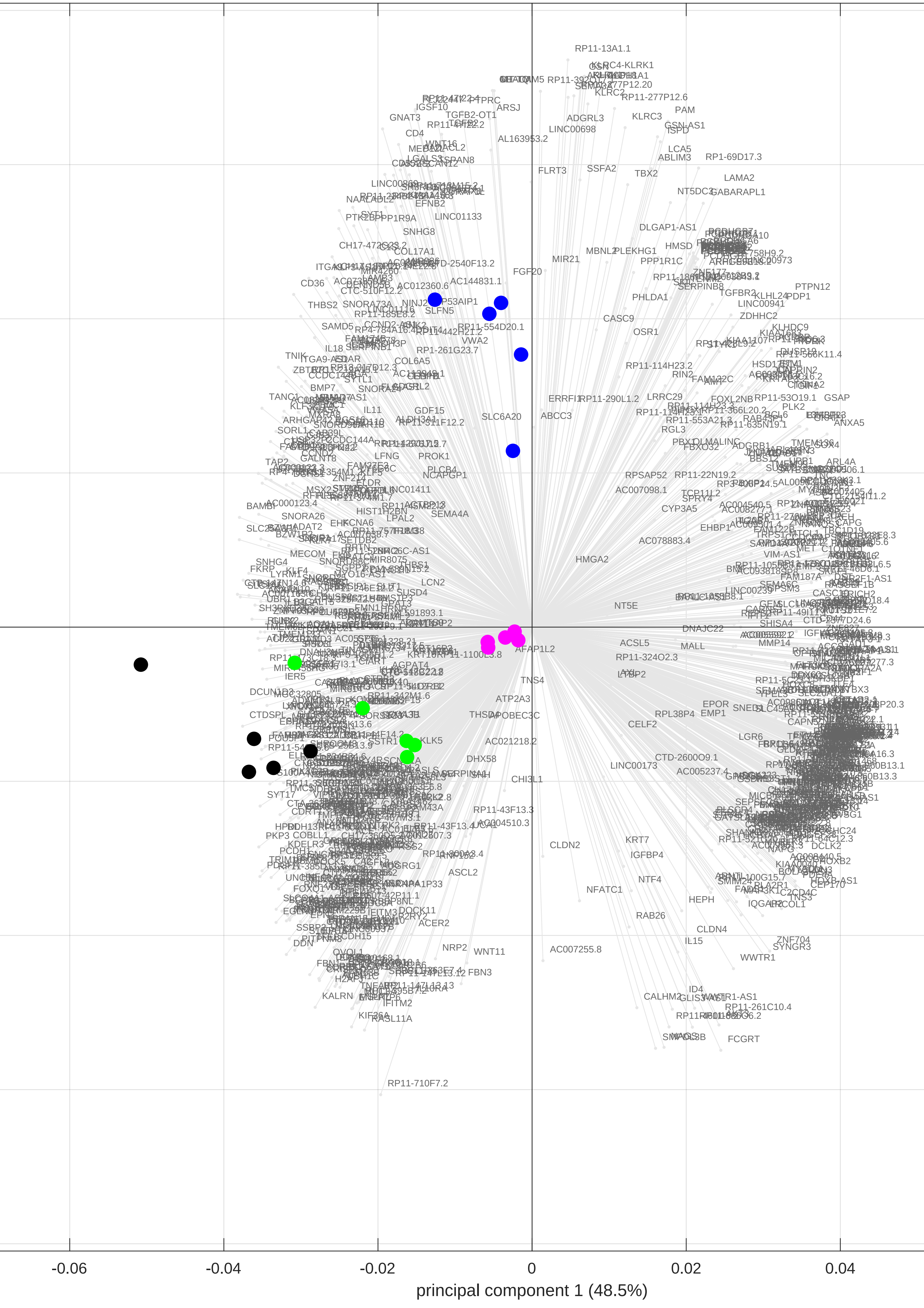

### pca_biplot.pdf

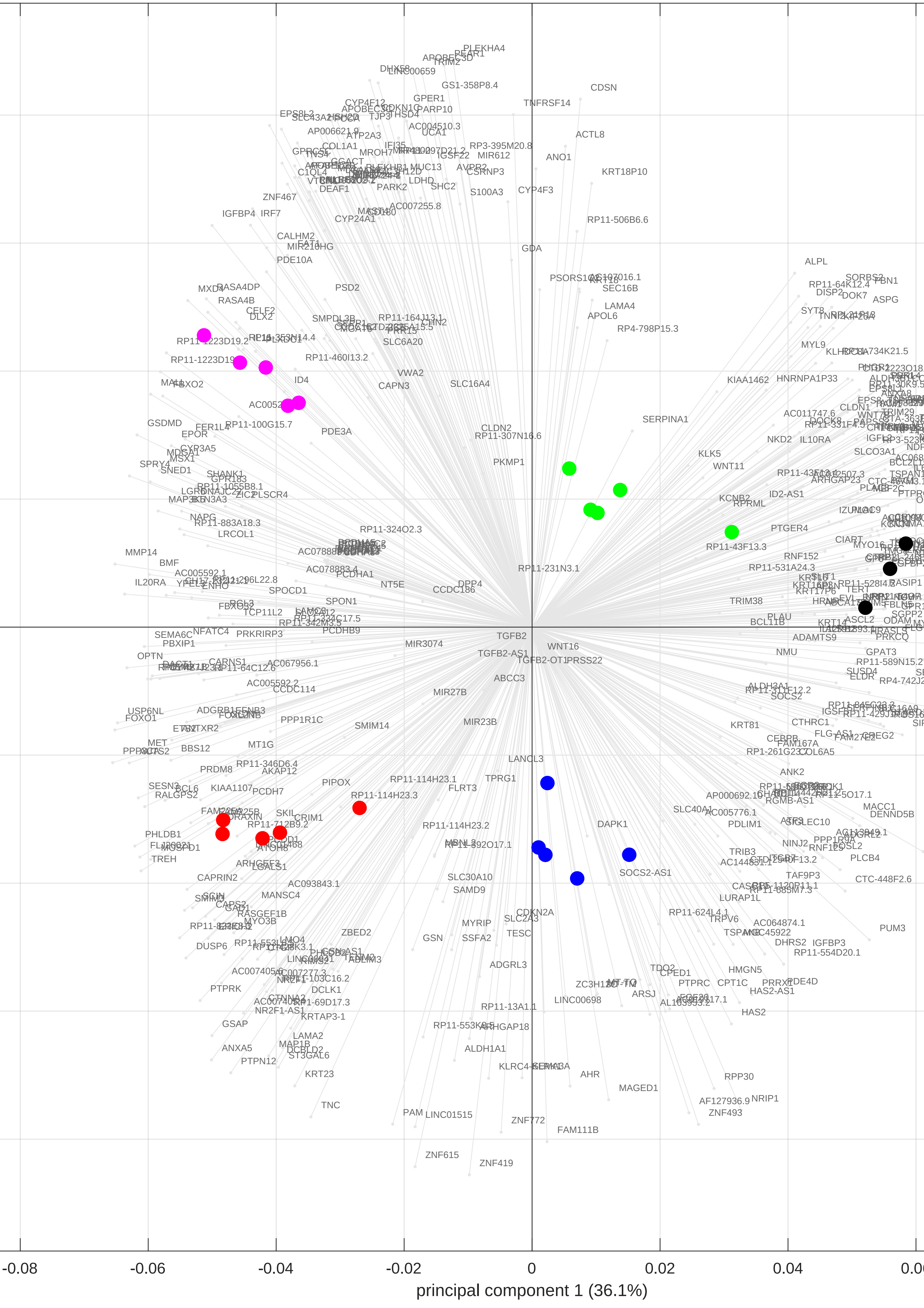

### pca_biplot.pdf

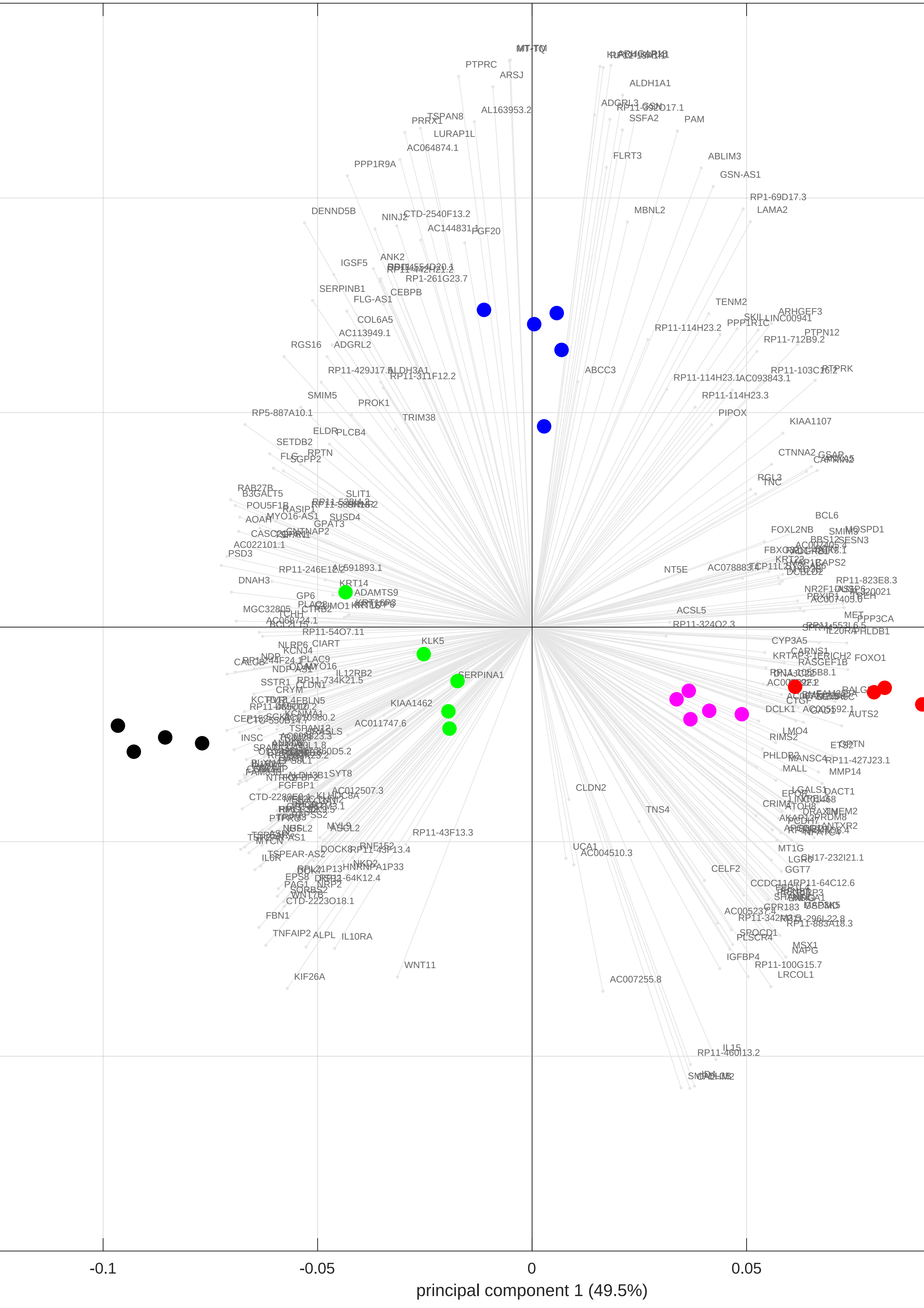

### pca_biplot.pdf

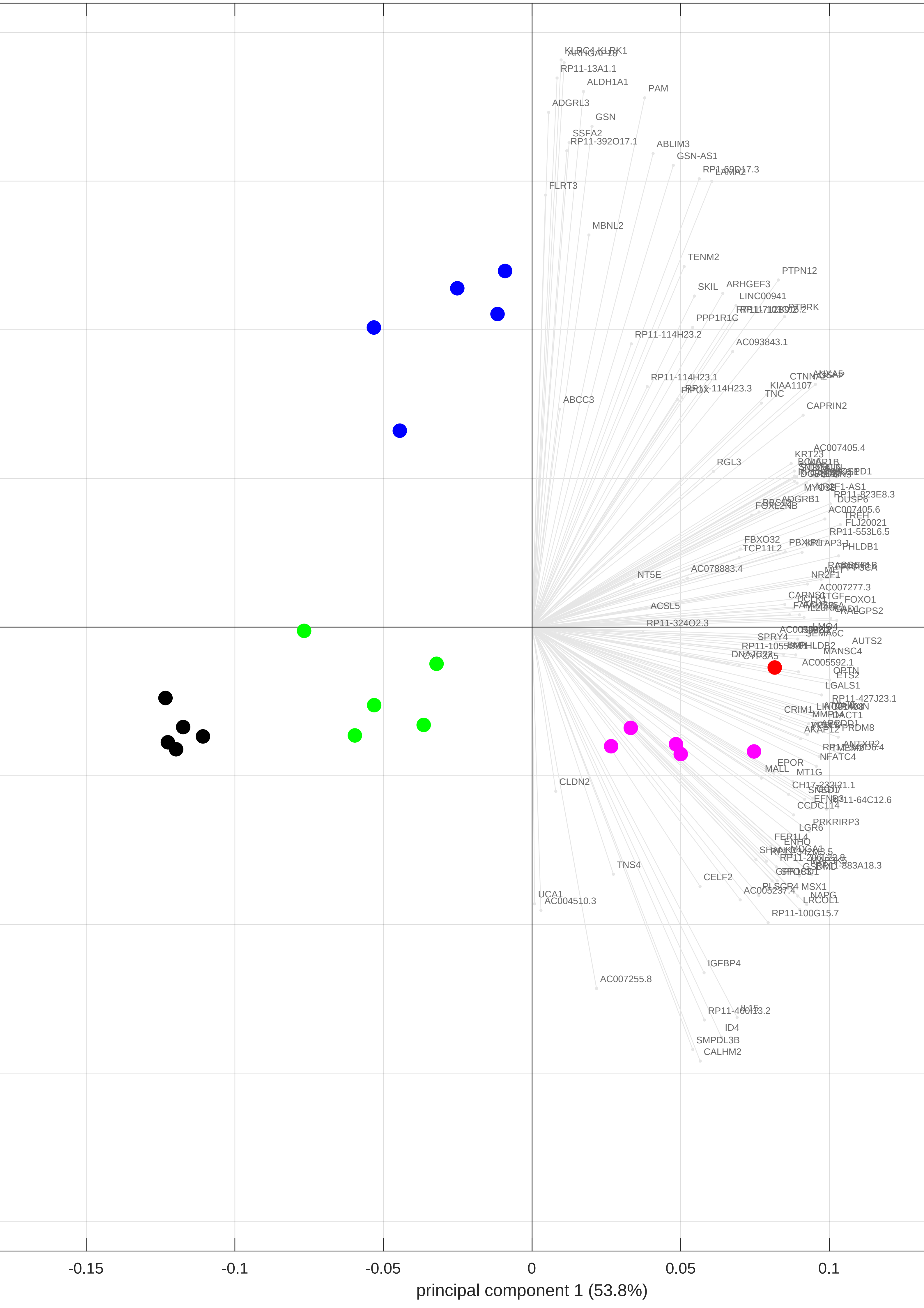

### pca_biplot.png

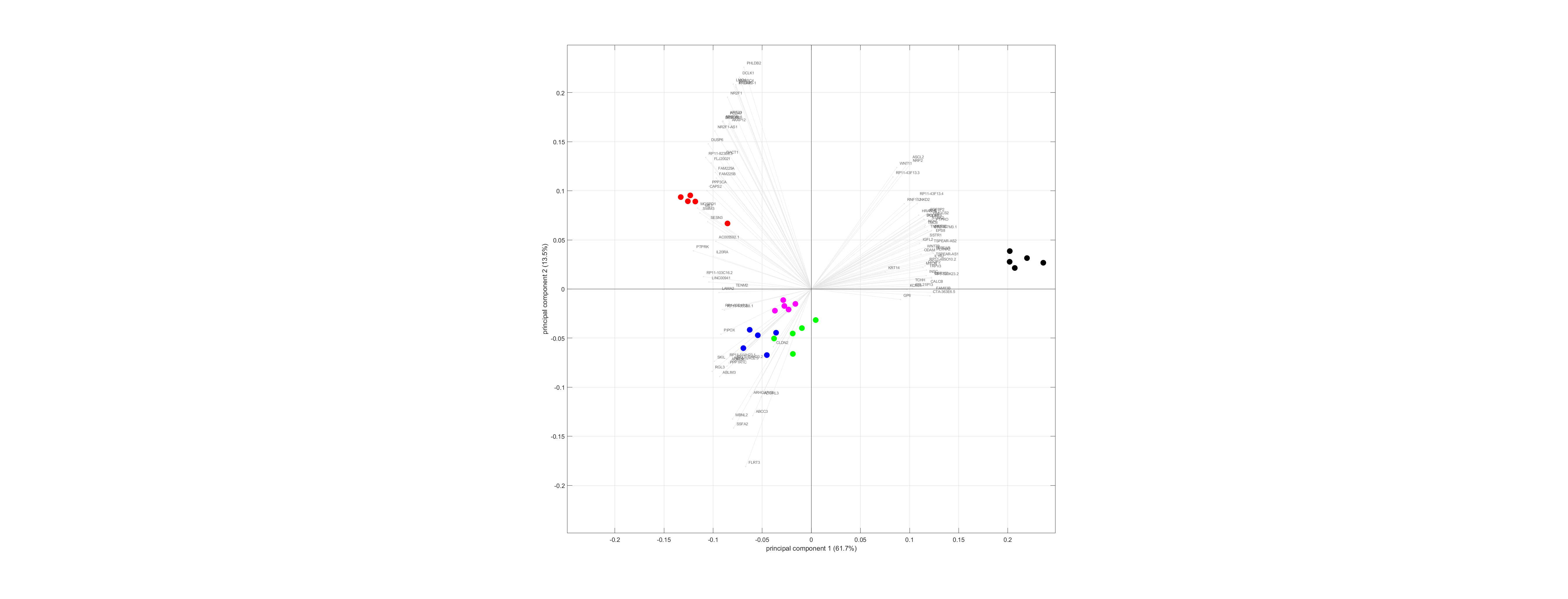

### pca_biplot.png

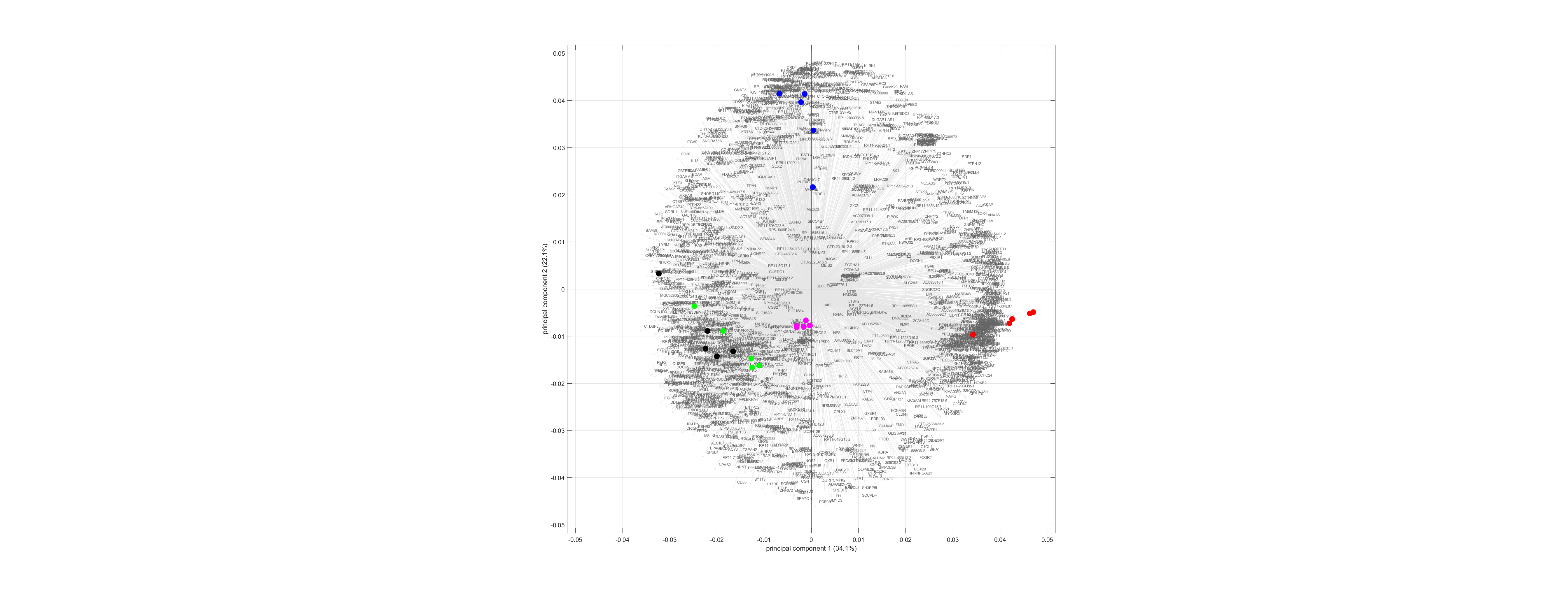

### pca_biplot.png

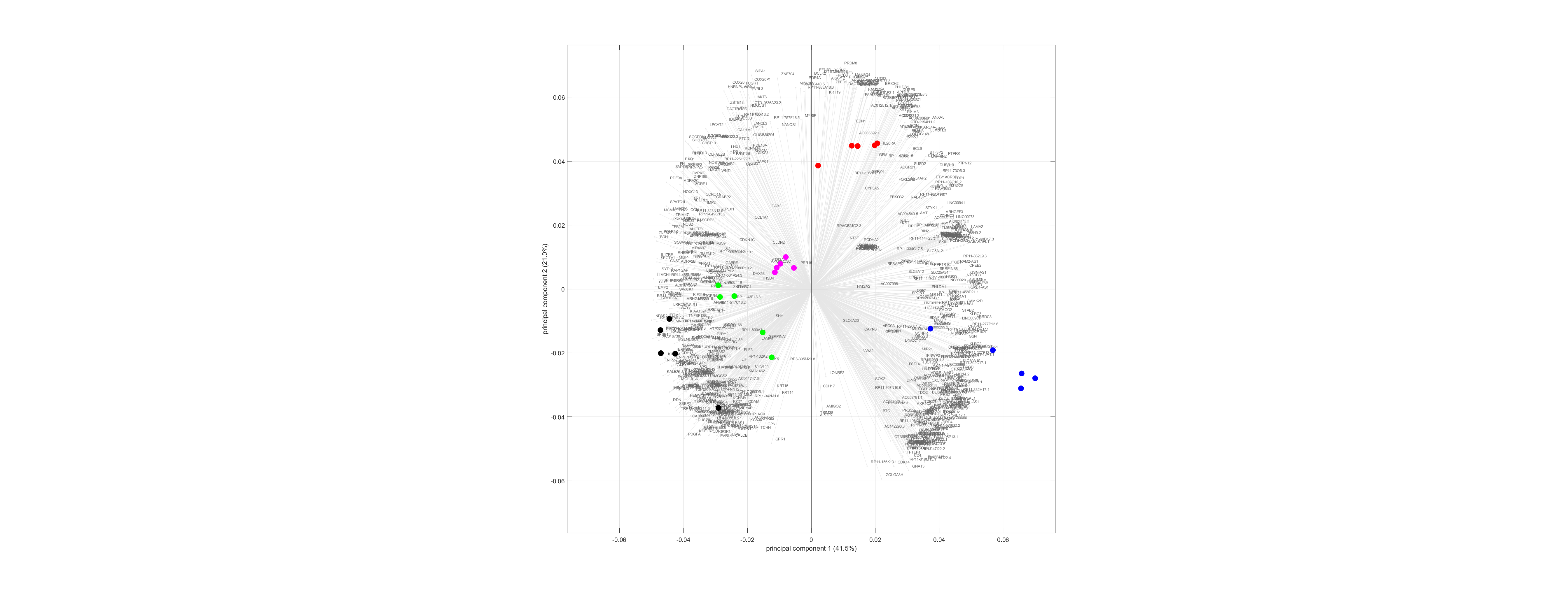

### pca_biplot.png

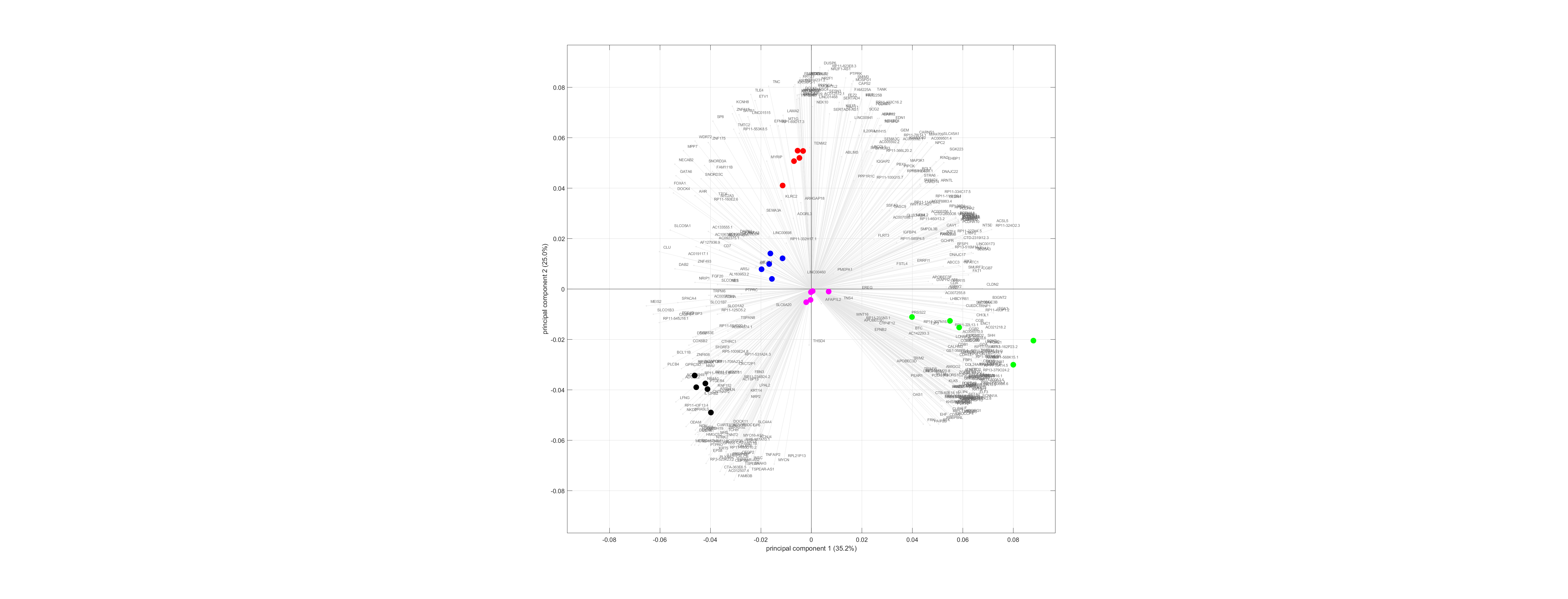

### pca_biplot.png

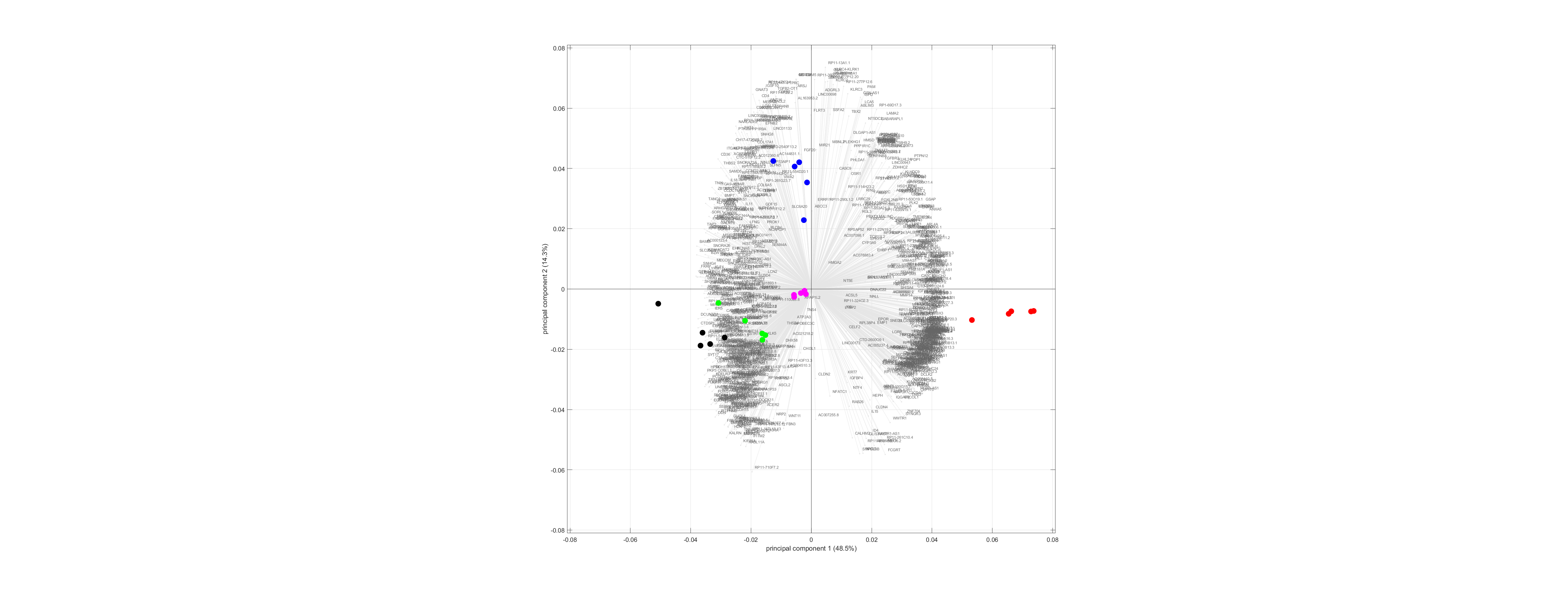

### pca_biplot.png

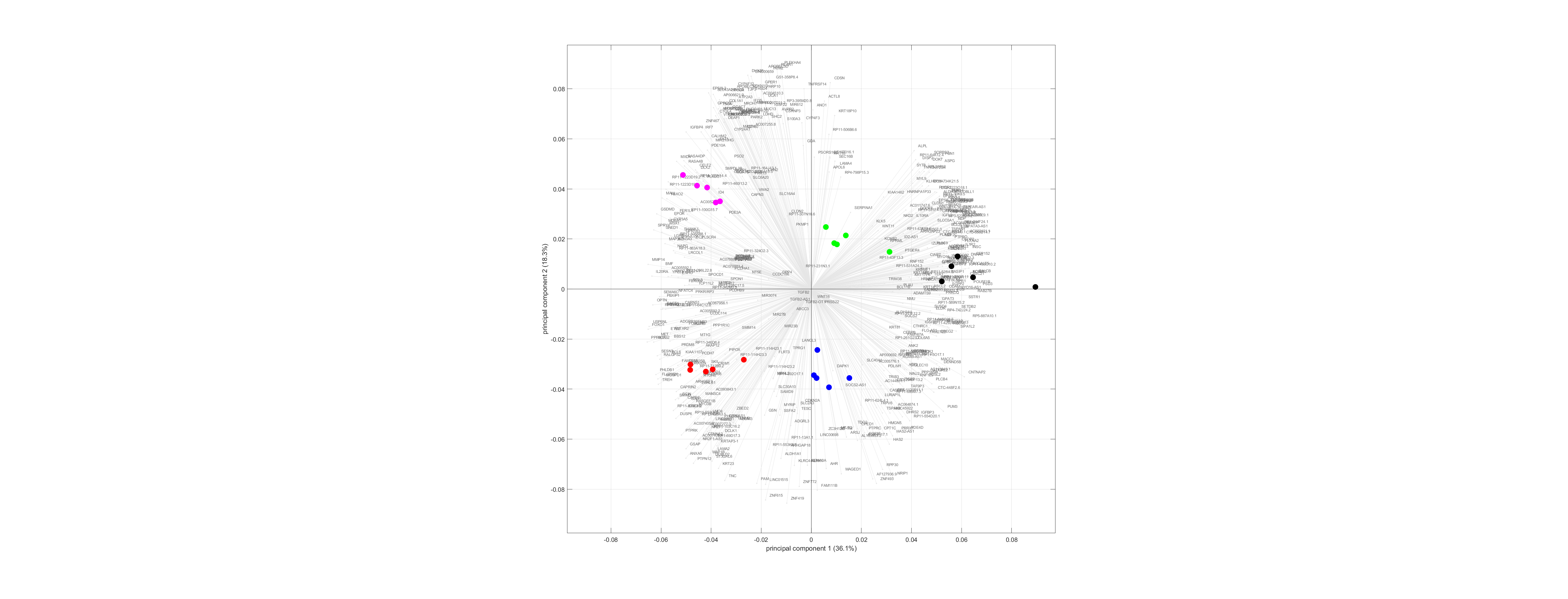

### pca_biplot.png

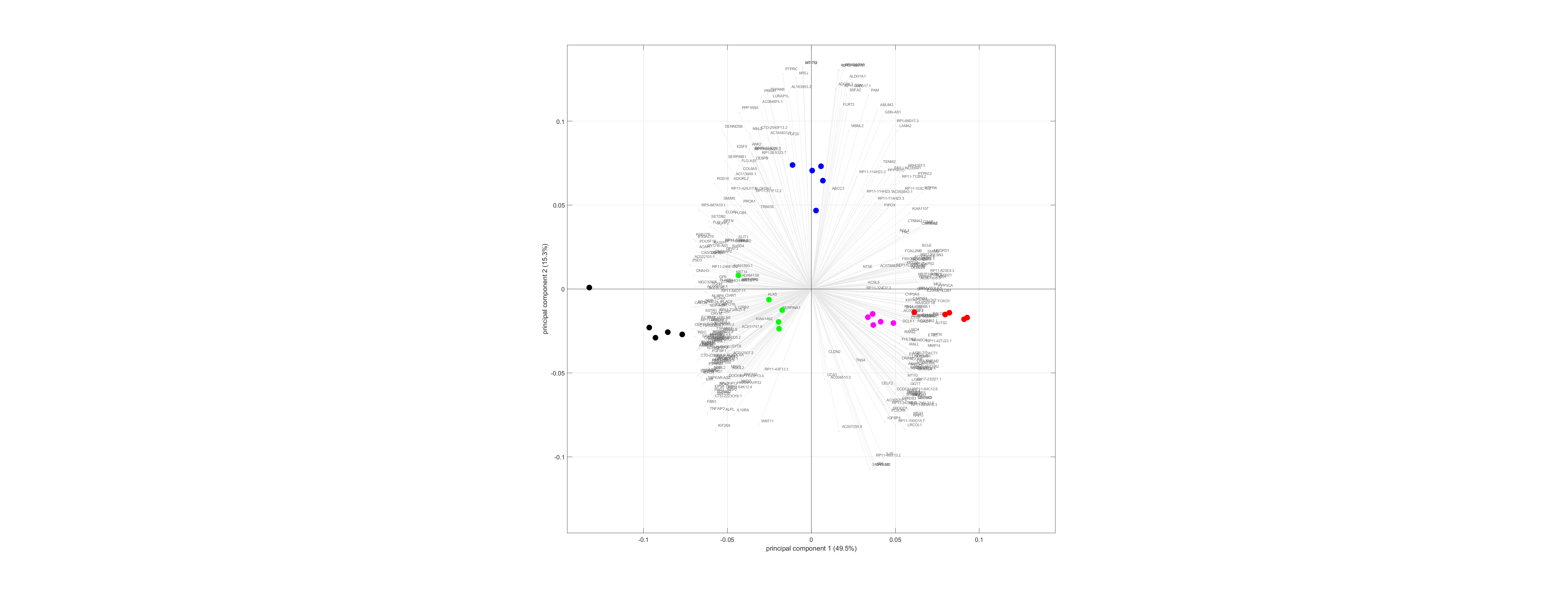

### pca_biplot.png

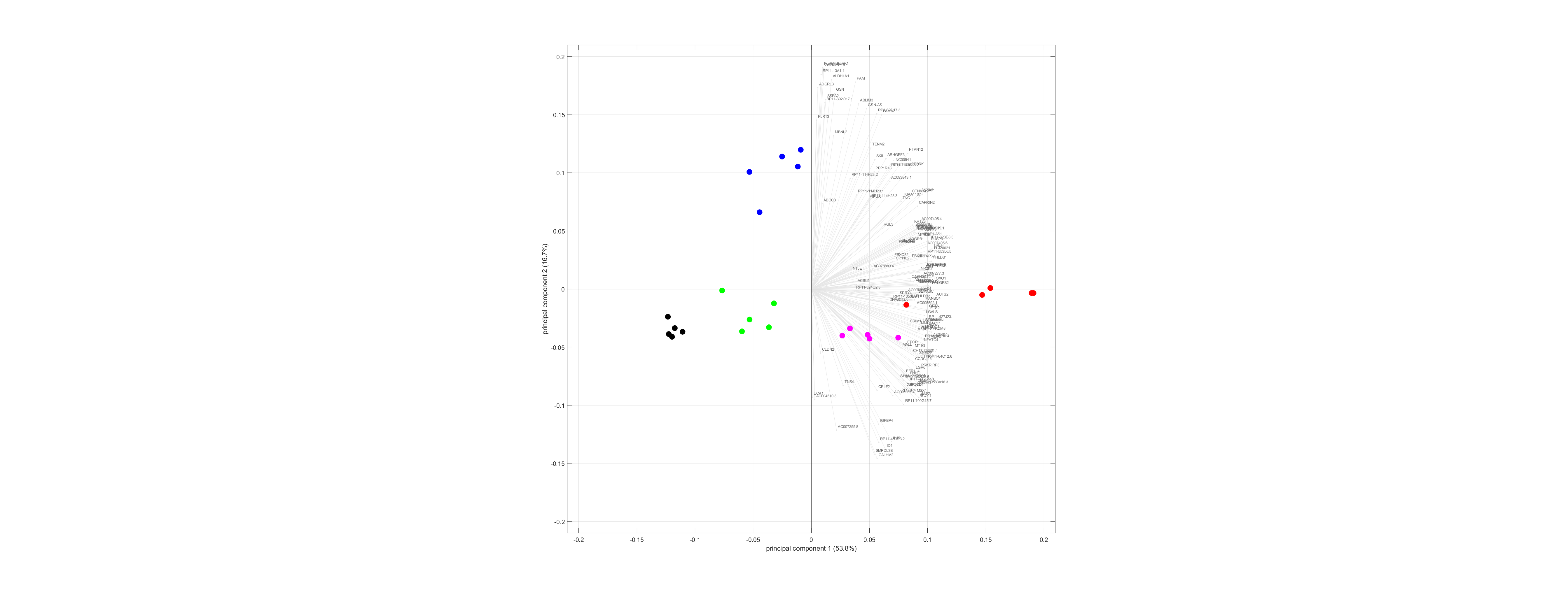

### vulcano.pdf

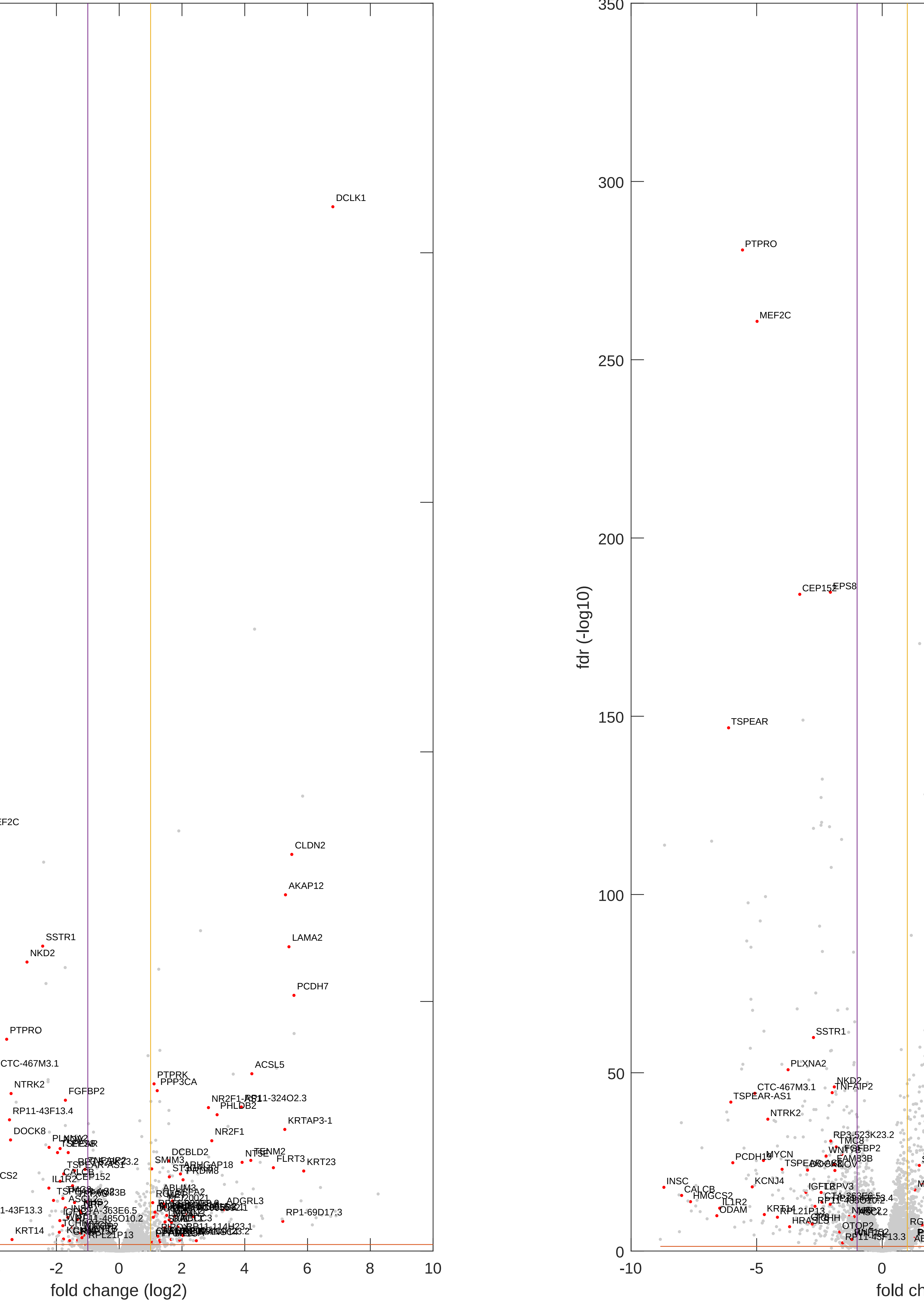

### vulcano.pdf

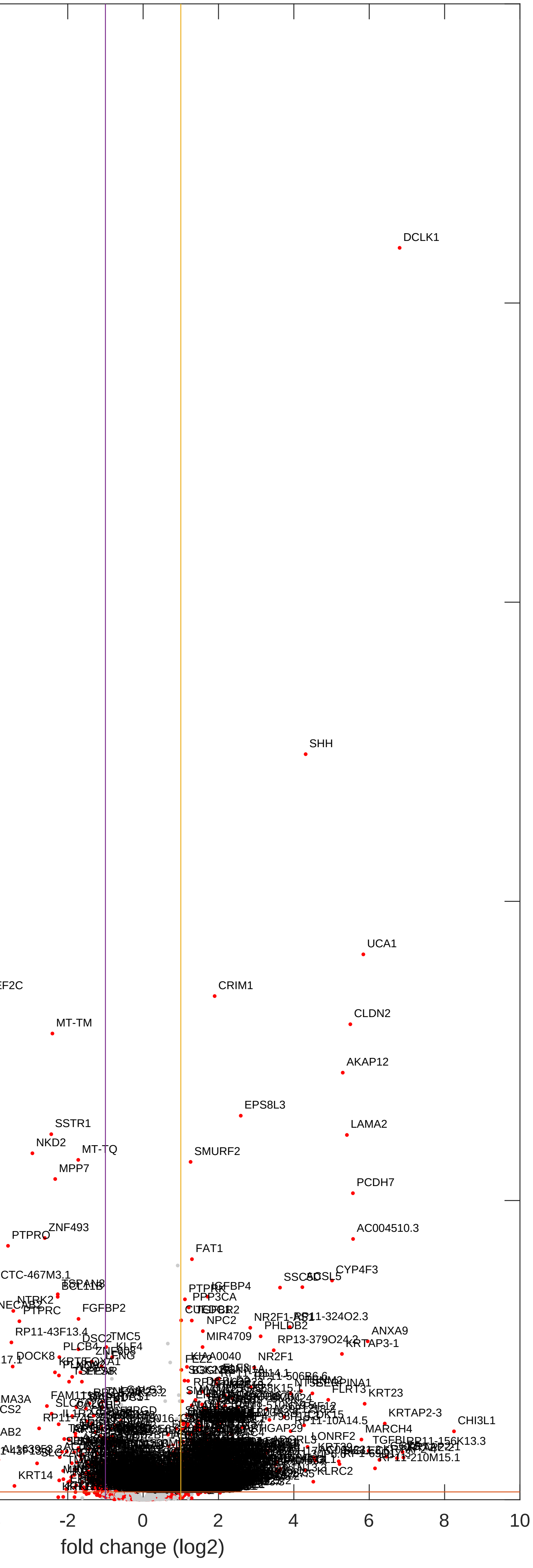

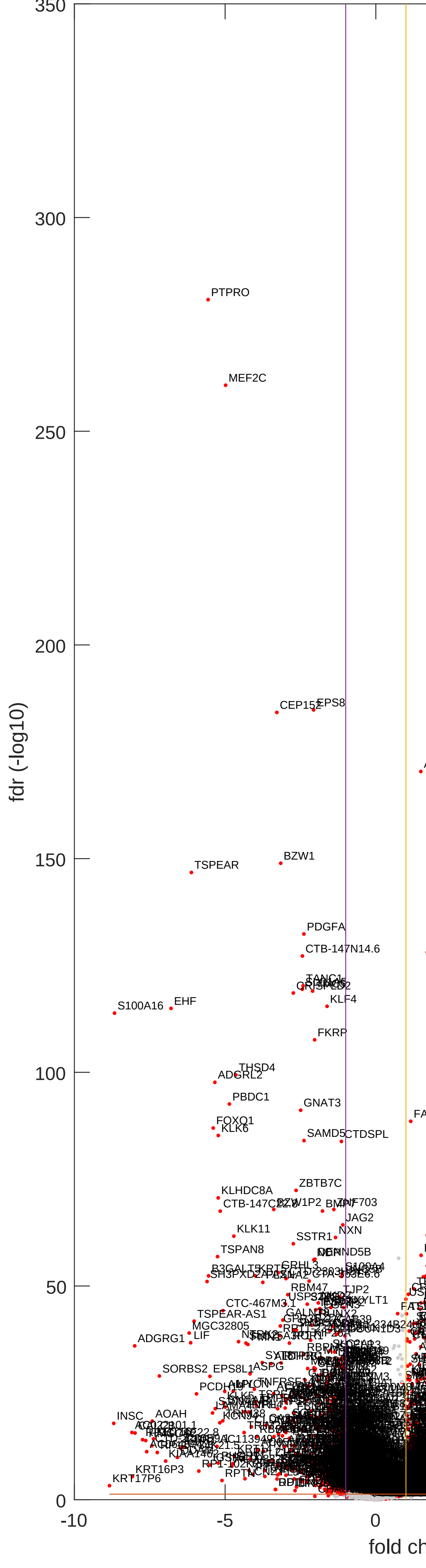

### vulcano.pdf

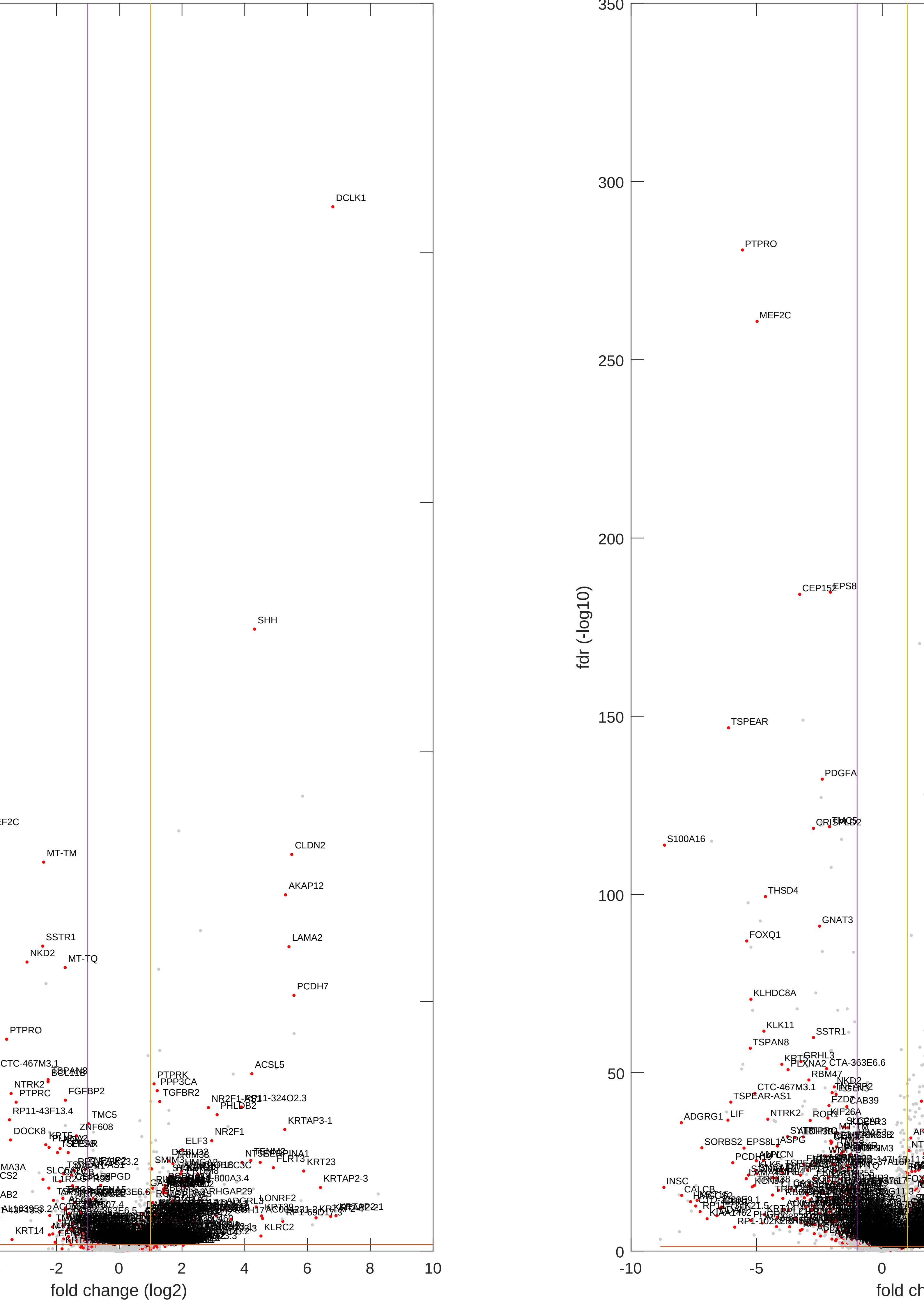

### vulcano.pdf

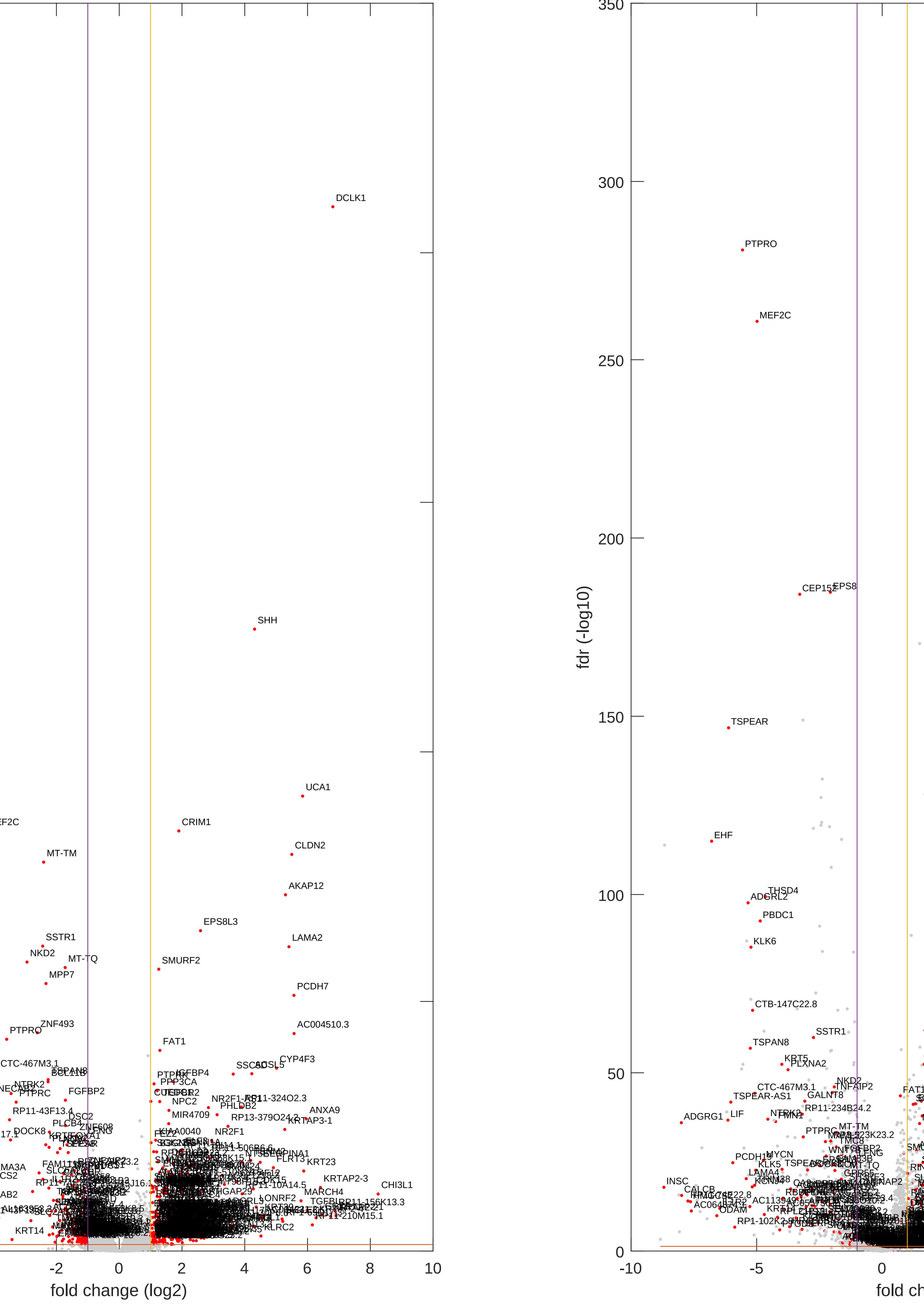

### vulcano.pdf

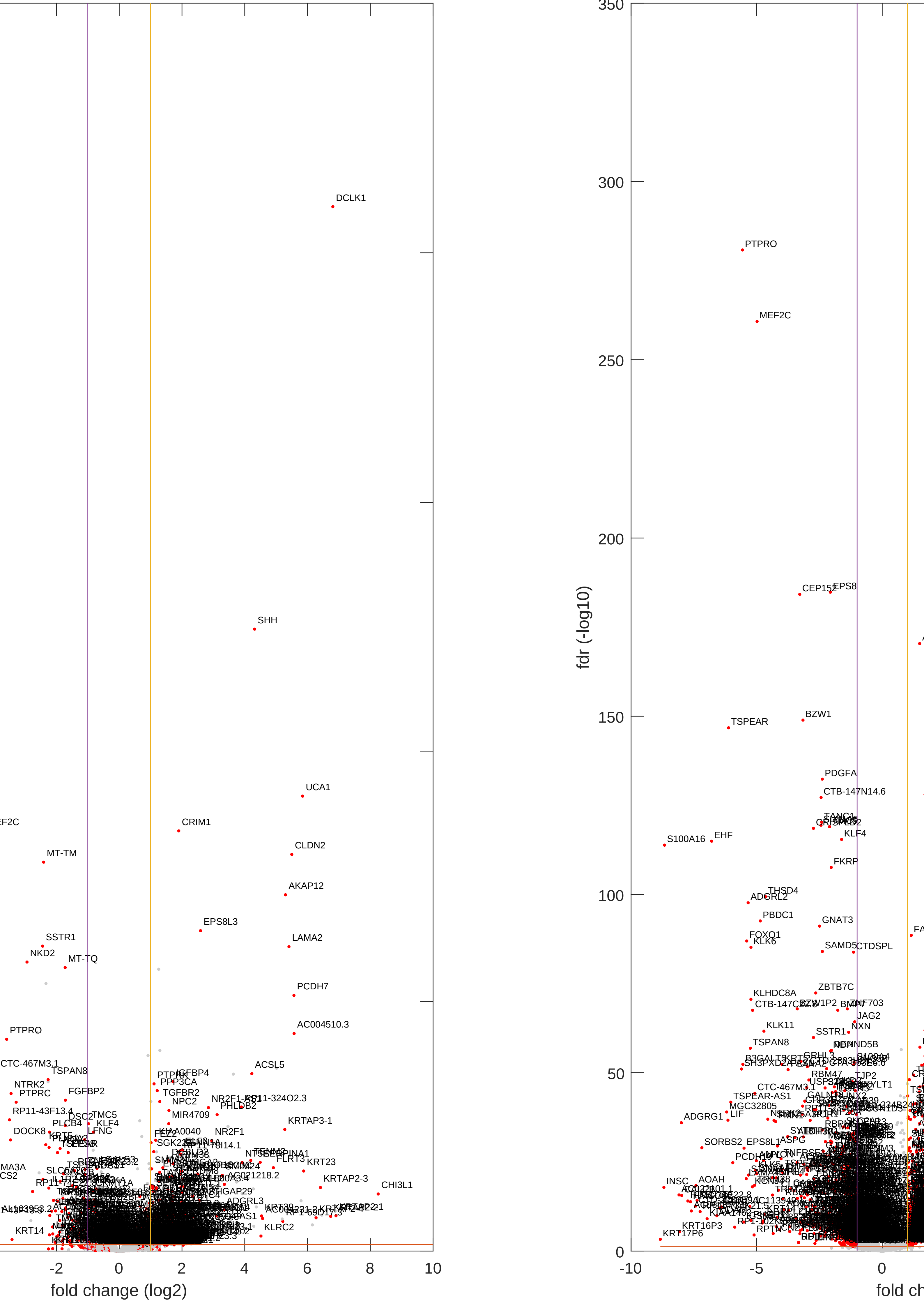

### vulcano.pdf

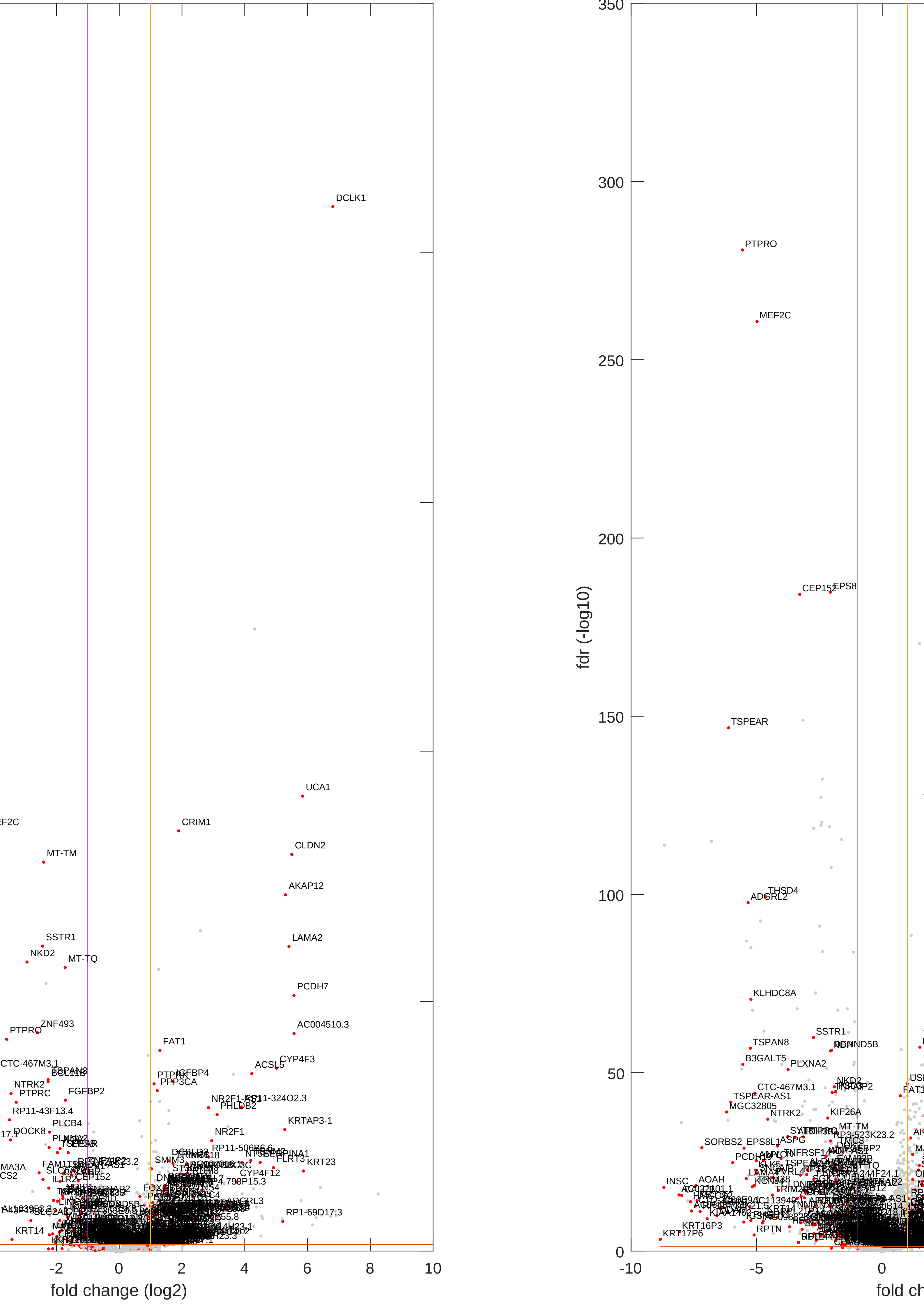

### vulcano.pdf

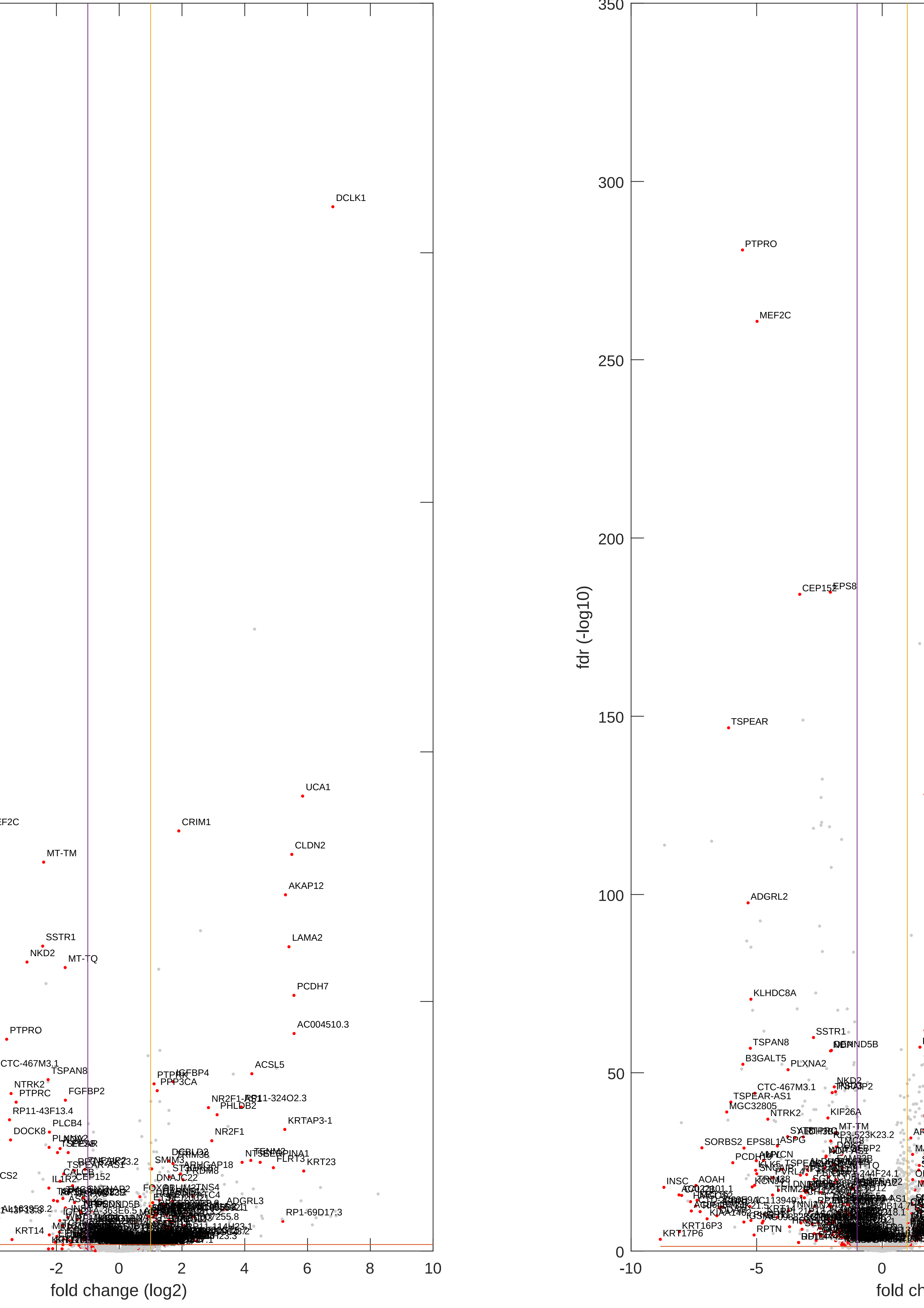

### vulcano.pdf

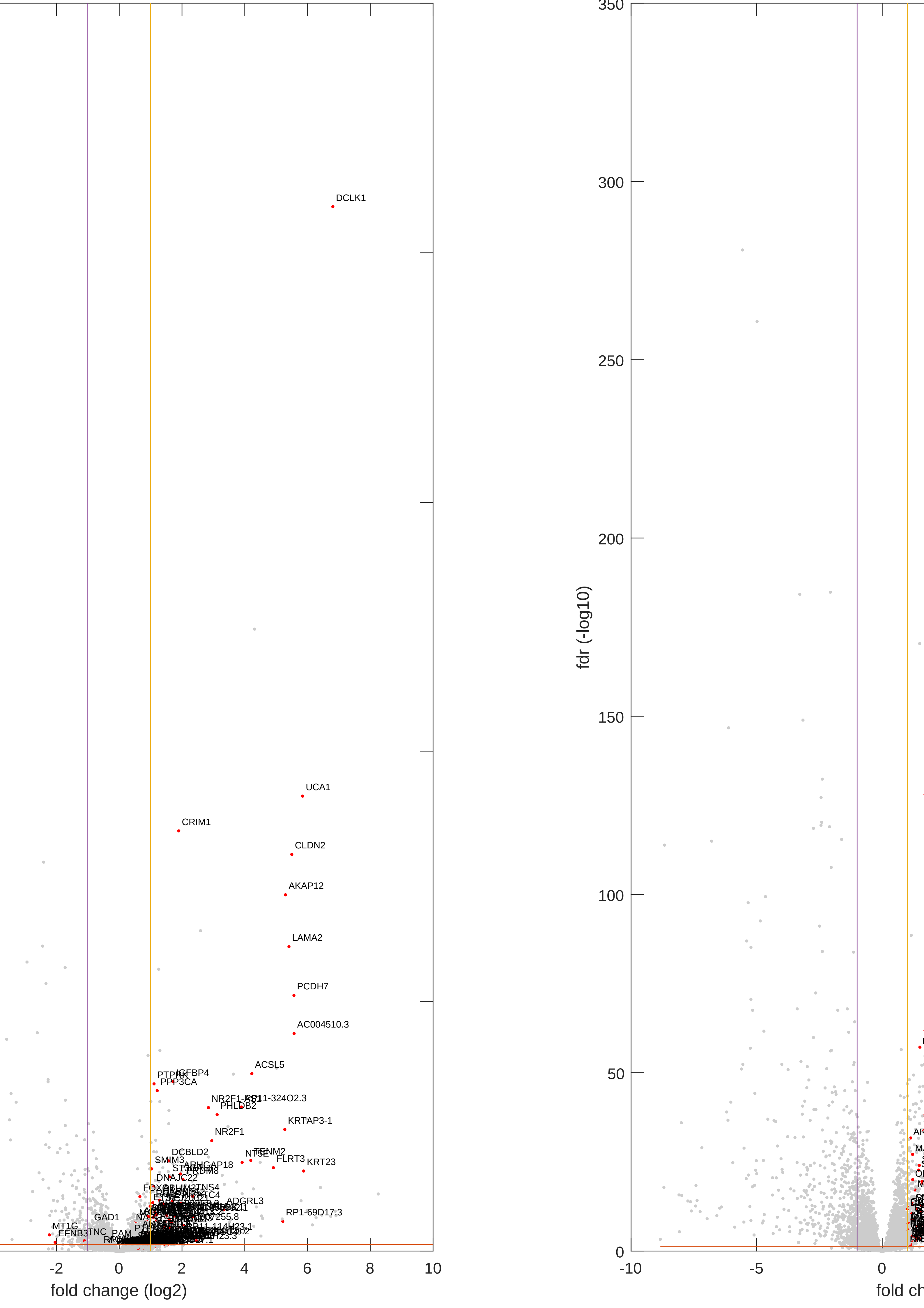

### vulcano.png

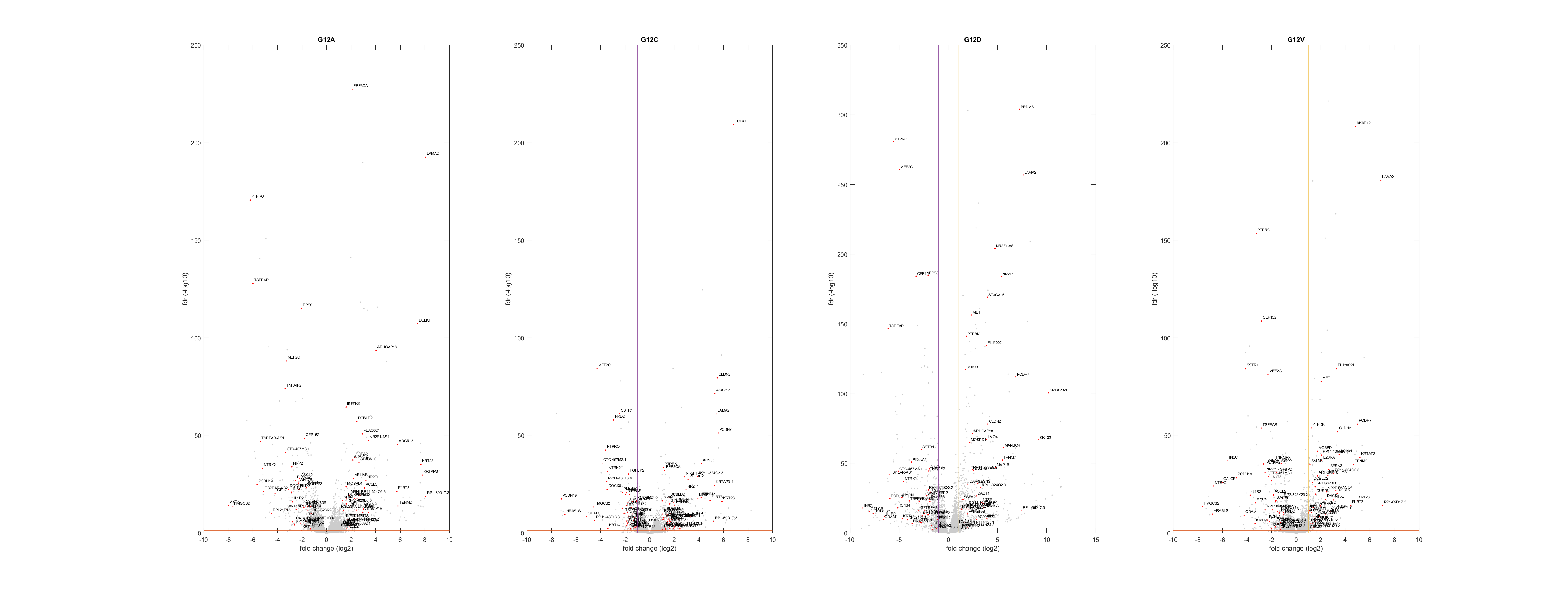

### vulcano.png

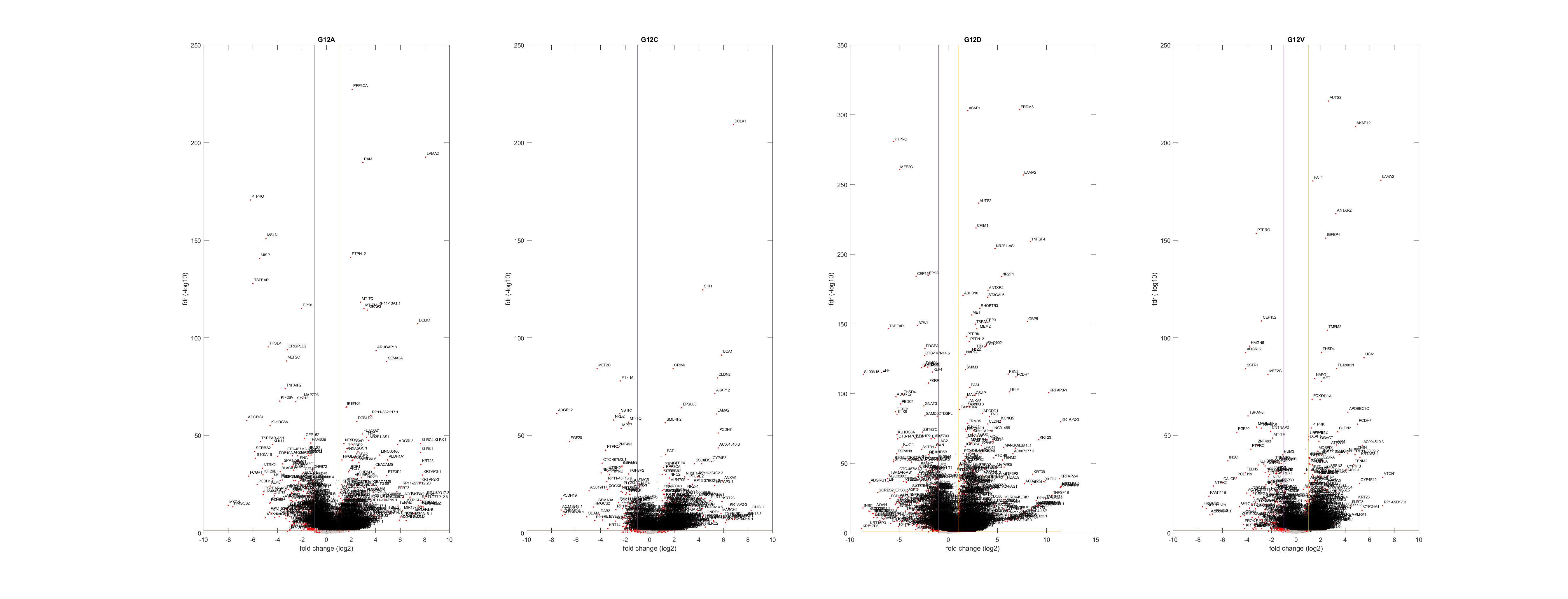

### vulcano.png

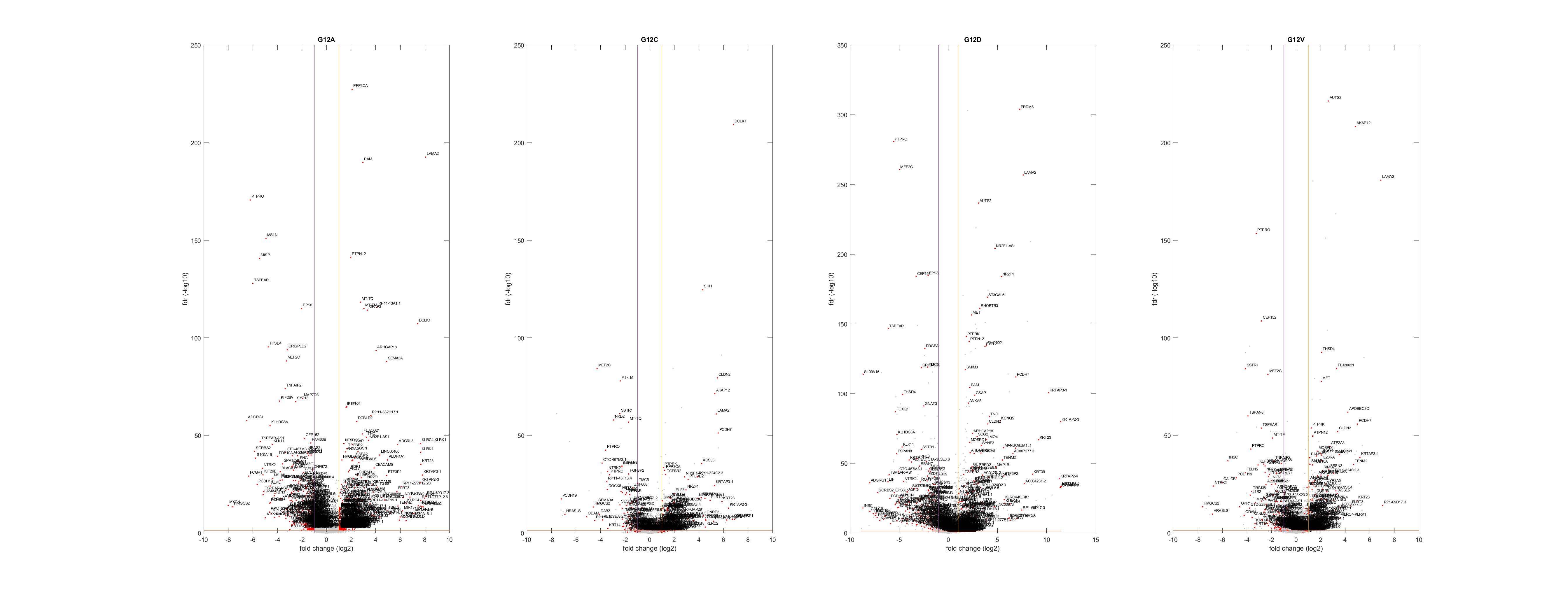

### vulcano.png

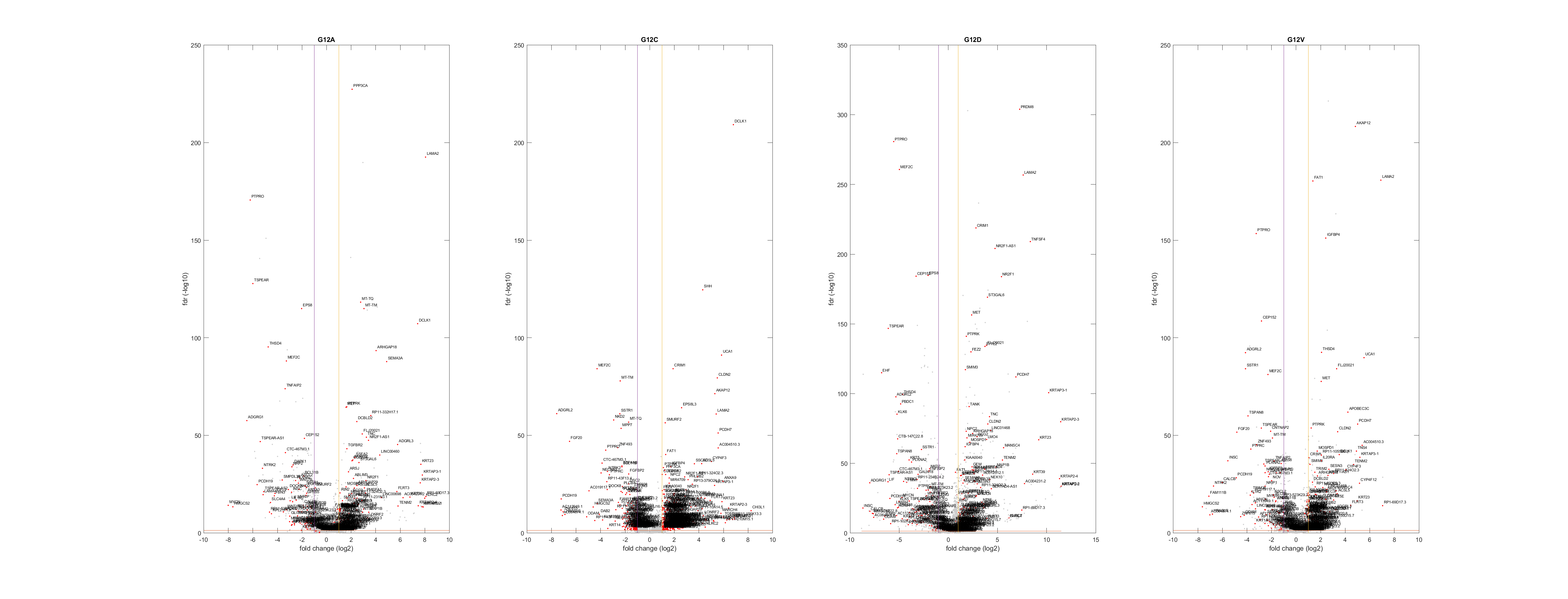

### vulcano.png

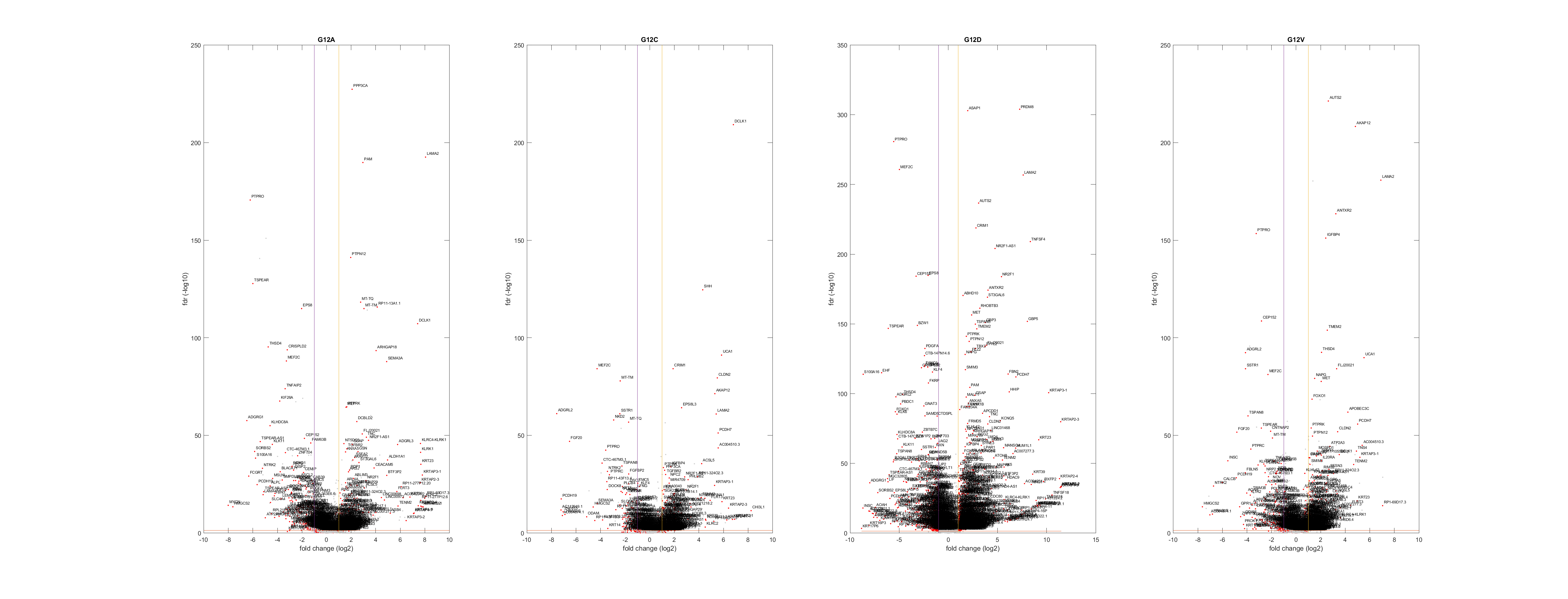

### vulcano.png

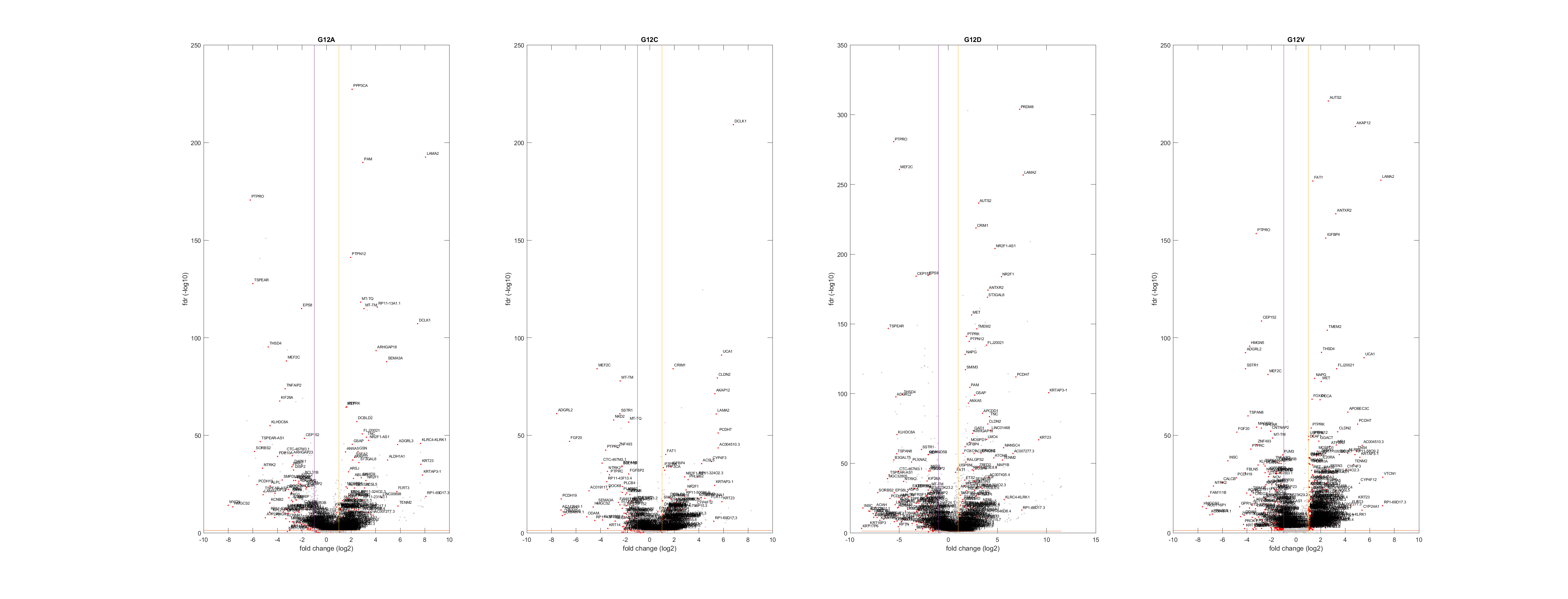

### vulcano.png

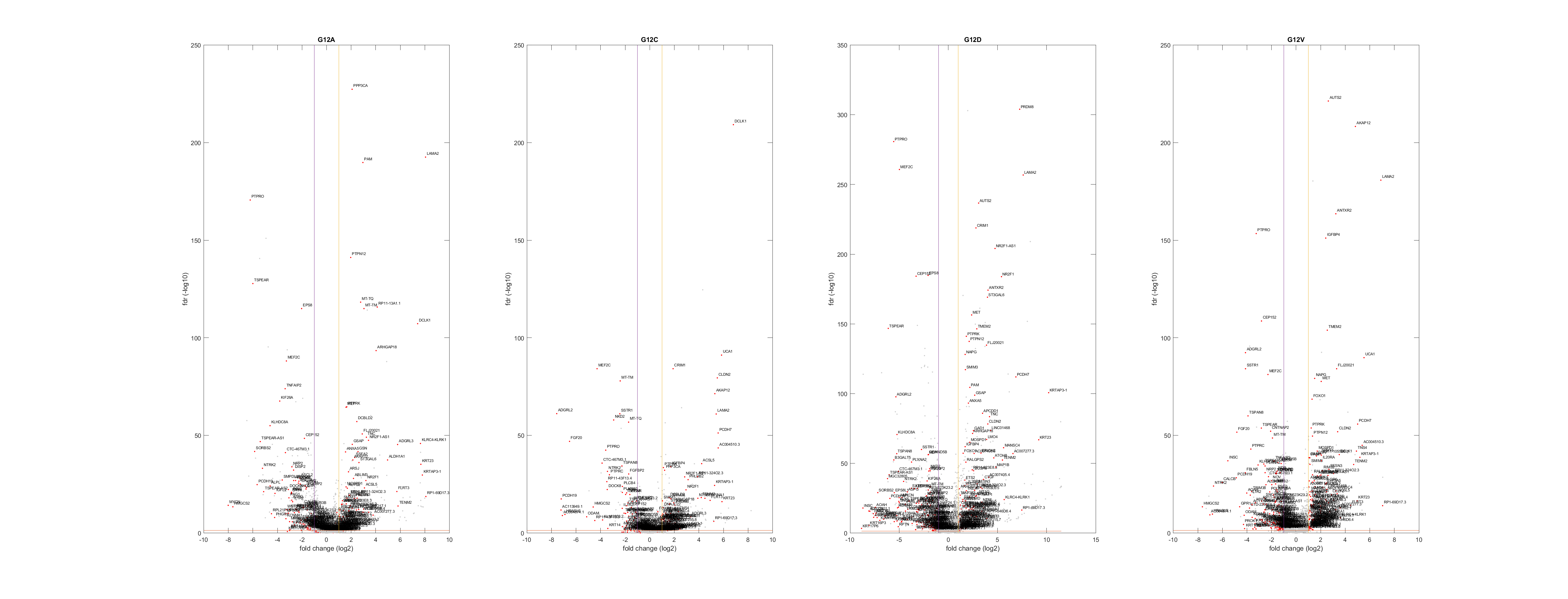

### vulcano.png

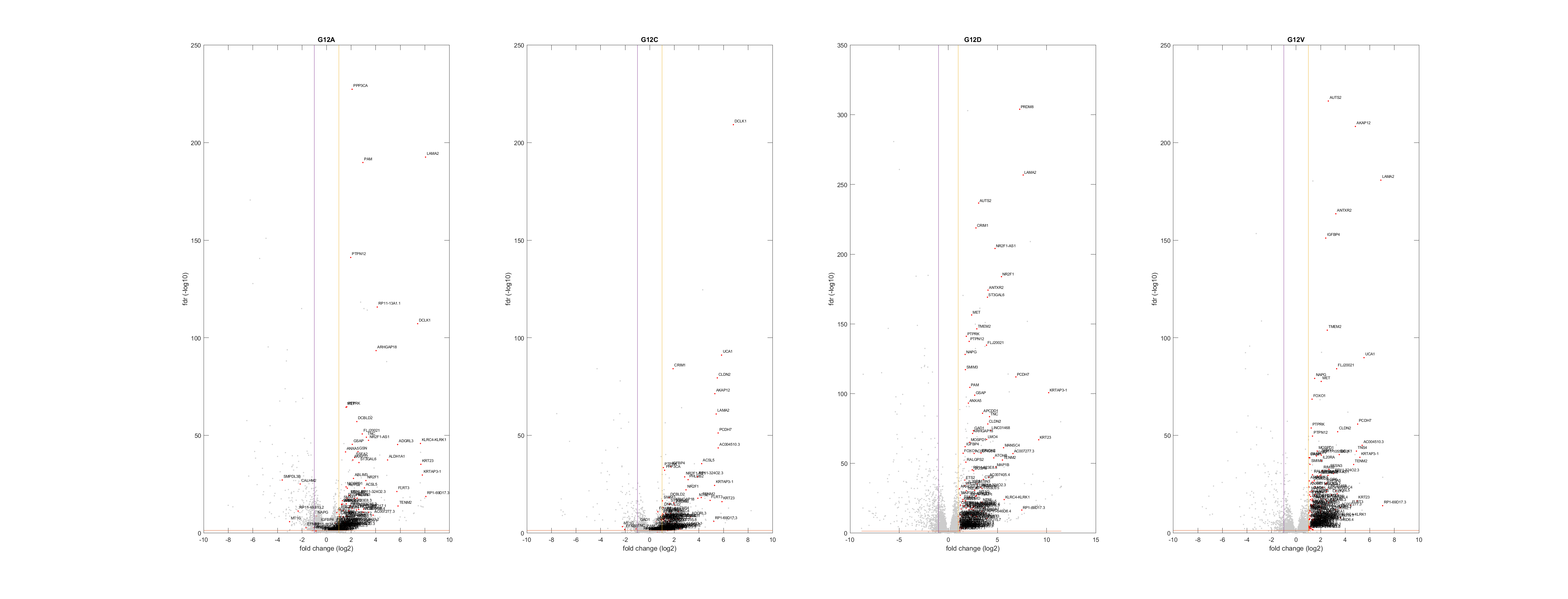
